# Methanotrophs are vigorous H_2_S oxidizers using a sulfide:quinone oxidoreductase and a *ba*_3_-type terminal oxidase

**DOI:** 10.1101/2022.08.31.505896

**Authors:** Rob A. Schmitz, Stijn H. Peeters, Sepehr S. Mohammadi, Tom Berben, Timo van Erven, Carmen A. Iosif, Theo van Alen, Wouter Versantvoort, Mike S.M. Jetten, Huub J.M. Op den Camp, Arjan Pol

## Abstract

Hydrogen sulfide (H_2_S) is produced in a wide range of anoxic environments where sulfate (SO_4_^2−^) reduction is coupled to decomposition of organic matter. In the same environments, methane (CH_4_) is the end product of an anaerobic food chain and both H_2_S and CH_4_ diffuse upwards into oxic zones where aerobic microorganisms can utilize these gases. Methane-oxidizing bacteria are known to oxidize a major part of the produced CH_4_ in these ecosystems, mitigating the emissions of this potent greenhouse gas to the atmosphere. However, how methanotrophy is affected by toxic H_2_S is largely unexplored. Here, we show that a single microorganism can oxidize CH_4_ and H_2_S simultaneously. By oxidizing H_2_S, the thermoacidophilic methanotroph *Methylacidiphilum fumariolicum* SolV can alleviate the inhibitory effects on CH_4_ oxidation. In response to H_2_S, strain SolV upregulated a type III sulfide:quinone oxidoreductase (SQR) and a sulfide-insensitive *ba*_3_-type terminal oxidase to dissipate the reducing equivalents derived from H_2_S oxidation. Through extensive chemostat cultivation of *M. fumariolicum* SolV we demonstrate that it converts high loads of H_2_S to elemental sulfur (S^0^). Moreover, we show chemolithoautotrophy by tracing ^13^CO_2_ fixation into new biomass by using H_2_S as sole energy source. Molecular surveys revealed several putative SQR sequences in a range of proteobacterial methanotrophs from various environments, suggesting that H_2_S detoxification is much more widespread in methanotrophs than previously assumed, enabling them to connect carbon and sulfur cycles in new ways.

## INTRODUCTION

Hydrogen sulfide (H_2_S) is the most reduced form of sulfur (S) and a potent energy source, toxicant and signaling molecule (Bagarinao, 1992; Kimura et al., 2012; Berben et al., 2019). It is a weak acid that easily diffuses through membranes and is known to inhibit various processes, including aerobic respiration by binding to cytochrome *c* oxidases as well as several other metabolic process that use copper- and iron-containing enzymes (Bagarinao, 1992; Pietri et al., 2011; Barton et al., 2014; Landry et al., 2021). Consequently, microorganisms living in environments in which H_2_S is present require effective mechanisms to detoxify H_2_S (Marcia et al., 2009; Xia et al., 2017). H_2_S is produced during sulfate (SO_4_^2−^) reduction coupled to the decomposition and mineralization of organic matter, in a myriad of environments, such as wetlands, marine sediments, landfills, and acidic geothermal environments (Jørgensen, 1982; Lomans et al., 2002; Lyimo et al., 2002; Muyzer and Stams, 2008; Pester et al., 2012; Xia et al., 2017; Fakhraee and Katsev, 2019; Jørgensen et al., 2019; Schmitz et al., 2021). Accordingly, emissions of H_2_S can lead to the chemical production of sulfur dioxides in the atmosphere, causing subsequent acid rain and acidification of the environment (Bates et al., 1992). Furthermore, H_2_S produced by microbes contributes to corrosion of pipelines, causing damage costing billions of dollars per year (Enning and Garrelfs, 2014).

After sulfate is depleted, the remaining organic matter is ultimately converted to methane (CH_4_) in oxygen depleted ecosystems (Jørgensen, 1982; Dar et al., 2008; Muyzer and Stams, 2008; Pester et al., 2012; Sela-Adler et al., 2017; Chen et al., 2019). When both H_2_S and CH_4_ diffuse into the overlaying oxic zones, CH_4_ can be utilized as energy source by aerobic methane-oxidizing bacteria, that are assumed to mitigate the majority of emissions of this potent greenhouse gas (Murrell and Jetten, 2009). Despite this effective methane biofilter, 548 to 736 Tg of CH_4_ is still annually released into the atmosphere from various natural and anthropogenic sources (Kirschke et al., 2013; Dean et al., 2018). Aerobic methanotrophs are part of several bacterial classes and families, including the ubiquitous Alpha- and Gammaproteobacteria (Hanson and Hanson, 1996; Op den Camp et al., 2009; Schmitz et al., 2021) and the extremely acidophilic *Methylacidiphilaceae* of the phylum Verrucomicrobia (Pol et al., 2007; Dunfield et al., 2007; Islam et al., 2008; Van Teeseling et al., 2014). The latter are acidophilic bacteria that all share a low pH optimum (2.0−3.5) and live between 35 and 60 °C (Op den Camp et al., 2009; Van Teeseling et al., 2014; Mohammadi et al., 2019). All known verrucomicrobial methanotrophs have been isolated from geothermal environments such as fumaroles and mudpots, from which large amounts of mostly thermogenic CH_4_ and H_2_S are emitted (Castaldi and Tedesco, 2005; Pol et al., 2007; Crognale et al., 2018; Picone et al., 2020a; Schmitz et al., 2021). Since geothermal environments are typically characterized by high H_2_S emissions, the verrucomicrobial methanotrophs isolated from these ecosystems are preeminent examples to study how methanotrophs would cope with H_2_S.

From recent studies it has become clear that methanotrophs are metabolically versatile and able to use environmentally relevant energy sources such as H_2_, propane, ethane, acetate, acetone, 2-propanol, and acetol (Hakobyan and Liesack, 2020; Picone et al., 2020b; Schmitz et al., 2021; Awala et al., 2021). The ability to utilize various energy sources is highly beneficial in environments with heavily fluctuating gas emissions. Recently, it was demonstrated that pure cultures of the verrucomicrobial methanotroph *Methylacidiphilum fumariolicum* SolV can consume methanethiol (CH3SH), with concomitant substoichiometric formation of H_2_S, which indicated that strain SolV partly metabolized this toxic intermediate (Schmitz et al., 2022). Subsequently, the elegant study of Gwak et al. (2022) demonstrated that also proteobacterial methanotrophs can oxidize H_2_S. They isolated the versatile alphaproteobacterium ‘*Methylovirgula thiovorans*’ strain HY1 from a peatland in South Korea that could grow on thiosulfate (S_2_O_3_^2−^), tetrathionate (S_4_O_6_^2−^), elemental sulfur (S^0^), and a range of carbon compounds. However, growth on H_2_S was not studied. In addition, strain HY1 cells grown on methane as sole energy source were not able to oxidize H_2_S, and H_2_S oxidation was only initiated and observed in cells grown in the presence of thiosulfate. Based on these observations it is paramount to investigate whether microorganisms exist that can oxidize the environmentally relevant gases CH_4_ and H_2_S simultaneously, how methanotrophs cope with H_2_S and whether such methanotrophs can conserve energy and produce biomass using H_2_S as energy source.

Here, through extensive chemostat cultivation we show for the first time that a microorganism can oxidize CH_4_ and H_2_S simultaneously. *M. fumariolicum* SolV is inhibited by the presence of elevated H_2_S concentrations and rapidly oxidizes H_2_S to elemental sulfur (S^0^) as a detoxification mechanism to alleviate the inhibitory effect of H_2_S on methane oxidation. Strain SolV responds to H_2_S with the upregulation of a Type III sulfide:quinone oxidoreductase (SQR) and a H_2_S-insensitive *ba*_3_-type cytochrome *c* oxidase, to create an electron transfer pathway from H_2_S to O_2_. Furthermore, strain SolV incorporates ^13^CO_2_ using energy derived from the oxidation of H_2_S as sole energy source. We propose that the H_2_S oxidation capacity of verrucomicrobial methanotrophs is essential to thrive in acidic geothermal ecosystems. In addition, we found SQR in a plethora of proteobacterial methanotrophs of various environments. Since CH_4_ and H_2_S co-occur in a myriad of oxygen-limited ecosystems, H_2_S oxidation could be a trait present among many aerobic methanotrophs.

## METHODS

### Phylogenetic analysis

All available genome sequences of known methylotrophs from the orders Methylococcales (Gammaproteobacteria) and Methylacidiphilales (Verrucomicrobia), the families *Methylocystaceae* and *Beijerinckia* (Alphaproteobacteria), and the genus *Methylomirabilis* were retrieved from the NCBI database. Genomes of methanotrophs were selected by blasting amino acid sequences of PmoA from *Methylococcus capsulatus* (SwissProt accession Q607G3) and sMMO from *Methylosinus acidophilus* (NCBI accession AAY83388.1) with an e-value threshold of 10^−3^ and a %-id threshold of >30%. Genomes containing a methane monooxygenase sequence were subsequently mined for putative SQR sequences by blasting a representative sequence of each of the SQR subtypes as defined by previous research (Marcia et al., 2010): type I, WP_010961392.1; type II, WP_011001489.1; type III, WP_009059890.1; type IV, WP_011372252.1; type V, WP_012502121.1; type VI, WP_011439951.1. Putative SQR sequences were aligned with those in the phylogenetic tree of Marcia et al. (2010) using Muscle 3.8.1551 (Edgar, 2004) with default settings. A maximum-likelihood phylogenetic tree with 500 bootstrap replicates was constructed using RAxML 8.2.10 (Stamatakis, 2014) using the rapid bootstrapping option and the LG amino acid substitution model (Le and Gascuel, 2008). The final tree was visualized using MEGA7 and the clade of flavocytochrome *c* sulfide dehydrogenase (FCSD) sequences was used as outgroup.

### Microorganism and culturing

*Methylacidiphilum fumariolicum* SolV used in this study was isolated from a mud pot near Naples, Italy (Pol et al., 2007). The growth medium was composed of 0.2 mM MgCl_2_ · 6 H_2_O; 0.2 mM CaCl_2_ · 2 H_2_O; 1 mM Na_2_SO_4_; 2 mM K_2_SO_4_; 7.5 mM (NH_4_)_2_SO_4_ and 1 mM NaH_2_PO_4_ · H_2_O and trace elements at final concentrations of 1 μM NiCl_2_ 1 μM CoCl_2_, 1 μM MoO_4_Na_2_, 1 μM ZnSO_4_, 1 μM CeCl_3_, 5 μM MnCl_2_, 5 μM FeSO_4_, 10 μM CuSO_4_ and 40–50 μM nitrilotriacetic acid (NTA). Cells were grown as methane-limited continuous culture at 55 °C as described before (Schmitz et al., 2020), except that the pH was regulated at 2.5–3.0, that a small 400 mL chemostat was used with medium as described above and that H_2_ was not supplemented. The oxygen concentration was regulated at 1% air saturation. In addition, a second chemostat was operated under similar conditions, but to which H_2_S was added through an additional gas inlet (**Supplementary Figure S1**). H_2_S was produced by mixing 100 mM anoxic Na2S and 210 mM HCl in a 50 mL bottle with a peristaltic pump. The argon/CO_2_ (95%/5%, v/v) gas stream to the reactor was led through this bottle. The influx of H_2_S was regulated by using a peristaltic pump for mixing Na_2_S and HCl. Since H_2_S was supplied through the gas inlet and therefore needs to be transferred to the liquid phase, the liquid H_2_S concentration will be close to or lower than its equilibrium concentration, which at 55 °C is 1.6 times the gas concentration (calculated from the Ostwald coefficient at 55 °C; Wilhelm et al., 1977). In addition, to observe whether *M. fumariolicum* SolV can grow on H_2_S as sole energy source a batch culture was operated in the same setup as the chemostat system. In this case the medium flow was stopped, and the argon/CO_2_ gas was changed for an argon only gas stream. At the same time equal amounts of a ^13^C-labelled bicarbonate solution (50 mM) and HCl solution (100 mM) were additionally added to the sulfide mixing bottle, creating a ^13^C-CO_2_ gas concentration of about 2%. 5 mL biomass samples from the batch culture were collected by centrifugation over several days and the pellets were washed with acidified water (pH 3). Pellets were then resuspended in small amounts of acidified water and samples were subsequently pipetted into tin cups and dried overnight at 70 °C under vacuum. ^13^CO_2_ incorporation into biomass was assessed by measuring the ^13/12^C ratio using a Finnigan DeltaPlus isotope-ratio mass spectrometer (IR-MS) as described by Khadem et al. (2011).

### Membrane-inlet mass spectrometry and respiration

To accurately measure dissolved gases, membrane-inlet mass spectrometry (MIMS) was performed as described previously (Schmitz et al., 2020), except that a 30 mL MIMS chamber was used. The inserted probe consisted of a blunt end stainless steel tube (diameter 3 mm) that was perforated with 4–16 holes of 1 mm diameter. The holes were covered with silicon tubing (Silastic, 50VMQ Q7-4750 Dow Corning, supplied by Freudenberg Medical via VWR international, 1.96 mm outer diameter x 1.47 mm inner diameter). For easy mounting the silicon tubing was soaked shortly in hexane, which causes silicone to swell. The metal part was wetted with iso-propanol as lubricant. The probe was directly connected via a 1/8 or 1/16 inch stainless steel tube to the MS that was operated 40 μA emission current. Medium with a pH equal to that of the culture (pH 2.5–3.0) added to the chamber was first flushed with 3% CO_2_ in argon gas after which the oxygen concentration was adjusted to the desired value by adding pure oxygen gas or air via the headspace. Mass 15 and 16 are both dominant masses for CH_4_ in the mass spectrometer, but mass 15 has a much lower background signal than mass 16 and was therefore chosen to measure CH_4_. Methane and hydrogen (mass 2) were added as a gas in the headspace or, in the case of calibration, from a saturated stock solution. These stock solutions were prepared in a closed bottle with water at room temperature and a headspace of pure gas with known pressure. For the solubility in water 1.47 mM and 0.80 mM was taken for methane and hydrogen, respectively (at 22 °C and 1 bar; Wilhelm et al., 1977). When CO_2_ production rates were to be measured, ^13^C-bicarbonate and equimolar amounts of sulfuric acid were added after flushing with argon. In this way the simultaneously occurring CO_2_ fixation is mainly from ^13^C-labelled CO_2_ (mass 45), leading to less interference with CO_2_ production. At the start, unlabeled CO_2_ (mass 44) is very low, and its increase reflects almost exclusively CO_2_ production from unlabeled methane or methanol.

The stoichiometry of H_2_S oxidation was determined through pulse-wise additions of a sulfide stock solution and O_2_ (as a gas from a syringe) in order to keep concentrations low at 1–3 μM H_2_S and 0–5 μM O_2_. In total, 0.7–1.4 mM of Na_2_S was added over a period of 1.5–3 h. During this experiment, equimolar amounts of a sulfuric acid stock solution were added simultaneously to limit the pH change within 0.2 units. The oxygen concentration was simultaneously measured in the MIMS chamber by means of a fiber-optic oxygen sensor spot (TROXSP5, PyroScience, Aachen, Germany) that was glued on the inside of the chamber. These spots could measure down to about 20 nM oxygen, which is much lower than can be measured with the mass 32 signal of MIMS.

### Batch incubations and gas chromatography

To determine kinetic parameters of H_2_S oxidation by sulfide-adapted cells, batch incubations were performed in 120 mL serum bottles containing 10 mL medium without any trace elements. Trace elements were omitted to minimize the effect of abiotic sulfide oxidation. The bottles were closed with butyl rubber stoppers. Incubations were performed at 55 °C and 350 rpm. H_2_S was prepared by mixing Na_2_S with HCl in a closed bottle. A volume headspace was taken and injected into 120 mL serum bottles and equilibrated for 30 min before initiating the assay by addition of cells. H_2_S was measured by injecting 100 μL of the headspace of the bottles with a Hamilton glass syringe into a GC (7890B GC systems Agilent technologies, Santa Clara, USA) equipped with a Carbopack BHT100 glass column (2 m, ID 2 mm) and a flame photometric detector (FPD). The areas obtained were used to calculate H_2_S amounts using calibration standard curves with Na2S · 9 H_2_O (newly purchased from Sigma).

### RNA isolation and data analysis

For each replicate, 10 mL was sampled from the chemostat, and cells were immediately pelleted for 3 min at 15,000 × *g*, snap-frozen in liquid nitrogen and stored at –80 °C. Total RNA was isolated using the RiboPure™ RNA Purification Kit for bacteria (Thermo Fisher Scientific, Waltham, MA, USA) according to the manufacturer’s protocol. Ribosomal RNA was removed from the total RNA samples to enrich for mRNA using the MICROBExpress™ Bacterial mRNA Enrichment Kit (Thermo Fisher Scientific) according to the manufacturer’s protocol. The QubitTM RNA HS Assay Kit (Thermo Fisher Scientific) and the Agilent RNA 6000 Nano Kit (Agilent Technologies, Waldbronn, Germany) and protocols were used for the quantitative and qualitative analysis of the extracted total RNA and enriched mRNA. The latter was used for library preparation by using the TruSeq Stranded mRNA Reference Guide (Illumina, San Diego, CA, USA) according to the manufacturer’s protocol. For quantitative and qualitative assessment of the synthesized cDNA, the QubitTM dsDNA HS Kit (Thermo Fisher Scientific) and the Agilent High Sensitivity DNA kit (Agilent Technologies) and protocols were used. Transcriptome reads were checked for quality using FastQC (Andrews, 2010) and subsequently trimmed 10 base pairs at the 5’ end and 5 base pairs at the 3’ end of each read. Reads were mapped against the *M. fumariolicum* SolV complete genome (Anvar et al., 2014; accession number LM997411) using Bowtie2 (Langmead and Salzberg, 2012). The remainder of the analysis and the production of images was performed in version 4.0.2 of the R environment (R core team, 2020). The mapped read counts per gene were determined using Rsubread (Liao et al., 2019) and fold change and dispersion were estimated using DEseq2 (Love et al., 2014). Before doing any statistics, principal component analysis on the top 1000 genes by variance of each sample was performed to check whether samples within the same condition were both similar to each sample part of the same condition, and dissimilar to any other sample. For differential expression, a Wald test was employed by DEseq2 to calculate adjusted p-values. Differences in counts were considered to be significant if the basemean was higher than 4, the log_2_-fold change was higher than [0.58] and the adjusted p-value was ≤ 0.05. For easy comparisons between samples, TPM (Transcripts Per Kilobase Million) values were calculated.

### TOC measurements

The total organic carbon (TOC) concentrations of the cultures were determined using a TOC-L CPH/CPN analyzer (Shimadzu, Duisburg, Germany). Samples were diluted three times in Milli-Q water before measurements and subsequently sparged for 20 min with ozone while stirring to remove CO_2_ from the liquid. Acidification of the solutions was not needed due to the low pH of the samples. An optical density of 1 measured at 600 nm is equivalent to approximately 450 mg dry weight (DW) per liter.

## RESULTS

### Genes encoding putative SQRs are widespread in methanotrophs

The detection of a sulfide:quinone oxidoreductase (SQR) in the genomes of verrucomicrobial methanotrophs prompted us to investigate the H_2_S oxidation capacity of these methanotrophs (Schmitz et al., 2021). Interestingly, genes encoding a putative SQR also seem to be widespread in proteobacterial methanotrophs of various environments (**Figure 1**). Different types of putative SQRs were detected in a large variety of proteobacterial genera in which a pMMO and/or sMMO was present, such as *Methylobacter*, *Methylocaldum*, *Methylocapsa*, *Methylococcus*, *Methylocystis*, *Methylomagnum*, *Methylomarinum*, *Methylomicrobium*, *Methylomonas*, *Methyloprofundus*, *Methylosinus*, *Methylospira*, *Methyloterricola*, *Methylotetracoccus*, *Methylotuvimicrobium* and *Methylovulum*. In addition, the recently isolated alphaproteobacterium ‘*Methylovirgula thiovorans*’ strain HY1 encodes a type I SQR (Gwak et al., 2022). In contrast, verrucomicrobial methanotrophs possess genes encoding a type III SQR, comprising bacterial and archaeal SQRs of which the least is known (Marcia et al., 2010). Methanotrophs in general could benefit from encoding an SQR for the detoxification of H_2_S, as this compound coexists with CH_4_ in a large variety of ecosystems.

**Figure 1:**
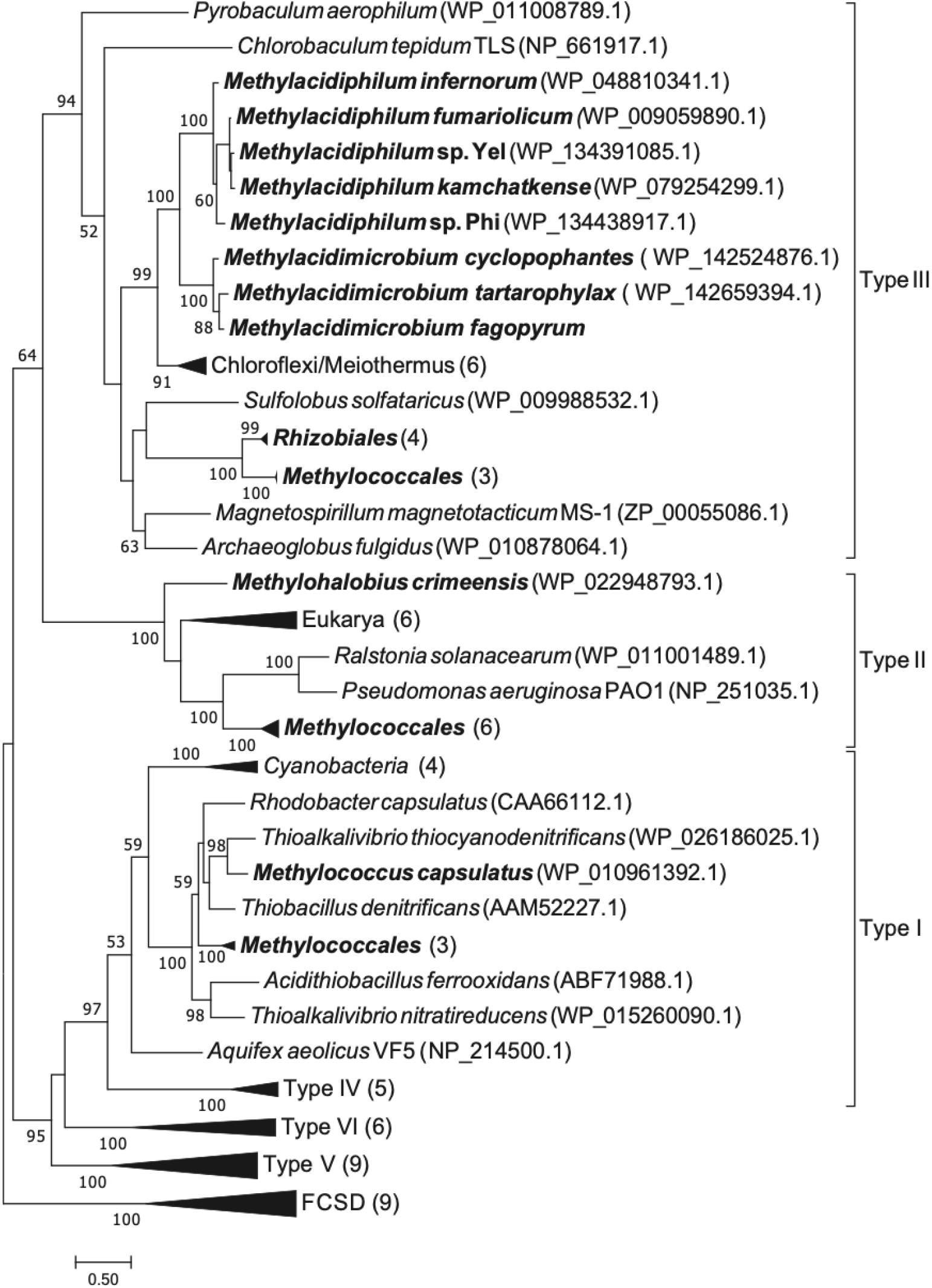
Phylogenetic tree of genes encoding different types of putative sulfide:quinone oxidoreductases (SQR) in a variety of microorganisms. A selection of microorganisms that also possess a particulate methane monooxygenase (pMMO) or soluble methane monooxygenase (sMMO) are indicated in bold.

### H_2_S inhibits methanotrophy in *M. fumariolicum* SolV

To investigate the effect of H_2_S on cells of *M. fumariolicum* SolV that were not yet adapted to H_2_S, a continuous culture was operated under low dissolved O_2_ concentrations (1% air saturation), receiving limiting amounts of CH_4_ (D = 0.016 h^−1^), resulting in a specific activity of 39 μmol CH_4_ · min^−1^ · g DW^−1^. To meticulously measure the conversion and respiration rate of various gases, cells of the continuous culture were studied with membrane-inlet mass spectrometry (MIMS). In the MIMS chamber, a maximum CH_4_ conversion rate of 200 ± 11 μmol CH_4_ · min^−1^ · g DW^−1^ was measured with a concomitant O_2_ consumption of 302 ± 9 μmol O_2_ · min^−1^ · g DW^−1^ (**Table 1**). Since one mole O_2_ is needed to activate one mole CH_4_, theoretically 102 μmol O_2_ · min^−1^ · g DW^−1^ should be used for respiration of CH_4_ to CO_2_. Clearly, the CH_4_ oxidation capacity of *M. fumariolicum* SolV cells was affected by H_2_S, from a concentration as low as 0.5 μM. At 10 μM H_2_S, the conversion of CH_4_ was inhibited by 95% and at 20 μM by over 99%.

**Table 1:**
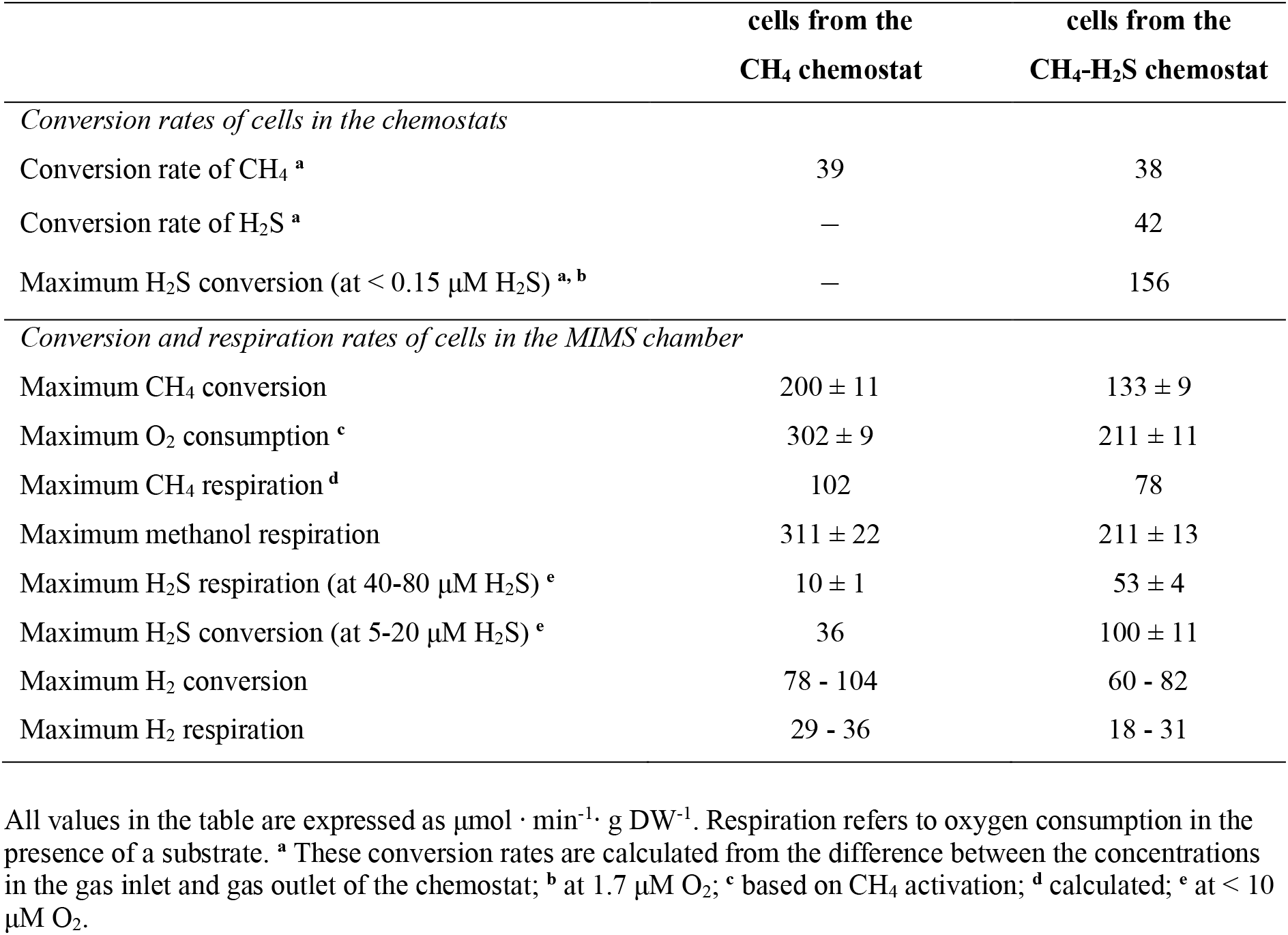
Comparison of conversion and respiration rates of *M. fumariolicum* SolV cells from the CH_4_ chemostat compared to the cells from the CH_4_/H_2_S chemostat.

High initial O_2_ consumption rates of 134–178 μmol O_2_ · min^−1^ · g DW^−1^ were measured when only H_2_S was administered to *M. fumariolicum* SolV cells in the MIMS chamber. However, these rates decreased rapidly within a few minutes. At high H_2_S concentrations (40*–* 80 μM H_2_S) and low O_2_ concentrations (< 10 μM O_2_) the final activity decreased to a stable value of 10 ± 1 μmol O_2_ · min^−1^ · g DW^−1^. This respiration rate suggested the presence of a sulfide-sensitive terminal oxidase (SSTO) quickly inactivated after addition of H_2_S and a sulfide-insensitive terminal oxidase (SITO) responsible for the remaining relatively low activity (Nicholls et al., 2013). When methanol (CH_3_OH) respiration of methane-grown *M. fumariolicum* SolV cells in the MIMS chamber was followed over time, a rate of 311 ± 22 μmol O_2_ · min^−1^ · g DW^−1^ was measured (**Table 1**). Hence, the maximum reaction rate of the SITO (10 ± 1 μmol O_2_ · min^−1^ · g DW^−1^) constitutes only 3% of the maximum respiration rates measured for the same culture with methanol. SITO activity was cyanide sensitive as 95% of the respiration rate was inhibited at 1 mM potassium cyanide. At 80 μM H_2_S and 80 μM O_2_ instead of low O_2_, the respiration rate increased from 10 to 14 μmol O_2_ · min^−1^ · g DW^−1^, suggesting that O_2_ is competing with H_2_S for the active site of the SSTO, thereby partly alleviating the H_2_S inhibition.

Interestingly, when methanol was added during respiration of >40 μM H_2_S by *M. fumariolicum* SolV in the MIMS chamber, H_2_S consumption ceased immediately while the respiration rate was not significantly altered. Conceivably, reducing equivalents of methanol oxidation were directed to the SITO and preferred over reducing equivalents yielded from H_2_S oxidation. In addition, in these incubations up to 7 μmol H_2_S · min^−1^ · g DW^−1^ was formed after addition of methanol, even in the presence of O_2_. This finding suggests the presence of a H_2_S-producing enzyme used by *M. fumariolicum* SolV, reducing theretofore produced polysulfides and/or elemental sulfur (S^0^; see results below) with methanol-derived reducing equivalents. Due to the low concentration of dissolved oxygen of the culture, *M. fumariolicum* SolV expressed a high hydrogenase activity of 78–104 μmol H_2_ · min^−1^ · g DW^−1^ (**Table 1**) (Mohammadi et al., 2017). The respiration rates with H_2_ as substrate were 29–36 μmol O_2_ · min^−1^ · g DW^−1^ (**Table 1**), as expected from an H_2_:O_2_ ratio of 1:0.35 (Mohammadi et al., 2017; Mohammadi et al., 2019). H_2_ conversion was lower in the presence of H_2_S, but H_2_ respiration (18–22 μmol O_2_ · min^−1^ · g DW^−1^) in the presence of 20 μM H_2_S continued at a level dependent on the SITO and the remaining SSTO activity due to incomplete inhibition. H_2_S conversion was low or absent in the presence of H_2_ and resumed after all H_2_ was consumed. The observation that reducing equivalents of H_2_ and methanol are preferred over those of H_2_S suggests a preference in SITO-dependent respiration for H_2_ and methanol over H_2_S. Similarly, consumption of a concentration of 20–40 μM H_2_S was reduced by about 80–90% upon addition of 200 μM formate (CHOOH), while the total respiration rate increased by 15%.

The H_2_S respiration rate at low (5–20 μM) H_2_S was about 11–18 μmol O_2_ · min^−1^ · g DW^−1^, which is higher than measured at concentrations above 40 μM H_2_S (10 ± 1 μmol O_2_ · min^−1^ · g DW^−1^) suggesting incomplete inhibition of SSTO at lower H_2_S concentrations. The observation that the respiration rate increased further to 22–27 μmol O_2_ · min^−1^ · g DW^−1^ in the presence of the additional substrates suggests that maximum H_2_S respiration was not only limited by activity of the terminal oxidases, but in a preceding metabolic step as well. Assuming that all H_2_S is converted to S^0^, this would result in a maximum H_2_S conversion capacity of 36 μmol H_2_S · min^−1^ · g DW^−1^ at 5–20 μM H_2_S by *M. fumariolicum* SolV cells not adapted to H_2_S (**Table 1**). The SITO activity of the culture was the same when the O_2_ concentration in the chemostat was increased from 1 to 20% air saturation, indicating that expression of SITO was not influenced by the O_2_ concentration.

### *M. fumariolicum* SolV efficiently detoxifies H_2_S to elemental sulfur after adaptation

To investigate whether *M. fumariolicum* SolV cells alter their physiology in response to H_2_S, cells in a continuous, methane-limited culture were exposed to gradually increasing concentrations of H_2_S over the course of a week (**Supplementary Figure S1**). Eventually, the H_2_S load was above that of CH_4_ and up to a point at which 38 μmol CH_4_ · min^−1^ · g DW^−1^ and 42 μmol H_2_S · min^−1^ · g DW^−1^ were consumed simultaneously in the chemostat (**Table 1**), while the H_2_S concentration in the gas outlet remained below 2 nmol/L. After a steady state in this dual CH_4_-H_2_S chemostat was achieved, MIMS experiments were performed to compare the sulfide-adapted cells with non-adapted cells (see section above). A stoichiometry of 1 H_2_S to 0.48 O_2_ (± 0.003; *n* = 4) was measured, which suggests conversion of H_2_S into elemental sulfur (S^0^). The turbidity in the MIMS chamber increased during H_2_S oxidation from 0.135 to 0.6 (measured at 600 nm) when a total of 1 mM H_2_S was converted, and microscopic particles (irregular rather than globular) were observed. As this process does not produce or consume protons, H_2_S addition accordingly did not lead to a significant pH change of the above culture that converted a sulfide concentration equivalent to 28 mM. The production of S^0^ was evident, as an increasing amount of yellow precipitate developed at the bottom and attached to the glass wall and metal parts of the chemostat (**Figure 2**). Formation of S^0^ occurred almost exclusively at these surfaces as sulfur particles were hardly observed in the culture liquid. Based on H_2_S additions to *M. fumariolicum* SolV cells in the MIMS chamber, maximum conversion rates were approximately 100 ± 11 μmol H_2_S · min^−1^ · g DW^−1^ when tested at 5–100 μM H_2_S and 5– 10 μM O_2_ while the maximum O_2_ consumption rates were 53 ± 4 μmol O_2_ · min^−1^ · g DW^−1^ (**Table 1**). As at 40 μM H_2_S the SSTO was shown to be almost completely inhibited, these values represent the rates of the SITO, which are more than five times higher compared to the non-adapted cells in the chemostat. Hence, the non-adapted cells and especially the sulfide-adapted cells efficiently detoxify H_2_S.

**Figure 2:**
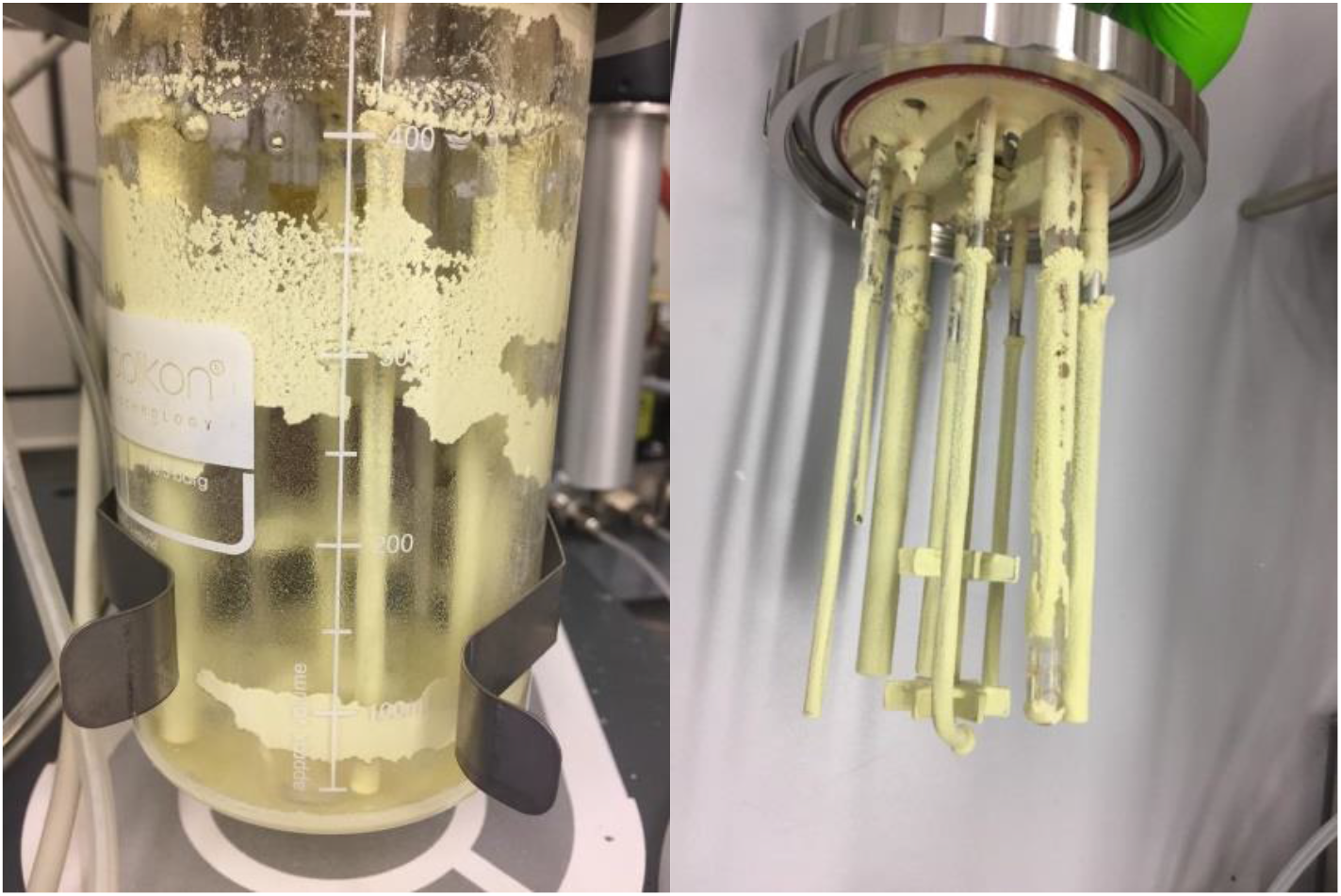
Yellow deposits of elemental sulfur from the oxidation of H_2_S by *Methylacidiphilum fumariolicum* SolV after growing for weeks in a continuous culture.

Above experiments were all conducted at low O_2_ concentrations in order to limit interference of chemical H_2_S oxidation. H_2_S oxidation rates increased with higher O_2_ concentrations (134–156 μmol H_2_S · min^−1^ · g DW^−1^ at 60–80 μM O_2_) measured in MIMS experiments. Again, this increase in rate can be attributed to competition between O_2_ and H_2_S at the active site of SSTO, at least partially alleviating H_2_S inhibition. Increasing non-specific biochemical and chemical reactions of H_2_S were considered as an explanation for increased rates but were dismissed since similarly high rates could also be obtained in the chemostat culture itself at very low O_2_ concentrations (1% air saturation) while keeping H_2_S at low, non-inhibiting concentrations. These conditions were obtained by gradually increasing the H_2_S to a final H_2_S load of 156 μmol H_2_S · min^−1^ · g DW^−1^. As a result, H_2_S in the outlet had only increased from 2 to 25 nmol/L and therefore remained below a liquid concentration of 40 nM, which was considered not to affect pMMO (as measured through MIMS incubations). At this high H_2_S load, CH_4_ conversion had decreased with about 40% but remained stable. When the maximum H_2_S conversion rate was reached, H_2_S in the outlet had increased to 0.1 μmol/L while CH_4_ conversion remained stable. When in a similar way the chemostat was given H_2_S only while methane was disconnected, the same maximum H_2_S conversion activity of 156 μmol H_2_S · min^−1^ · g DW^−1^ was measured. As respiration is not the limiting factor in this setup (**Table 1**), this rate seems to be the maximum H_2_S conversion rate and equal to the SITO activity. The above measured maximum H_2_S consumption rate in the MIMS exceeded that of CH_4_ conversion (133 ± 9 μmol CH_4_ · min^−1^ · g DW^−1^). From the oxygen consumption rate a CH_4_ respiration rate of 78 μmol O_2_ · min^−1^ · g DW^−1^ could be calculated. This respiration rate was about 24% lower compared to the cells of the chemostat culture receiving CH_4_ only (102 μmol O_2_ · min^−1^ · g DW^−1^). Accordingly, the increased H_2_S oxidation capacity seems to be at the expense of the CH_4_ conversion capacity. CH_4_ conversion in MIMS incubations was affected in a similar way as for the culture on CH_4_ only. Inhibition was about 60% when H_2_S was added around detection level of 1 μM and increased to about 97% at 10 μM H_2_S and 99% at 20 μM.

The effect of H_2_S on methanol oxidation of the sulfide-adapted cells was measured through MIMS and compared with the non-adapted cells. As for CH_4_, the initial maximum methanol respiration rates (211 ± 13 μmol O_2_ · min^−1^ · g DW^−1^) were about 30% lower than measured for the culture grown on CH_4_ only (**Table 1**). The rates dropped immediately upon addition of 20–30 μM H_2_S, but the remaining activity (69 μmol O_2_ · min^−1^ · g DW^−1^) was higher than expected from the maximum H_2_S respiration rate (53 ± 4 μmol O_2_ · min^−1^ · g DW^−1^), indicating that at least some methanol, as opposed to methane at these sulfide concentrations, was still respired. This was confirmed by the CO_2_ production rate that continued at 20–30%. Conversely, when the MIMS experiment was started with H_2_S, the conversion rate decreased about 50% when methanol was added. Hence, methanol and H_2_S were respired simultaneously and seem to compete for the same terminal oxidase. Similarly, H_2_ conversion continued for 70–80% after H_2_S additions and SITO activity would be high enough to account for maximum H_2_ conversion rates of 82 μmol H_2_ · min^−1^ · g DW^−1^ (**Table 1**). Inhibition of CH_4_ conversion seemed reversible. When H_2_S was consumed or flushed out of a short-term incubation the CH_4_ conversion and CO_2_ production from methanol resumed immediately at their previous rates. After longer periods (2 h) of inhibition by H_2_S (10–20 μM), CH_4_ conversion rates were 25– 35% lower. It could not be concluded that the lower rates were caused by inhibition of pMMO or elsewhere in the respiratory chain as also methanol conversion (as measured from O_2_ consumption and CO_2_ production) was impaired after such H_2_S exposures. When O_2_ and H_2_S additions were stopped after these H_2_S oxidation periods and the incubation became anoxic, H_2_S production occurred as before but at a slower rate of about 4 μmol H_2_S produced · min^−1^ · g DW^−1^. H_2_S production was stimulated when methanol or H_2_ gas was present up to 13 μmol H_2_S produced · min^−1^ · g DW^−1^, presumably from sulfur/polysulfide previously accumulated.

### Kinetics of H_2_S oxidation and respiration

The oxidation of 10–25 μM H_2_S measured in the MIMS chamber revealed a quite constant oxidation rate down to 2 μM H_2_S and dropped sharply below 1 μM, suggesting an apparent affinity constant for H_2_S at or even below 0.5 μM, which was the detection level. Therefore, kinetics of H_2_S oxidation was also studied in shaking bottles of which the headspace was measured by GC analyses. Starting at 4 μM H_2_S, an almost linear decrease down to 1 μM H_2_S was observed with a rate of 167–223 μmol H_2_S · min^−1^ · g DW^−1^ (**Figure 3a**). These rates are slightly higher than the maximum sulfide conversion rates measured in the MIMS chamber, which could be explained by the ambient air present in the serum bottles for GC measurements. Michaelis-Menten modelling of the H_2_S consumption resulted in an apparent affinity of 0.32 μM H_2_S. However, the H_2_S trace below 1 μM H_2_S did not follow the predicted curve and remained well above it, presumably due to respiration limitations. When the O_2_ consumption was followed down to zero at H_2_S concentrations of 15–20 μM the oxygen consumption followed Michaelis-Menten kinetics and resulted in an apparent affinity constant of 0.14 ± 0.01 μM O_2_ (**Figure 3b**). This suggests the rate is not limited by SQR, but possibly by respiration at the higher sulfide concentrations, when SSTO becomes inhibited. Consequently, the apparent affinity constant would be higher. Assuming only one terminal oxidase to be active under these conditions and H_2_S conversion not being the limiting factor, this may be considered a reliable value for the sulfide-insensitive terminal oxidase.

**Figure 3:**
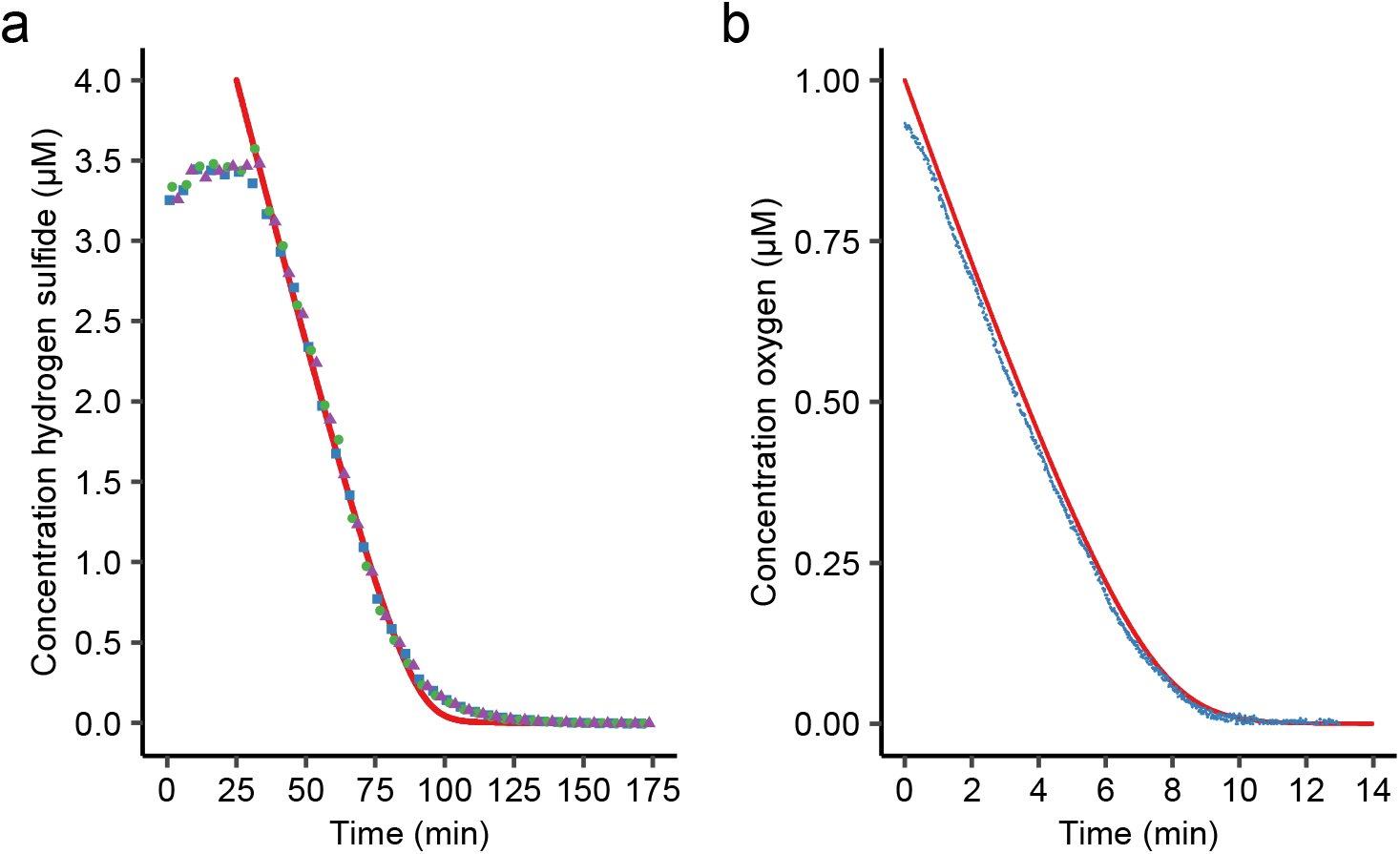
Kinetics of (**a**) H_2_S consumption measured in serum bottles through GC. Cells of *M. fumariolicum* SolV were added 33 min after the start of the experiment. (**b**) H_2_S respiration measured through a fiber-optic oxygen sensor spot. Different symbols in (**a**) indicate different replicates (*n* = 3). Red lines indicate calculated Michaelis-Menten curve fitting.

### *M. fumariolicum* SolV alters its electron transport chain in response to H_2_S

To understand which genes and enzymes are involved in the adaption to H_2_S, mRNA from the CH_4_ and the CH_4_-H_2_S continuous culture was isolated and gene expression was analyzed (**Table 2 and Supplementary File S1**). In the presence of H_2_S, the operon Mfumv2_0219-21 is upregulated about 1.7-fold. The genes in this operon are annotated as NAD(FAD)-dependent dehydrogenase (Mfumv2_0219), a protein homologous to the sulfur carrier protein TusA (Mfumv2_0220) and a possible sulfur carrier protein DsrE2 (Mfumv2_0221), respectively. A more detailed investigation of the upregulated gene Mfumv2_0219 revealed this gene to encode a type III sulfide:quinone oxidoreductase (SQR) (**Figure 1**) that could be responsible for the observed H_2_S oxidation (Marcia et al., 2010). The genes Mfumv2_0942 and Mfumv2_0943 were upregulated 2-fold and 8-fold and their products show high similarity to the cytochrome *c* protein SorB and sulfite:cytochrome *c* oxidoreductase SorA of *Thiobacillus novellus*, respectively (Kappler et al., 2000). An enzymatic pathway from elemental sulfur to sulfite (SO_3_^2−^) could not be resolved, but sulfite might also be produced chemically. The operon Mfumv2_1257-61 encodes a cytochrome *ba*_3_-type oxidase that was upregulated approximately 5-fold in the presence of H_2_S. Since through MIMS experiments it was shown that SITO respiration was five times higher in sulfide-adapted cells, it is conceivable that this *ba*_3_-type oxidase is the dedicated sulfide-insensitive terminal oxidase. Interestingly, the highest upregulated gene (15-fold) is an 89 kDa heptahaem cytochrome *c* protein of unknown function (Mfumv2_1950), which shows highest similarity to genes found in thermophilic sulfide-oxidizers. In the presence of H_2_S, several genes encoding enzymes involved in the biosynthesis of H_2_S for production of sulfur-containing metabolites were heavily downregulated (**Table 2**). In addition, downregulation of genes involved in CH_4_ oxidation and subsequent electron transfer in the respiratory chain was observed (**Table 2**). The downregulation of genes involved in methane respiration is in accordance with a decreased maximum methane conversation and respiration rate in the chemostat with high H_2_S influx.

**Table 2:** Regulation of *Methylacidiphilum fumariolicum* SolV genes of cells grown with CH_4_ and H_2_S in comparison with cells grown with CH_4_ as sole energy source. Listed genes have a basemean higher than 4, an upregulation or downregulation factor (averages of triplicates) higher than 1.5 and an adjusted p-value ≤ 0.05.

| ORF | Annotation | Upregulation factor |
| --- | --- | --- |
| ***Genes involved in the oxidation of sulfur compounds and respiratory chain*** |  |  |
| Mfumv2_0219 | NAD(FAD)-dependent dehydrogenase (hcaD) | 1.81 |
| Mfumv2_0220 | Putative sulfur carrier protein | 1.70 |
| Mfumv2_0221 | Peroxioredoxin family protein | 1.66 |
| Mfumv2_0942 | Cytochrome *c* family protein | 1.93 |
| Mfumv2_0943 | Sulfite oxidase or related enzyme | 8.33 |
| Mfumv2_1257 | Conserved protein of unknown function | 4.47 |
| Mfumv2_1258 | Conserved protein of unknown function | 5.44 |
| Mfumv2_1259 | Cytochrome *c* oxidase (B(O/a)3-type) chain II | 5.23 |
| Mfumv2_1260 | Cytochrome *c* oxidase subunit 1 (cbaA) | 4.46 |
| Mfumv2_1261 | Conserved protein of unknown function | 3.16 |
| Mfumv2_1950 | Hepta-haem-containing protein | 15.73 |
| Mfumv2_1951 | Putative starvation-inducible outer membrane lipoprotein | 9.70 |
| ORF | Annotation | Downregulation factor |
| ***Genes involved in assimilatory H<sub>2</sub>S production*** |  |  |
| Mfumv2_0525 | Sulfate adenylyltransferase subunit 1 | 11.63 |
| Mfumv2_0526 | Sulfate adenylyltransferase subunit 2 (cysD) | 18.65 |
| Mfumv2_0527 | Phosphoadenosine phosphosulfate reductase (cysH) | 29.39 |
| Mfumv2_0528 | Homocitrate synthase 1 (nifV) | 8.59 |
| Mfumv2_0573 | Polysulfide reductase | 2.17 |
| Mfumv2_0815 | Sulfite reductase [NADPH] hemoprotein beta-component (cysI) | 33.40 |
| Mfumv2_2220 | O-acetylserine sulfhydrylase A (cysK) | 3.01 |
| ***Genes involved in methane oxidation*** |  |  |
| Mfumv2_1464 | PqqA peptide cyclase PqqE | 2.08 |
| Mfumv2_1604 | Methane monooxygenase subunit alpha (PmoB3) | 3.55 |
| Mfumv2_1605 | Methane monooxygenase subunit beta (PmoA3) | 2.95 |
| Mfumv2_1606 | Methane monooxygenase subunit gamma (PmoC3) | 2.88 |
| Mfumv2_1791 | Methane monooxygenase subunit alpha (PmoB1) | 1.64 |
| Mfumv2_1792 | Methane monooxygenase subunit beta (PmoA1) | 1.70 |
| Mfumv2_1793 | Methane monooxygenase (PmoC1) | 1.62 |
| ***Genes involved in the respiratory chain*** |  |  |
| Mfumv2_2064 | Succinate dehydrogenase flavoprotein subunit | 1.58 |
| Mfumv2_2065 | Succinate dehydrogenase cytochrome *b* subunit | 2.49 |
| Mfumv2_2239 | NADH-quinone oxidoreductase subunit D (nuoD) | 1.58 |
| Mfumv2_2240 | NADH-quinone oxidoreductase subunit C (nuoC) | 1.64 |
| Mfumv2_2241 | NADH-quinone oxidoreductase subunit B (nuoB) | 1.59 |
| Mfumv2_2458 | ATP synthase F1 complex subunit alpha (atpA) | 1.66 |
| Mfumv2_2459 | ATP synthase F1 complex subunit gamma (atpG) | 1.62 |
| Mfumv2_2460 | ATP synthase F1 complex subunit beta (atpD) | 1.56 |
| Mfumv2_1602 | Phosphoenolpyruvate synthetase (ppsA) | 2.66 |

### *M. fumariolicum* SolV incorporates CO_2_ with H_2_S as sole energy source

Through physiological chemostat measurements, MIMS and transcriptomics, *M. fumariolicum* SolV was shown to efficiently detoxify H_2_S to S^0^. Consequently, electrons derived from H_2_S could be used for energy conservation and autotrophic growth. To assess incorporation of CO_2_ with H_2_S as sole energy source, a batch reactor inoculated with a diluted culture (OD_600_ = 0.05) from the CH_4_-H_2_S chemostat was used. H_2_S and ^13^CO_2_ were added and their fate was followed over time, while CH_4_ was not supplemented. A sulfide load close to the maximum capacity (147 μmol H_2_S · min^−1^ · g DW^−1^) was applied. Clearly, *M. fumariolicum* SolV fixes ^13^CO_2_ by using H_2_S as sole energy source, since the percentage ^13^C-labeled biomass increased over time (**Figure 4**). This increase in ^13^C-containing biomass was accompanied by an increase in optical density (measured at 600 nm). Through microscopy, in the liquid only minute amounts of sulfur particles were observed as opposed to bacterial cells. When the H_2_S load was increased about 2-fold, the ^13^C incorporation rate also approximately doubled and the optical density increased sharply. By measuring the gas inlet and outlet of the reactor throughout the incubation period, an H_2_S conversion efficiency of approximately 100% to 98% could be determined.

**Figure 4:**
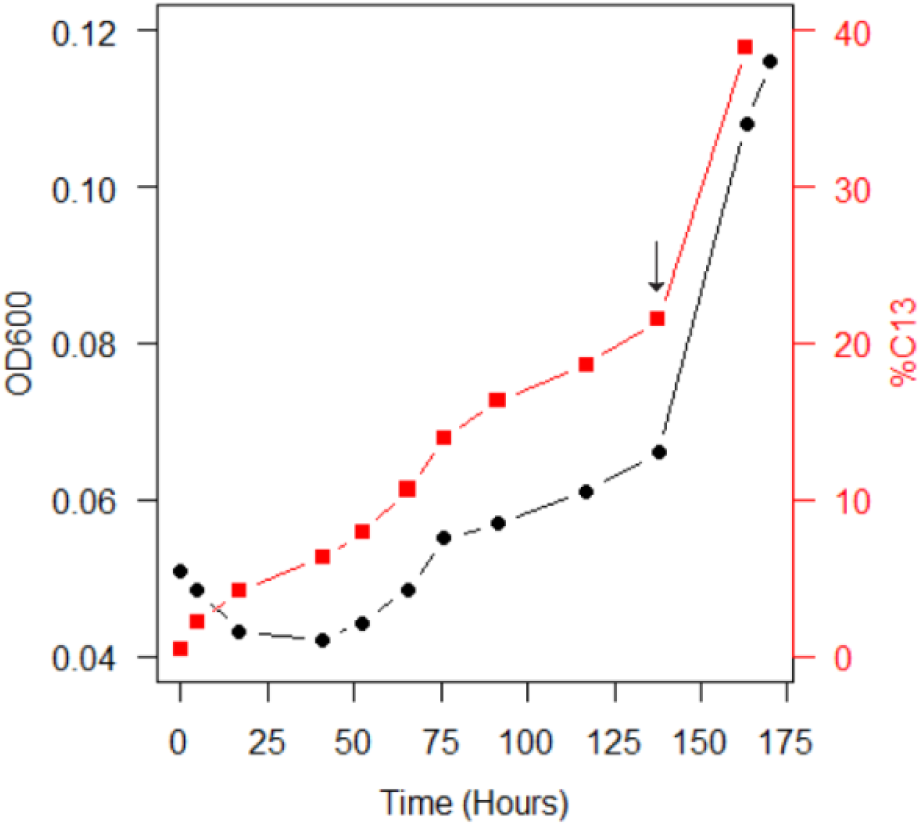
Increase in OD_600_ and percentage ^13^C in the biomass over time by cells of *Methylacidiphilum fumariolicum* SolV in fed-batch mode with H_2_S as sole energy source and ^13^CO_2_ as carbon source. The arrow indicates the time point at which the H_2_S load was almost doubled.

## DISCUSSION

In this study, we showed that the thermoacidophilic methanotroph *Methylacidiphilum fumariolicum* SolV efficiently detoxifies H_2_S to elemental sulfur. At low pH, sulfide is present in fully protonated state and can easily pass the membrane bilayer. Detoxification is a necessity since we showed that H_2_S inhibits both respiration and methane oxidation. Cells respond to the presence of H_2_S by upregulating a type III sulfide:quinone oxidoreductase (SQR) and a sulfide-insensitive *ba*_3_-type oxidase (SITO). To the best of our knowledge, no one has shown before that a microorganism can oxidize CH_4_ and H_2_S simultaneously, and that a methanotroph can produce biomass from CO_2_ with H_2_S as sole energy source. In addition, we provide proof for an H_2_S detoxification mechanism in methanotrophs, which, according to genomic information and the co-occurrence of methane and sulfide in a myriad of environments, could be much more widespread.

Understanding the involvement of microorganisms in biogeochemical cycles is important in future climate modelling (Cavicchioli et al., 2019). By oxidizing methane, microorganisms mitigate the emissions of this strong greenhouse gas from various environments (Stein, 2020). Typically, geothermal environments are characterized by morphological features such as fumaroles, hot springs and mud pools that emit CH_4_ and H_2_S (Picone et al., 2020a; Schmitz et al., 2021). All genomes of known isolated verrucomicrobial methanotrophs and many different proteobacterial methanotrophs possess a gene encoding a putative SQR. This is not surprising, since CH_4_ and H_2_S coexist in many environments. Remarkably, very little is known about the effect of H_2_S on methanotrophy. A methanotrophic consortium sampled from a landfill showed decreased methanotrophic activity in the presence of H_2_S (Lee et al., 2015). In addition, methane oxidation by *Methylocaldum* sp. SAD2, isolated from a sulfide-rich anaerobic digester, was slightly inhibited (44-60% decrease in methanol production) in the presence of 0.1% H_2_S, but the mechanism was not explored (Zhang et al., 2016; Wei et al., 2017). ‘*Methylovirgula thiovorans*’ strain HY1A was recently shown to be able to consume various reduced sulfur compounds together with CH_4_, but a simultaneous oxidation of CH_4_ and H_2_S could not be observed (Gwak et al., 2022). The H_2_S concentration was below the detection limit in the peatland where strain HY1A was isolated from, suggesting that a vigorous H_2_S detoxification might not be necessary. In contrast, the geothermal environment where *M. fumariolicum* SolV thrives, is characterized by high concentrations of sulfide (from < 50 ppm to 20000 ppm; Tedesco and Sabroux, 1987; Pol et al., 2007; Picone et al., 2020a). In this light, the observation that *M. fumariolicum* SolV fixes CO_2_ with H_2_S as sole energy source and efficiently oxidizes H_2_S to S^0^ could be highly advantageous in such harsh systems. Considering that in the natural environment multiple substrates are present at the same time rather than a single energy source, a mixotrophic lifestyle is expected, in which CH_4_, H_2_ and H_2_S are utilized simultaneously (Carere et al., 2017; Mohammadi et al., 2019).

H_2_S is known to bind to metals such as copper and iron, which could lead to inhibition of the CH_4_ oxidation capacity of the copper-dependent pMMO and terminal oxidases involved in the reduction of O_2_ (Bagarinao, 1992; Pietri et al., 2011; Barton et al., 2014; Landry et al., 2021). Interestingly, ‘*Methylovirgula thiovorans*’ strain HY1A only encodes an iron-dependent sMMO (Gwak et al., 2022), whereas *M. fumariolicum* SolV encodes multiple copper-dependent pMMOs (Schmitz et al., 2021). The former strain does not simultaneously oxidize H_2_S and CH_4_, while the latter has a rapid H_2_S detoxification system to alleviate inhibition of methanotrophy. Hence, the extent to which a type of methane monooxygenase is inhibited could influence the need for an H_2_S detoxification system. Since in *M. fumariolicum* SolV the gene encoding a type III SQR was upregulated in the presence of H_2_S, we propose that this enzyme is responsible for the observed oxidation of H_2_S to elemental sulfur. Indeed, type III SQRs were shown to couple the oxidation of H_2_S to the reduction of quinones in several archaea and bacteria (Lencina et al., 2013; Han and Perner, 2016). Still, purification of this enzyme is necessary to accurately understand its kinetics and how the catalysis is coupled to the quinone pool. In verrucomicrobial methanotrophs, three different types of terminal oxidases are found: an *aa*_3_-type, a *ba*_3_-type, and a *cbb*_3_-type (Schmitz et al., 2021). Possessing multiple types of terminal oxidases allows a branched electron transport chain, which is highly advantageous in environments with fluctuating conditions and varying substrate and oxygen availability. Through respiration studies, we showed that *M. fumariolicum* SolV possesses one or more sulfide-sensitive terminal oxidases (SSTO) and a sulfide-insensitive terminal oxidase (SITO). Since a *ba*_3_-type terminal oxidase is upregulated in cells growing at high H_2_S loads, we propose this specific enzyme complex to be the dedicated SITO in verrucomicrobial methanotrophs. Similarly, in sulfur-grown cells of *Acidithiobacillus ferrooxidans* this *ba*_3_-type oxidase was upregulated (Brasseur et al., 2004). It is tempting to speculate on the upregulation of the multihaem cytochrome *c* protein in *M. fumariolicum* SolV, which may be involved as electron carrier from SQR to the electron transport chain. This putative electron carrier might explain why sulfide respiration still partly continues upon addition of methanol in the sulfide-adapted cells and not in the non-adapted cells. In non-adapted cells, the putative multihaem electron carrier could be the limiting factor for sulfide respiration, being overruled by the relatively large amounts of the electron carrier XoxG, mediating electron transfer from methanol to the terminal oxidase. In contrast, in sulfide-adapted cells the ratio between the putative multihaem electron carrier and XoxG is much larger than in non-adapted cells, and sulfide respiration can occur concurrently with methanol oxidation, using the same terminal oxidase.

As CH_4_ conversion was almost completely inactivated in the presence of 20 μM H_2_S while methanol, formate and H_2_ conversion could still proceed at the same H_2_S concentration, it can be concluded that H_2_S is impeding both the SSTO and the reaction catalyzed by pMMO. The observed decrease in CH_4_ conversion in the chemostat at high an H_2_S load of 156 μmol H_2_S · min^−1^ · g DW^−1^ was much more than can be expected from inhibition studies and may indicate that a large portion of the respiratory chain is used for electrons generated by H_2_S oxidation. *M. fumariolicum* SolV converts H_2_S to elemental sulfur, but the reducing equivalents produced from this oxidation seem to muddle the electron transport chain. Similarly, it was proposed that an overactive SQR in *Rhodobacter capsulatus* could lead to an overreduction of the quinone pool (Schütz et al., 1999). The upregulated *ba*3-type oxidase could alleviate this problem by oxidizing quinol and reducing the terminal electron acceptor O_2_. In *Aquifex aeolicus*, a related *ba*_3_-type oxidase was found in a supercomplex with SQR (Prunetti et al., 2010). This terminal oxidase was shown to not only oxidize reduced cytochrome *c*, but also ubiquinol directly (Gao et al., 2012). A branched electron transport chain with different terminal oxidases enables metabolic versatility and adaptations. For example, *E. coli* uses the proton-pumping *bo*_3_-type oxidase during growth but requires the sulfide-insensitive *bd*-type oxidases to keep growing in the presence of H_2_S (Forte et al., 2016). The measured stoichiometry of 1 H_2_S : 0.48 O_2_ in MIMS experiments and the visible production of elemental sulfur clearly show that H_2_S is not oxidized further. In line with this, downstream genes involved in the oxidation of elemental sulfur to sulfite are absent in strain SolV. Whether genes involved in this pathway were lost during evolution or never acquired by verrucomicrobial methanotrophs remains to be investigated.

Cells of *M. fumariolicum* SolV were shown to rapidly oxidize H_2_S with a high apparent affinity constant below 1 μM H_2_S. The observed kinetic values are not surprising, since H_2_S already inhibits methanotrophy at such low concentrations. Through gas chromatography, an exact apparent affinity constant for whole cells could not be determined, as H_2_S consumption did not follow a typical Michaelis-Menten curve. A limitation of the respiratory capacity for H_2_S oxidation above about 1 μM may explain such deviation and could be resolved by purification of SQR. The observation that *M. fumariolicum* SolV reduced elemental sulfur to H_2_S in the presence of H_2_ or methanol is intriguing. ‘*Methylovirgula thiovorans*’ strain HY1A grown on thiosulfate increasingly produced an enzyme that resembles a protein known to possess sulfhydrogenase activity (Gwak et al., 2022). Interestingly, this enzyme clusters with the group 3b [NiFe] hydrogenase of *M. fumariolicum* SolV, thought to be involved in the production of NAD(P)H for CO_2_ fixation (Carere et al., 2017). Indeed, it is proposed that these hydrogenases can have sulfhydrogenase activity, which could be a mechanism to dispose of reducing equivalents (Ma et al., 1993; Søndergaard et al., 2016). Hence, the group 3b [NiFe] hydrogenase of *M. fumariolicum* SolV might be responsible for the conversion of S^0^ to H_2_S. Culturing in a continuous culture again proved to be a very powerful tool to investigate the metabolism of methanotrophs (Mohammadi et al., 2017; Carere et al., 2021). Through adaption, *M. fumariolicum* SolV was able to respire H_2_S at a rate five times higher than non-adapted cells, presumably due to the upregulation of SQR and the *ba*_3_-type terminal oxidase.

Verrucomicrobial methanotrophs that thrive in geothermal environments have a clear mechanism to cope with H_2_S. Accordingly, SQR and a sulfide-insensitive terminal oxidase could enable these methanotrophs to thrive in H_2_S-rich environments. Indeed, pyrosequencing showed that *Methylacidimicrobium*-related 16S rRNA gene sequences were abundantly present in the crown of concrete sewage pipes rich in CH_4_ and H_2_S (Pagaling et al., 2014). Concerning proteobacterial methanotrophs, the effect of H_2_S warrants further investigation. Since aerobic methanotrophs live in environments in which H_2_S is often present, we propose that the mechanism of H_2_S detoxification is widespread in methanotrophs in various environments.

## AUTHOR CONTRIBUTIONS

RAS, SSM, AP and HJMOdC designed the project and experiments. RAS, SSM, TvE, CAI, TvA, WV and AP conducted the experiments. TB conducted phylogenetic analyses. RAS, SHP, SSM, TvE, CAI and AP performed data analyses. RAS, MSMJ, HJMOdC and AP wrote the manuscript. RAS, MSMJ, HJMOdC and AP supervised the research.

## ACKNOWLEDGEMENTS

We want to thank Dr. Marianne Guiral and Dr. Frauke Baymann (CNRS, Aix-Marseille University, Marseille, France) for fruitful discussions.

## SUPPLEMENTARY INFORMATION

**Supplementary Figure 1:**
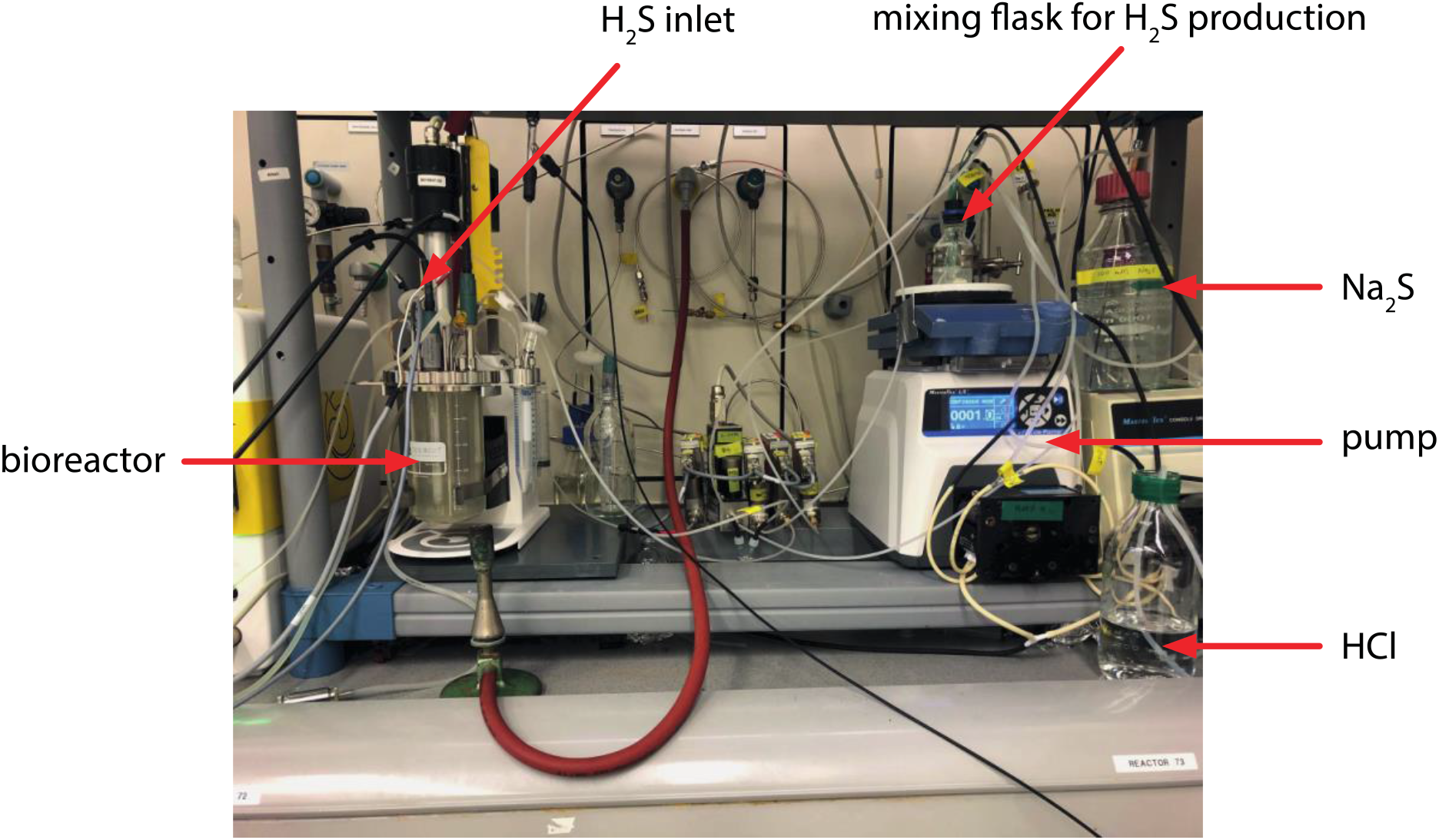
Culturing of *Methylacidiphilum fumariolicum* SolV in a continuous culture on H_2_S and CH_4_. H_2_S is produced externally in a mixing flask by mixing sodium sulfide (Na_2_S) and hydrochloric acid (HCl) and fed into the chemostat through a gas inlet.

