## Supplementary File S1 for "Methanotrophs are vigorous H_2_S oxidizers using a sulfide:quinone oxidoreductase and a *ba*_3_-type terminal oxidase"

| locus.tag | baseMean | log2FoldChange | lfcSE | stat | pvalue | padj |
|---|---|---|---|---|---|---|
| Mfumv2_0001 | 636.9937784 | -0.022534492 | 0.136365 | -0.16525 | 0.868746 | 0.939349 |
| Mfumv2_0002 | 678.3503298 | -0.249651531 | 0.115083 | -2.16933 | 0.030058 | 0.114096 |
| Mfumv2_0003 | 160.3069359 | 0.036577691 | 0.186446 | 0.196184 | 0.844466 | 0.926055 |
| Mfumv2_0004 | 0 | NA | NA | NA | NA | NA |
| Mfumv2_0005 | 18.22034109 | 0.325361894 | 0.415559 | 0.78295 | 0.433656 | NA |
| Mfumv2_0006 | 261.1488522 | 0.083832294 | 0.155605 | 0.53875 | 0.590059 | 0.793808 |
| Mfumv2_0007 | 140.9586241 | 0.016186916 | 0.181368 | 0.089249 | 0.928884 | 0.972652 |
| Mfumv2_0008 | 178.7140796 | 0.182962951 | 0.163717 | 1.117558 | 0.263756 | 0.501785 |
| Mfumv2_0009 | 921.956769 | 0.429900547 | 0.128839 | 3.336728 | 0.000848 | 0.007278 |
| Mfumv2_0010 | 588.2583813 | 0.016067254 | 0.126896 | 0.126618 | 0.899243 | 0.954347 |
| Mfumv2_0011 | 529.5893716 | -0.006944235 | 0.118172 | -0.05876 | 0.95314 | 0.981476 |
| Mfumv2_0012 | 148.4189096 | 0.11697844 | 0.226429 | 0.516623 | 0.605419 | 0.80336 |
| Mfumv2_0013 | 87.93988091 | 0.243533746 | 0.242929 | 1.002491 | 0.316107 | 0.55903 |
| Mfumv2_0014 | 433.8555041 | 0.081994602 | 0.128009 | 0.64054 | 0.521822 | 0.743504 |
| Mfumv2_0015 | 1734.728954 | -0.160727161 | 0.095173 | -1.68879 | 0.09126 | 0.253962 |
| Mfumv2_0016 | 522.6863307 | -0.758169692 | 0.14544 | -5.21293 | 1.86E-07 | 4.72E-06 |
| Mfumv2_0017 | 2948.591633 | -0.377171914 | 0.136863 | -2.75583 | 0.005854 | 0.033224 |
| Mfumv2_0018 | 1057.99336 | -0.151172208 | 0.104501 | -1.44661 | 0.148006 | 0.342958 |
| Mfumv2_0019 | 834.1355659 | 0.087889546 | 0.128614 | 0.683357 | 0.494381 | 0.722336 |
| Mfumv2_0020 | 21.48058566 | 0.071655426 | 0.42022 | 0.170519 | 0.864602 | NA |
| Mfumv2_0021 | 468.9062225 | 0.055660905 | 0.118444 | 0.469933 | 0.638403 | 0.823866 |
| Mfumv2_0022 | 431.5638071 | 0.056161001 | 0.117253 | 0.478971 | 0.631959 | 0.82074 |
| Mfumv2_0023 | 60.46930312 | -0.039344502 | 0.241568 | -0.16287 | 0.87062 | 0.940363 |
| Mfumv2_0024 | 674.0846969 | -0.255235989 | 0.118645 | -2.15126 | 0.031456 | 0.117342 |
| Mfumv2_0025 | 467.3130923 | -0.0315565 | 0.118156 | -0.26707 | 0.789412 | 0.897706 |
| Mfumv2_0026 | 633.9603822 | -0.34021269 | 0.114328 | -2.97577 | 0.002923 | 0.019545 |
| Mfumv2_0027 | 274.089617 | -0.334188598 | 0.153543 | -2.17651 | 0.029517 | 0.112732 |
| Mfumv2_0028 | 48.46906249 | -0.402306846 | 0.287309 | -1.40026 | 0.161436 | 0.368134 |
| Mfumv2_0029 | 210.6094306 | -0.575739092 | 0.170286 | -3.38101 | 0.000722 | 0.006448 |
| Mfumv2_0030 | 188.5333145 | -0.583666406 | 0.249248 | -2.34171 | 0.019196 | 0.084201 |
| Mfumv2_0031 | 175.0438387 | 0.025073634 | 0.157233 | 0.159468 | 0.8733 | 0.942751 |
| Mfumv2_0032 | 276.1425736 | -0.409382151 | 0.179178 | -2.28478 | 0.022326 | 0.09432 |
| Mfumv2_0033 | 132.8821673 | 0.328527662 | 0.176292 | 1.86354 | 0.062386 | 0.192822 |
| Mfumv2_0034 | 17.69277523 | 0.593857473 | 0.42683 | 1.391322 | 0.164128 | NA |
| Mfumv2_0035 | 5.742666062 | 0.520259191 | 0.75095 | 0.692802 | 0.488434 | NA |
| Mfumv2_0036 | 231.2410346 | -0.38551082 | 0.150136 | -2.56775 | 0.010236 | 0.05217 |
| Mfumv2_0037 | 181.7294493 | -0.242207349 | 0.158374 | -1.52934 | 0.126181 | 0.311041 |
| Mfumv2_0038 | 9.530099087 | 0.420678344 | 0.59822 | 0.703217 | 0.481921 | NA |
| Mfumv2_0039 | 633.1262411 | -0.009640804 | 0.129639 | -0.07437 | 0.940719 | 0.977742 |
| Mfumv2_0040 | 72.9379726 | 0.644155192 | 0.272861 | 2.360743 | 0.018238 | 0.080885 |
| Mfumv2_0041 | 383.7721793 | 0.015340083 | 0.450623 | 0.034042 | 0.972844 | 0.990093 |
| Mfumv2_0042 | 7.991829394 | 0.42519168 | 0.636744 | 0.66776 | 0.504287 | NA |
| Mfumv2_0043 | 532.11599 | -0.327202253 | 0.116238 | -2.81492 | 0.004879 | 0.029346 |
| Mfumv2_0044 | 433.6336839 | 0.038744228 | 0.127191 | 0.304614 | 0.76066 | 0.884498 |
| Mfumv2_0045 | 1051.041862 | 0.120425468 | 0.116948 | 1.029737 | 0.303133 | 0.544232 |
| Mfumv2_0046 | 110.7787049 | 0.385535895 | 0.202607 | 1.902877 | 0.057057 | 0.181031 |
| Mfumv2_0047 | 352.0219728 | 0.481779695 | 0.14515 | 3.319191 | 0.000903 | 0.007685 |
| Mfumv2_0048 | 233.8413303 | 0.140284565 | 0.152607 | 0.919252 | 0.357964 | 0.59836 |
| Mfumv2_0049 | 420.2516478 | 0.074126744 | 0.129949 | 0.57043 | 0.568386 | 0.778914 |
| Mfumv2_0050 | 154.6288109 | 0.09497915 | 0.163593 | 0.580582 | 0.561522 | 0.77267 |

| | | | | | | | |
|---|---|---|---|---|---|---|---|
| Mfumv2_0051 | 110.9686666 | | -0.195983954 | 0.192865 | -1.01617 | 0.309548 | 0.551803 |
| Mfumv2_0052 | 238.7028421 | | -0.040454518 | 0.143046 | -0.28281 | 0.777324 | 0.892523 |
| Mfumv2_0053 | 297.4966169 | | 0.206739989 | 0.133475 | 1.548908 | 0.121404 | 0.304876 |
| Mfumv2_0054 | 389.5596161 | | 0.062171767 | 0.121918 | 0.509946 | 0.610089 | 0.806892 |
| Mfumv2_0055 | 58.25563598 | | 0.512478175 | 0.25106 | 2.041257 | 0.041225 | 0.14304 |
| Mfumv2_0056 | 10.32977349 | | 0.163442107 | 0.542242 | 0.301419 | 0.763095 | NA |
| Mfumv2_0057 | 2.592360259 | | -0.298557353 | 1.058567 | -0.28204 | 0.777914 | NA |
| Mfumv2_0058 | 0 | NA | | NA | NA | NA | NA |
| Mfumv2_0059 | 0.223916542 | | -1.639710324 | 3.825436 | -0.42863 | 0.66819 | NA |
| Mfumv2_0060 | 1127.769285 | | -1.056741977 | 0.14802 | -7.13918 | 9.39E-13 | 5.55E-11 |
| Mfumv2_0061 | 437.3028835 | | -0.887670247 | 0.156404 | -5.67551 | 1.38E-08 | 4.37E-07 |
| Mfumv2_0062 | 286.3566468 | | -0.886612312 | 0.157811 | -5.6182 | 1.93E-08 | 5.79E-07 |
| Mfumv2_0063 | 109.480757 | | -0.835281346 | 0.194399 | -4.29675 | 1.73E-05 | 0.00029 |
| Mfumv2_0064 | 354.3583101 | | -0.821247354 | 0.137691 | -5.96442 | 2.46E-09 | 9.31E-08 |
| Mfumv2_0065 | 61.69179851 | | -0.048848009 | 0.244406 | -0.19986 | 0.841587 | 0.924411 |
| Mfumv2_0066 | 48.82775522 | | -0.187711456 | 0.282891 | -0.66355 | 0.50698 | 0.732316 |
| Mfumv2_0067 | 175.4451567 | | -0.156846604 | 0.256684 | -0.61105 | 0.541167 | 0.76074 |
| Mfumv2_0068 | 5.722784193 | | 0.986196975 | 0.827014 | 1.192479 | 0.233073 | NA |
| Mfumv2_0069 | 0.651796466 | | 2.489940654 | 2.123985 | 1.172297 | 0.241078 | NA |
| Mfumv2_0070 | 55.04736123 | | 0.437357377 | 0.270348 | 1.617756 | 0.105715 | 0.282156 |
| Mfumv2_0071 | 150.580419 | | 0.098480759 | 0.202796 | 0.485614 | 0.627241 | 0.819861 |
| Mfumv2_0073 | 35.60685573 | | 0.483394561 | 0.311164 | 1.553504 | 0.120303 | 0.304013 |
| Mfumv2_0074 | 1391.413285 | | -0.088639016 | 0.099538 | -0.8905 | 0.373197 | 0.610542 |
| Mfumv2_0075 | 1013.391423 | | 0.062291699 | 0.119222 | 0.522483 | 0.601334 | 0.801694 |
| Mfumv2_0076 | 1684.8113 | | -0.982777022 | 0.339175 | -2.89755 | 0.003761 | 0.024063 |
| Mfumv2_0077 | 593.7251022 | | -0.715506269 | 0.154291 | -4.63737 | 3.53E-06 | 7.02E-05 |
| Mfumv2_0078 | 337.544397 | | -0.898872262 | 0.16782 | -5.35617 | 8.50E-08 | 2.31E-06 |
| Mfumv2_0079 | 137.9496733 | | -0.099394679 | 0.197702 | -0.50275 | 0.615139 | 0.81037 |
| Mfumv2_0080 | 72.85094249 | | 0.09807307 | 0.234062 | 0.419005 | 0.675213 | 0.842548 |
| Mfumv2_0081 | 58.15098766 | | 0.507006419 | 0.244734 | 2.071664 | 0.038297 | 0.136699 |
| Mfumv2_0082 | 178.7254188 | | 0.09390623 | 0.169481 | 0.554082 | 0.579523 | 0.788261 |
| Mfumv2_0083 | 72.6462913 | | 0.221463861 | 0.230458 | 0.960975 | 0.336565 | 0.579896 |
| Mfumv2_0084 | 374.2168573 | | -0.062417298 | 0.135206 | -0.46164 | 0.644336 | 0.82661 |
| Mfumv2_0085 | 472.1494838 | | -0.121055394 | 0.122293 | -0.98988 | 0.322234 | 0.566462 |
| Mfumv2_0086 | 758.3831832 | | -0.158086093 | 0.131259 | -1.20438 | 0.228443 | 0.465042 |
| Mfumv2_0087 | 750.5211797 | | 0.059304956 | 0.124883 | 0.474885 | 0.634869 | 0.822746 |
| Mfumv2_0088 | 1124.361523 | | -0.058066764 | 0.143714 | -0.40404 | 0.686181 | 0.845728 |
| Mfumv2_0089 | 539.8108496 | | -0.120168255 | 0.131336 | -0.91497 | 0.360207 | 0.598929 |
| Mfumv2_0090 | 104.5859051 | | -0.051382219 | 0.195402 | -0.26296 | 0.792584 | 0.898083 |
| Mfumv2_0091 | 1445.586609 | | -0.408200589 | 0.117255 | -3.48132 | 0.000499 | 0.004847 |
| Mfumv2_0092 | 478.0509526 | | -0.207039393 | 0.124891 | -1.65776 | 0.097365 | 0.267588 |
| Mfumv2_0093 | 968.1448602 | | -0.25183557 | 0.108065 | -2.33041 | 0.019785 | 0.085709 |
| Mfumv2_0094 | 354.7535879 | | -0.500181313 | 0.134922 | -3.7072 | 0.00021 | 0.002365 |
| Mfumv2_0095 | 1536.492429 | | -0.10766597 | 0.110839 | -0.97138 | 0.331361 | 0.573388 |
| Mfumv2_0096 | 88.3578946 | | 0.466876851 | 0.216397 | 2.157498 | 0.030967 | 0.116503 |
| Mfumv2_0097 | 0.140690113 | | 0.283845181 | 4.080473 | 0.069562 | 0.944542 | NA |
| Mfumv2_0098 | 0 | NA | | NA | NA | NA | NA |
| Mfumv2_0099 | 2.321755906 | | 2.323489646 | 1.404672 | 1.654115 | 0.098104 | NA |
| Mfumv2_0100 | 0 | NA | | NA | NA | NA | NA |
| Mfumv2_0101 | 29.88867005 | | 0.734330183 | 0.347668 | 2.112157 | 0.034673 | 0.126651 |
| Mfumv2_0102 | 169.0974345 | | 0.296169143 | 0.162932 | 1.817752 | 0.069102 | 0.20941 |
| Mfumv2_0104 | 151.0041109 | | 0.131864143 | 0.176473 | 0.747219 | 0.454931 | 0.683588 |

| | | | | | | |
|---|---|---|---|---|---|---|
| Mfumv2_0105 | 150.7678946 | 0.294445796 | 0.169613 | 1.73599 | 0.082566 | 0.235617 |
| Mfumv2_0106 | 354.5712377 | 0.105366696 | 0.128804 | 0.818041 | 0.413334 | 0.647422 |
| Mfumv2_0107 | 5.689015796 | 0.579801451 | 0.769633 | 0.753348 | 0.451241 | NA |
| Mfumv2_0109 | 1182.438352 | -0.680838654 | 0.170389 | -3.99579 | 6.45E-05 | 0.000881 |
| Mfumv2_0110 | 20.63464056 | 0.430193505 | 0.411201 | 1.046187 | 0.295475 | NA |
| Mfumv2_0111 | 1582.575801 | 0.300591256 | 0.105147 | 2.858772 | 0.004253 | 0.026534 |
| Mfumv2_0112 | 1097.972549 | 0.242222029 | 0.113018 | 2.143216 | 0.032096 | 0.118968 |
| Mfumv2_0113 | 1335.174555 | -0.215330098 | 0.143424 | -1.50135 | 0.133265 | 0.321018 |
| Mfumv2_0114 | 41.72771275 | 0.441528767 | 0.288114 | 1.532478 | 0.125404 | 0.311034 |
| Mfumv2_0115 | 334.2566692 | -0.321792207 | 0.174672 | -1.84226 | 0.065437 | 0.200706 |
| Mfumv2_0116 | 70.85587561 | 0.456866719 | 0.227015 | 2.012495 | 0.044168 | 0.149634 |
| Mfumv2_0117 | 77.66179401 | 0.548111856 | 0.227526 | 2.409003 | 0.015996 | 0.072871 |
| Mfumv2_0118 | 209.004824 | -0.082653273 | 0.153523 | -0.53838 | 0.590318 | 0.793808 |
| Mfumv2_0120 | 143.0828119 | 0.150825678 | 0.175345 | 0.860164 | 0.389699 | 0.622615 |
| Mfumv2_0121 | 176.3436787 | 0.19063258 | 0.156494 | 1.218146 | 0.223169 | 0.457963 |
| Mfumv2_0122 | 144.0263955 | 0.052026979 | 0.170659 | 0.304859 | 0.760474 | 0.884498 |
| Mfumv2_0123 | 51.02461777 | -0.295733374 | 0.272411 | -1.08561 | 0.27765 | 0.517918 |
| Mfumv2_0124 | 629.7688284 | -0.400229994 | 0.132014 | -3.03172 | 0.002432 | 0.016904 |
| Mfumv2_0125 | 55.18174608 | 0.556248862 | 0.274153 | 2.028975 | 0.042461 | 0.146046 |
| Mfumv2_0126 | 4.724573927 | 0.957866385 | 0.943374 | 1.015362 | 0.309933 | NA |
| Mfumv2_0127 | 323.7945106 | -0.019373562 | 0.134777 | -0.14375 | 0.885701 | 0.947191 |
| Mfumv2_0128 | 2523.607417 | -0.328359804 | 0.112811 | -2.91071 | 0.003606 | 0.023146 |
| Mfumv2_0129 | 825.398179 | -0.038396424 | 0.113159 | -0.33931 | 0.734373 | 0.872254 |
| Mfumv2_0130 | 2444.136171 | 0.127743649 | 0.120733 | 1.05807 | 0.290023 | 0.527769 |
| Mfumv2_0131 | 2098.025579 | -0.101103808 | 0.104479 | -0.9677 | 0.333195 | 0.576066 |
| Mfumv2_0132 | 1034.40919 | 0.095069779 | 0.159166 | 0.597298 | 0.550308 | 0.76593 |
| Mfumv2_0133 | 669.2377901 | -0.300479628 | 0.362344 | -0.82927 | 0.406954 | 0.641232 |
| Mfumv2_0134 | 409.1135576 | -0.283399899 | 0.131473 | -2.15557 | 0.031117 | 0.116632 |
| Mfumv2_0135 | 379.9810887 | 0.021161403 | 0.139373 | 0.151833 | 0.879319 | 0.945691 |
| Mfumv2_0136 | 240.6830324 | -0.303493292 | 0.142776 | -2.12566 | 0.033532 | 0.123482 |
| Mfumv2_0137 | 0.703434705 | -2.046170433 | 2.190034 | -0.93431 | 0.350144 | NA |
| Mfumv2_0138 | 142.1367385 | -0.084290125 | 0.193586 | -0.43541 | 0.663262 | 0.835454 |
| Mfumv2_0139 | 239.6113051 | -0.54394469 | 0.155752 | -3.49237 | 0.000479 | 0.004761 |
| Mfumv2_0140 | 377.5442115 | -0.103326485 | 0.133608 | -0.77335 | 0.439313 | 0.669636 |
| Mfumv2_0141 | 306.8825137 | -0.004784648 | 0.15281 | -0.03131 | 0.975021 | 0.990165 |
| Mfumv2_0142 | 882.0406195 | -0.403568128 | 0.122494 | -3.29459 | 0.000986 | 0.008149 |
| Mfumv2_0143 | 208.8684581 | 0.060037548 | 0.152602 | 0.393426 | 0.694005 | 0.849183 |
| Mfumv2_0144 | 310.1415178 | 0.173391002 | 0.145202 | 1.194138 | 0.232424 | 0.468815 |
| Mfumv2_0145 | 155.3594323 | 0.016546668 | 0.18189 | 0.09097 | 0.927516 | 0.972652 |
| Mfumv2_0146 | 2741.062451 | -0.070769468 | 0.099029 | -0.71464 | 0.474834 | 0.701944 |
| Mfumv2_0147 | 249.8036688 | 0.037423141 | 0.149756 | 0.249894 | 0.802669 | 0.903903 |
| Mfumv2_0148 | 327.0293598 | 0.251185281 | 0.127695 | 1.96707 | 0.049175 | 0.163294 |
| Mfumv2_0149 | 379.5110487 | 0.076023743 | 0.139392 | 0.545395 | 0.585482 | 0.791543 |
| Mfumv2_0150 | 105.327358 | 0.062777522 | 0.18892 | 0.332296 | 0.739665 | 0.875859 |
| Mfumv2_0151 | 336.2910868 | 0.282330572 | 0.130551 | 2.1626 | 0.030572 | 0.115449 |
| Mfumv2_0152 | 0 | NA | NA | NA | NA | NA |
| Mfumv2_0153 | 0.682231785 | 1.19019749 | 2.048169 | 0.581103 | 0.561171 | NA |
| Mfumv2_0155 | 9.192359397 | 1.286962077 | 0.624458 | 2.060928 | 0.03931 | NA |
| Mfumv2_0156 | 8.426490642 | 0.406277577 | 0.613642 | 0.662076 | 0.507922 | NA |
| Mfumv2_0157 | 17.2132504 | -0.060396719 | 0.419339 | -0.14403 | 0.885478 | NA |
| Mfumv2_0158 | 17.39018992 | 0.363718484 | 0.450578 | 0.807227 | 0.419536 | NA |
| Mfumv2_0159 | 2.558750227 | 0.431450223 | 1.135492 | 0.379968 | 0.703969 | NA |

| | | | | | | |
|---|---|---|---|---|---|---|
| Mfumv2_0161 | 25.59676364 | 0.136932091 | 0.353926 | 0.386894 | 0.698834 | 0.850369 |
| Mfumv2_0162 | 0 | NA | NA | NA | NA | NA |
| Mfumv2_0163 | 14.83191737 | 0.843218873 | 0.481224 | 1.752236 | 0.079733 | NA |
| Mfumv2_0165 | 3.445352096 | -0.713633829 | 0.932849 | -0.765 | 0.444269 | NA |
| Mfumv2_0166 | 9.001124896 | 0.949538392 | 0.617442 | 1.537858 | 0.124083 | NA |
| Mfumv2_0167 | 329.9487062 | 0.25505003 | 0.131414 | 1.940807 | 0.052282 | 0.169683 |
| Mfumv2_0168 | 80.39670281 | 0.089530928 | 0.208582 | 0.429235 | 0.667752 | 0.83651 |
| Mfumv2_0169 | 3066.531113 | -1.00936894 | 0.125125 | -8.06687 | 7.21E-16 | 5.17E-14 |
| Mfumv2_0170 | 233.2014177 | -0.638214314 | 0.163714 | -3.89835 | 9.69E-05 | 0.001239 |
| Mfumv2_0171 | 10.74291488 | 0.109891393 | 0.536683 | 0.20476 | 0.837759 | NA |
| Mfumv2_0172 | 251.0442864 | -0.225937715 | 0.153169 | -1.47509 | 0.14019 | 0.331845 |
| Mfumv2_0173 | 284.6374666 | 0.347731791 | 0.141465 | 2.45807 | 0.013969 | 0.066186 |
| Mfumv2_0174 | 220.6621862 | -0.510462531 | 0.151684 | -3.3653 | 0.000765 | 0.006708 |
| Mfumv2_0175 | 79.36070023 | 0.383663524 | 0.217118 | 1.767074 | 0.077216 | 0.226506 |
| Mfumv2_0176 | 69.87296593 | 0.165879904 | 0.229412 | 0.723067 | 0.469639 | 0.697712 |
| Mfumv2_0177 | 54.48625114 | 0.417070982 | 0.256427 | 1.626474 | 0.103849 | 0.279294 |
| Mfumv2_0178 | 2.968912379 | 0.298604183 | 1.094263 | 0.272882 | 0.784944 | NA |
| Mfumv2_0179 | 1.338891152 | 2.256972675 | 1.740608 | 1.296658 | 0.194749 | NA |
| Mfumv2_0180 | 21.26618099 | -0.514853653 | 0.391975 | -1.31349 | 0.189019 | NA |
| Mfumv2_0181 | 0 | NA | NA | NA | NA | NA |
| Mfumv2_0182 | 7.894268306 | -0.374441466 | 0.645495 | -0.58008 | 0.561858 | NA |
| Mfumv2_0183 | 30.14586451 | -0.09795295 | 0.370813 | -0.26416 | 0.791659 | 0.897706 |
| Mfumv2_0184 | 151.9631466 | 0.510684368 | 0.200585 | 2.545973 | 0.010897 | 0.054732 |
| Mfumv2_0185 | 14.80515582 | 0.25117971 | 0.462776 | 0.542768 | 0.58729 | NA |
| Mfumv2_0186 | 122.5548314 | -0.080663159 | 0.203476 | -0.39643 | 0.691791 | 0.847961 |
| Mfumv2_0187 | 195.3507906 | -0.184804766 | 0.212353 | -0.87027 | 0.384152 | 0.617611 |
| Mfumv2_0188 | 62.46696433 | 0.520598776 | 0.250585 | 2.077531 | 0.037753 | 0.135864 |
| Mfumv2_0189 | 29.38178017 | -0.101340181 | 0.33286 | -0.30445 | 0.760783 | 0.884498 |
| Mfumv2_0190 | 254.8124016 | -0.279571921 | 0.154903 | -1.80482 | 0.071103 | 0.214484 |
| Mfumv2_0191 | 2.075419266 | 0.89678937 | 1.326568 | 0.676022 | 0.499027 | NA |
| Mfumv2_0192 | 11.5128306 | -0.468832316 | 0.583665 | -0.80326 | 0.421827 | NA |
| Mfumv2_0193 | 6407.157645 | -0.428878278 | 0.133591 | -3.21039 | 0.001326 | 0.010403 |
| Mfumv2_0194 | 395.9385646 | -0.24039916 | 0.121523 | -1.97823 | 0.047903 | 0.159863 |
| Mfumv2_0195 | 90.99114009 | 0.462160392 | 0.216412 | 2.135553 | 0.032716 | 0.120913 |
| Mfumv2_0196 | 590.74393 | -0.044119482 | 0.144865 | -0.30456 | 0.760704 | 0.884498 |
| Mfumv2_0197 | 229.1109707 | 0.146759203 | 0.170094 | 0.862813 | 0.38824 | 0.621494 |
| Mfumv2_0198 | 408.992568 | -0.365071253 | 0.152809 | -2.38907 | 0.016891 | 0.075578 |
| Mfumv2_0199 | 1567.755035 | -0.054369967 | 0.107352 | -0.50647 | 0.61253 | 0.808872 |
| Mfumv2_0200 | 853.2494676 | -1.10859441 | 0.177235 | -6.25494 | 3.98E-10 | 1.78E-08 |
| Mfumv2_0202 | 354.3634742 | -1.274888737 | 0.182391 | -6.98986 | 2.75E-12 | 1.54E-10 |
| Mfumv2_0203 | 477.2475845 | -1.362930716 | 0.153078 | -8.9035 | 5.41E-19 | 4.18E-17 |
| Mfumv2_0204 | 325.4505198 | 0.162825049 | 0.169791 | 0.958974 | 0.337572 | 0.581133 |
| Mfumv2_0205 | 8.594121595 | 0.502038943 | 0.603611 | 0.831726 | 0.405564 | NA |
| Mfumv2_0206 | 117.0007644 | 0.148662023 | 0.20074 | 0.74057 | 0.458954 | 0.687278 |
| Mfumv2_0207 | 159.1034297 | -0.457148086 | 0.172144 | -2.65561 | 0.007916 | 0.042984 |
| Mfumv2_0208 | 9.211463135 | 0.423805635 | 0.609262 | 0.695604 | 0.486677 | NA |
| Mfumv2_0209 | 101.2453219 | -0.279425087 | 0.2107 | -1.32618 | 0.184782 | 0.40507 |
| Mfumv2_0210 | 475.6052243 | 0.098111524 | 0.129217 | 0.759279 | 0.447686 | 0.677564 |
| Mfumv2_0211 | 1602.116281 | 0.125037354 | 0.120111 | 1.041015 | 0.297869 | 0.53718 |
| Mfumv2_0212 | 3.371635919 | 0.009876769 | 0.932095 | 0.010596 | 0.991546 | NA |
| Mfumv2_0213 | 18.23665523 | 0.730070954 | 0.428801 | 1.702587 | 0.088645 | NA |
| Mfumv2_0214 | 280.4983349 | -0.189221827 | 0.196069 | -0.96508 | 0.334506 | 0.57734 |

| | | | | | | |
|---|---|---|---|---|---|---|
| Mfumv2_0215 | 13.53276763 | 0.146256236 | 0.501185 | 0.291821 | 0.770423 | NA |
| Mfumv2_0216 | 270.3011722 | 0.289328695 | 0.146919 | 1.969311 | 0.048917 | 0.162707 |
| Mfumv2_0217 | 295.0130594 | 0.030805337 | 0.153403 | 0.200814 | 0.840844 | 0.924411 |
| Mfumv2_0218 | 8.686022339 | 0.49825901 | 0.621108 | 0.80221 | 0.422432 | NA |
| Mfumv2_0219 | 1347.341572 | 0.853107615 | 0.121635 | 7.013641 | 2.32E-12 | 1.33E-10 |
| Mfumv2_0220 | 423.4537348 | 0.762708096 | 0.140012 | 5.447466 | 5.11E-08 | 1.47E-06 |
| Mfumv2_0221 | 1737.381 | 0.731360634 | 0.157544 | 4.64227 | 3.45E-06 | 6.92E-05 |
| Mfumv2_0222 | 364.2482116 | 0.064494152 | 0.142357 | 0.453045 | 0.650516 | 0.829769 |
| Mfumv2_0223 | 58.54248799 | 0.057228755 | 0.247279 | 0.231434 | 0.816978 | 0.909463 |
| Mfumv2_0224 | 218.4841544 | 0.177647391 | 0.152344 | 1.166092 | 0.243577 | 0.48259 |
| Mfumv2_0226 | 48.70488069 | 0.652379488 | 0.268278 | 2.431729 | 0.015027 | 0.069709 |
| Mfumv2_0227 | 49.04416465 | 0.377961224 | 0.276894 | 1.365002 | 0.172252 | 0.385806 |
| Mfumv2_0228 | 120.4071647 | 0.014235836 | 0.206905 | 0.068804 | 0.945146 | 0.979433 |
| Mfumv2_0229 | 240.9240327 | 0.07613653 | 0.165639 | 0.459653 | 0.645766 | 0.827387 |
| Mfumv2_0230 | 744.7622736 | 0.249212204 | 0.153742 | 1.620978 | 0.105022 | 0.281695 |
| Mfumv2_0231 | 95.33608036 | -0.040988324 | 0.19635 | -0.20875 | 0.834642 | 0.920306 |
| Mfumv2_0232 | 814.9140644 | 0.016450068 | 0.114792 | 0.143303 | 0.886051 | 0.947191 |
| Mfumv2_0233 | 131.8831075 | 0.356358484 | 0.183432 | 1.942726 | 0.052049 | 0.169476 |
| Mfumv2_0234 | 391.1461205 | -0.106587996 | 0.178444 | -0.59732 | 0.550295 | 0.76593 |
| Mfumv2_0235 | 182.4461698 | 0.121998717 | 0.161457 | 0.755609 | 0.449884 | 0.679561 |
| Mfumv2_0236 | 1394.254896 | -0.02089161 | 0.107565 | -0.19422 | 0.846001 | 0.927178 |
| Mfumv2_0239 | 210.2300806 | 0.290483777 | 0.152075 | 1.910138 | 0.056115 | 0.17838 |
| Mfumv2_0240 | 119.576157 | 0.056047281 | 0.204096 | 0.274613 | 0.783614 | 0.89556 |
| Mfumv2_0241 | 64.54947275 | -0.000871996 | 0.251745 | -0.00346 | 0.997236 | 0.99869 |
| Mfumv2_0242 | 103.5943496 | -0.00954686 | 0.202998 | -0.04703 | 0.96249 | 0.987056 |
| Mfumv2_0243 | 221.8548211 | -0.061894563 | 0.142881 | -0.43319 | 0.664878 | 0.835454 |
| Mfumv2_0244 | 153.3748666 | 0.09243958 | 0.18296 | 0.505244 | 0.613388 | 0.808872 |
| Mfumv2_0245 | 104.1993207 | 0.346857206 | 0.2016 | 1.720522 | 0.085338 | 0.24181 |
| Mfumv2_0246 | 301.7066252 | 0.173843116 | 0.157987 | 1.100363 | 0.271174 | 0.510102 |
| Mfumv2_0247 | 228.483477 | -0.593017607 | 0.185451 | -3.19771 | 0.001385 | 0.010787 |
| Mfumv2_0248 | 250.2932234 | -0.668855372 | 0.18918 | -3.53555 | 0.000407 | 0.00413 |
| Mfumv2_0249 | 196.9802603 | -0.42589551 | 0.206025 | -2.0672 | 0.038715 | 0.137517 |
| Mfumv2_0250 | 1577.212079 | 0.085230819 | 0.096434 | 0.883824 | 0.376791 | 0.611746 |
| Mfumv2_0251 | 7351.74556 | 0.023455322 | 0.117162 | 0.200195 | 0.841328 | 0.924411 |
| Mfumv2_0252 | 517.8201683 | -0.611813713 | 0.118518 | -5.16222 | 2.44E-07 | 5.98E-06 |
| Mfumv2_0253 | 479.7037658 | -0.587055669 | 0.114587 | -5.12323 | 3.00E-07 | 7.10E-06 |
| Mfumv2_0254 | 754.2005196 | 0.057390068 | 0.127019 | 0.451822 | 0.651398 | 0.82984 |
| Mfumv2_0255 | 1959.304164 | -0.22898586 | 0.106489 | -2.15033 | 0.03153 | 0.117342 |
| Mfumv2_0256 | 1359.532514 | -0.209462737 | 0.107098 | -1.9558 | 0.050489 | 0.16683 |
| Mfumv2_0257 | 387.5336498 | 0.11914624 | 0.134329 | 0.886975 | 0.375092 | 0.61116 |
| Mfumv2_0258 | 2248.406033 | -0.295418822 | 0.142928 | -2.06691 | 0.038743 | 0.137517 |
| Mfumv2_0259 | 1234.938873 | -0.185627886 | 0.115043 | -1.61356 | 0.106624 | 0.283025 |
| Mfumv2_0260 | 179.3223764 | -0.301661507 | 0.179492 | -1.68064 | 0.092832 | 0.257954 |
| Mfumv2_0261 | 52.01622744 | 0.350948499 | 0.277141 | 1.266316 | 0.2054 | 0.434665 |
| Mfumv2_0262 | 140.2270875 | -0.197282782 | 0.20074 | -0.98278 | 0.325716 | 0.568517 |
| Mfumv2_0263 | 602.4640975 | 0.061131558 | 0.127179 | 0.480675 | 0.630748 | 0.82074 |
| Mfumv2_0264 | 1735.319851 | 0.00202387 | 0.103984 | 0.019463 | 0.984472 | 0.992996 |
| Mfumv2_0266 | 901.5377349 | -0.026135967 | 0.108681 | -0.24048 | 0.809955 | 0.907508 |
| Mfumv2_0267 | 116.3958796 | 0.466530643 | 0.303251 | 1.538428 | 0.123944 | 0.308846 |
| Mfumv2_0268 | 27.21187851 | -0.232899687 | 0.367529 | -0.63369 | 0.526283 | 0.748798 |
| Mfumv2_0269 | 143.1312878 | 0.186593291 | 0.177987 | 1.048353 | 0.294476 | 0.533937 |
| Mfumv2_0270 | 631.0718477 | -0.01905786 | 0.130805 | -0.1457 | 0.884161 | 0.946455 |

| | | | | | | |
|---|---|---|---|---|---|---|
| Mfumv2_0271 | 279.4020812 | 0.008990013 | 0.150286 | 0.05982 | 0.952299 | 0.981373 |
| Mfumv2_0272 | 354.441656 | -0.154198227 | 0.145155 | -1.0623 | 0.2881 | 0.525698 |
| Mfumv2_0273 | 43.52764215 | 0.314821883 | 0.281396 | 1.118785 | 0.263232 | 0.501264 |
| Mfumv2_0274 | 668.6611248 | -0.675627147 | 0.21506 | -3.14157 | 0.00168 | 0.012504 |
| Mfumv2_0275 | 99.34918725 | -1.066504587 | 0.64282 | -1.6591 | 0.097095 | 0.26733 |
| Mfumv2_0276 | 2746.095604 | -0.246837423 | 0.088939 | -2.77535 | 0.005514 | 0.032118 |
| Mfumv2_0277 | 194.1496625 | 0.023093643 | 0.156558 | 0.147509 | 0.882731 | 0.946455 |
| Mfumv2_0280 | 128.6477892 | 0.058548673 | 0.204388 | 0.286458 | 0.774527 | 0.890175 |
| Mfumv2_0281 | 13.56149705 | 0.464524992 | 0.493594 | 0.941107 | 0.34665 | NA |
| Mfumv2_0282 | 208.2855654 | 0.217919669 | 0.162798 | 1.338589 | 0.180704 | 0.401144 |
| Mfumv2_0283 | 21.66623365 | 0.104983056 | 0.393341 | 0.266901 | 0.789545 | NA |
| Mfumv2_0284 | 2.998998181 | 0.420542351 | 1.030376 | 0.408145 | 0.683168 | NA |
| Mfumv2_0285 | 9.345610499 | 0.659602591 | 0.618076 | 1.067188 | 0.285887 | NA |
| Mfumv2_0286 | 497.4814589 | -0.119529884 | 0.207854 | -0.57507 | 0.565246 | 0.776198 |
| Mfumv2_0287 | 15.90359792 | 0.317602146 | 0.466887 | 0.680254 | 0.496343 | NA |
| Mfumv2_0288 | 4.372296641 | 0.745058985 | 0.856958 | 0.869423 | 0.384616 | NA |
| Mfumv2_0289 | 6.096813947 | 0.756120772 | 0.759404 | 0.995677 | 0.319407 | NA |
| Mfumv2_0290 | 39.20696347 | -0.236257591 | 0.30745 | -0.76844 | 0.442225 | 0.672462 |
| Mfumv2_0291 | 1.399761789 | 0.412728827 | 1.492316 | 0.276569 | 0.782111 | NA |
| Mfumv2_0292 | 431.1406756 | 0.086135893 | 0.124281 | 0.693075 | 0.488263 | 0.716 |
| Mfumv2_0293 | 136.0460204 | -0.304691585 | 0.173627 | -1.75486 | 0.079283 | 0.22984 |
| Mfumv2_0294 | 161.7789679 | 0.239379952 | 0.162408 | 1.473945 | 0.140496 | 0.331845 |
| Mfumv2_0295 | 549.9969754 | 0.071073324 | 0.127961 | 0.555431 | 0.5786 | 0.788261 |
| Mfumv2_0296 | 106.7473206 | -0.242236295 | 0.198307 | -1.22152 | 0.221889 | 0.456269 |
| Mfumv2_0297 | 308.296412 | 0.247615376 | 0.13257 | 1.86781 | 0.061789 | 0.191971 |
| Mfumv2_0298 | 292.9499439 | 0.109896498 | 0.135 | 0.814046 | 0.415618 | 0.648274 |
| Mfumv2_0299 | 360.6202428 | 0.313376747 | 0.142655 | 2.196741 | 0.028039 | 0.108955 |
| Mfumv2_0300 | 70.70537668 | 0.185301269 | 0.243959 | 0.75956 | 0.447518 | 0.677564 |
| Mfumv2_0302 | 8.522656639 | 1.327146276 | 0.682877 | 1.943463 | 0.05196 | NA |
| Mfumv2_0303 | 76.93172584 | 0.67871892 | 0.244797 | 2.772578 | 0.005561 | 0.032292 |
| Mfumv2_0304 | 94.91881536 | 0.547261288 | 0.20092 | 2.723776 | 0.006454 | 0.036017 |
| Mfumv2_0306 | 354.1805161 | 0.131765467 | 0.144078 | 0.914544 | 0.360431 | 0.598929 |
| Mfumv2_0307 | 324.7256799 | -0.059357163 | 0.142746 | -0.41582 | 0.677539 | 0.842824 |
| Mfumv2_0308 | 84.3574133 | 0.061021883 | 0.21038 | 0.290055 | 0.771774 | 0.888026 |
| Mfumv2_0309 | 231.9282277 | 0.211436862 | 0.19449 | 1.087133 | 0.276978 | 0.517627 |
| Mfumv2_0310 | 241.88087 | -0.069089899 | 0.18677 | -0.36992 | 0.711443 | 0.858431 |
| Mfumv2_0311 | 152.5139349 | -0.054731833 | 0.176313 | -0.31043 | 0.756238 | 0.884498 |
| Mfumv2_0312 | 11.93095723 | 0.246496852 | 0.548653 | 0.449276 | 0.653232 | NA |
| Mfumv2_0313 | 1.585205398 | 1.498760894 | 1.612133 | 0.929676 | 0.352539 | NA |
| Mfumv2_0314 | 42.79979583 | 0.174012838 | 0.306997 | 0.566822 | 0.570835 | 0.780673 |
| Mfumv2_0315 | 126.5114976 | 0.213435261 | 0.183781 | 1.161354 | 0.245498 | 0.48434 |
| Mfumv2_0316 | 382.8247753 | 0.240094487 | 0.1482 | 1.620073 | 0.105217 | 0.28184 |
| Mfumv2_0317 | 734.4057752 | -0.270384397 | 0.150584 | -1.79557 | 0.072563 | 0.215979 |
| Mfumv2_0318 | 211.1024547 | 0.183142521 | 0.15302 | 1.196853 | 0.231364 | 0.468609 |
| Mfumv2_0319 | 819.7922781 | -0.147302438 | 0.103511 | -1.42307 | 0.154717 | 0.355229 |
| Mfumv2_0320 | 8.796421846 | 0.15351193 | 0.594112 | 0.258389 | 0.796107 | NA |
| Mfumv2_0321 | 13.06055454 | 0.484814452 | 0.534675 | 0.906746 | 0.364541 | NA |
| Mfumv2_0323 | 284.8789152 | 0.183681648 | 0.151976 | 1.208622 | 0.226808 | 0.463067 |
| Mfumv2_0324 | 245.2928389 | 0.321065061 | 0.13892 | 2.311155 | 0.020824 | 0.089203 |
| Mfumv2_0325 | 140.9151049 | 0.148578436 | 0.183372 | 0.810257 | 0.417792 | 0.650655 |
| Mfumv2_0326 | 319.2679598 | -0.169990895 | 0.130815 | -1.29947 | 0.193781 | 0.421298 |
| Mfumv2_0327 | 8.239447756 | 0.20016044 | 0.612375 | 0.326859 | 0.743774 | NA |

| | | | | | | |
|---|---|---|---|---|---|---|
| Mfumv2_0328 | 223.7389654 | -0.008708461 | 0.157776 | -0.0552 | 0.955983 | 0.982524 |
| Mfumv2_0329 | 1118.932863 | -0.079001665 | 0.135018 | -0.58512 | 0.558466 | 0.771231 |
| Mfumv2_0330 | 102.6133878 | 0.049807726 | 0.194445 | 0.256153 | 0.797832 | 0.901995 |
| Mfumv2_0331 | 948.5206136 | -0.046083114 | 0.105263 | -0.43779 | 0.661539 | 0.835454 |
| Mfumv2_0332 | 382.8974304 | -0.046090766 | 0.151204 | -0.30483 | 0.760499 | 0.884498 |
| Mfumv2_0333 | 3784.545907 | 0.185462534 | 0.453453 | 0.409001 | 0.682539 | 0.843309 |
| Mfumv2_0334 | 968.6851495 | 0.611291504 | 0.194156 | 3.148451 | 0.001641 | 0.012259 |
| Mfumv2_0335 | 114.6870812 | -0.26666433 | 0.232231 | -1.14827 | 0.250856 | 0.49072 |
| Mfumv2_0336 | 124.089351 | -0.295253944 | 0.186736 | -1.58113 | 0.113848 | 0.294651 |
| Mfumv2_0337 | 108.795859 | 0.210973582 | 0.21364 | 0.98752 | 0.323388 | 0.566462 |
| Mfumv2_0338 | 207.7767073 | -0.419838605 | 0.162627 | -2.5816 | 0.009834 | 0.050659 |
| Mfumv2_0339 | 160.1199382 | 0.022933063 | 0.170237 | 0.134713 | 0.892839 | 0.949055 |
| Mfumv2_0340 | 49.54278888 | 0.608663294 | 0.280649 | 2.168774 | 0.0301 | 0.114096 |
| Mfumv2_0341 | 444.9196744 | -0.404704931 | 0.165262 | -2.44887 | 0.01433 | 0.067309 |
| Mfumv2_0342 | 111.8105033 | -0.007079488 | 0.185672 | -0.03813 | 0.969585 | 0.98909 |
| Mfumv2_0343 | 362.5955662 | -0.352871839 | 0.136114 | -2.59247 | 0.009529 | 0.04955 |
| Mfumv2_0344 | 635.4691332 | -0.594780239 | 0.135109 | -4.40224 | 1.07E-05 | 0.000196 |
| Mfumv2_0345 | 129.6198982 | 0.181092819 | 0.177271 | 1.021561 | 0.306989 | 0.548772 |
| Mfumv2_0346 | 419.6338427 | 0.139818108 | 0.127441 | 1.097119 | 0.27259 | 0.511806 |
| Mfumv2_0347 | 329.8526792 | 0.339735264 | 0.149576 | 2.271315 | 0.023128 | 0.096467 |
| Mfumv2_0348 | 27.162525 | -0.064973464 | 0.347154 | -0.18716 | 0.851535 | 0.931265 |
| Mfumv2_0349 | 52.72299706 | -0.283151704 | 0.262274 | -1.0796 | 0.280318 | 0.520962 |
| Mfumv2_0350 | 46.06868675 | 0.347119475 | 0.293075 | 1.184407 | 0.236252 | 0.474072 |
| Mfumv2_0351 | 41.71826582 | 0.325774005 | 0.288849 | 1.127834 | 0.25939 | 0.498197 |
| Mfumv2_0352 | 5.952361315 | -0.243519373 | 0.720346 | -0.33806 | 0.735319 | NA |
| Mfumv2_0353 | 936.0582292 | 0.17726936 | 0.101385 | 1.748473 | 0.080382 | 0.23169 |
| Mfumv2_0354 | 147.543631 | 0.054639449 | 0.183827 | 0.297232 | 0.766289 | 0.887428 |
| Mfumv2_0355 | 81.01689803 | 0.528864307 | 0.237788 | 2.224104 | 0.026141 | 0.103325 |
| Mfumv2_0356 | 561.7908793 | 0.280445525 | 0.151413 | 1.852185 | 0.063999 | 0.1972 |
| Mfumv2_0357 | 676.5269912 | 0.045699322 | 0.108011 | 0.423098 | 0.672224 | 0.83934 |
| Mfumv2_0358 | 613.9860301 | -0.850890982 | 0.184718 | -4.60644 | 4.10E-06 | 8.07E-05 |
| Mfumv2_0359 | 2282.412793 | -0.207010381 | 0.112801 | -1.83518 | 0.066479 | 0.203281 |
| Mfumv2_0360 | 3990.263834 | -0.5717193 | 0.100401 | -5.69433 | 1.24E-08 | 4.08E-07 |
| Mfumv2_0361 | 140.3276597 | 0.249111626 | 0.186615 | 1.334893 | 0.181911 | 0.402228 |
| Mfumv2_0363 | 1272.083844 | -0.122552593 | 0.107817 | -1.13668 | 0.255673 | 0.493892 |
| Mfumv2_0364 | 1678.220328 | 0.154137388 | 0.09922 | 1.553499 | 0.120304 | 0.304013 |
| Mfumv2_0365 | 2138.356012 | -0.237227656 | 0.10276 | -2.30857 | 0.020968 | 0.089625 |
| Mfumv2_0366 | 1699.086532 | -0.067854163 | 0.104035 | -0.65222 | 0.514258 | 0.737285 |
| Mfumv2_0367 | 1343.857103 | -0.185535692 | 0.109735 | -1.69076 | 0.090883 | 0.253962 |
| Mfumv2_0368 | 1601.947913 | -0.279032752 | 0.100286 | -2.78238 | 0.005396 | 0.031792 |
| Mfumv2_0369 | 684.6342983 | -0.17462615 | 0.163355 | -1.069 | 0.28507 | 0.523113 |
| Mfumv2_0370 | 8.987633127 | 0.545662474 | 0.606274 | 0.900026 | 0.368106 | NA |
| Mfumv2_0371 | 675.622312 | -0.494680076 | 0.124646 | -3.96867 | 7.23E-05 | 0.000975 |
| Mfumv2_0372 | 1229.443503 | -0.416630471 | 0.10843 | -3.8424 | 0.000122 | 0.00153 |
| Mfumv2_0373 | 308.6611768 | -0.233498171 | 0.130761 | -1.78569 | 0.074149 | 0.220039 |
| Mfumv2_0374 | 176.8166463 | -0.168863421 | 0.180781 | -0.93408 | 0.350264 | 0.594325 |
| Mfumv2_0375 | 2.72765416 | 1.113117759 | 1.103596 | 1.008628 | 0.313153 | NA |
| Mfumv2_0376 | 5.584799994 | 0.754354766 | 0.749242 | 1.006824 | 0.314019 | NA |
| Mfumv2_0377 | 177.7904518 | 0.259106974 | 0.162092 | 1.59852 | 0.109927 | 0.289821 |
| Mfumv2_0378 | 936.6379662 | 0.01531272 | 0.1042 | 0.146955 | 0.883167 | 0.946455 |
| Mfumv2_0379 | 255.0725143 | -0.042282949 | 0.137537 | -0.30743 | 0.758517 | 0.884498 |
| Mfumv2_0380 | 102.6883718 | 0.108132935 | 0.19615 | 0.551275 | 0.581445 | 0.788739 |

| | | | | | | |
|---|---|---|---|---|---|---|
| Mfumv2_0381 | 127.6441015 | 0.119418248 | 0.185582 | 0.643479 | 0.519913 | 0.741836 |
| Mfumv2_0382 | 193.3967 | 0.092467808 | 0.153203 | 0.603565 | 0.546133 | 0.764552 |
| Mfumv2_0383 | 132.6945783 | 0.21710788 | 0.178738 | 1.214672 | 0.224491 | 0.458802 |
| Mfumv2_0384 | 205.1932905 | 0.066336698 | 0.16108 | 0.411824 | 0.680468 | 0.842824 |
| Mfumv2_0385 | 118.6327524 | 0.139297738 | 0.182888 | 0.761654 | 0.446266 | 0.676641 |
| Mfumv2_0386 | 167.047053 | 0.42554232 | 0.182129 | 2.336483 | 0.019466 | 0.085016 |
| Mfumv2_0387 | 1857.350864 | -0.068822434 | 0.117186 | -0.58729 | 0.557008 | 0.771212 |
| Mfumv2_0388 | 1624.557915 | -0.142673853 | 0.126012 | -1.13222 | 0.25754 | 0.496067 |
| Mfumv2_0389 | 452.632529 | -0.245344276 | 0.139253 | -1.76186 | 0.078093 | 0.227347 |
| Mfumv2_0390 | 657.5641289 | -0.303707953 | 0.120029 | -2.53029 | 0.011397 | 0.056394 |
| Mfumv2_0391 | 309.7147976 | -0.284814621 | 0.147148 | -1.93557 | 0.052921 | 0.170701 |
| Mfumv2_0392 | 1225.64822 | -0.401969207 | 0.130539 | -3.07931 | 0.002075 | 0.01494 |
| Mfumv2_0393 | 254.2547715 | -0.167462627 | 0.15792 | -1.06043 | 0.28895 | 0.526769 |
| Mfumv2_0394 | 118.443252 | 0.035700696 | 0.180564 | 0.197718 | 0.843266 | 0.925749 |
| Mfumv2_0395 | 333.9413283 | -0.144717739 | 0.135334 | -1.06933 | 0.284919 | 0.523113 |
| Mfumv2_0396 | 41.60495623 | -0.158675687 | 0.358883 | -0.44214 | 0.658389 | 0.834776 |
| Mfumv2_0397 | 5236.152689 | -0.396276716 | 0.098173 | -4.03651 | 5.43E-05 | 0.000777 |
| Mfumv2_0399 | 224.1878364 | -0.092665376 | 0.151224 | -0.61277 | 0.540029 | 0.76028 |
| Mfumv2_0400 | 385.9949437 | -0.403626889 | 0.135202 | -2.98536 | 0.002832 | 0.019095 |
| Mfumv2_0401 | 214.0348659 | 0.146602646 | 0.154394 | 0.949534 | 0.342349 | 0.585344 |
| Mfumv2_0402 | 12.82108077 | 0.419741164 | 0.508257 | 0.825844 | 0.408893 | NA |
| Mfumv2_0403 | 150.9731196 | 0.143297073 | 0.175405 | 0.816948 | 0.413958 | 0.647422 |
| Mfumv2_0404 | 760.6542112 | -0.241104249 | 0.119662 | -2.01488 | 0.043917 | 0.14929 |
| Mfumv2_0405 | 1210.372775 | -0.038053346 | 0.097651 | -0.38969 | 0.696767 | 0.850369 |
| Mfumv2_0406 | 177.5735351 | -0.255939352 | 0.170428 | -1.50175 | 0.133162 | 0.321018 |
| Mfumv2_0407 | 683.5586068 | -0.366281947 | 0.121132 | -3.02383 | 0.002496 | 0.017291 |
| Mfumv2_0408 | 1324.493743 | -0.261992332 | 0.105568 | -2.48174 | 0.013074 | 0.062689 |
| Mfumv2_0409 | 54.86903607 | -0.143380648 | 0.264657 | -0.54176 | 0.587983 | 0.792791 |
| Mfumv2_0410 | 55.24076416 | 0.045916767 | 0.279558 | 0.164248 | 0.869536 | 0.939698 |
| Mfumv2_0411 | 113.1684007 | 0.057571391 | 0.209877 | 0.27431 | 0.783846 | 0.89556 |
| Mfumv2_0412 | 255.2348237 | 0.004199698 | 0.136076 | 0.030863 | 0.975379 | 0.990165 |
| Mfumv2_0413 | 403.727307 | 0.183819176 | 0.12202 | 1.506474 | 0.131946 | 0.320143 |
| Mfumv2_0414 | 672.0064123 | -0.216836328 | 0.139466 | -1.55476 | 0.120003 | 0.304013 |
| Mfumv2_0415 | 862.3033364 | -0.364054984 | 0.136331 | -2.67038 | 0.007576 | 0.041475 |
| Mfumv2_0416 | 436.7886732 | -0.18986638 | 0.128725 | -1.47498 | 0.140218 | 0.331845 |
| Mfumv2_0417 | 89.54255235 | -0.386283209 | 0.214397 | -1.80172 | 0.07159 | 0.215306 |
| Mfumv2_0418 | 11.20515653 | -0.570203807 | 0.534754 | -1.06629 | 0.286291 | NA |
| Mfumv2_0419 | 19.56674841 | -0.383872028 | 0.406583 | -0.94414 | 0.345097 | NA |
| Mfumv2_0420 | 86.58477722 | -0.318005391 | 0.212512 | -1.49641 | 0.134547 | 0.323319 |
| Mfumv2_0421 | 1444.836265 | -0.351233994 | 0.121491 | -2.89104 | 0.00384 | 0.024489 |
| Mfumv2_0422 | 52.9084082 | -0.544120748 | 0.26163 | -2.07974 | 0.03755 | 0.13562 |
| Mfumv2_0423 | 115.1553509 | -0.772445113 | 0.190251 | -4.06014 | 4.90E-05 | 0.000709 |
| Mfumv2_0424 | 567.8523803 | -0.926071539 | 0.128035 | -7.23297 | 4.73E-13 | 2.88E-11 |
| Mfumv2_0425 | 512.9955227 | -0.917369877 | 0.132717 | -6.91223 | 4.77E-12 | 2.59E-10 |
| Mfumv2_0426 | 380.6531205 | -0.873698124 | 0.142127 | -6.1473 | 7.88E-10 | 3.37E-08 |
| Mfumv2_0427 | 318.9992162 | -0.876286208 | 0.145009 | -6.04296 | 1.51E-09 | 6.08E-08 |
| Mfumv2_0428 | 862.037707 | -0.642620841 | 0.149067 | -4.31096 | 1.63E-05 | 0.000279 |
| Mfumv2_0429 | 29.11065868 | -0.245893425 | 0.341898 | -0.7192 | 0.472017 | 0.699323 |
| Mfumv2_0430 | 29.09613421 | -0.015229087 | 0.345971 | -0.04402 | 0.96489 | 0.988508 |
| Mfumv2_0431 | 233.0891578 | 0.191613404 | 0.172067 | 1.113594 | 0.265453 | 0.50406 |
| Mfumv2_0432 | 177.5537361 | -0.375638398 | 0.177034 | -2.12184 | 0.033851 | 0.124328 |
| Mfumv2_0433 | 247.4050442 | 0.010195965 | 0.139482 | 0.073099 | 0.941728 | 0.977742 |

| | | | | | | |
|---|---|---|---|---|---|---|
| Mfumv2_0434 | 39.14196261 | 0.100180301 | 0.29129 | 0.343919 | 0.730907 | 0.871449 |
| Mfumv2_0435 | 106.5302078 | 0.126084412 | 0.210281 | 0.599601 | 0.548772 | 0.76593 |
| Mfumv2_0436 | 630.4758059 | -0.054114834 | 0.115613 | -0.46807 | 0.639737 | 0.823866 |
| Mfumv2_0437 | 147.2061954 | -0.131124291 | 0.187355 | -0.69987 | 0.484009 | 0.711319 |
| Mfumv2_0438 | 501.1951209 | -0.418854941 | 0.130542 | -3.20858 | 0.001334 | 0.010428 |
| Mfumv2_0439 | 488.1276311 | -0.325400655 | 0.130807 | -2.48764 | 0.012859 | 0.062102 |
| Mfumv2_0440 | 1225.959841 | -0.397328229 | 0.122813 | -3.23523 | 0.001215 | 0.009665 |
| Mfumv2_0441 | 8.742500632 | 0.128134145 | 0.587601 | 0.218063 | 0.82738 | NA |
| Mfumv2_0442 | 238.4477995 | 0.0517675 | 0.140424 | 0.36865 | 0.712388 | 0.859057 |
| Mfumv2_0443 | 928.6277917 | -0.603793972 | 0.146836 | -4.11204 | 3.92E-05 | 0.000611 |
| Mfumv2_0444 | 9.299823413 | 0.331843306 | 0.588174 | 0.564192 | 0.572623 | NA |
| Mfumv2_0445 | 1.359855302 | 2.391763981 | 1.679539 | 1.424059 | 0.154429 | NA |
| Mfumv2_0447 | 107.4838567 | 0.292910934 | 0.203279 | 1.440929 | 0.149605 | 0.345864 |
| Mfumv2_0448 | 6442.133457 | -0.446470285 | 0.128912 | -3.46336 | 0.000533 | 0.005079 |
| Mfumv2_0449 | 822.9167775 | 0.318523476 | 0.113574 | 2.804549 | 0.005039 | 0.030217 |
| Mfumv2_0450 | 984.0396336 | 0.060744513 | 0.098453 | 0.616992 | 0.53724 | 0.757946 |
| Mfumv2_0451 | 143.1061232 | 0.36746401 | 0.179376 | 2.048565 | 0.040505 | 0.140785 |
| Mfumv2_0452 | 137.2102775 | 0.239622999 | 0.180982 | 1.324017 | 0.185497 | 0.40507 |
| Mfumv2_0453 | 319.678253 | 0.108198323 | 0.131018 | 0.825825 | 0.408903 | 0.642476 |
| Mfumv2_0454 | 202.6860025 | -0.173361449 | 0.239711 | -0.72321 | 0.46955 | 0.697712 |
| Mfumv2_0455 | 33.9442449 | 0.538017456 | 0.341116 | 1.577227 | 0.114743 | 0.295159 |
| Mfumv2_0456 | 168.8401516 | 0.294112611 | 0.221393 | 1.328465 | 0.184025 | 0.40405 |
| Mfumv2_0457 | 662.3095536 | -0.089244017 | 0.133847 | -0.66676 | 0.504926 | 0.730833 |
| Mfumv2_0458 | 379.0049821 | -0.084033351 | 0.134725 | -0.62374 | 0.532799 | 0.754861 |
| Mfumv2_0459 | 161.9536206 | -0.20549874 | 0.159951 | -1.28476 | 0.198875 | 0.428432 |
| Mfumv2_0460 | 488.2864467 | 0.158935427 | 0.119546 | 1.329497 | 0.183684 | 0.403743 |
| Mfumv2_0461 | 5906.336217 | -0.243238869 | 0.11954 | -2.03479 | 0.041872 | 0.144788 |
| Mfumv2_0462 | 33.47847895 | 0.478100535 | 0.406845 | 1.175143 | 0.239938 | 0.479636 |
| Mfumv2_0463 | 110.605093 | -0.101541858 | 0.186492 | -0.54448 | 0.586108 | 0.791857 |
| Mfumv2_0464 | 80.92703098 | -0.288451111 | 0.267499 | -1.07833 | 0.280888 | 0.521056 |
| Mfumv2_0465 | 0.479518162 | -1.365869326 | 2.681855 | -0.5093 | 0.610542 | NA |
| Mfumv2_0466 | 7.490575826 | 0.762156218 | 0.694735 | 1.097046 | 0.272621 | NA |
| Mfumv2_0467 | 12.02969761 | 0.69534428 | 0.566838 | 1.226707 | 0.219933 | NA |
| Mfumv2_0468 | 0.11657447 | 0.283845181 | 4.080473 | 0.069562 | 0.944542 | NA |
| Mfumv2_0469 | 12.33329345 | -0.511730721 | 0.495297 | -1.03318 | 0.30152 | NA |
| Mfumv2_0470 | 36.23930858 | -0.252069594 | 0.300833 | -0.8379 | 0.402084 | 0.635053 |
| Mfumv2_0472 | 64.54643209 | 0.009323596 | 0.242457 | 0.038455 | 0.969325 | 0.98909 |
| Mfumv2_0473 | 308.8135932 | -0.001113956 | 0.138366 | -0.00805 | 0.993576 | 0.996553 |
| Mfumv2_0474 | 16.16468837 | 0.315680868 | 0.462567 | 0.682454 | 0.494952 | NA |
| Mfumv2_0475 | 946.3731317 | 0.03375302 | 0.16594 | 0.203405 | 0.838818 | 0.923896 |
| Mfumv2_0476 | 312.0149491 | 0.068668182 | 0.13195 | 0.520409 | 0.602778 | 0.802506 |
| Mfumv2_0477 | 399.0297993 | 0.108903395 | 0.124463 | 0.874986 | 0.381581 | 0.61574 |
| Mfumv2_0478 | 134.4119653 | -0.38673363 | 0.171487 | -2.25518 | 0.024122 | 0.098475 |
| Mfumv2_0479 | 294.0723924 | 0.289405447 | 0.141163 | 2.050149 | 0.04035 | 0.140533 |
| Mfumv2_0480 | 308.4400814 | -0.441256192 | 0.130011 | -3.394 | 0.000689 | 0.006205 |
| Mfumv2_0481 | 550.2023919 | -0.003900929 | 0.130882 | -0.0298 | 0.976223 | 0.990521 |
| Mfumv2_0482 | 91.08913133 | 0.243340024 | 0.227154 | 1.071257 | 0.284054 | 0.523113 |
| Mfumv2_0483 | 271.0472157 | 0.080796722 | 0.136412 | 0.592298 | 0.553651 | 0.767622 |
| Mfumv2_0484 | 283.1701153 | -0.145544391 | 0.181518 | -0.80182 | 0.422657 | 0.65569 |
| Mfumv2_0485 | 181.3394049 | 0.512374928 | 0.158531 | 3.23201 | 0.001229 | 0.009723 |
| Mfumv2_0487 | 2.741494826 | 1.216226604 | 1.137572 | 1.069142 | 0.285005 | NA |
| Mfumv2_0488 | 437.898606 | -0.30440087 | 0.16935 | -1.79746 | 0.072262 | 0.215979 |

| | | | | | | |
|---|---|---|---|---|---|---|
| Mfumv2_0489 | 206.4720537 | -0.42118691 | 0.248095 | -1.69768 | 0.089568 | 0.251667 |
| Mfumv2_0490 | 23.17234121 | -0.107486251 | 0.451028 | -0.23831 | 0.811637 | NA |
| Mfumv2_0491 | 2.005803101 | 0.354297613 | 1.337621 | 0.264872 | 0.791108 | NA |
| Mfumv2_0492 | 199.9956977 | -7.97E-05 | 0.179837 | -0.00044 | 0.999646 | 0.999646 |
| Mfumv2_0493 | 441.0531668 | 0.111038872 | 0.126699 | 0.876397 | 0.380814 | 0.615513 |
| Mfumv2_0495 | 204.5033452 | -0.35622943 | 0.183158 | -1.94493 | 0.051783 | 0.169157 |
| Mfumv2_0496 | 18.30479206 | 0.362850952 | 0.426728 | 0.850311 | 0.395152 | NA |
| Mfumv2_0497 | 29.06852233 | 0.093864545 | 0.349954 | 0.26822 | 0.78853 | 0.897539 |
| Mfumv2_0498 | 349.8141018 | 0.090668175 | 0.165246 | 0.548688 | 0.58322 | 0.79008 |
| Mfumv2_0499 | 1293.143695 | -0.518092157 | 0.114756 | -4.51473 | 6.34E-06 | 0.000119 |
| Mfumv2_0500 | 660.4702772 | -0.18359179 | 0.119508 | -1.53623 | 0.124482 | 0.30951 |
| Mfumv2_0501 | 718.954635 | -0.191884665 | 0.140577 | -1.36498 | 0.172259 | 0.385806 |
| Mfumv2_0502 | 1250.191874 | -0.237685294 | 0.140625 | -1.6902 | 0.090989 | 0.253962 |
| Mfumv2_0503 | 206.8266536 | 0.155333576 | 0.146773 | 1.058328 | 0.289906 | 0.527769 |
| Mfumv2_0505 | 124.2497501 | 0.266096366 | 0.179132 | 1.485481 | 0.137417 | 0.327557 |
| Mfumv2_0506 | 35.090619 | 0.908405134 | 0.325455 | 2.791185 | 0.005252 | 0.031214 |
| Mfumv2_0507 | 49.83272041 | -0.384107534 | 0.282795 | -1.35825 | 0.174384 | 0.389263 |
| Mfumv2_0508 | 874.0720951 | -0.008827888 | 0.105465 | -0.0837 | 0.933292 | 0.97503 |
| Mfumv2_0509 | 261.0216787 | 0.08902293 | 0.166895 | 0.533407 | 0.593752 | 0.795297 |
| Mfumv2_0510 | 192.266985 | -0.281989307 | 0.14989 | -1.88131 | 0.05993 | 0.187829 |
| Mfumv2_0511 | 1.748561738 | -0.334876461 | 1.306193 | -0.25638 | 0.797661 | NA |
| Mfumv2_0512 | 49.2331152 | 0.138610382 | 0.278213 | 0.498216 | 0.618332 | 0.813809 |
| Mfumv2_0513 | 338.3968833 | -0.044632684 | 0.149327 | -0.29889 | 0.765023 | 0.887034 |
| Mfumv2_0514 | 132.1618324 | 0.215536761 | 0.182888 | 1.178517 | 0.238591 | 0.477419 |
| Mfumv2_0515 | 69.45778983 | 0.213906484 | 0.23328 | 0.916952 | 0.359168 | 0.598398 |
| Mfumv2_0516 | 388.8210322 | 0.074152534 | 0.121431 | 0.610654 | 0.541429 | 0.76074 |
| Mfumv2_0517 | 203.0502202 | -0.153687953 | 0.16157 | -0.95121 | 0.341496 | 0.584519 |
| Mfumv2_0518 | 1838.530063 | 0.033116156 | 0.106773 | 0.310153 | 0.756444 | 0.884498 |
| Mfumv2_0519 | 2112.77392 | -0.295049991 | 0.12042 | -2.45018 | 0.014279 | 0.067309 |
| Mfumv2_0520 | 151.3555049 | 0.125097045 | 0.189557 | 0.659943 | 0.50929 | 0.733977 |
| Mfumv2_0521 | 442.1542043 | -0.146058621 | 0.124719 | -1.1711 | 0.241559 | 0.480964 |
| Mfumv2_0522 | 95.60939592 | 0.039471184 | 0.205889 | 0.191711 | 0.847969 | 0.928376 |
| Mfumv2_0523 | 90.81206314 | -0.071081783 | 0.212466 | -0.33456 | 0.737959 | 0.875183 |
| Mfumv2_0524 | 57.63598873 | -0.118562862 | 0.287674 | -0.41214 | 0.680234 | 0.842824 |
| Mfumv2_0525 | 1733.210426 | -3.53985142 | 0.481521 | -7.3514 | 1.96E-13 | 1.23E-11 |
| Mfumv2_0526 | 1469.745139 | -4.220755728 | 0.199425 | -21.1646 | 2.03E-99 | 1.02E-96 |
| Mfumv2_0527 | 535.1840594 | -4.877044614 | 0.224797 | -21.6953 | 2.27E-104 | 1.52E-101 |
| Mfumv2_0528 | 72.54085426 | -3.102304975 | 0.602076 | -5.15268 | 2.57E-07 | 6.22E-06 |
| Mfumv2_0530 | 81.23822166 | 0.171318565 | 0.210062 | 0.815564 | 0.41475 | 0.647422 |
| Mfumv2_0531 | 86.68309963 | 0.204353474 | 0.204848 | 0.997587 | 0.31848 | 0.561743 |
| Mfumv2_0532 | 99.05114264 | -0.116823848 | 0.19347 | -0.60384 | 0.545953 | 0.764552 |
| Mfumv2_0533 | 103.3607882 | 0.270470768 | 0.196442 | 1.376848 | 0.168559 | 0.380062 |
| Mfumv2_0534 | 72.2534298 | 0.101050046 | 0.222864 | 0.453417 | 0.650249 | 0.829769 |
| Mfumv2_0535 | 16.06640042 | -0.631040603 | 0.437472 | -1.44247 | 0.14917 | NA |
| Mfumv2_0536 | 48.97433776 | -0.282932344 | 0.265128 | -1.06715 | 0.285903 | 0.523113 |
| Mfumv2_0537 | 4.066067801 | 0.882612385 | 0.965038 | 0.914589 | 0.360408 | NA |
| Mfumv2_0538 | 975.134422 | 0.00864438 | 0.118242 | 0.073108 | 0.94172 | 0.977742 |
| Mfumv2_0539 | 164.7153233 | 0.147768405 | 0.165011 | 0.895506 | 0.370516 | 0.608143 |
| Mfumv2_0540 | 81.34591571 | 0.245230635 | 0.226512 | 1.08264 | 0.278968 | 0.518933 |
| Mfumv2_0541 | 27.1419085 | 0.319862992 | 0.345644 | 0.925411 | 0.354752 | 0.59836 |
| Mfumv2_0543 | 7.739269346 | -0.237672117 | 0.617968 | -0.3846 | 0.700532 | NA |
| Mfumv2_0544 | 11.70507012 | 0.303656567 | 0.516258 | 0.588188 | 0.556406 | NA |

| | | | | | | |
|---|---|---|---|---|---|---|
| Mfumv2_0545 | 24.71819639 | 0.150221485 | 0.36391 | 0.412799 | 0.679754 | 0.842824 |
| Mfumv2_0546 | 74.3155074 | 0.191999862 | 0.240351 | 0.798832 | 0.424388 | 0.656853 |
| Mfumv2_0547 | 15.979298 | 0.435664051 | 0.467053 | 0.932794 | 0.350926 | NA |
| Mfumv2_0548 | 139.6866324 | 0.114002734 | 0.170631 | 0.668126 | 0.504053 | 0.730096 |
| Mfumv2_0550 | 42.56960149 | 0.018417797 | 0.299935 | 0.061406 | 0.951036 | 0.981373 |
| Mfumv2_0551 | 272.2663322 | -0.380853344 | 0.139083 | -2.73831 | 0.006176 | 0.034863 |
| Mfumv2_0552 | 61.50573361 | -0.35176889 | 0.349998 | -1.00506 | 0.314868 | 0.557331 |
| Mfumv2_0553 | 13.46081719 | 0.648140835 | 0.492029 | 1.317282 | 0.187744 | NA |
| Mfumv2_0555 | 10.6385405 | 0.314934245 | 0.567664 | 0.55479 | 0.579038 | NA |
| Mfumv2_0556 | 36.34606699 | 0.351027521 | 0.315546 | 1.112446 | 0.265947 | 0.50452 |
| Mfumv2_0557 | 7.518153396 | 0.175743744 | 0.667734 | 0.263194 | 0.792401 | NA |
| Mfumv2_0558 | 42.90751845 | -0.051955051 | 0.30612 | -0.16972 | 0.865229 | 0.938198 |
| Mfumv2_0559 | 17.76683394 | 0.58835958 | 0.437291 | 1.345465 | 0.178475 | NA |
| Mfumv2_0560 | 27.79715101 | 0.360387315 | 0.349857 | 1.0301 | 0.302963 | 0.544232 |
| Mfumv2_0561 | 46.42022717 | 0.368630181 | 0.288494 | 1.277774 | 0.201329 | 0.432589 |
| Mfumv2_0562 | 95.10266062 | 0.040476748 | 0.205632 | 0.196841 | 0.843952 | 0.925996 |
| Mfumv2_0563 | 76.85352248 | -0.279613183 | 0.222879 | -1.25455 | 0.209641 | 0.440553 |
| Mfumv2_0564 | 2283.903829 | -0.610674237 | 0.119346 | -5.11682 | 3.11E-07 | 7.26E-06 |
| Mfumv2_0565 | 739.4069641 | 0.075972154 | 0.121227 | 0.626692 | 0.530861 | 0.752646 |
| Mfumv2_0566 | 316.6985131 | -0.15812032 | 0.151222 | -1.04562 | 0.295738 | 0.534877 |
| Mfumv2_0567 | 261.9128624 | -0.767727087 | 0.190327 | -4.03373 | 5.49E-05 | 0.000777 |
| Mfumv2_0568 | 115.2985402 | -0.238213607 | 0.192871 | -1.23509 | 0.216796 | 0.44809 |
| Mfumv2_0569 | 214.7216633 | 0.100627695 | 0.162716 | 0.618426 | 0.536294 | 0.75729 |
| Mfumv2_0570 | 44.24644703 | 0.216336383 | 0.320264 | 0.675494 | 0.499362 | 0.727497 |
| Mfumv2_0571 | 4.780239957 | -0.634869581 | 0.852642 | -0.74459 | 0.456519 | NA |
| Mfumv2_0572 | 3.904032719 | 0.300304082 | 0.90374 | 0.33229 | 0.73967 | NA |
| Mfumv2_0573 | 171.4002476 | -1.115836182 | 0.278868 | -4.00131 | 6.30E-05 | 0.00087 |
| Mfumv2_0574 | 226.091633 | -0.167197065 | 0.187522 | -0.89161 | 0.372601 | 0.610542 |
| Mfumv2_0575 | 216.1681345 | -0.45086009 | 0.166488 | -2.70807 | 0.006768 | 0.037662 |
| Mfumv2_0577 | 359.3758204 | -0.032575839 | 0.137539 | -0.23685 | 0.812775 | 0.907755 |
| Mfumv2_0578 | 891.597098 | -0.772463607 | 0.160566 | -4.81087 | 1.50E-06 | 3.14E-05 |
| Mfumv2_0579 | 194.7479349 | -0.003568673 | 0.154032 | -0.02317 | 0.981516 | 0.992996 |
| Mfumv2_0580 | 68.29848379 | -0.137807148 | 0.230843 | -0.59697 | 0.550524 | 0.76593 |
| Mfumv2_0581 | 153.5213123 | -0.195162214 | 0.167098 | -1.16795 | 0.242828 | 0.482533 |
| Mfumv2_0582 | 703.5575308 | 0.344914262 | 0.113194 | 3.047117 | 0.00231 | 0.01632 |
| Mfumv2_0583 | 434.3583368 | 0.379635269 | 0.152569 | 2.488284 | 0.012836 | 0.062102 |
| Mfumv2_0584 | 1694.544063 | -0.421469863 | 0.11537 | -3.6532 | 0.000259 | 0.002812 |
| Mfumv2_0585 | 780.57163 | -0.079252461 | 0.125986 | -0.62906 | 0.529312 | 0.75151 |
| Mfumv2_0586 | 13.01781918 | -0.105479442 | 0.608104 | -0.17346 | 0.862293 | NA |
| Mfumv2_0587 | 3.477363471 | -0.738748727 | 0.948664 | -0.77873 | 0.436142 | NA |
| Mfumv2_0588 | 62.43345939 | 0.023771522 | 0.239765 | 0.099145 | 0.921023 | 0.9689 |
| Mfumv2_0589 | 255.7911179 | -0.530074211 | 0.149459 | -3.54662 | 0.00039 | 0.004 |
| Mfumv2_0590 | 46.66051702 | 0.244270942 | 0.271699 | 0.89905 | 0.368626 | 0.607278 |
| Mfumv2_0591 | 230.2822957 | 0.091450136 | 0.155056 | 0.589786 | 0.555334 | 0.769425 |
| Mfumv2_0592 | 942.0223167 | -0.001818648 | 0.126796 | -0.01434 | 0.988556 | 0.994355 |
| Mfumv2_0593 | 17.23082125 | 0.653321355 | 0.451034 | 1.448497 | 0.147478 | NA |
| Mfumv2_0594 | 81.90450309 | 0.456124065 | 0.310394 | 1.469502 | 0.141697 | 0.333336 |
| Mfumv2_0595 | 557.5206256 | -0.291191219 | 0.11772 | -2.4736 | 0.013376 | 0.06383 |
| Mfumv2_0596 | 175.2448545 | 0.52990409 | 0.188128 | 2.816727 | 0.004852 | 0.02927 |
| Mfumv2_0597 | 2261.568843 | -0.409301805 | 0.119344 | -3.42958 | 0.000605 | 0.005622 |
| Mfumv2_0598 | 4.436341991 | 0.094670096 | 0.840008 | 0.112701 | 0.910267 | NA |
| Mfumv2_0599 | 1.685934079 | -0.705997671 | 1.359701 | -0.51923 | 0.6036 | NA |

| | | | | | | |
|---|---|---|---|---|---|---|
| Mfumv2_0600 | 9.445606663 | 0.049921925 | 0.594888 | 0.083918 | 0.933121 | NA |
| Mfumv2_0601 | 144.9544828 | 0.111279428 | 0.173139 | 0.642717 | 0.520407 | 0.742015 |
| Mfumv2_0602 | 294.128278 | 0.045808002 | 0.135363 | 0.338408 | 0.735056 | 0.872254 |
| Mfumv2_0603 | 182.3847379 | -0.32428847 | 0.180108 | -1.80052 | 0.071778 | 0.215548 |
| Mfumv2_0604 | 124.4963376 | -0.081366553 | 0.198421 | -0.41007 | 0.681754 | 0.843309 |
| Mfumv2_0605 | 401.6792306 | -0.094063217 | 0.142139 | -0.66177 | 0.508118 | 0.733339 |
| Mfumv2_0606 | 42.3828377 | 0.546793882 | 0.329084 | 1.661564 | 0.0966 | 0.266579 |
| Mfumv2_0607 | 2440.434755 | -0.17993763 | 0.186696 | -0.9638 | 0.335146 | 0.577948 |
| Mfumv2_0608 | 1990.574838 | 0.115413082 | 0.095911 | 1.203334 | 0.228847 | 0.465042 |
| Mfumv2_0609 | 185.3925566 | -0.017666035 | 0.190821 | -0.09258 | 0.926238 | 0.972211 |
| Mfumv2_0610 | 345.8659887 | 0.214776816 | 0.142764 | 1.50442 | 0.132473 | 0.320289 |
| Mfumv2_0611 | 262.702005 | 0.384489813 | 0.172887 | 2.22394 | 0.026152 | 0.103325 |
| Mfumv2_0612 | 43.74734479 | -0.025254953 | 0.292432 | -0.08636 | 0.931179 | 0.973858 |
| Mfumv2_0613 | 536.4884325 | -0.276623857 | 0.125968 | -2.19599 | 0.028093 | 0.108955 |
| Mfumv2_0614 | 150.1506324 | -0.083200038 | 0.221883 | -0.37497 | 0.70768 | 0.855433 |
| Mfumv2_0615 | 664.9690883 | -0.113452969 | 0.120236 | -0.94358 | 0.345382 | 0.588836 |
| Mfumv2_0616 | 359.9770199 | 0.030202837 | 0.126498 | 0.23876 | 0.811291 | 0.907508 |
| Mfumv2_0617 | 101.4703955 | 0.059998638 | 0.199952 | 0.300065 | 0.764127 | 0.887034 |
| Mfumv2_0618 | 133.8851463 | 0.049584063 | 0.180991 | 0.273959 | 0.784116 | 0.89556 |
| Mfumv2_0619 | 311.8085532 | 0.070109832 | 0.13591 | 0.515853 | 0.605957 | 0.803543 |
| Mfumv2_0620 | 1014.962601 | -0.358200577 | 0.108283 | -3.30801 | 0.00094 | 0.007898 |
| Mfumv2_0621 | 134.9120998 | 0.394713219 | 0.204395 | 1.931127 | 0.053467 | 0.172141 |
| Mfumv2_0622 | 148.1762216 | 0.044590487 | 0.190652 | 0.233885 | 0.815074 | 0.908704 |
| Mfumv2_0623 | 396.0682408 | -0.162272687 | 0.138198 | -1.1742 | 0.240314 | 0.47971 |
| Mfumv2_0624 | 936.0078135 | -0.075900101 | 0.107724 | -0.70458 | 0.481071 | 0.708557 |
| Mfumv2_0625 | 465.73769 | -0.19244534 | 0.127566 | -1.50859 | 0.131403 | 0.319212 |
| Mfumv2_0626 | 414.6436002 | -0.571002685 | 0.125876 | -4.53624 | 5.73E-06 | 0.000109 |
| Mfumv2_0627 | 352.3189561 | -0.294937212 | 0.128526 | -2.29477 | 0.021746 | 0.092365 |
| Mfumv2_0628 | 324.6578142 | -0.167639079 | 0.128014 | -1.30954 | 0.190351 | 0.414768 |
| Mfumv2_0629 | 499.7826456 | 0.004460577 | 0.122454 | 0.036427 | 0.970942 | 0.98916 |
| Mfumv2_0630 | 0 | NA | NA | NA | NA | NA |
| Mfumv2_0631 | 166.2817323 | 0.343016788 | 0.166231 | 2.063495 | 0.039066 | 0.137931 |
| Mfumv2_0632 | 989.199523 | -0.35023152 | 0.137782 | -2.54192 | 0.011024 | 0.054822 |
| Mfumv2_0633 | 0.532309273 | -1.508460162 | 2.414938 | -0.62464 | 0.532209 | NA |
| Mfumv2_0634 | 149.8921814 | -0.011283419 | 0.164602 | -0.06855 | 0.945348 | 0.979433 |
| Mfumv2_0635 | 72.024156 | 0.227769032 | 0.233865 | 0.973932 | 0.33009 | 0.57267 |
| Mfumv2_0636 | 31.87747162 | 0.949190215 | 0.336047 | 2.824578 | 0.004734 | 0.028735 |
| Mfumv2_0637 | 557.4835238 | 0.008260544 | 0.132032 | 0.062565 | 0.950113 | 0.981373 |
| Mfumv2_0638 | 9.009598935 | 0.648787018 | 0.603525 | 1.074995 | 0.282377 | NA |
| Mfumv2_0639 | 567.1851713 | -0.085351214 | 0.125328 | -0.68102 | 0.495857 | 0.72344 |
| Mfumv2_0640 | 89.88689482 | -0.191818845 | 0.24663 | -0.77776 | 0.436711 | 0.667188 |
| Mfumv2_0641 | 361.0127951 | -0.209016818 | 0.165996 | -1.25917 | 0.20797 | 0.437957 |
| Mfumv2_0642 | 303.1066925 | 0.231964111 | 0.141357 | 1.640985 | 0.100801 | 0.275147 |
| Mfumv2_0643 | 185.7967123 | -0.022650888 | 0.164818 | -0.13743 | 0.890691 | 0.948277 |
| Mfumv2_0644 | 48.60906061 | -0.102679527 | 0.270536 | -0.37954 | 0.704286 | 0.854415 |
| Mfumv2_0645 | 4.400184163 | 1.299415701 | 0.894743 | 1.452278 | 0.146424 | NA |
| Mfumv2_0646 | 213.6964161 | 0.083274853 | 0.147941 | 0.562891 | 0.573509 | 0.783263 |
| Mfumv2_0647 | 176.8727031 | 0.466377929 | 0.166483 | 2.801348 | 0.005089 | 0.030368 |
| Mfumv2_0648 | 204.6912781 | 0.390418536 | 0.147969 | 2.638524 | 0.008327 | 0.044609 |
| Mfumv2_0649 | 103.8524377 | 0.032192883 | 0.190747 | 0.168772 | 0.865976 | 0.938374 |
| Mfumv2_0650 | 328.9802535 | 0.10008672 | 0.128321 | 0.779969 | 0.435409 | 0.666666 |
| Mfumv2_0651 | 135.4720983 | 0.284316495 | 0.171757 | 1.655343 | 0.097855 | 0.267968 |

| | | | | | | |
|---|---|---|---|---|---|---|
| Mfumv2_0653 | 129.842501 | 0.22610502 | 0.180531 | 1.252448 | 0.210407 | 0.441239 |
| Mfumv2_0654 | 156.0493333 | -0.04141958 | 0.178832 | -0.23161 | 0.81684 | 0.909463 |
| Mfumv2_0655 | 102.4966249 | -0.093185546 | 0.207903 | -0.44822 | 0.653996 | 0.832096 |
| Mfumv2_0656 | 101.8856826 | 0.191688986 | 0.201372 | 0.951916 | 0.341139 | 0.584519 |
| Mfumv2_0657 | 98.44060213 | 0.354841655 | 0.223138 | 1.590235 | 0.111782 | 0.291291 |
| Mfumv2_0658 | 1515.125128 | -0.973105441 | 0.124451 | -7.8192 | 5.32E-15 | 3.56E-13 |
| Mfumv2_0659 | 506.4734913 | -0.913908064 | 0.173328 | -5.2727 | 1.34E-07 | 3.51E-06 |
| Mfumv2_0660 | 237.2083045 | -0.866269122 | 0.202006 | -4.28833 | 1.80E-05 | 0.000299 |
| Mfumv2_0661 | 429.9154886 | 0.041955469 | 0.131722 | 0.318514 | 0.750095 | 0.880222 |
| Mfumv2_0662 | 307.7859649 | -0.041217116 | 0.140072 | -0.29426 | 0.768561 | 0.887638 |
| Mfumv2_0663 | 741.1613284 | 0.174499854 | 0.113196 | 1.541568 | 0.123179 | 0.307411 |
| Mfumv2_0664 | 124.0586885 | 0.012248954 | 0.189806 | 0.064534 | 0.948545 | 0.980261 |
| Mfumv2_0665 | 2454.746123 | -0.117295798 | 0.1162 | -1.00943 | 0.312766 | 0.555077 |
| Mfumv2_0666 | 536.1831337 | 0.055222212 | 0.128704 | 0.429065 | 0.667876 | 0.83651 |
| Mfumv2_0667 | 141.375346 | -0.204592685 | 0.187675 | -1.09014 | 0.275651 | 0.516133 |
| Mfumv2_0668 | 1149.630502 | -0.172153629 | 0.108939 | -1.58027 | 0.114045 | 0.294651 |
| Mfumv2_0669 | 219.6643921 | -0.046593996 | 0.151238 | -0.30808 | 0.758018 | 0.884498 |
| Mfumv2_0670 | 726.6463449 | -0.017305489 | 0.115053 | -0.15041 | 0.880439 | 0.946389 |
| Mfumv2_0671 | 2119.726116 | -0.23417186 | 0.100153 | -2.33815 | 0.01938 | 0.084823 |
| Mfumv2_0672 | 1323.553601 | -0.097645567 | 0.095284 | -1.02479 | 0.305464 | 0.54695 |
| Mfumv2_0673 | 183.8468005 | 0.017690748 | 0.155925 | 0.113457 | 0.909668 | 0.962868 |
| Mfumv2_0674 | 1306.234971 | 0.617792574 | 0.165943 | 3.722914 | 0.000197 | 0.002248 |
| Mfumv2_0675 | 210.2998059 | 0.339247958 | 0.170526 | 1.989416 | 0.046655 | 0.156478 |
| Mfumv2_0676 | 778.49985 | -0.033971726 | 0.125393 | -0.27092 | 0.786451 | 0.89568 |
| Mfumv2_0677 | 995.2236386 | -0.001227562 | 0.118974 | -0.01032 | 0.991768 | 0.995456 |
| Mfumv2_0678 | 205.0589492 | -0.045125677 | 0.153795 | -0.29341 | 0.769205 | 0.887638 |
| Mfumv2_0679 | 95.58616902 | -0.227641939 | 0.230209 | -0.98885 | 0.322737 | 0.566462 |
| Mfumv2_0680 | 155.2211837 | -0.372747614 | 0.181826 | -2.05002 | 0.040362 | 0.140533 |
| Mfumv2_0681 | 40.35871581 | 0.132674795 | 0.282677 | 0.469351 | 0.638819 | 0.823866 |
| Mfumv2_0682 | 558.6790748 | -0.074101349 | 0.123825 | -0.59844 | 0.549549 | 0.76593 |
| Mfumv2_0683 | 47.2161633 | -0.006228758 | 0.285154 | -0.02184 | 0.982573 | 0.992996 |
| Mfumv2_0684 | 101.1922175 | 0.027709741 | 0.199415 | 0.138955 | 0.889486 | 0.94819 |
| Mfumv2_0685 | 136.1984892 | -0.02272961 | 0.173236 | -0.13121 | 0.895612 | 0.951499 |
| Mfumv2_0686 | 44.02174904 | 0.105253068 | 0.297956 | 0.353251 | 0.7239 | 0.86775 |
| Mfumv2_0687 | 127.4855645 | 0.202230282 | 0.177388 | 1.140047 | 0.254267 | 0.493394 |
| Mfumv2_0688 | 879.0939555 | 0.083280292 | 0.106843 | 0.779465 | 0.435706 | 0.666666 |
| Mfumv2_0689 | 129.4891331 | 0.347389672 | 0.249605 | 1.391759 | 0.163995 | 0.372279 |
| Mfumv2_0690 | 1981.136554 | 0.057605799 | 0.106115 | 0.542862 | 0.587225 | 0.7923 |
| Mfumv2_0691 | 678.5322259 | -0.546645496 | 0.14409 | -3.79377 | 0.000148 | 0.001733 |
| Mfumv2_0692 | 2223.668599 | -0.123230599 | 0.103391 | -1.19189 | 0.233306 | 0.470122 |
| Mfumv2_0693 | 657.7522793 | -0.148076992 | 0.13183 | -1.12324 | 0.261334 | 0.499274 |
| Mfumv2_0694 | 904.3513135 | 0.179841608 | 0.11118 | 1.617568 | 0.105756 | 0.282156 |
| Mfumv2_0695 | 517.2352389 | 0.410997955 | 0.136671 | 3.007197 | 0.002637 | 0.018203 |
| Mfumv2_0696 | 390.9014462 | 0.192042405 | 0.143591 | 1.337423 | 0.181085 | 0.401544 |
| Mfumv2_0697 | 676.0634776 | -0.075441849 | 0.11636 | -0.64835 | 0.516758 | 0.738908 |
| Mfumv2_0698 | 430.5255769 | 0.042457184 | 0.121643 | 0.34903 | 0.727067 | 0.869969 |
| Mfumv2_0699 | 775.5490882 | -0.009422862 | 0.114079 | -0.0826 | 0.93417 | 0.97544 |
| Mfumv2_0700 | 1082.505249 | -0.244013627 | 0.133932 | -1.82192 | 0.068466 | 0.208408 |
| Mfumv2_0701 | 172.521394 | -0.056062672 | 0.179068 | -0.31308 | 0.754219 | 0.884029 |
| Mfumv2_0702 | 211.1090856 | -0.066730975 | 0.153175 | -0.43565 | 0.66309 | 0.835454 |
| Mfumv2_0703 | 555.3890881 | -0.161506739 | 0.116545 | -1.38579 | 0.16581 | 0.37555 |
| Mfumv2_0704 | 179.4012318 | -0.146142005 | 0.161254 | -0.90628 | 0.364786 | 0.60367 |

| | | | | | | |
|---|---|---|---|---|---|---|
| Mfumv2_0705 | 1218.23012 | 0.316515945 | 0.105335 | 3.004842 | 0.002657 | 0.018282 |
| Mfumv2_0706 | 1549.532375 | -0.154544926 | 0.124717 | -1.23916 | 0.215286 | 0.446347 |
| Mfumv2_0707 | 5.657118027 | -0.993895574 | 0.797359 | -1.24649 | 0.212586 | NA |
| Mfumv2_0708 | 0.829241574 | 0.418343462 | 2.049175 | 0.204152 | 0.838235 | NA |
| Mfumv2_0709 | 47.31453199 | 0.190114653 | 0.277311 | 0.685565 | 0.492988 | 0.721349 |
| Mfumv2_0710 | 24.01671686 | 0.191858496 | 0.385341 | 0.497893 | 0.61856 | 0.813809 |
| Mfumv2_0711 | 122.7830992 | 0.328011983 | 0.21103 | 1.55434 | 0.120103 | 0.304013 |
| Mfumv2_0712 | 229.3093083 | -0.031201385 | 0.150504 | -0.20731 | 0.835766 | 0.921039 |
| Mfumv2_0713 | 2888.132594 | -0.092094906 | 0.110833 | -0.83093 | 0.406011 | 0.640249 |
| Mfumv2_0714 | 3911.503684 | -0.2409677 | 0.098448 | -2.44767 | 0.014379 | 0.067334 |
| Mfumv2_0715 | 252.7923597 | -0.054497212 | 0.156538 | -0.34814 | 0.727734 | 0.870248 |
| Mfumv2_0716 | 263.8643332 | -0.276012183 | 0.137427 | -2.00842 | 0.044599 | 0.150839 |
| Mfumv2_0717 | 193.223125 | -0.016795121 | 0.151343 | -0.11097 | 0.911637 | 0.963869 |
| Mfumv2_0718 | 992.1670303 | -0.392699888 | 0.116493 | -3.37102 | 0.000749 | 0.006628 |
| Mfumv2_0719 | 2885.585178 | -0.203758198 | 0.120598 | -1.68957 | 0.09111 | 0.253962 |
| Mfumv2_0720 | 1316.191815 | -0.271881922 | 0.14132 | -1.92387 | 0.05437 | 0.174768 |
| Mfumv2_0721 | 552.2480447 | -0.145684479 | 0.12779 | -1.14003 | 0.254275 | 0.493394 |
| Mfumv2_0722 | 1420.046359 | -0.164278025 | 0.104543 | -1.57139 | 0.116091 | 0.297864 |
| Mfumv2_0723 | 1282.983776 | -0.216076438 | 0.098543 | -2.19272 | 0.028328 | 0.109329 |
| Mfumv2_0724 | 434.7668293 | -0.088707462 | 0.161099 | -0.55064 | 0.58188 | 0.788797 |
| Mfumv2_0725 | 1.598361373 | -1.630680036 | 1.40106 | -1.16389 | 0.244469 | NA |
| Mfumv2_0726 | 0.394531882 | 1.763048078 | 2.763706 | 0.637929 | 0.52352 | NA |
| Mfumv2_0727 | 0 | NA | NA | NA | NA | NA |
| Mfumv2_0728 | 0.874050045 | 0.487960987 | 1.895991 | 0.257365 | 0.796897 | NA |
| Mfumv2_0729 | 480.8767679 | 0.225715233 | 0.125605 | 1.797024 | 0.072332 | 0.215979 |
| Mfumv2_0730 | 1.899869361 | 1.019428639 | 1.326717 | 0.768385 | 0.442259 | NA |
| Mfumv2_0731 | 0.420358933 | 1.837350107 | 2.71198 | 0.677494 | 0.498093 | NA |
| Mfumv2_0732 | 8.208383683 | 0.65652929 | 0.638329 | 1.028512 | 0.303709 | NA |
| Mfumv2_0733 | 8.769361922 | -0.261888456 | 0.624 | -0.41969 | 0.67471 | NA |
| Mfumv2_0734 | 157.6917548 | 0.212861736 | 0.232165 | 0.916857 | 0.359217 | 0.598398 |
| Mfumv2_0735 | 0.907859735 | 0.352631006 | 1.867036 | 0.188872 | 0.850193 | NA |
| Mfumv2_0736 | 1639.302043 | -0.073437728 | 0.126303 | -0.58144 | 0.560944 | 0.77267 |
| Mfumv2_0737 | 0.926563302 | 0.314832243 | 1.856113 | 0.169619 | 0.86531 | NA |
| Mfumv2_0738 | 0 | NA | NA | NA | NA | NA |
| Mfumv2_0739 | 2.304398366 | 1.26501189 | 1.274212 | 0.992779 | 0.320817 | NA |
| Mfumv2_0740 | 1242.149168 | 0.004073956 | 0.101416 | 0.040171 | 0.967957 | 0.98909 |
| Mfumv2_0741 | 1148.058596 | -0.126712728 | 0.100095 | -1.26592 | 0.205541 | 0.434665 |
| Mfumv2_0742 | 276.4289985 | -0.135843095 | 0.142189 | -0.95537 | 0.339391 | 0.583765 |
| Mfumv2_0743 | 311.2338638 | 0.128954966 | 0.15145 | 0.851467 | 0.39451 | 0.627033 |
| Mfumv2_0745 | 362.5920435 | -0.093073946 | 0.146599 | -0.63489 | 0.525503 | 0.748217 |
| Mfumv2_0746 | 87.32316111 | -0.269321725 | 0.272688 | -0.98765 | 0.323322 | 0.566462 |
| Mfumv2_0747 | 2.758112699 | 0.479263793 | 1.107107 | 0.432897 | 0.665089 | NA |
| Mfumv2_0748 | 47.36249618 | 0.836461341 | 0.291809 | 2.866465 | 0.004151 | 0.025978 |
| Mfumv2_0749 | 372.7319222 | 0.080000289 | 0.153383 | 0.521573 | 0.601968 | 0.801958 |
| Mfumv2_0750 | 128.9723911 | -0.133690633 | 0.174964 | -0.7641 | 0.444807 | 0.675447 |
| Mfumv2_0751 | 240.9526452 | -0.126464576 | 0.19402 | -0.65181 | 0.514521 | 0.737285 |
| Mfumv2_0752 | 44.12982383 | -0.34906279 | 0.396487 | -0.88039 | 0.378648 | 0.613471 |
| Mfumv2_0753 | 34.98903063 | 0.158714181 | 0.321026 | 0.494397 | 0.621026 | 0.815985 |
| Mfumv2_0754 | 74.69643919 | 0.067338118 | 0.238829 | 0.281951 | 0.777981 | 0.892523 |
| Mfumv2_0755 | 131.985828 | 0.049047406 | 0.185808 | 0.263968 | 0.791805 | 0.897706 |
| Mfumv2_0756 | 74.35279592 | 0.059006056 | 0.232924 | 0.253328 | 0.800015 | 0.902938 |
| Mfumv2_0757 | 40.98003188 | 0.26221017 | 0.304948 | 0.859853 | 0.38987 | 0.622615 |

| | | | | | | |
|---|---|---|---|---|---|---|
| Mfumv2_0758 | 4.861715039 | 0.8181066 | 0.854096 | 0.957862 | 0.338132 | NA |
| Mfumv2_0759 | 39.40289973 | 0.285927685 | 0.332915 | 0.858862 | 0.390417 | 0.622992 |
| Mfumv2_0760 | 1155.869442 | 0.475517814 | 0.162779 | 2.921243 | 0.003486 | 0.022594 |
| Mfumv2_0761 | 601.2224862 | 0.457738372 | 0.160236 | 2.856649 | 0.004281 | 0.026629 |
| Mfumv2_0762 | 261.8269008 | 0.447378503 | 0.173457 | 2.579194 | 0.009903 | 0.050883 |
| Mfumv2_0763 | 315.7990388 | 0.268245697 | 0.153572 | 1.746715 | 0.080687 | 0.232235 |
| Mfumv2_0764 | 133.7410602 | 0.39797607 | 0.179278 | 2.219882 | 0.026427 | 0.104101 |
| Mfumv2_0765 | 124.1750624 | -0.196228393 | 0.213615 | -0.91861 | 0.358301 | 0.59836 |
| Mfumv2_0766 | 26.75619565 | 0.162188564 | 0.349735 | 0.463746 | 0.642829 | 0.825732 |
| Mfumv2_0767 | 111.2126325 | 0.09959653 | 0.195632 | 0.509101 | 0.610682 | 0.807144 |
| Mfumv2_0768 | 173.822926 | 0.701105003 | 0.182949 | 3.832246 | 0.000127 | 0.001565 |
| Mfumv2_0769 | 198.316434 | 0.24815372 | 0.157123 | 1.579364 | 0.114253 | 0.294651 |
| Mfumv2_0770 | 49.46172944 | 0.277742583 | 0.34046 | 0.815786 | 0.414623 | 0.647422 |
| Mfumv2_0771 | 29.37646133 | 0.0703475 | 0.33667 | 0.208951 | 0.834487 | 0.920306 |
| Mfumv2_0772 | 180.7171336 | 0.158851757 | 0.171481 | 0.926353 | 0.354262 | 0.59836 |
| Mfumv2_0773 | 47.36568268 | -0.011252223 | 0.269865 | -0.0417 | 0.966741 | 0.988892 |
| Mfumv2_0774 | 153.5623964 | 0.330769481 | 0.164124 | 2.015361 | 0.043867 | 0.14929 |
| Mfumv2_0775 | 69.28241869 | 0.268382654 | 0.243403 | 1.102626 | 0.27019 | 0.508726 |
| Mfumv2_0776 | 9.174782261 | 0.155834459 | 0.610848 | 0.255112 | 0.798637 | NA |
| Mfumv2_0777 | 91.47722582 | 0.740412966 | 0.228078 | 3.246312 | 0.001169 | 0.009395 |
| Mfumv2_0778 | 7.376180942 | 0.480215289 | 0.64248 | 0.74744 | 0.454798 | NA |
| Mfumv2_0779 | 648.7399951 | -0.129802714 | 0.125401 | -1.0351 | 0.300622 | 0.540689 |
| Mfumv2_0780 | 5.353672895 | -0.111588542 | 0.74864 | -0.14906 | 0.88151 | NA |
| Mfumv2_0781 | 1.178916541 | 2.106015297 | 1.933563 | 1.089189 | 0.276071 | NA |
| Mfumv2_0782 | 92.23518543 | 0.362222681 | 0.203187 | 1.782702 | 0.074635 | 0.220827 |
| Mfumv2_0783 | 81.13609885 | 0.081378927 | 0.218809 | 0.371917 | 0.709955 | 0.857151 |
| Mfumv2_0784 | 434.9223593 | -0.268157272 | 0.129385 | -2.07255 | 0.038214 | 0.136699 |
| Mfumv2_0785 | 706.6434573 | -0.035805784 | 0.108536 | -0.3299 | 0.741477 | 0.876167 |
| Mfumv2_0786 | 316.5558704 | -0.041280651 | 0.129124 | -0.3197 | 0.749198 | 0.880198 |
| Mfumv2_0787 | 38.86068897 | 0.126602965 | 0.305687 | 0.414158 | 0.678758 | 0.842824 |
| Mfumv2_0788 | 279.9078501 | 0.020333277 | 0.146498 | 0.138796 | 0.889612 | 0.94819 |
| Mfumv2_0789 | 72.57135612 | 0.331613769 | 0.225024 | 1.473681 | 0.140568 | 0.331845 |
| Mfumv2_0790 | 127.1243086 | -0.013602317 | 0.180442 | -0.07538 | 0.93991 | 0.977742 |
| Mfumv2_0791 | 89.98022329 | 0.606855729 | 0.203866 | 2.976741 | 0.002913 | 0.019545 |
| Mfumv2_0792 | 334.1363265 | 0.044094024 | 0.147621 | 0.298697 | 0.765171 | 0.887034 |
| Mfumv2_0793 | 73.15069966 | 0.454559709 | 0.226558 | 2.006374 | 0.044816 | 0.151321 |
| Mfumv2_0794 | 263.8398283 | -0.165949971 | 0.155131 | -1.06974 | 0.284735 | 0.523113 |
| Mfumv2_0795 | 2557.984574 | -0.062718996 | 0.096036 | -0.65308 | 0.513706 | 0.737168 |
| Mfumv2_0796 | 0 | NA | NA | NA | NA | NA |
| Mfumv2_0797 | 106.1223858 | -0.01475598 | 0.191375 | -0.07711 | 0.93854 | 0.977463 |
| Mfumv2_0798 | 201.6681781 | -0.257361769 | 0.179246 | -1.4358 | 0.151059 | 0.348825 |
| Mfumv2_0799 | 443.446698 | -0.064326357 | 0.124512 | -0.51663 | 0.605416 | 0.80336 |
| Mfumv2_0800 | 769.9292352 | -0.175915655 | 0.112173 | -1.56826 | 0.116821 | 0.299354 |
| Mfumv2_0801 | 536.1691146 | -0.093167459 | 0.225297 | -0.41353 | 0.679217 | 0.842824 |
| Mfumv2_0802 | 689.3888302 | -0.467269893 | 0.13467 | -3.46974 | 0.000521 | 0.004984 |
| Mfumv2_0803 | 62.2711298 | 0.203571507 | 0.248981 | 0.817618 | 0.413576 | 0.647422 |
| Mfumv2_0804 | 58.73819491 | -0.165566413 | 0.274558 | -0.60303 | 0.546489 | 0.764552 |
| Mfumv2_0805 | 169.0403944 | 0.049895971 | 0.173571 | 0.287467 | 0.773755 | 0.889796 |
| Mfumv2_0806 | 153.2019524 | -0.113289493 | 0.164345 | -0.68934 | 0.49061 | 0.718917 |
| Mfumv2_0807 | 65.67196934 | 0.095392695 | 0.268558 | 0.355203 | 0.722438 | 0.867012 |
| Mfumv2_0808 | 180.3565615 | -0.05554418 | 0.154367 | -0.35982 | 0.718982 | 0.865449 |
| Mfumv2_0809 | 195.6436398 | -0.058354691 | 0.177146 | -0.32942 | 0.741842 | 0.876167 |

| | | | | | | |
|---|---|---|---|---|---|---|
| Mfumv2_0810 | 1278.417254 | -0.755478092 | 0.344641 | -2.19207 | 0.028374 | 0.109329 |
| Mfumv2_0811 | 130.4950221 | 0.199844609 | 0.177508 | 1.125833 | 0.260236 | 0.499079 |
| Mfumv2_0812 | 3.765244339 | -0.284779476 | 0.88056 | -0.32341 | 0.746387 | NA |
| Mfumv2_0813 | 211.5286251 | -0.251475817 | 0.154277 | -1.63003 | 0.103095 | 0.278385 |
| Mfumv2_0814 | 3253.773277 | -1.20651684 | 0.11411 | -10.5733 | 3.96E-26 | 4.98E-24 |
| Mfumv2_0815 | 4687.963863 | -5.06159242 | 0.161058 | -31.4272 | 8.61E-217 | 1.73E-213 |
| Mfumv2_0816 | 336.1063777 | 0.079747827 | 0.126964 | 0.628114 | 0.529929 | 0.751856 |
| Mfumv2_0817 | 675.9320309 | -0.067703884 | 0.130531 | -0.51868 | 0.603984 | 0.802869 |
| Mfumv2_0818 | 134.5830675 | 0.136694369 | 0.181531 | 0.753008 | 0.451445 | 0.680331 |
| Mfumv2_0819 | 305.9537326 | -0.125345396 | 0.139468 | -0.89874 | 0.368791 | 0.607278 |
| Mfumv2_0820 | 272.2652726 | -0.032592247 | 0.146879 | -0.2219 | 0.824393 | 0.914526 |
| Mfumv2_0821 | 1003.397981 | 0.168054041 | 0.133883 | 1.255231 | 0.209395 | 0.440497 |
| Mfumv2_0822 | 62.26823515 | -0.1949119 | 0.244997 | -0.79557 | 0.426283 | 0.659278 |
| Mfumv2_0823 | 987.205073 | -0.617666714 | 0.118536 | -5.21079 | 1.88E-07 | 4.72E-06 |
| Mfumv2_0824 | 1986.007578 | -0.231293879 | 0.102209 | -2.26295 | 0.023639 | 0.09769 |
| Mfumv2_0825 | 208.9177465 | -0.195953557 | 0.150861 | -1.2989 | 0.193977 | 0.421298 |
| Mfumv2_0826 | 5.310042353 | -0.595801078 | 0.800888 | -0.74393 | 0.456922 | NA |
| Mfumv2_0827 | 0.169414025 | -1.639708171 | 4.080473 | -0.40184 | 0.6878 | NA |
| Mfumv2_0828 | 0.257264584 | 1.152229179 | 3.235444 | 0.356127 | 0.721745 | NA |
| Mfumv2_0830 | 6.474608302 | 1.031789252 | 0.716124 | 1.440798 | 0.149642 | NA |
| Mfumv2_0831 | 219.5299847 | -0.444886772 | 0.150329 | -2.95942 | 0.003082 | 0.020236 |
| Mfumv2_0832 | 60.81898822 | -0.378955809 | 0.248307 | -1.52616 | 0.126971 | 0.312603 |
| Mfumv2_0833 | 30.77354086 | -0.527370022 | 0.41525 | -1.27001 | 0.204083 | 0.433912 |
| Mfumv2_0834 | 43.94415681 | 0.160838858 | 0.306 | 0.525617 | 0.599154 | 0.799801 |
| Mfumv2_0835 | 121.060859 | 0.262442 | 0.204674 | 1.282242 | 0.199758 | 0.429672 |
| Mfumv2_0837 | 1.536974112 | 1.453788138 | 1.626366 | 0.893888 | 0.371382 | NA |
| Mfumv2_0838 | 24.62008283 | 0.054530697 | 0.371639 | 0.14673 | 0.883345 | 0.946455 |
| Mfumv2_0839 | 56.98627454 | 0.125869953 | 0.245079 | 0.513589 | 0.60754 | 0.804604 |
| Mfumv2_0840 | 7.425781477 | 0.144644911 | 0.63573 | 0.227526 | 0.820015 | NA |
| Mfumv2_0841 | 1.913041198 | 0.310142621 | 1.242704 | 0.249571 | 0.802919 | NA |
| Mfumv2_0842 | 26.21624986 | 0.025394597 | 0.389607 | 0.06518 | 0.948031 | 0.980261 |
| Mfumv2_0843 | 53.98993448 | 0.380484625 | 0.273778 | 1.389754 | 0.164603 | 0.373237 |
| Mfumv2_0844 | 312.7443987 | -0.105445416 | 0.155842 | -0.67662 | 0.498649 | 0.726986 |
| Mfumv2_0845 | 112.2532026 | -0.004641499 | 0.193402 | -0.024 | 0.980853 | 0.992996 |
| Mfumv2_0846 | 237.2572412 | 0.363669832 | 0.155992 | 2.331338 | 0.019736 | 0.085709 |
| Mfumv2_0847 | 333.4168683 | 0.378899982 | 0.142402 | 2.660769 | 0.007796 | 0.042446 |
| Mfumv2_0848 | 207.2610193 | 0.121259878 | 0.157897 | 0.767968 | 0.442506 | 0.672462 |
| Mfumv2_0849 | 53.4515785 | -0.015510643 | 0.258961 | -0.0599 | 0.952239 | 0.981373 |
| Mfumv2_0850 | 36.58024983 | 0.083909449 | 0.315587 | 0.265884 | 0.790329 | 0.897706 |
| Mfumv2_0851 | 58.25739606 | 0.278530516 | 0.244279 | 1.140217 | 0.254196 | 0.493394 |
| Mfumv2_0852 | 374.0952423 | -0.155963601 | 0.137462 | -1.13459 | 0.256545 | 0.4951 |
| Mfumv2_0853 | 145.3271909 | 0.12959193 | 0.186709 | 0.694084 | 0.48763 | 0.715594 |
| Mfumv2_0854 | 167.6777469 | 0.038820408 | 0.168093 | 0.230946 | 0.817357 | 0.909463 |
| Mfumv2_0855 | 218.1284102 | -0.167817949 | 0.157223 | -1.06739 | 0.285796 | 0.523113 |
| Mfumv2_0856 | 21.51215716 | 0.179121959 | 0.384349 | 0.46604 | 0.641187 | NA |
| Mfumv2_0857 | 175.3081528 | 0.306174351 | 0.16163 | 1.894293 | 0.058186 | 0.183799 |
| Mfumv2_0858 | 89.31657513 | 0.112381507 | 0.202369 | 0.555331 | 0.578668 | 0.788261 |
| Mfumv2_0859 | 229.93448 | -0.355551284 | 0.148341 | -2.39685 | 0.016537 | 0.074323 |
| Mfumv2_0860 | 241.2264999 | -0.131467825 | 0.142223 | -0.92438 | 0.35529 | 0.59836 |
| Mfumv2_0861 | 137.5520906 | 0.437066031 | 0.178233 | 2.45222 | 0.014198 | 0.067114 |
| Mfumv2_0862 | 199.3698264 | 0.048611466 | 0.151934 | 0.319951 | 0.749006 | 0.880198 |
| Mfumv2_0863 | 11.86755994 | 0.415650038 | 0.561268 | 0.740555 | 0.458963 | NA |

| | | | | | | | |
|---|---|---|---|---|---|---|---|
| Mfumv2_0864 | 0.561049046 | | 2.254428531 | 2.470651 | 0.912483 | 0.361514 | NA |
| Mfumv2_0865 | 0.83752729 | | -0.006987464 | 2.08715 | -0.00335 | 0.997329 | NA |
| Mfumv2_0866 | 0.11657447 | | 0.283845181 | 4.080473 | 0.069562 | 0.944542 | NA |
| Mfumv2_0867 | 2.015074583 | | 3.025853302 | 1.559867 | 1.939814 | 0.052402 | NA |
| Mfumv2_0868 | 18.12921305 | | 0.120642512 | 0.424523 | 0.284184 | 0.77627 | NA |
| Mfumv2_0869 | 0 | NA | | NA | NA | NA | NA |
| Mfumv2_0870 | 69.01740439 | | -0.010332129 | 0.247333 | -0.04177 | 0.966679 | 0.988892 |
| Mfumv2_0871 | 1854.981418 | | -0.192099 | 0.107442 | -1.78793 | 0.073788 | 0.21929 |
| Mfumv2_0872 | 215.5059491 | | 0.384915608 | 0.163315 | 2.356888 | 0.018429 | 0.08155 |
| Mfumv2_0873 | 1423.057884 | | -0.044764945 | 0.107659 | -0.4158 | 0.677555 | 0.842824 |
| Mfumv2_0874 | 10977.64142 | | -0.42020381 | 0.104565 | -4.0186 | 5.85E-05 | 0.000817 |
| Mfumv2_0875 | 233.0454149 | | 0.039873357 | 0.205855 | 0.193696 | 0.846414 | 0.927178 |
| Mfumv2_0876 | 90.69382605 | | -0.007995198 | 0.211807 | -0.03775 | 0.969889 | 0.98909 |
| Mfumv2_0877 | 36.72314694 | | 0.090417969 | 0.295291 | 0.3062 | 0.759452 | 0.884498 |
| Mfumv2_0878 | 58.31251802 | | -0.24438855 | 0.248593 | -0.98309 | 0.325564 | 0.568517 |
| Mfumv2_0879 | 206.9219185 | | -0.056297172 | 0.15071 | -0.37355 | 0.708742 | 0.856202 |
| Mfumv2_0880 | 401.0644958 | | -0.215781682 | 0.13214 | -1.63298 | 0.102473 | 0.277826 |
| Mfumv2_0881 | 328.7818744 | | -0.077518207 | 0.142532 | -0.54387 | 0.586534 | 0.791899 |
| Mfumv2_0883 | 68.52302352 | | 0.244671178 | 0.233894 | 1.046077 | 0.295525 | 0.534877 |
| Mfumv2_0884 | 104.0419804 | | 0.247909722 | 0.19159 | 1.293963 | 0.195678 | 0.424075 |
| Mfumv2_0885 | 164.6638815 | | 0.27741523 | 0.164274 | 1.688738 | 0.09127 | 0.253962 |
| Mfumv2_0886 | 622.1023442 | | -0.115187159 | 0.116364 | -0.98989 | 0.322229 | 0.566462 |
| Mfumv2_0887 | 179.373474 | | 0.213894389 | 0.201103 | 1.063604 | 0.287508 | 0.525338 |
| Mfumv2_0888 | 234.511139 | | 0.084473561 | 0.164486 | 0.513562 | 0.607558 | 0.804604 |
| Mfumv2_0889 | 37.47858812 | | -0.253228338 | 0.298848 | -0.84735 | 0.396801 | 0.628189 |
| Mfumv2_0890 | 94.59277897 | | 0.195025202 | 0.231334 | 0.843046 | 0.399203 | 0.631495 |
| Mfumv2_0891 | 105.1532157 | | -0.337399738 | 0.206452 | -1.63428 | 0.102201 | 0.277826 |
| Mfumv2_0892 | 46.53673833 | | 0.1571528 | 0.27458 | 0.572338 | 0.567093 | 0.777672 |
| Mfumv2_0893 | 95.21136351 | | 0.146417688 | 0.204602 | 0.715623 | 0.474224 | 0.701558 |
| Mfumv2_0894 | 830.5340724 | | -0.644549751 | 0.165191 | -3.90184 | 9.55E-05 | 0.001237 |
| Mfumv2_0895 | 1.788181011 | | 0.328032875 | 1.333228 | 0.246044 | 0.805648 | NA |
| Mfumv2_0896 | 4.28929312 | | -0.409373836 | 0.828589 | -0.49406 | 0.621263 | NA |
| Mfumv2_0898 | 11.4861887 | | 0.979351836 | 0.544818 | 1.797576 | 0.072244 | NA |
| Mfumv2_0899 | 0 | NA | | NA | NA | NA | NA |
| Mfumv2_0900 | 125.8532586 | | 0.628646145 | 0.182503 | 3.444585 | 0.000572 | 0.00542 |
| Mfumv2_0902 | 1091.670921 | | 1.15238191 | 0.147752 | 7.799441 | 6.22E-15 | 4.03E-13 |
| Mfumv2_0903 | 55.22693879 | | 0.607577228 | 0.312635 | 1.943408 | 0.051967 | 0.169476 |
| Mfumv2_0904 | 9.727393759 | | -0.108782794 | 0.563077 | -0.19319 | 0.846807 | NA |
| Mfumv2_0905 | 28.43764665 | | 0.116952876 | 0.337503 | 0.346524 | 0.728949 | 0.871183 |
| Mfumv2_0906 | 25.64686453 | | 0.097582943 | 0.358622 | 0.272105 | 0.785541 | 0.89568 |
| Mfumv2_0907 | 92.53122371 | | 0.35134502 | 0.224368 | 1.565932 | 0.117364 | 0.300167 |
| Mfumv2_0908 | 437.5286492 | | 0.102028938 | 0.123528 | 0.82596 | 0.408827 | 0.642476 |
| Mfumv2_0909 | 187.126847 | | -0.240829074 | 0.162452 | -1.48247 | 0.138217 | 0.328735 |
| Mfumv2_0910 | 427.556853 | | -0.116787964 | 0.130025 | -0.89819 | 0.369082 | 0.607278 |
| Mfumv2_0911 | 91.13046155 | | 0.614828836 | 0.222311 | 2.76562 | 0.005681 | 0.032483 |
| Mfumv2_0912 | 150.9728434 | | 0.092170241 | 0.209409 | 0.440145 | 0.659832 | 0.835288 |
| Mfumv2_0913 | 85.80332254 | | -0.097178528 | 0.209986 | -0.46279 | 0.643517 | 0.826087 |
| Mfumv2_0914 | 288.8429471 | | 0.066291252 | 0.139723 | 0.474447 | 0.635181 | 0.822746 |
| Mfumv2_0915 | 570.7640601 | | -0.002675622 | 0.115426 | -0.02318 | 0.981506 | 0.992996 |
| Mfumv2_0916 | 1.750870311 | | 1.522538317 | 1.518032 | 1.002968 | 0.315876 | NA |
| Mfumv2_0917 | 55.40898081 | | -0.178591197 | 0.2556 | -0.69871 | 0.484731 | 0.71186 |
| Mfumv2_0918 | 996.5420463 | | -0.155922413 | 0.107077 | -1.45617 | 0.145346 | 0.338354 |

| | | | | | | |
|---|---|---|---|---|---|---|
| Mfumv2_0919 | 624.9067415 | -0.089530491 | 0.121989 | -0.73392 | 0.462996 | 0.691569 |
| Mfumv2_0920 | 549.004244 | 0.104672338 | 0.115569 | 0.905716 | 0.365086 | 0.60367 |
| Mfumv2_0921 | 2571.142886 | -0.271413455 | 0.091313 | -2.97235 | 0.002955 | 0.01966 |
| Mfumv2_0922 | 555.0331363 | -0.403620292 | 0.159782 | -2.52607 | 0.011535 | 0.056658 |
| Mfumv2_0923 | 423.2806142 | -0.132252932 | 0.154886 | -0.85387 | 0.393176 | 0.625684 |
| Mfumv2_0924 | 831.6017019 | 0.492058306 | 0.113147 | 4.34884 | 1.37E-05 | 0.000243 |
| Mfumv2_0925 | 1333.174642 | -0.040010768 | 0.104013 | -0.38467 | 0.700481 | 0.851856 |
| Mfumv2_0926 | 154.8611205 | 0.063971798 | 0.205296 | 0.311607 | 0.755339 | 0.884498 |
| Mfumv2_0927 | 3.873751473 | -1.661837572 | 0.939504 | -1.76885 | 0.07692 | NA |
| Mfumv2_0928 | 0.695125769 | -0.609248876 | 2.100287 | -0.29008 | 0.771756 | NA |
| Mfumv2_0929 | 0.82919313 | 1.449141952 | 2.083264 | 0.695611 | 0.486672 | NA |
| Mfumv2_0930 | 72.33392082 | -0.162274147 | 0.225225 | -0.7205 | 0.471218 | 0.698655 |
| Mfumv2_0931 | 490.5763022 | 0.281783961 | 0.13341 | 2.112162 | 0.034673 | 0.126651 |
| Mfumv2_0933 | 1639.990641 | -0.331758305 | 0.114844 | -2.88877 | 0.003868 | 0.024511 |
| Mfumv2_0934 | 1270.114058 | -0.298851552 | 0.115303 | -2.59189 | 0.009545 | 0.04955 |
| Mfumv2_0935 | 596.0338839 | 0.028003468 | 0.118294 | 0.236728 | 0.812868 | 0.907755 |
| Mfumv2_0936 | 740.13308 | -0.369924744 | 0.106051 | -3.48819 | 0.000486 | 0.004807 |
| Mfumv2_0937 | 443.9102595 | -0.266963703 | 0.13892 | -1.92171 | 0.054642 | 0.175082 |
| Mfumv2_0938 | 776.6652999 | -0.187019722 | 0.126419 | -1.47937 | 0.139042 | 0.330184 |
| Mfumv2_0939 | 435.6943383 | -0.312532435 | 0.138337 | -2.25922 | 0.02387 | 0.097866 |
| Mfumv2_0940 | 2794.547883 | -0.092336372 | 0.090769 | -1.01727 | 0.309024 | 0.551359 |
| Mfumv2_0941 | 2429.14768 | -0.345305711 | 0.088532 | -3.90034 | 9.61E-05 | 0.001237 |
| Mfumv2_0942 | 103.598841 | 0.945644866 | 0.230651 | 4.099899 | 4.13E-05 | 0.000624 |
| Mfumv2_0943 | 384.7272663 | 3.05843653 | 0.173842 | 17.59316 | 2.78E-69 | 1.12E-66 |
| Mfumv2_0944 | 251.9388523 | -0.052370026 | 0.146416 | -0.35768 | 0.720584 | 0.866005 |
| Mfumv2_0945 | 72.13040455 | -0.123352421 | 0.257566 | -0.47892 | 0.631998 | 0.82074 |
| Mfumv2_0946 | 63.20326054 | 0.39048826 | 0.237337 | 1.645293 | 0.099909 | 0.273086 |
| Mfumv2_0947 | 143.0722328 | 0.029355384 | 0.16594 | 0.176904 | 0.859584 | 0.936499 |
| Mfumv2_0948 | 197.7457169 | 0.438100503 | 0.158369 | 2.766335 | 0.005669 | 0.032483 |
| Mfumv2_0949 | 2252.21684 | 0.059634294 | 0.131214 | 0.454483 | 0.649482 | 0.829769 |
| Mfumv2_0950 | 223.1863158 | 0.078368261 | 0.168629 | 0.464738 | 0.642119 | 0.825347 |
| Mfumv2_0951 | 2371.093959 | -0.127194626 | 0.13011 | -0.97759 | 0.328275 | 0.571494 |
| Mfumv2_0952 | 5975.641949 | -0.360190394 | 0.134575 | -2.67651 | 0.007439 | 0.040914 |
| Mfumv2_0953 | 530.4231339 | -0.153004593 | 0.122091 | -1.2532 | 0.210133 | 0.441126 |
| Mfumv2_0954 | 1055.355722 | -0.285454193 | 0.11271 | -2.53264 | 0.011321 | 0.056156 |
| Mfumv2_0955 | 860.829259 | -0.262381966 | 0.102367 | -2.56316 | 0.010372 | 0.052489 |
| Mfumv2_0956 | 260.2965182 | -0.270129774 | 0.139465 | -1.9369 | 0.052757 | 0.170676 |
| Mfumv2_0957 | 1668.451246 | -0.22565386 | 0.107307 | -2.10288 | 0.035476 | 0.128883 |
| Mfumv2_0958 | 316.6670561 | -0.325504051 | 0.131399 | -2.47723 | 0.013241 | 0.063335 |
| Mfumv2_0959 | 543.8834612 | -0.361047564 | 0.115486 | -3.12633 | 0.00177 | 0.012978 |
| Mfumv2_0960 | 1747.784487 | -0.239396683 | 0.099362 | -2.40933 | 0.015982 | 0.072871 |
| Mfumv2_0961 | 775.1813075 | -0.380051662 | 0.1198 | -3.17239 | 0.001512 | 0.01147 |
| Mfumv2_0962 | 395.4004373 | -0.044758944 | 0.129895 | -0.34458 | 0.730412 | 0.871376 |
| Mfumv2_0963 | 500.6318084 | -0.025449472 | 0.114194 | -0.22286 | 0.823644 | 0.914199 |
| Mfumv2_0964 | 586.9583241 | -0.178829077 | 0.11791 | -1.51666 | 0.129352 | 0.316912 |
| Mfumv2_0965 | 781.7671759 | -0.140027478 | 0.113218 | -1.2368 | 0.216163 | 0.447702 |
| Mfumv2_0966 | 1666.030341 | -0.297613325 | 0.107525 | -2.76786 | 0.005643 | 0.032482 |
| Mfumv2_0967 | 654.4150476 | -0.331140082 | 0.116541 | -2.8414 | 0.004492 | 0.027511 |
| Mfumv2_0968 | 877.9659197 | -0.234318521 | 0.100991 | -2.32018 | 0.020331 | 0.087463 |
| Mfumv2_0969 | 652.6631001 | -0.217948747 | 0.127248 | -1.71278 | 0.086753 | 0.245129 |
| Mfumv2_0970 | 1710.736462 | -0.214670634 | 0.103948 | -2.06517 | 0.038907 | 0.137855 |
| Mfumv2_0971 | 1666.35148 | -0.250985861 | 0.105661 | -2.37539 | 0.01753 | 0.07809 |

| | | | | | | |
|---|---|---|---|---|---|---|
| Mfumv2_0972 | 1852.198124 | -0.177320751 | 0.100625 | -1.7622 | 0.078036 | 0.227347 |
| Mfumv2_0973 | 255.2773672 | -0.170482292 | 0.148988 | -1.14426 | 0.252514 | 0.492248 |
| Mfumv2_0974 | 513.878774 | -0.085211185 | 0.117032 | -0.7281 | 0.466552 | 0.694814 |
| Mfumv2_0975 | 673.367577 | -0.386850983 | 0.146425 | -2.64198 | 0.008242 | 0.044275 |
| Mfumv2_0977 | 25.93839228 | 0.190610091 | 0.364852 | 0.522431 | 0.60137 | 0.801694 |
| Mfumv2_0978 | 10486.71234 | -0.444609757 | 0.118687 | -3.74608 | 0.00018 | 0.002062 |
| Mfumv2_0979 | 9300.521347 | -0.259407758 | 0.13403 | -1.93545 | 0.052935 | 0.170701 |
| Mfumv2_0980 | 357.2446677 | 0.144021245 | 0.137255 | 1.049294 | 0.294043 | 0.533634 |
| Mfumv2_0981 | 411.4905484 | 0.061398979 | 0.153256 | 0.40063 | 0.688693 | 0.846498 |
| Mfumv2_0982 | 260.7615332 | 0.123049953 | 0.14495 | 0.848915 | 0.395929 | 0.627515 |
| Mfumv2_0984 | 4.881675724 | 0.282994712 | 0.795093 | 0.355927 | 0.721895 | NA |
| Mfumv2_0985 | 99.98746777 | 0.217024905 | 0.208738 | 1.0397 | 0.298479 | 0.537798 |
| Mfumv2_0986 | 43.93443211 | -0.125404813 | 0.288549 | -0.4346 | 0.663849 | 0.835454 |
| Mfumv2_0988 | 1424.700424 | -0.475316329 | 0.133885 | -3.55019 | 0.000385 | 0.003966 |
| Mfumv2_0989 | 342.2492482 | 0.064570285 | 0.132137 | 0.488661 | 0.625082 | 0.818317 |
| Mfumv2_0990 | 252.6860752 | -0.135727515 | 0.153627 | -0.88348 | 0.376974 | 0.611746 |
| Mfumv2_0991 | 812.9598205 | -0.087731581 | 0.109562 | -0.80075 | 0.423277 | 0.656145 |
| Mfumv2_0992 | 1670.508304 | -0.042723803 | 0.102269 | -0.41776 | 0.676122 | 0.842824 |
| Mfumv2_0993 | 1127.648077 | 0.038542847 | 0.118674 | 0.324779 | 0.745348 | 0.878227 |
| Mfumv2_0994 | 557.9483658 | -0.091266521 | 0.112326 | -0.81252 | 0.416496 | 0.649139 |
| Mfumv2_0995 | 18.03119865 | -0.041176479 | 0.417416 | -0.09865 | 0.921419 | NA |
| Mfumv2_0996 | 26.31018061 | 0.662152244 | 0.357929 | 1.849955 | 0.06432 | 0.197885 |
| Mfumv2_0997 | 271.4553203 | 0.079599351 | 0.225626 | 0.352793 | 0.724244 | 0.86775 |
| Mfumv2_0998 | 32.69360818 | -0.071143936 | 0.351193 | -0.20258 | 0.839465 | 0.924101 |
| Mfumv2_0999 | 6.052777185 | -0.137742998 | 0.739256 | -0.18633 | 0.852189 | NA |
| Mfumv2_1000 | 59.68194664 | -0.193287099 | 0.246881 | -0.78292 | 0.433676 | 0.666097 |
| Mfumv2_1001 | 356.7020353 | -0.15926826 | 0.209855 | -0.75894 | 0.447887 | 0.677564 |
| Mfumv2_1002 | 0.829241574 | 0.418343462 | 2.049175 | 0.204152 | 0.838235 | NA |
| Mfumv2_1003 | 0.618170572 | 0.743117098 | 2.264584 | 0.328147 | 0.7428 | NA |
| Mfumv2_1004 | 0.402562965 | 0.092878996 | 2.826527 | 0.03286 | 0.973786 | NA |
| Mfumv2_1005 | 300.5058659 | -0.002126943 | 0.136079 | -0.01563 | 0.987529 | 0.994355 |
| Mfumv2_1006 | 14.26691669 | 0.00503286 | 0.493747 | 0.010193 | 0.991867 | NA |
| Mfumv2_1007 | 323.8442981 | 0.016865323 | 0.150926 | 0.111745 | 0.911025 | 0.963797 |
| Mfumv2_1008 | 711.0378215 | 0.146278219 | 0.109601 | 1.334642 | 0.181994 | 0.402228 |
| Mfumv2_1009 | 346.6019362 | 0.113256727 | 0.172627 | 0.656079 | 0.511773 | 0.735445 |
| Mfumv2_1010 | 519.0024972 | 0.008794959 | 0.117712 | 0.074716 | 0.940441 | 0.977742 |
| Mfumv2_1011 | 826.0074543 | -0.403067032 | 0.16857 | -2.3911 | 0.016798 | 0.075329 |
| Mfumv2_1012 | 409.1899109 | 0.004049314 | 0.146653 | 0.027612 | 0.977972 | 0.991795 |
| Mfumv2_1013 | 132.0923851 | 0.30617011 | 0.184563 | 1.658889 | 0.097138 | 0.26733 |
| Mfumv2_1014 | 446.9299759 | 0.135337387 | 0.12493 | 1.083302 | 0.278674 | 0.518866 |
| Mfumv2_1015 | 320.9229438 | -0.034133235 | 0.133911 | -0.25489 | 0.798805 | 0.902079 |
| Mfumv2_1016 | 489.5484546 | -0.408962107 | 0.117536 | -3.47945 | 0.000502 | 0.004853 |
| Mfumv2_1017 | 186.0845831 | -0.167006683 | 0.171166 | -0.9757 | 0.329213 | 0.572136 |
| Mfumv2_1018 | 97.17226376 | -0.222431369 | 0.196156 | -1.13395 | 0.256814 | 0.495144 |
| Mfumv2_1019 | 454.9347052 | 0.072946208 | 0.119777 | 0.609015 | 0.542514 | 0.761318 |
| Mfumv2_1020 | 76.49601784 | -0.103516981 | 0.23146 | -0.44723 | 0.654706 | 0.832295 |
| Mfumv2_1021 | 17.24473865 | -0.313729196 | 0.429603 | -0.73028 | 0.46522 | NA |
| Mfumv2_1022 | 21.64731878 | -0.351283548 | 0.378538 | -0.928 | 0.353407 | NA |
| Mfumv2_1023 | 2.815863597 | -0.244808429 | 1.047824 | -0.23363 | 0.815268 | NA |
| Mfumv2_1024 | 11755.56163 | -0.379321372 | 0.096283 | -3.93966 | 8.16E-05 | 0.001071 |
| Mfumv2_1025 | 1380.450489 | -0.119590875 | 0.09385 | -1.27428 | 0.202565 | 0.43339 |
| Mfumv2_1026 | 3306.219528 | 0.007142279 | 0.087823 | 0.081326 | 0.935183 | 0.975484 |

| | | | | | | |
|---|---|---|---|---|---|---|
| Mfumv2_1027 | 867.9464599 | -0.112597039 | 0.104741 | -1.075 | 0.282373 | 0.522845 |
| Mfumv2_1028 | 1776.986271 | -0.42975916 | 0.105666 | -4.06716 | 4.76E-05 | 0.000698 |
| Mfumv2_1029 | 718.5747071 | -0.479830435 | 0.141719 | -3.3858 | 0.00071 | 0.006365 |
| Mfumv2_1030 | 1565.252707 | -0.55989434 | 0.104823 | -5.34133 | 9.23E-08 | 2.47E-06 |
| Mfumv2_1031 | 6675.286705 | -0.37005071 | 0.097238 | -3.8056 | 0.000141 | 0.001666 |
| Mfumv2_1032 | 6628.44012 | -0.327728741 | 0.091255 | -3.59136 | 0.000329 | 0.003497 |
| Mfumv2_1033 | 432.9249098 | 0.050571718 | 0.123645 | 0.409009 | 0.682533 | 0.843309 |
| Mfumv2_1034 | 72.39265709 | -0.025624245 | 0.219333 | -0.11683 | 0.906996 | 0.960546 |
| Mfumv2_1035 | 748.3283697 | -0.089827801 | 0.113543 | -0.79114 | 0.428864 | 0.662643 |
| Mfumv2_1036 | 1081.176439 | -0.123503056 | 0.107429 | -1.14963 | 0.250297 | 0.490502 |
| Mfumv2_1037 | 1798.553064 | -0.325679896 | 0.117351 | -2.77527 | 0.005516 | 0.032118 |
| Mfumv2_1038 | 16.38030547 | 0.901631505 | 0.452595 | 1.992136 | 0.046356 | NA |
| Mfumv2_1039 | 0.57197699 | -0.591129077 | 2.523381 | -0.23426 | 0.814783 | NA |
| Mfumv2_1041 | 2.702337201 | 0.181451801 | 1.046372 | 0.17341 | 0.862329 | NA |
| Mfumv2_1042 | 0.867914164 | -0.166206782 | 1.928429 | -0.08619 | 0.931317 | NA |
| Mfumv2_1043 | 112.2962413 | 0.252149917 | 0.206027 | 1.223869 | 0.221002 | 0.455785 |
| Mfumv2_1044 | 584.3561392 | -0.360181724 | 0.118403 | -3.04201 | 0.00235 | 0.016508 |
| Mfumv2_1045 | 3432.531736 | -0.020198794 | 0.13679 | -0.14766 | 0.882609 | 0.946455 |
| Mfumv2_1046 | 385.3174764 | 0.034749904 | 0.128036 | 0.271407 | 0.786078 | 0.89568 |
| Mfumv2_1047 | 123.7793923 | -0.057705311 | 0.182517 | -0.31616 | 0.751877 | 0.881799 |
| Mfumv2_1048 | 346.4492871 | 0.058925648 | 0.124733 | 0.472412 | 0.636632 | 0.823208 |
| Mfumv2_1049 | 320.0858216 | -0.062711059 | 0.13192 | -0.47537 | 0.634521 | 0.822746 |
| Mfumv2_1050 | 444.0566308 | -0.179601977 | 0.156647 | -1.14654 | 0.251571 | 0.49164 |
| Mfumv2_1051 | 286.3275456 | -0.202981099 | 0.162695 | -1.24762 | 0.212172 | 0.443494 |
| Mfumv2_1052 | 379.1557919 | 0.117354008 | 0.127683 | 0.919103 | 0.358042 | 0.59836 |
| Mfumv2_1053 | 397.4836418 | -0.208359095 | 0.140188 | -1.48628 | 0.137204 | 0.327557 |
| Mfumv2_1054 | 89.95861229 | -0.031943953 | 0.206413 | -0.15476 | 0.877013 | 0.945235 |
| Mfumv2_1056 | 425.6763642 | 0.093623905 | 0.12514 | 0.748151 | 0.454369 | 0.683254 |
| Mfumv2_1057 | 75.93697188 | 0.326643602 | 0.244557 | 1.335652 | 0.181663 | 0.402228 |
| Mfumv2_1058 | 255.1443983 | 0.241519081 | 0.157871 | 1.529848 | 0.126054 | 0.311041 |
| Mfumv2_1060 | 561.5144394 | -0.157602521 | 0.116435 | -1.35357 | 0.175874 | 0.392155 |
| Mfumv2_1061 | 494.0937405 | -0.038501042 | 0.11676 | -0.32974 | 0.741593 | 0.876167 |
| Mfumv2_1062 | 193.9443247 | 0.157943418 | 0.166071 | 0.951057 | 0.341575 | 0.584519 |
| Mfumv2_1063 | 0 | NA | NA | NA | NA | NA |
| Mfumv2_1064 | 379.7176927 | -0.05158333 | 0.158809 | -0.32481 | 0.745322 | 0.878227 |
| Mfumv2_1065 | 88.14946677 | -0.084701016 | 0.210895 | -0.40163 | 0.687959 | 0.846498 |
| Mfumv2_1066 | 190.4006506 | 0.149512563 | 0.159017 | 0.940229 | 0.3471 | 0.590453 |
| Mfumv2_1067 | 255.6457569 | 0.124036451 | 0.156869 | 0.790701 | 0.429118 | 0.662643 |
| Mfumv2_1068 | 740.0815763 | -0.615200095 | 0.113611 | -5.41499 | 6.13E-08 | 1.69E-06 |
| Mfumv2_1069 | 2.055442718 | 0.488122856 | 1.247063 | 0.391418 | 0.695488 | NA |
| Mfumv2_1070 | 31.04043271 | 0.261041513 | 0.338125 | 0.772026 | 0.440099 | 0.670326 |
| Mfumv2_1071 | 0.479518162 | -1.365869326 | 2.681855 | -0.5093 | 0.610542 | NA |
| Mfumv2_1072 | 1.444777275 | -0.971745365 | 1.530082 | -0.63509 | 0.525367 | NA |
| Mfumv2_1073 | 316.5598553 | -0.398015376 | 0.15112 | -2.63377 | 0.008444 | 0.044999 |
| Mfumv2_1074 | 427.2922502 | 0.257819452 | 0.137575 | 1.874022 | 0.060927 | 0.190659 |
| Mfumv2_1075 | 67.90340911 | 0.140773228 | 0.240644 | 0.584986 | 0.558557 | 0.771231 |
| Mfumv2_1076 | 40.99347204 | 0.192376471 | 0.286976 | 0.670358 | 0.50263 | 0.730021 |
| Mfumv2_1077 | 74.05037437 | 0.268763276 | 0.220948 | 1.216411 | 0.223828 | 0.45844 |
| Mfumv2_1078 | 63.23556906 | -0.557433271 | 0.244745 | -2.27761 | 0.02275 | 0.095616 |
| Mfumv2_1079 | 205.9738007 | -0.237493171 | 0.149344 | -1.59024 | 0.11178 | 0.291291 |
| Mfumv2_1080 | 301.2760601 | -0.097828075 | 0.133198 | -0.73445 | 0.462672 | 0.691569 |
| Mfumv2_1081 | 207.9862304 | 0.098160899 | 0.149403 | 0.657022 | 0.511167 | 0.735445 |

| | | | | | | |
|---|---|---|---|---|---|---|
| Mfumv2_1082 | 600.3231398 | 0.121192616 | 0.115425 | 1.049968 | 0.293733 | 0.533553 |
| Mfumv2_1083 | 52.98080898 | 0.793889616 | 0.270699 | 2.932743 | 0.00336 | 0.021844 |
| Mfumv2_1084 | 587.4670161 | 0.470913065 | 0.115872 | 4.064063 | 4.82E-05 | 0.000702 |
| Mfumv2_1085 | 223.090242 | 0.133785092 | 0.152658 | 0.876373 | 0.380827 | 0.615513 |
| Mfumv2_1086 | 388.793162 | -0.097279949 | 0.123344 | -0.78869 | 0.430293 | 0.663946 |
| Mfumv2_1087 | 107.8934346 | -0.11087561 | 0.186801 | -0.59355 | 0.552815 | 0.766992 |
| Mfumv2_1088 | 5.021012452 | 1.667975387 | 0.883192 | 1.888576 | 0.058949 | NA |
| Mfumv2_1089 | 723.0164258 | -0.06433853 | 0.116668 | -0.55147 | 0.581315 | 0.788739 |
| Mfumv2_1090 | 233.5164472 | 0.148033345 | 0.15961 | 0.927468 | 0.353684 | 0.598106 |
| Mfumv2_1091 | 299.2412888 | -0.145671467 | 0.135833 | -1.07243 | 0.283526 | 0.523113 |
| Mfumv2_1092 | 1256.274304 | -0.364887151 | 0.095663 | -3.81428 | 0.000137 | 0.001633 |
| Mfumv2_1093 | 126.7555636 | -0.284887065 | 0.179184 | -1.58991 | 0.111855 | 0.291291 |
| Mfumv2_1094 | 85.46224124 | -0.399614255 | 0.21418 | -1.86578 | 0.062072 | 0.192145 |
| Mfumv2_1095 | 152.3274756 | -0.085124313 | 0.193088 | -0.44086 | 0.659316 | 0.835162 |
| Mfumv2_1097 | 160.6978141 | -0.01701587 | 0.162756 | -0.10455 | 0.916734 | 0.967289 |
| Mfumv2_1098 | 626.9105522 | -0.088205205 | 0.115572 | -0.7632 | 0.445342 | 0.675749 |
| Mfumv2_1099 | 5.483205655 | 0.15936131 | 0.775548 | 0.205482 | 0.837195 | NA |
| Mfumv2_1100 | 4.545436557 | 0.560435667 | 0.936419 | 0.598488 | 0.549514 | NA |
| Mfumv2_1101 | 1038.455169 | 0.04126979 | 0.105831 | 0.389958 | 0.696567 | 0.850369 |
| Mfumv2_1102 | 136.8559816 | -0.045987343 | 0.179068 | -0.25681 | 0.797322 | 0.901925 |
| Mfumv2_1103 | 208.9044438 | -0.101339391 | 0.15657 | -0.64725 | 0.517472 | 0.739403 |
| Mfumv2_1104 | 908.675255 | -0.062602077 | 0.116241 | -0.53855 | 0.590196 | 0.793808 |
| Mfumv2_1105 | 346.9167254 | 0.008000835 | 0.133789 | 0.059802 | 0.952313 | 0.981373 |
| Mfumv2_1106 | 629.5062812 | 0.050198481 | 0.123479 | 0.406536 | 0.684349 | 0.844506 |
| Mfumv2_1107 | 617.729662 | -0.168343249 | 0.127082 | -1.32468 | 0.185276 | 0.40507 |
| Mfumv2_1108 | 244.155525 | -0.22255297 | 0.151147 | -1.47243 | 0.140904 | 0.33186 |
| Mfumv2_1109 | 591.88183 | -0.089373676 | 0.146805 | -0.60879 | 0.542661 | 0.761318 |
| Mfumv2_1110 | 107.7477337 | -0.104713835 | 0.18774 | -0.55776 | 0.577008 | 0.787506 |
| Mfumv2_1111 | 459.5450963 | -0.500445599 | 0.155157 | -3.22542 | 0.001258 | 0.00991 |
| Mfumv2_1112 | 390.4909106 | 0.009129358 | 0.16358 | 0.05581 | 0.955493 | 0.982524 |
| Mfumv2_1113 | 1209.986709 | -0.088587378 | 0.10182 | -0.87004 | 0.384277 | 0.617611 |
| Mfumv2_1114 | 2581.577597 | 0.099850581 | 0.093905 | 1.06331 | 0.287641 | 0.525338 |
| Mfumv2_1115 | 275.4629421 | -0.182305742 | 0.144331 | -1.26311 | 0.206549 | 0.436338 |
| Mfumv2_1116 | 106.6728041 | 0.092794512 | 0.196014 | 0.473407 | 0.635923 | 0.823176 |
| Mfumv2_1117 | 512.852944 | -0.142811013 | 0.133686 | -1.06826 | 0.285403 | 0.523113 |
| Mfumv2_1118 | 538.6905376 | 0.14508508 | 0.143078 | 1.014029 | 0.310569 | 0.55222 |
| Mfumv2_1119 | 422.4811867 | -0.279028652 | 0.125237 | -2.228 | 0.02588 | 0.103162 |
| Mfumv2_1120 | 327.2862437 | 0.078004455 | 0.146257 | 0.533337 | 0.5938 | 0.795297 |
| Mfumv2_1121 | 33.50230116 | 0.578121362 | 0.331808 | 1.742337 | 0.08145 | 0.234023 |
| Mfumv2_1122 | 17.09510638 | 0.27854973 | 0.448568 | 0.620976 | 0.534616 | NA |
| Mfumv2_1123 | 28.76024274 | 0.750357902 | 0.340661 | 2.202651 | 0.027619 | 0.107533 |
| Mfumv2_1124 | 224.646415 | 0.061534294 | 0.148298 | 0.414936 | 0.678189 | 0.842824 |
| Mfumv2_1127 | 6864.866 | -0.409394495 | 0.123945 | -3.30302 | 0.000956 | 0.008003 |
| Mfumv2_1132 | 34.68382319 | 0.413498514 | 0.324345 | 1.274874 | 0.202354 | 0.43339 |
| Mfumv2_1133 | 127.8183649 | 0.16536266 | 0.179463 | 0.921429 | 0.356826 | 0.59836 |
| Mfumv2_1134 | 110.7996952 | 0.284316435 | 0.19717 | 1.441988 | 0.149306 | 0.34557 |
| Mfumv2_1135 | 995.6059162 | -0.545686739 | 0.133238 | -4.09559 | 4.21E-05 | 0.000627 |
| Mfumv2_1136 | 326.1613235 | -0.12154606 | 0.14012 | -0.86744 | 0.385699 | 0.6194 |
| Mfumv2_1137 | 207.2644035 | 0.045082775 | 0.152211 | 0.296186 | 0.767088 | 0.887638 |
| Mfumv2_1138 | 363.6398537 | -0.075575984 | 0.12308 | -0.61404 | 0.539189 | 0.759628 |
| Mfumv2_1139 | 171.076838 | 0.092206658 | 0.158395 | 0.582129 | 0.56048 | 0.77267 |
| Mfumv2_1140 | 59.91837849 | -0.037859789 | 0.247986 | -0.15267 | 0.878659 | 0.945691 |

| | | | | | | |
|---|---|---|---|---|---|---|
| Mfumv2_1141 | 368.9419643 | -0.066893708 | 0.136956 | -0.48843 | 0.625245 | 0.818317 |
| Mfumv2_1142 | 234.61458 | 0.359426789 | 0.142983 | 2.513765 | 0.011945 | 0.058388 |
| Mfumv2_1143 | 57.17077504 | 0.744148422 | 0.284999 | 2.611056 | 0.009026 | 0.047518 |
| Mfumv2_1144 | 8.572551911 | 0.711350675 | 0.667457 | 1.065762 | 0.286531 | NA |
| Mfumv2_1145 | 1.869482486 | 1.090960237 | 1.364017 | 0.799815 | 0.423818 | NA |
| Mfumv2_1146 | 1.987823339 | 0.617866112 | 1.31565 | 0.469628 | 0.638621 | NA |
| Mfumv2_1147 | 6.590500909 | -0.228231907 | 0.673361 | -0.33894 | 0.734652 | NA |
| Mfumv2_1148 | 1137.182512 | -0.316123072 | 0.097211 | -3.25194 | 0.001146 | 0.009248 |
| Mfumv2_1149 | 131.8587126 | -0.098870566 | 0.210054 | -0.47069 | 0.637861 | 0.823866 |
| Mfumv2_1150 | 368.4319784 | -0.395888827 | 0.127969 | -3.09362 | 0.001977 | 0.014369 |
| Mfumv2_1151 | 287.5229313 | -0.169597407 | 0.134664 | -1.25941 | 0.207881 | 0.437957 |
| Mfumv2_1152 | 69.55412591 | -0.049343887 | 0.223098 | -0.22118 | 0.824956 | 0.914645 |
| Mfumv2_1153 | 82.73530847 | -0.193536631 | 0.213575 | -0.90618 | 0.364842 | 0.60367 |
| Mfumv2_1154 | 227.8922894 | -0.318988761 | 0.145549 | -2.19162 | 0.028407 | 0.109329 |
| Mfumv2_1155 | 630.3134427 | -0.1269556 | 0.111495 | -1.13866 | 0.254844 | 0.493715 |
| Mfumv2_1156 | 158.7786405 | 0.361770896 | 0.167761 | 2.15647 | 0.031047 | 0.116586 |
| Mfumv2_1157 | 82.42180666 | 0.327557308 | 0.262658 | 1.247089 | 0.212365 | 0.443494 |
| Mfumv2_1158 | 361.1784664 | 0.351303719 | 0.141528 | 2.482224 | 0.013057 | 0.062689 |
| Mfumv2_1159 | 281.2832713 | -0.096673224 | 0.146456 | -0.66009 | 0.509199 | 0.733977 |
| Mfumv2_1160 | 105.0309294 | 0.206523526 | 0.199323 | 1.036127 | 0.300143 | 0.540311 |
| Mfumv2_1161 | 452.5252934 | 0.372983187 | 0.153431 | 2.430953 | 0.015059 | 0.069709 |
| Mfumv2_1162 | 234.2481445 | 0.319768646 | 0.154233 | 2.073278 | 0.038146 | 0.136699 |
| Mfumv2_1163 | 136.1824937 | 0.14436519 | 0.194099 | 0.743771 | 0.457015 | 0.686205 |
| Mfumv2_1164 | 114.7866324 | 0.045475012 | 0.197637 | 0.230094 | 0.818019 | 0.909463 |
| Mfumv2_1165 | 977.2738132 | -0.195985257 | 0.109907 | -1.78318 | 0.074556 | 0.220827 |
| Mfumv2_1166 | 431.699135 | -0.247354923 | 0.126827 | -1.95034 | 0.051136 | 0.167862 |
| Mfumv2_1167 | 210.887971 | 0.057985783 | 0.171263 | 0.338578 | 0.734928 | 0.872254 |
| Mfumv2_1168 | 108.8316091 | 0.189744893 | 0.185998 | 1.020143 | 0.307661 | 0.549414 |
| Mfumv2_1169 | 199.7842382 | 0.319835908 | 0.154395 | 2.07154 | 0.038308 | 0.136699 |
| Mfumv2_1170 | 408.2383769 | 0.137454194 | 0.128167 | 1.072465 | 0.283511 | 0.523113 |
| Mfumv2_1171 | 112.3762874 | -0.04895782 | 0.201792 | -0.24262 | 0.808303 | 0.907196 |
| Mfumv2_1172 | 93.35870128 | 0.162205273 | 0.206012 | 0.787357 | 0.431073 | 0.664639 |
| Mfumv2_1173 | 42.083192 | 0.25701075 | 0.28929 | 0.88842 | 0.374315 | 0.610542 |
| Mfumv2_1174 | 612.2205276 | -0.54983466 | 0.154764 | -3.55274 | 0.000381 | 0.003948 |
| Mfumv2_1175 | 184.7481039 | -0.258398306 | 0.158308 | -1.63225 | 0.102628 | 0.277869 |
| Mfumv2_1176 | 487.5898528 | -0.265957239 | 0.140749 | -1.88959 | 0.058813 | 0.185197 |
| Mfumv2_1177 | 317.1586592 | 0.084874098 | 0.130774 | 0.649014 | 0.516329 | 0.738821 |
| Mfumv2_1178 | 114.3793785 | -0.2519576 | 0.181928 | -1.38493 | 0.166074 | 0.375723 |
| Mfumv2_1179 | 41.00828186 | 0.238423135 | 0.287715 | 0.828677 | 0.407287 | 0.641254 |
| Mfumv2_1180 | 148.7485596 | 0.328844057 | 0.195918 | 1.678479 | 0.093254 | 0.258242 |
| Mfumv2_1181 | 82.43204038 | 0.649220129 | 0.229385 | 2.830265 | 0.004651 | 0.028314 |
| Mfumv2_1182 | 138.7218397 | 0.139509968 | 0.168977 | 0.825615 | 0.409023 | 0.642476 |
| Mfumv2_1183 | 38002.2232 | -0.499779862 | 0.114299 | -4.37258 | 1.23E-05 | 0.00022 |
| Mfumv2_1184 | 2905.168076 | -0.413292587 | 0.100876 | -4.09702 | 4.19E-05 | 0.000627 |
| Mfumv2_1185 | 1498.216645 | -0.548040482 | 0.130048 | -4.21415 | 2.51E-05 | 0.000403 |
| Mfumv2_1186 | 1404.498978 | -0.265472141 | 0.096097 | -2.76254 | 0.005735 | 0.032641 |
| Mfumv2_1188 | 8.830846039 | -0.12975487 | 0.654107 | -0.19837 | 0.842756 | NA |
| Mfumv2_1189 | 8.1109916 | -0.842645922 | 0.738176 | -1.14152 | 0.253652 | NA |
| Mfumv2_1191 | 6137.01656 | -0.605383379 | 0.111658 | -5.42177 | 5.90E-08 | 1.65E-06 |
| Mfumv2_1192 | 1067.440328 | -0.137549306 | 0.099991 | -1.37562 | 0.16894 | 0.380068 |
| Mfumv2_1193 | 1670.467485 | -0.235652939 | 0.116506 | -2.02266 | 0.043108 | 0.147788 |
| Mfumv2_1195 | 323.5206386 | 0.137527185 | 0.136707 | 1.005997 | 0.314417 | 0.557023 |

| | | | | | | |
|---|---|---|---|---|---|---|
| Mfumv2_1196 | 1208.801688 | -0.270772403 | 0.138909 | -1.94928 | 0.051262 | 0.168001 |
| Mfumv2_1197 | 2164.69934 | -0.201545331 | 0.108464 | -1.85818 | 0.063143 | 0.194862 |
| Mfumv2_1198 | 259.0937727 | 0.181160293 | 0.145543 | 1.24472 | 0.213235 | 0.444386 |
| Mfumv2_1199 | 35.96628252 | 0.606483154 | 0.324152 | 1.870982 | 0.061348 | 0.191378 |
| Mfumv2_1200 | 7.040359675 | 1.288077266 | 0.708672 | 1.817593 | 0.069126 | NA |
| Mfumv2_1201 | 3.850198038 | 0.319310511 | 0.899017 | 0.355177 | 0.722457 | NA |
| Mfumv2_1202 | 1.917697909 | -0.61700323 | 1.240229 | -0.49749 | 0.618843 | NA |
| Mfumv2_1203 | 21.99385937 | 0.866323323 | 0.424436 | 2.041115 | 0.041239 | NA |
| Mfumv2_1204 | 195.3806305 | 0.068159279 | 0.157085 | 0.433901 | 0.66436 | 0.835454 |
| Mfumv2_1205 | 1130.407119 | -0.303501519 | 0.13118 | -2.31362 | 0.020689 | 0.088811 |
| Mfumv2_1206 | 1883.808148 | -0.299648484 | 0.118701 | -2.52439 | 0.01159 | 0.05679 |
| Mfumv2_1207 | 742.0041302 | -0.309723715 | 0.14192 | -2.18239 | 0.029081 | 0.111496 |
| Mfumv2_1208 | 446.5713569 | -0.166851011 | 0.16454 | -1.01405 | 0.31056 | 0.55222 |
| Mfumv2_1209 | 590.696018 | -0.380270027 | 0.174834 | -2.17503 | 0.029628 | 0.112732 |
| Mfumv2_1210 | 1317.796937 | -0.416881895 | 0.139601 | -2.98625 | 0.002824 | 0.019095 |
| Mfumv2_1211 | 305.7576891 | -0.041450593 | 0.149891 | -0.27654 | 0.782134 | 0.895332 |
| Mfumv2_1212 | 0.140690113 | 0.283845181 | 4.080473 | 0.069562 | 0.944542 | NA |
| Mfumv2_1213 | 8.258555786 | 0.78010119 | 0.623372 | 1.251423 | 0.21078 | NA |
| Mfumv2_1214 | 905.663652 | -0.248601344 | 0.109941 | -2.26123 | 0.023745 | 0.09769 |
| Mfumv2_1215 | 75.27919103 | -0.252052867 | 0.216034 | -1.16673 | 0.24332 | 0.48259 |
| Mfumv2_1216 | 2.12674707 | 1.351853785 | 1.32468 | 1.020514 | 0.307485 | NA |
| Mfumv2_1217 | 29.94913597 | 0.044004497 | 0.399646 | 0.110109 | 0.912323 | 0.963869 |
| Mfumv2_1218 | 571.1922267 | 0.050145383 | 0.126204 | 0.397337 | 0.691119 | 0.847961 |
| Mfumv2_1219 | 1305.543462 | -0.015013665 | 0.117814 | -0.12744 | 0.898596 | 0.954165 |
| Mfumv2_1221 | 173.253712 | 0.51100754 | 0.170932 | 2.98953 | 0.002794 | 0.019028 |
| Mfumv2_1222 | 141.9897541 | 0.57538629 | 0.173609 | 3.314258 | 0.000919 | 0.007789 |
| Mfumv2_1223 | 460.2080232 | 0.124515862 | 0.140126 | 0.8886 | 0.374218 | 0.610542 |
| Mfumv2_1225 | 198.3952526 | 0.507873021 | 0.164149 | 3.093972 | 0.001975 | 0.014369 |
| Mfumv2_1226 | 371.401567 | -0.231515958 | 0.149899 | -1.54448 | 0.122473 | 0.30641 |
| Mfumv2_1227 | 2217.08227 | -0.09931116 | 0.131537 | -0.755 | 0.450246 | 0.679597 |
| Mfumv2_1228 | 69.6982094 | 0.468720549 | 0.226544 | 2.069005 | 0.038546 | 0.137302 |
| Mfumv2_1229 | 160.3802888 | 0.614192979 | 0.183574 | 3.345744 | 0.000821 | 0.007076 |
| Mfumv2_1230 | 88.62781073 | -0.004202892 | 0.202495 | -0.02076 | 0.983441 | 0.992996 |
| Mfumv2_1231 | 577.1063293 | -0.028028295 | 0.113264 | -0.24746 | 0.804552 | 0.904502 |
| Mfumv2_1232 | 1647.802715 | -0.050898139 | 0.105499 | -0.48245 | 0.629486 | 0.82074 |
| Mfumv2_1233 | 3937.681358 | -0.418918094 | 0.127879 | -3.2759 | 0.001053 | 0.008672 |
| Mfumv2_1234 | 193.343796 | -0.425690691 | 0.16841 | -2.52771 | 0.011481 | 0.056532 |
| Mfumv2_1235 | 86.38504118 | 0.206121007 | 0.205885 | 1.001147 | 0.316756 | 0.559685 |
| Mfumv2_1236 | 120.6852734 | 0.137445666 | 0.184978 | 0.743039 | 0.457458 | 0.686358 |
| Mfumv2_1237 | 13.35424046 | 0.735642826 | 0.511413 | 1.438452 | 0.150306 | NA |
| Mfumv2_1238 | 592.5356097 | 0.712588097 | 0.161986 | 4.399064 | 1.09E-05 | 0.000197 |
| Mfumv2_1239 | 1545.157125 | 0.185904582 | 0.119407 | 1.556895 | 0.119495 | 0.304013 |
| Mfumv2_1240 | 998.9975624 | 0.140593948 | 0.11683 | 1.20341 | 0.228818 | 0.465042 |
| Mfumv2_1241 | 998.5413236 | 0.265207572 | 0.116133 | 2.283655 | 0.022392 | 0.09432 |
| Mfumv2_1242 | 224.1731739 | 0.516435857 | 0.162423 | 3.179568 | 0.001475 | 0.011353 |
| Mfumv2_1243 | 469.340891 | -0.016821373 | 0.185603 | -0.09063 | 0.927786 | 0.972652 |
| Mfumv2_1244 | 410.3965108 | -0.872980915 | 0.655761 | -1.33125 | 0.183107 | 0.403743 |
| Mfumv2_1245 | 256.9904953 | -0.607035403 | 0.211109 | -2.87547 | 0.004034 | 0.025407 |
| Mfumv2_1246 | 380.2518293 | -0.003720195 | 0.168646 | -0.02206 | 0.982401 | 0.992996 |
| Mfumv2_1247 | 1867.643396 | 0.231315631 | 0.101294 | 2.283607 | 0.022395 | 0.09432 |
| Mfumv2_1248 | 422.7879761 | 0.26853315 | 0.133411 | 2.012827 | 0.044133 | 0.149634 |
| Mfumv2_1249 | 206.0618045 | 0.418097911 | 0.152691 | 2.738194 | 0.006178 | 0.034863 |

| | | | | | | | |
|---|---|---|---|---|---|---|---|
| Mfumv2_1250 | 23.09307055 | | 0.085869841 | 0.463 | 0.185464 | 0.852865 | NA |
| Mfumv2_1251 | 7.966644957 | | 0.452841302 | 0.633876 | 0.7144 | 0.47498 | NA |
| Mfumv2_1252 | 98.37124975 | | 0.791985016 | 0.207134 | 3.823537 | 0.000132 | 0.001592 |
| Mfumv2_1253 | 332.2027975 | | 0.176275319 | 0.139141 | 1.266883 | 0.205197 | 0.434665 |
| Mfumv2_1254 | 0 | NA | | NA | NA | NA | NA |
| Mfumv2_1255 | 778.9184758 | | 0.628093899 | 0.110691 | 5.674315 | 1.39E-08 | 4.37E-07 |
| Mfumv2_1256 | 6477.196687 | | -0.058278679 | 0.103132 | -0.56509 | 0.572015 | 0.781754 |
| Mfumv2_1257 | 456.6321988 | | 2.160425732 | 0.156439 | 13.81002 | 2.22E-43 | 4.46E-41 |
| Mfumv2_1258 | 157.9876092 | | 2.444800767 | 0.209679 | 11.65973 | 2.05E-31 | 3.16E-29 |
| Mfumv2_1259 | 549.6227584 | | 2.385955267 | 0.177361 | 13.45251 | 2.98E-41 | 5.44E-39 |
| Mfumv2_1260 | 1463.842205 | | 2.157673289 | 0.145082 | 14.87205 | 5.01E-50 | 1.26E-47 |
| Mfumv2_1261 | 55.78010269 | | 1.661345737 | 0.315475 | 5.266175 | 1.39E-07 | 3.59E-06 |
| Mfumv2_1262 | 743.7048875 | | -0.450856518 | 0.125636 | -3.5886 | 0.000332 | 0.003515 |
| Mfumv2_1263 | 1.861713393 | | -0.12530436 | 1.36695 | -0.09167 | 0.926963 | NA |
| Mfumv2_1264 | 2.01505872 | | 0.71399145 | 1.262726 | 0.565436 | 0.571777 | NA |
| Mfumv2_1265 | 0.373839054 | | 1.700247813 | 2.80949 | 0.60518 | 0.545059 | NA |
| Mfumv2_1266 | 8.229751873 | | 0.830835285 | 0.701295 | 1.184717 | 0.23613 | NA |
| Mfumv2_1267 | 69.15660508 | | 0.61646163 | 0.25384 | 2.428543 | 0.01516 | 0.070013 |
| Mfumv2_1268 | 133.7747113 | | 0.555857116 | 0.175158 | 3.173465 | 0.001506 | 0.01147 |
| Mfumv2_1269 | 31.70163709 | | 1.434793223 | 0.398824 | 3.597557 | 0.000321 | 0.003433 |
| Mfumv2_1270 | 473.0853331 | | 0.837845603 | 0.133778 | 6.262954 | 3.78E-10 | 1.72E-08 |
| Mfumv2_1271 | 315.5472971 | | -0.24048453 | 0.154528 | -1.55626 | 0.119647 | 0.304013 |
| Mfumv2_1272 | 214.4051609 | | 0.601920042 | 0.149475 | 4.026894 | 5.65E-05 | 0.000794 |
| Mfumv2_1273 | 564.6673868 | | -0.38271148 | 0.144314 | -2.65194 | 0.008003 | 0.043221 |
| Mfumv2_1274 | 967.037493 | | -0.070530833 | 0.144372 | -0.48853 | 0.625171 | 0.818317 |
| Mfumv2_1275 | 624.7675959 | | -1.092000832 | 0.627887 | -1.73917 | 0.082005 | 0.234775 |
| Mfumv2_1276 | 403.796385 | | 0.295036864 | 0.136592 | 2.159988 | 0.030774 | 0.115993 |
| Mfumv2_1277 | 157.5672048 | | 0.158341154 | 0.197238 | 0.802791 | 0.422096 | 0.655325 |
| Mfumv2_1278 | 107.6644816 | | 0.684177247 | 0.216821 | 3.155486 | 0.001602 | 0.012011 |
| Mfumv2_1279 | 724.2532173 | | 0.653223108 | 0.119462 | 5.468035 | 4.55E-08 | 1.32E-06 |
| Mfumv2_1280 | 19.16205913 | | 0.552147601 | 0.423211 | 1.304664 | 0.192007 | NA |
| Mfumv2_1281 | 1796.27132 | | 0.13615943 | 0.14091 | 0.966286 | 0.333901 | 0.57679 |
| Mfumv2_1282 | 712.5923144 | | 0.029060808 | 0.138337 | 0.210073 | 0.833611 | 0.920178 |
| Mfumv2_1283 | 722.8858654 | | 0.242452033 | 0.1165 | 2.081141 | 0.037421 | 0.135457 |
| Mfumv2_1284 | 295.54189 | | 0.082064423 | 0.154982 | 0.529509 | 0.596453 | 0.798317 |
| Mfumv2_1285 | 230.5078823 | | 0.059607771 | 0.149065 | 0.399877 | 0.689247 | 0.846498 |
| Mfumv2_1286 | 162.6022346 | | 0.497845723 | 0.179244 | 2.77747 | 0.005478 | 0.032118 |
| Mfumv2_1287 | 188.0211785 | | 0.402418425 | 0.192108 | 2.094754 | 0.036193 | 0.131248 |
| Mfumv2_1288 | 392.7328722 | | -0.167391181 | 0.160389 | -1.04366 | 0.296643 | 0.53545 |
| Mfumv2_1289 | 322.9426667 | | 1.618943603 | 0.153385 | 10.55478 | 4.83E-26 | 5.71E-24 |
| Mfumv2_1290 | 86.5175 | | -0.050320805 | 0.230285 | -0.21852 | 0.827028 | 0.915426 |
| Mfumv2_1291 | 102.742597 | | 0.627567981 | 0.198444 | 3.162436 | 0.001565 | 0.011772 |
| Mfumv2_1292 | 213.7877786 | | 0.583985798 | 0.157037 | 3.718776 | 0.0002 | 0.002272 |
| Mfumv2_1293 | 112.124146 | | 0.704176741 | 0.194186 | 3.626298 | 0.000288 | 0.003089 |
| Mfumv2_1294 | 41.10882627 | | 0.452135454 | 0.28405 | 1.591748 | 0.111441 | 0.291291 |
| Mfumv2_1295 | 10.9964597 | | 0.855333059 | 0.550952 | 1.552463 | 0.120551 | NA |
| Mfumv2_1296 | 40.30409584 | | 0.476528184 | 0.299286 | 1.592218 | 0.111336 | 0.291291 |
| Mfumv2_1297 | 28.60993626 | | 1.389140347 | 0.361981 | 3.837605 | 0.000124 | 0.00155 |
| Mfumv2_1299 | 25.46386196 | | 0.013433424 | 0.420004 | 0.031984 | 0.974485 | 0.990165 |
| Mfumv2_1300 | 114.0007077 | | 0.409185345 | 0.199378 | 2.052312 | 0.040139 | 0.140243 |
| Mfumv2_1301 | 465.1296479 | | 0.401460468 | 0.14457 | 2.776933 | 0.005487 | 0.032118 |
| Mfumv2_1302 | 156.3863659 | | 0.574265241 | 0.168543 | 3.407233 | 0.000656 | 0.00602 |

| | | | | | | | |
|---|---|---|---|---|---|---|---|
| Mfumv2_1303 | 387.4776757 | | 0.26773277 | 0.171007 | 1.565623 | 0.117437 | 0.300167 |
| Mfumv2_1305 | 191.5101674 | | -0.481795151 | 0.556272 | -0.86611 | 0.386428 | 0.620075 |
| Mfumv2_1306 | 490.1213659 | | 0.333163223 | 0.124061 | 2.685478 | 0.007243 | 0.040084 |
| Mfumv2_1307 | 107.0595032 | | 0.516390766 | 0.202977 | 2.544083 | 0.010956 | 0.054771 |
| Mfumv2_1310 | 689.1865318 | | -0.286669358 | 0.149145 | -1.92208 | 0.054595 | 0.175082 |
| Mfumv2_1311 | 43.85938089 | | 0.838937014 | 0.29451 | 2.848587 | 0.004391 | 0.027229 |
| Mfumv2_1312 | 0.938246609 | | 0.202414507 | 1.972481 | 0.102619 | 0.918265 | NA |
| Mfumv2_1313 | 18.97084768 | | 0.100562959 | 0.439888 | 0.228611 | 0.819172 | NA |
| Mfumv2_1314 | 6.893837447 | | -1.313376336 | 0.742664 | -1.76847 | 0.076983 | NA |
| Mfumv2_1315 | 3.352167627 | | 0.512613492 | 1.008764 | 0.50816 | 0.611341 | NA |
| Mfumv2_1316 | 139.2120291 | | 0.644815327 | 0.176141 | 3.660793 | 0.000251 | 0.002757 |
| Mfumv2_1317 | 651.3187052 | | 0.197004302 | 0.134995 | 1.459342 | 0.144471 | 0.337099 |
| Mfumv2_1318 | 908.6578665 | | 0.452652957 | 0.109914 | 4.118235 | 3.82E-05 | 0.000599 |
| Mfumv2_1319 | 90.88085579 | | 0.276197823 | 0.207726 | 1.329625 | 0.183642 | 0.403743 |
| Mfumv2_1320 | 158.2513547 | | 0.384411223 | 0.173456 | 2.216191 | 0.026678 | 0.104886 |
| Mfumv2_1321 | 132.5289837 | | 0.113117635 | 0.198944 | 0.56859 | 0.569635 | 0.779561 |
| Mfumv2_1322 | 50.14151573 | | -0.039492274 | 0.276367 | -0.1429 | 0.886371 | 0.947191 |
| Mfumv2_1323 | 10.05327952 | | 1.281915109 | 0.637415 | 2.011117 | 0.044313 | NA |
| Mfumv2_1324 | 7.75541621 | | 0.180118742 | 0.627897 | 0.28686 | 0.774219 | NA |
| Mfumv2_1325 | 20.21696859 | | 0.119041666 | 0.444748 | 0.267661 | 0.78896 | NA |
| Mfumv2_1326 | 2.296423085 | | -0.418116143 | 1.200781 | -0.3482 | 0.727687 | NA |
| Mfumv2_1327 | 1.129787016 | | -0.324810311 | 1.63723 | -0.19839 | 0.84274 | NA |
| Mfumv2_1328 | 0.393330567 | | -2.461665841 | 2.808251 | -0.87658 | 0.380713 | NA |
| Mfumv2_1329 | 0 | NA | | NA | NA | NA | NA |
| Mfumv2_1330 | 0.397954697 | | 1.772791437 | 2.756505 | 0.64313 | 0.52014 | NA |
| Mfumv2_1331 | 337.7236216 | | 0.700660924 | 0.142914 | 4.902674 | 9.45E-07 | 2.04E-05 |
| Mfumv2_1332 | 837.3621214 | | 0.301847199 | 0.125897 | 2.397564 | 0.016504 | 0.074323 |
| Mfumv2_1333 | 1130.617786 | | -0.041017219 | 0.151047 | -0.27155 | 0.785966 | 0.89568 |
| Mfumv2_1334 | 1408.024979 | | 0.208972998 | 0.115883 | 1.803311 | 0.071339 | 0.214874 |
| Mfumv2_1335 | 19.04607907 | | 0.705968487 | 0.439067 | 1.607882 | 0.107861 | NA |
| Mfumv2_1336 | 13.98854141 | | 0.593148598 | 0.485396 | 1.221988 | 0.221712 | NA |
| Mfumv2_1337 | 2.235616754 | | 1.569403698 | 1.323541 | 1.185762 | 0.235716 | NA |
| Mfumv2_1338 | 0.514529168 | | 2.151912868 | 2.498393 | 0.861319 | 0.389062 | NA |
| Mfumv2_1339 | 18.21102832 | | 0.230897273 | 0.420963 | 0.548498 | 0.58335 | NA |
| Mfumv2_1340 | 617.5774182 | | -0.001793059 | 0.120117 | -0.01493 | 0.98809 | 0.994355 |
| Mfumv2_1341 | 665.3303845 | | -0.337726928 | 0.151859 | -2.22395 | 0.026152 | 0.103325 |
| Mfumv2_1342 | 191.5704996 | | 0.619819024 | 0.187711 | 3.301986 | 0.00096 | 0.008003 |
| Mfumv2_1343 | 393.6208143 | | 0.379455032 | 0.133536 | 2.841604 | 0.004489 | 0.027511 |
| Mfumv2_1344 | 751.491931 | | 0.228894284 | 0.110878 | 2.064372 | 0.038982 | 0.13788 |
| Mfumv2_1345 | 2479.944955 | | -0.070879101 | 0.112375 | -0.63074 | 0.528214 | 0.750482 |
| Mfumv2_1346 | 628.0542629 | | 0.359378666 | 0.117801 | 3.050729 | 0.002283 | 0.01632 |
| Mfumv2_1347 | 1.20669377 | | -0.353299135 | 1.59604 | -0.22136 | 0.824812 | NA |
| Mfumv2_1348 | 132.7023776 | | 1.037882136 | 0.18334 | 5.660964 | 1.51E-08 | 4.65E-07 |
| Mfumv2_1349 | 153.236996 | | 0.40009193 | 0.176044 | 2.272677 | 0.023046 | 0.096467 |
| Mfumv2_1350 | 95.88473745 | | 0.742992644 | 0.218503 | 3.400377 | 0.000673 | 0.006145 |
| Mfumv2_1351 | 740.8115943 | | 0.008826117 | 0.126091 | 0.069998 | 0.944195 | 0.979292 |
| Mfumv2_1352 | 895.8732061 | | 0.25508383 | 0.109758 | 2.324056 | 0.020122 | 0.086751 |
| Mfumv2_1353 | 575.8667737 | | 0.059329864 | 0.123455 | 0.480577 | 0.630817 | 0.82074 |
| Mfumv2_1354 | 176.276285 | | -0.15395051 | 0.188752 | -0.81562 | 0.414716 | 0.647422 |
| Mfumv2_1355 | 1240.009085 | | -0.603782704 | 0.129829 | -4.6506 | 3.31E-06 | 6.72E-05 |
| Mfumv2_1357 | 3925.699697 | | -0.994427009 | 0.106608 | -9.32785 | 1.08E-20 | 9.44E-19 |
| Mfumv2_1358 | 211.9133668 | | 0.265614187 | 0.154335 | 1.721024 | 0.085247 | 0.24181 |

| | | | | | | | |
|---|---|---|---|---|---|---|---|
| Mfumv2_1359 | 798.425512 | | 0.303754932 | 0.13434 | 2.261096 | 0.023753 | 0.09769 |
| Mfumv2_1360 | 189.4306011 | | 0.083844543 | 0.161646 | 0.518691 | 0.603976 | 0.802869 |
| Mfumv2_1361 | 2102.819546 | | -0.262974551 | 0.121442 | -2.16544 | 0.030354 | 0.114843 |
| Mfumv2_1364 | 15.96907009 | | 0.503164178 | 0.517423 | 0.972443 | 0.33083 | NA |
| Mfumv2_1365 | 15.38000408 | | 0.182482563 | 0.461448 | 0.395457 | 0.692506 | NA |
| Mfumv2_1366 | 2.176854866 | | 0.570613934 | 1.318618 | 0.432736 | 0.665206 | NA |
| Mfumv2_1367 | 0 | NA | | NA | NA | NA | NA |
| Mfumv2_1368 | 23.84457252 | | 0.120758129 | 0.371745 | 0.324841 | 0.745302 | 0.878227 |
| Mfumv2_1369 | 41.46406506 | | 0.686478548 | 0.300182 | 2.286872 | 0.022203 | 0.094106 |
| Mfumv2_1370 | 30.89881178 | | 0.301033472 | 0.330778 | 0.910077 | 0.362782 | 0.602338 |
| Mfumv2_1371 | 20.78958001 | | 0.461049281 | 0.390355 | 1.181102 | 0.237562 | NA |
| Mfumv2_1372 | 22.02257201 | | 0.426108495 | 0.392363 | 1.086006 | 0.277476 | NA |
| Mfumv2_1373 | 18.75807895 | | 0.173372882 | 0.42748 | 0.405569 | 0.685059 | NA |
| Mfumv2_1374 | 17.67773263 | | 0.334152057 | 0.446085 | 0.749077 | 0.453811 | NA |
| Mfumv2_1375 | 47.91101048 | | 0.033579031 | 0.278461 | 0.120588 | 0.904017 | 0.957896 |
| Mfumv2_1376 | 94.58341553 | | 0.180541921 | 0.2028 | 0.890245 | 0.373334 | 0.610542 |
| Mfumv2_1377 | 41.55773648 | | 0.460641169 | 0.29106 | 1.582636 | 0.113505 | 0.294233 |
| Mfumv2_1378 | 112.4900941 | | 0.695038188 | 0.199312 | 3.487187 | 0.000488 | 0.004807 |
| Mfumv2_1379 | 23.61672872 | | 1.036433341 | 0.395336 | 2.621653 | 0.00875 | 0.046507 |
| Mfumv2_1380 | 2.787504447 | | 0.118522212 | 1.131308 | 0.104766 | 0.916562 | NA |
| Mfumv2_1381 | 51.21711746 | | 0.244114847 | 0.264608 | 0.922553 | 0.35624 | 0.59836 |
| Mfumv2_1382 | 44.29622892 | | 0.117318811 | 0.276221 | 0.424728 | 0.671035 | 0.838377 |
| Mfumv2_1383 | 90.30996043 | | -0.013736525 | 0.204877 | -0.06705 | 0.946544 | 0.979704 |
| Mfumv2_1384 | 0.415456949 | | -2.513930232 | 2.758893 | -0.91121 | 0.362185 | NA |
| Mfumv2_1386 | 183.2190391 | | 0.064396726 | 0.163348 | 0.39423 | 0.693411 | 0.849183 |
| Mfumv2_1387 | 40.1893587 | | 0.488777262 | 0.303505 | 1.61044 | 0.107302 | 0.284392 |
| Mfumv2_1388 | 3.837074645 | | 0.437657127 | 0.917626 | 0.476945 | 0.633401 | NA |
| Mfumv2_1389 | 11.20469202 | | 0.490593188 | 0.540869 | 0.907046 | 0.364383 | NA |
| Mfumv2_1390 | 58.6156202 | | 0.166653465 | 0.247173 | 0.674237 | 0.500161 | 0.727605 |
| Mfumv2_1391 | 16.53722571 | | 0.034843297 | 0.447122 | 0.077928 | 0.937885 | NA |
| Mfumv2_1392 | 20.63146867 | | 0.315143516 | 0.393608 | 0.800652 | 0.423333 | NA |
| Mfumv2_1393 | 76.28591674 | | 0.496628011 | 0.22624 | 2.195137 | 0.028154 | 0.108981 |
| Mfumv2_1394 | 80.52739351 | | 0.201263087 | 0.219382 | 0.917409 | 0.358928 | 0.598398 |
| Mfumv2_1395 | 124.4201212 | | -0.13662883 | 0.189063 | -0.72266 | 0.469888 | 0.697712 |
| Mfumv2_1396 | 55.44949184 | | -0.127904586 | 0.266966 | -0.4791 | 0.631865 | 0.82074 |
| Mfumv2_1397 | 64.35024108 | | 0.130871757 | 0.23918 | 0.547168 | 0.584263 | 0.790427 |
| Mfumv2_1398 | 4.755313148 | | -0.084883489 | 0.798785 | -0.10627 | 0.915372 | NA |
| Mfumv2_1399 | 12.71126893 | | 0.192532921 | 0.500178 | 0.384929 | 0.70029 | NA |
| Mfumv2_1400 | 31.42089828 | | 0.206649886 | 0.331731 | 0.622943 | 0.533322 | 0.755069 |
| Mfumv2_1401 | 22.06056772 | | -0.147265575 | 0.380714 | -0.38681 | 0.698894 | NA |
| Mfumv2_1402 | 111.3254136 | | 0.541983807 | 0.193495 | 2.801028 | 0.005094 | 0.030368 |
| Mfumv2_1403 | 34.60244822 | | -0.022663948 | 0.305001 | -0.07431 | 0.940766 | 0.977742 |
| Mfumv2_1404 | 88.05956384 | | 0.377440398 | 0.237415 | 1.589792 | 0.111882 | 0.291291 |
| Mfumv2_1405 | 103.6681568 | | 0.231697873 | 0.202977 | 1.1415 | 0.253662 | 0.493394 |
| Mfumv2_1406 | 58.68485194 | | 0.611081387 | 0.260639 | 2.344554 | 0.01905 | 0.083744 |
| Mfumv2_1407 | 8.855595102 | | 0.875882368 | 0.635714 | 1.377793 | 0.168267 | NA |
| Mfumv2_1409 | 12.7839971 | | 0.630754226 | 0.528059 | 1.194477 | 0.232291 | NA |
| Mfumv2_1410 | 11.60894303 | | -0.540212639 | 0.507447 | -1.06457 | 0.287071 | NA |
| Mfumv2_1411 | 152.4150959 | | 0.205510263 | 0.180657 | 1.137572 | 0.255299 | 0.493892 |
| Mfumv2_1412 | 99.01931012 | | 0.188090704 | 0.192925 | 0.97494 | 0.32959 | 0.572296 |
| Mfumv2_1413 | 31.27753018 | | 0.244448614 | 0.348626 | 0.701177 | 0.483193 | 0.711161 |
| Mfumv2_1415 | 90.28815445 | | 0.817475452 | 0.222667 | 3.671287 | 0.000241 | 0.002664 |

| | | | | | | |
|---|---|---|---|---|---|---|
| Mfumv2_1416 | 1364.931772 | -0.060483892 | 0.093895 | -0.64416 | 0.51947 | 0.741731 |
| Mfumv2_1417 | 515.2872721 | -0.090074786 | 0.123011 | -0.73225 | 0.464014 | 0.692574 |
| Mfumv2_1418 | 0.642564068 | 0.861621498 | 2.248468 | 0.383204 | 0.701569 | NA |
| Mfumv2_1419 | 966.4258941 | -0.431212522 | 0.112118 | -3.84607 | 0.00012 | 0.001517 |
| Mfumv2_1420 | 3445.892202 | -0.473554271 | 0.092916 | -5.09661 | 3.46E-07 | 7.89E-06 |
| Mfumv2_1423 | 246.5482304 | 0.207877204 | 0.142164 | 1.462237 | 0.143676 | 0.336025 |
| Mfumv2_1424 | 49.50208901 | 0.227377863 | 0.267771 | 0.849151 | 0.395797 | 0.627515 |
| Mfumv2_1425 | 331.3389544 | 0.209837372 | 0.14836 | 1.414381 | 0.15725 | 0.360223 |
| Mfumv2_1426 | 542.6949356 | -0.091931476 | 0.122146 | -0.75263 | 0.451669 | 0.680331 |
| Mfumv2_1427 | 319.6650635 | 0.134957359 | 0.144115 | 0.936453 | 0.34904 | 0.592749 |
| Mfumv2_1428 | 326.1063926 | 0.044726356 | 0.14515 | 0.308139 | 0.757977 | 0.884498 |
| Mfumv2_1429 | 198.5488915 | 0.857297543 | 0.164863 | 5.200074 | 1.99E-07 | 4.94E-06 |
| Mfumv2_1430 | 1305.942841 | -0.130845225 | 0.098803 | -1.3243 | 0.185403 | 0.40507 |
| Mfumv2_1431 | 631.9541571 | -0.324315663 | 0.134029 | -2.41975 | 0.015531 | 0.071237 |
| Mfumv2_1432 | 198.7570701 | -0.039644934 | 0.150165 | -0.26401 | 0.791773 | 0.897706 |
| Mfumv2_1433 | 1043.568535 | 0.083078729 | 0.107728 | 0.771187 | 0.440596 | 0.670574 |
| Mfumv2_1434 | 404.9025385 | -0.048313842 | 0.141446 | -0.34157 | 0.732673 | 0.872002 |
| Mfumv2_1435 | 2.755503052 | -0.411311089 | 1.059559 | -0.38819 | 0.697875 | NA |
| Mfumv2_1436 | 4.302868966 | 1.263025023 | 0.911379 | 1.385839 | 0.165796 | NA |
| Mfumv2_1437 | 8.745730291 | 0.72526578 | 0.601363 | 1.206037 | 0.227803 | NA |
| Mfumv2_1438 | 0.255553176 | 1.14483936 | 3.242772 | 0.353043 | 0.724056 | NA |
| Mfumv2_1439 | 0.661499874 | -3.162282258 | 2.353633 | -1.34357 | 0.179086 | NA |
| Mfumv2_1440 | 0.138978706 | 0.283845181 | 4.080473 | 0.069562 | 0.944542 | NA |
| Mfumv2_1441 | 0.618496868 | -0.440505629 | 2.29634 | -0.19183 | 0.847876 | NA |
| Mfumv2_1442 | 0 | NA | NA | NA | NA | NA |
| Mfumv2_1443 | 0 | NA | NA | NA | NA | NA |
| Mfumv2_1444 | 4.450388847 | 0.492059545 | 0.823137 | 0.597786 | 0.549983 | NA |
| Mfumv2_1445 | 8.536497335 | 0.448138651 | 0.609372 | 0.73541 | 0.46209 | NA |
| Mfumv2_1446 | 0.947488369 | -2.551971814 | 1.999852 | -1.27608 | 0.201927 | NA |
| Mfumv2_1447 | 0.246042925 | -1.639711413 | 3.689798 | -0.44439 | 0.65676 | NA |
| Mfumv2_1448 | 10.04593483 | 0.094894848 | 0.552489 | 0.171759 | 0.863627 | NA |
| Mfumv2_1449 | 228.1344272 | 0.033669748 | 0.149897 | 0.22462 | 0.822275 | 0.913574 |
| Mfumv2_1450 | 329.5065233 | 0.47854095 | 0.157034 | 3.04737 | 0.002309 | 0.01632 |
| Mfumv2_1451 | 721.9032171 | -0.131236964 | 0.119495 | -1.09826 | 0.272091 | 0.511349 |
| Mfumv2_1452 | 151.7122169 | 0.368751049 | 0.189919 | 1.941626 | 0.052182 | 0.169635 |
| Mfumv2_1453 | 104.3062999 | 0.424226273 | 0.205912 | 2.060234 | 0.039376 | 0.138299 |
| Mfumv2_1455 | 429.9376389 | -0.074951823 | 0.148285 | -0.50546 | 0.613236 | 0.808872 |
| Mfumv2_1456 | 198.9180023 | -0.326997764 | 0.154693 | -2.11385 | 0.034528 | 0.126583 |
| Mfumv2_1457 | 3580.44967 | -0.340836057 | 0.10426 | -3.26908 | 0.001079 | 0.008776 |
| Mfumv2_1458 | 1436.069294 | -0.482961027 | 0.10004 | -4.82768 | 1.38E-06 | 2.93E-05 |
| Mfumv2_1459 | 461.0017426 | -0.525467846 | 0.138099 | -3.80501 | 0.000142 | 0.001666 |
| Mfumv2_1460 | 208.571382 | 0.1767541 | 0.16504 | 1.07098 | 0.284178 | 0.523113 |
| Mfumv2_1461 | 1518.83733 | -0.150635491 | 0.102717 | -1.46651 | 0.14251 | 0.334337 |
| Mfumv2_1462 | 444.8645005 | -0.424116461 | 0.139672 | -3.03651 | 0.002393 | 0.016695 |
| Mfumv2_1463 | 49.22675847 | -0.654442108 | 0.284782 | -2.29805 | 0.021559 | 0.091763 |
| Mfumv2_1464 | 1142.729022 | -1.05888868 | 0.112375 | -9.42284 | 4.39E-21 | 4.41E-19 |
| Mfumv2_1465 | 2786.381328 | -0.39397291 | 0.114595 | -3.43795 | 0.000586 | 0.005477 |
| Mfumv2_1466 | 2331.578566 | -0.19132313 | 0.09802 | -1.95188 | 0.050953 | 0.167809 |
| Mfumv2_1467 | 1465.565735 | -0.174296912 | 0.096141 | -1.81293 | 0.069843 | 0.211318 |
| Mfumv2_1468 | 216.1559263 | 0.023089808 | 0.158627 | 0.14556 | 0.884269 | 0.946455 |
| Mfumv2_1469 | 79.46878734 | 0.32772251 | 0.216718 | 1.512209 | 0.130481 | 0.318384 |
| Mfumv2_1470 | 1146.822469 | -0.363725732 | 0.109363 | -3.32585 | 0.000881 | 0.007536 |

| | | | | | | |
|---|---|---|---|---|---|---|
| Mfumv2_1471 | 36.19080657 | 0.453621467 | 0.311184 | 1.457729 | 0.144915 | 0.337743 |
| Mfumv2_1472 | 24.49303062 | 0.320734504 | 0.384173 | 0.83487 | 0.403791 | 0.637247 |
| Mfumv2_1473 | 3.01594235 | -0.419960032 | 1.036469 | -0.40518 | 0.685343 | NA |
| Mfumv2_1474 | 17.3881323 | -0.349461448 | 0.427458 | -0.81753 | 0.413623 | NA |
| Mfumv2_1475 | 0 | NA | NA | NA | NA | NA |
| Mfumv2_1476 | 0.843614726 | 1.56387336 | 1.945553 | 0.803819 | 0.421501 | NA |
| Mfumv2_1478 | 153.5376149 | -0.062407548 | 0.16554 | -0.37699 | 0.706178 | 0.855161 |
| Mfumv2_1479 | 271.4729892 | 0.040299923 | 0.135647 | 0.297093 | 0.766395 | 0.887428 |
| Mfumv2_1480 | 0 | NA | NA | NA | NA | NA |
| Mfumv2_1481 | 780.2985213 | -0.569213068 | 0.143018 | -3.98002 | 6.89E-05 | 0.000935 |
| Mfumv2_1482 | 685.1830633 | -0.52443071 | 0.153812 | -3.40956 | 0.000651 | 0.006014 |
| Mfumv2_1483 | 495.5741201 | -0.438889091 | 0.144481 | -3.0377 | 0.002384 | 0.016687 |
| Mfumv2_1484 | 404.0706852 | -0.495307607 | 0.129167 | -3.83462 | 0.000126 | 0.00156 |
| Mfumv2_1485 | 2255.303113 | -0.276067509 | 0.132777 | -2.07918 | 0.037601 | 0.13562 |
| Mfumv2_1486 | 82.24399146 | 0.313408872 | 0.214028 | 1.464335 | 0.143102 | 0.335073 |
| Mfumv2_1487 | 2939.606299 | -0.32961167 | 0.089384 | -3.68757 | 0.000226 | 0.002523 |
| Mfumv2_1488 | 1203.234657 | -0.455691957 | 0.140871 | -3.23482 | 0.001217 | 0.009665 |
| Mfumv2_1489 | 193.1743135 | 0.127348462 | 0.180375 | 0.706021 | 0.480175 | 0.707756 |
| Mfumv2_1490 | 192.3650413 | -0.019021163 | 0.19473 | -0.09768 | 0.922187 | 0.969478 |
| Mfumv2_1491 | 601.1968161 | 0.003294651 | 0.125645 | 0.026222 | 0.97908 | 0.992418 |
| Mfumv2_1492 | 8940.875591 | -0.291905763 | 0.095817 | -3.0465 | 0.002315 | 0.01632 |
| Mfumv2_1493 | 3580.010607 | -0.347078713 | 0.094147 | -3.68655 | 0.000227 | 0.002523 |
| Mfumv2_1494 | 11582.96657 | -0.279343975 | 0.104394 | -2.67586 | 0.007454 | 0.040914 |
| Mfumv2_1495 | 6163.12507 | -0.417415639 | 0.123994 | -3.36643 | 0.000761 | 0.006708 |
| Mfumv2_1496 | 28895.06366 | -0.49277215 | 0.12869 | -3.82914 | 0.000129 | 0.001575 |
| Mfumv2_1497 | 488.0232982 | -0.24644842 | 0.142085 | -1.73452 | 0.082827 | 0.236026 |
| Mfumv2_1498 | 2573.476511 | -0.321739992 | 0.095927 | -3.35402 | 0.000796 | 0.006927 |
| Mfumv2_1499 | 1900.644291 | -0.396325396 | 0.103833 | -3.81696 | 0.000135 | 0.001625 |
| Mfumv2_1500 | 1843.151067 | -0.417856249 | 0.162353 | -2.57376 | 0.01006 | 0.051427 |
| Mfumv2_1501 | 90.89273493 | -0.24089064 | 0.209628 | -1.14913 | 0.2505 | 0.490502 |
| Mfumv2_1502 | 231.6645539 | 0.680020416 | 0.194649 | 3.49357 | 0.000477 | 0.004761 |
| Mfumv2_1504 | 13.58407855 | 0.372041214 | 0.487215 | 0.763608 | 0.445101 | NA |
| Mfumv2_1505 | 1.442803877 | 1.317611037 | 1.557438 | 0.846012 | 0.397546 | NA |
| Mfumv2_1506 | 0 | NA | NA | NA | NA | NA |
| Mfumv2_1507 | 0 | NA | NA | NA | NA | NA |
| Mfumv2_1508 | 0 | NA | NA | NA | NA | NA |
| Mfumv2_1509 | 13.23165679 | -0.322004729 | 0.525679 | -0.61255 | 0.540174 | NA |
| Mfumv2_1510 | 396.8122059 | -0.320250391 | 0.211004 | -1.51775 | 0.129078 | 0.316627 |
| Mfumv2_1511 | 596.0829887 | -0.400958373 | 0.143708 | -2.79009 | 0.005269 | 0.031227 |
| Mfumv2_1512 | 1802.408803 | -0.274548888 | 0.122816 | -2.23544 | 0.025388 | 0.101603 |
| Mfumv2_1513 | 2557.91482 | -0.20225757 | 0.090698 | -2.23002 | 0.025746 | 0.102831 |
| Mfumv2_1514 | 819.3439698 | -0.111007708 | 0.114131 | -0.97264 | 0.330734 | 0.573291 |
| Mfumv2_1515 | 1212.985163 | -0.488013195 | 0.124906 | -3.90706 | 9.34E-05 | 0.001219 |
| Mfumv2_1516 | 1352.425574 | -0.15539098 | 0.113105 | -1.37386 | 0.169484 | 0.380866 |
| Mfumv2_1517 | 2125.518551 | -0.189287327 | 0.138384 | -1.36784 | 0.171363 | 0.384658 |
| Mfumv2_1518 | 549.9688715 | -0.126854977 | 0.127837 | -0.99232 | 0.321044 | 0.565769 |
| Mfumv2_1519 | 2610.005918 | -0.194443065 | 0.117481 | -1.6551 | 0.097904 | 0.267968 |
| Mfumv2_1520 | 3619.977636 | -0.144438267 | 0.123046 | -1.17386 | 0.240452 | 0.47971 |
| Mfumv2_1521 | 519.2003604 | 0.05758711 | 0.148511 | 0.387765 | 0.69819 | 0.850369 |
| Mfumv2_1522 | 172.9593001 | -0.21096187 | 0.163558 | -1.28983 | 0.19711 | 0.426259 |
| Mfumv2_1523 | 670.8350767 | -0.403078171 | 0.110892 | -3.63487 | 0.000278 | 0.003004 |
| Mfumv2_1524 | 1014.598655 | -0.043163971 | 0.11038 | -0.39105 | 0.69576 | 0.850369 |

| | | | | | | |
|---|---|---|---|---|---|---|
| Mfumv2_1525 | 938.4625301 | -0.026628617 | 0.121263 | -0.21959 | 0.826188 | 0.915001 |
| Mfumv2_1526 | 325.4872429 | 0.141664919 | 0.132113 | 1.072301 | 0.283585 | 0.523113 |
| Mfumv2_1527 | 82.94193986 | 0.088188872 | 0.2223 | 0.396712 | 0.69158 | 0.847961 |
| Mfumv2_1528 | 908.5666913 | -0.198110794 | 0.12993 | -1.52475 | 0.127322 | 0.313084 |
| Mfumv2_1529 | 112.9089643 | 0.235278906 | 0.189667 | 1.240484 | 0.214796 | 0.445791 |
| Mfumv2_1530 | 484.3563729 | -0.13878434 | 0.114157 | -1.21574 | 0.224086 | 0.45844 |
| Mfumv2_1531 | 344.1835838 | 1.099019971 | 0.183716 | 5.982154 | 2.20E-09 | 8.51E-08 |
| Mfumv2_1532 | 37.28020719 | 0.400817993 | 0.33821 | 1.185115 | 0.235972 | 0.474068 |
| Mfumv2_1533 | 2.117498809 | -0.879808035 | 1.20085 | -0.73265 | 0.463769 | NA |
| Mfumv2_1534 | 138.8434686 | 0.070372536 | 0.271432 | 0.259264 | 0.795431 | 0.900294 |
| Mfumv2_1535 | 4.689722112 | 0.557870892 | 0.862231 | 0.647008 | 0.517627 | NA |
| Mfumv2_1536 | 10.76398007 | -0.020105596 | 0.610796 | -0.03292 | 0.973741 | NA |
| Mfumv2_1537 | 101.6374189 | -0.076528459 | 0.197676 | -0.38714 | 0.698652 | 0.850369 |
| Mfumv2_1538 | 66.91967582 | -0.224918165 | 0.244782 | -0.91885 | 0.358173 | 0.59836 |
| Mfumv2_1539 | 642.1117974 | -2.35000174 | 0.137133 | -17.1367 | 7.90E-66 | 2.64E-63 |
| Mfumv2_1540 | 80.80017891 | -1.052785306 | 0.225741 | -4.66368 | 3.11E-06 | 6.37E-05 |
| Mfumv2_1541 | 31.51699386 | 0.184224172 | 0.381024 | 0.483498 | 0.628742 | 0.82074 |
| Mfumv2_1542 | 11.60790675 | 0.673547049 | 0.542231 | 1.242176 | 0.214172 | NA |
| Mfumv2_1543 | 2.837615239 | 0.286726836 | 1.045284 | 0.274305 | 0.78385 | NA |
| Mfumv2_1544 | 3.431366718 | 0.464132869 | 0.945942 | 0.490657 | 0.623669 | NA |
| Mfumv2_1545 | 5.493820335 | 1.194730976 | 0.78111 | 1.529529 | 0.126133 | NA |
| Mfumv2_1546 | 303.3365452 | -0.17399223 | 0.177039 | -0.98279 | 0.325711 | 0.568517 |
| Mfumv2_1547 | 425.9322086 | 0.18025259 | 0.127966 | 1.408602 | 0.158953 | 0.363709 |
| Mfumv2_1549 | 28.96229181 | 0.568255068 | 0.379897 | 1.495812 | 0.134703 | 0.323319 |
| Mfumv2_1550 | 0.364606656 | -0.677928345 | 2.932995 | -0.23114 | 0.817207 | NA |
| Mfumv2_1551 | 15.12694244 | 0.471990476 | 0.464648 | 1.015803 | 0.309723 | NA |
| Mfumv2_1552 | 297.9640486 | -0.42422645 | 0.137705 | -3.08069 | 0.002065 | 0.014925 |
| Mfumv2_1553 | 11.28836423 | -0.024540486 | 0.542045 | -0.04527 | 0.963889 | NA |
| Mfumv2_1556 | 731.7651176 | -0.235913686 | 0.133883 | -1.76209 | 0.078054 | 0.227347 |
| Mfumv2_1557 | 136.4759743 | 0.181419924 | 0.212417 | 0.854073 | 0.393065 | 0.625684 |
| Mfumv2_1558 | 267.4824099 | -0.129255342 | 0.135617 | -0.95309 | 0.340545 | 0.584519 |
| Mfumv2_1559 | 48.70679225 | -0.336674128 | 0.343518 | -0.98008 | 0.327049 | 0.569853 |
| Mfumv2_1560 | 152.6815963 | 0.215384378 | 0.169831 | 1.268226 | 0.204717 | 0.434665 |
| Mfumv2_1561 | 123.8623997 | 0.281757856 | 0.186527 | 1.510549 | 0.130903 | 0.318384 |
| Mfumv2_1562 | 39.68782926 | 0.397855386 | 0.3126 | 1.27273 | 0.203114 | 0.433912 |
| Mfumv2_1563 | 53.43497444 | 0.667072867 | 0.260924 | 2.556583 | 0.010571 | 0.053224 |
| Mfumv2_1564 | 232.2321512 | 0.440701438 | 0.169042 | 2.607051 | 0.009133 | 0.047904 |
| Mfumv2_1565 | 366.4102193 | 0.472436551 | 0.148509 | 3.181196 | 0.001467 | 0.011333 |
| Mfumv2_1567 | 861.8968415 | 0.087741775 | 0.557977 | 0.15725 | 0.875048 | 0.943624 |
| Mfumv2_1568 | 43.37011803 | 0.532109042 | 0.282073 | 1.886424 | 0.059238 | 0.185951 |
| Mfumv2_1570 | 332.8193756 | -0.098717785 | 0.132992 | -0.74228 | 0.457917 | 0.686534 |
| Mfumv2_1571 | 1.430197108 | -0.241330894 | 1.459022 | -0.16541 | 0.868625 | NA |
| Mfumv2_1572 | 43.16877835 | 0.076610996 | 0.289718 | 0.264433 | 0.791446 | 0.897706 |
| Mfumv2_1573 | 34.23701181 | 0.047209117 | 0.340613 | 0.138601 | 0.889766 | 0.94819 |
| Mfumv2_1575 | 846.8999788 | -0.1272872 | 0.105798 | -1.20311 | 0.228933 | 0.465042 |
| Mfumv2_1576 | 248.5243918 | 0.306407015 | 0.163615 | 1.872729 | 0.061106 | 0.19092 |
| Mfumv2_1577 | 618.7843325 | -0.230135509 | 0.158284 | -1.45394 | 0.145962 | 0.339396 |
| Mfumv2_1578 | 553.2896731 | -0.108061205 | 0.12092 | -0.89366 | 0.371504 | 0.609266 |
| Mfumv2_1579 | 367.4714386 | 0.374949957 | 0.128781 | 2.911538 | 0.003597 | 0.023146 |
| Mfumv2_1580 | 558.5614728 | -0.267983752 | 0.136869 | -1.95795 | 0.050236 | 0.166266 |
| Mfumv2_1581 | 1774.356151 | -0.200844305 | 0.114825 | -1.74914 | 0.080267 | 0.23169 |
| Mfumv2_1582 | 640.5691474 | -0.373185575 | 0.137891 | -2.70638 | 0.006802 | 0.03775 |

| | | | | | | |
|---|---|---|---|---|---|---|
| Mfumv2_1583 | 411.4756507 | -0.278910491 | 0.163741 | -1.70336 | 0.0885 | 0.249016 |
| Mfumv2_1584 | 7.194160148 | 0.267004266 | 0.651485 | 0.409839 | 0.681924 | NA |
| Mfumv2_1585 | 842.0059197 | -0.206879366 | 0.110713 | -1.86861 | 0.061677 | 0.191971 |
| Mfumv2_1586 | 395.1217427 | 0.265877927 | 0.1251 | 2.125323 | 0.03356 | 0.123482 |
| Mfumv2_1587 | 582.8379002 | 0.01997968 | 0.110244 | 0.181231 | 0.856187 | 0.934825 |
| Mfumv2_1588 | 224.4837467 | -0.265674092 | 0.152527 | -1.74181 | 0.081541 | 0.234023 |
| Mfumv2_1589 | 24.6777987 | 0.260314499 | 0.365368 | 0.712471 | 0.476173 | 0.702889 |
| Mfumv2_1590 | 136.2648479 | 0.212266671 | 0.17223 | 1.232457 | 0.217778 | 0.449657 |
| Mfumv2_1591 | 10.26070183 | 0.90902488 | 0.637092 | 1.426836 | 0.153627 | NA |
| Mfumv2_1592 | 49.77167674 | 0.2179746 | 0.265624 | 0.820613 | 0.411867 | 0.646438 |
| Mfumv2_1593 | 53.94674038 | 0.428891519 | 0.263425 | 1.628135 | 0.103496 | 0.278943 |
| Mfumv2_1594 | 143.692831 | 0.045898992 | 0.172322 | 0.266355 | 0.789966 | 0.897706 |
| Mfumv2_1595 | 11.10154484 | 0.088652553 | 0.558466 | 0.158743 | 0.873871 | NA |
| Mfumv2_1596 | 88.05555607 | 0.250794146 | 0.209779 | 1.195515 | 0.231886 | 0.468721 |
| Mfumv2_1597 | 3.571388375 | 0.040430677 | 0.989807 | 0.040847 | 0.967418 | NA |
| Mfumv2_1598 | 1.067253416 | 0.655204111 | 1.691436 | 0.387366 | 0.698486 | NA |
| Mfumv2_1599 | 152.4667454 | -0.163856295 | 0.178853 | -0.91615 | 0.359589 | 0.59852 |
| Mfumv2_1600 | 187.8841799 | -0.111837832 | 0.172005 | -0.6502 | 0.515562 | 0.738249 |
| Mfumv2_1601 | 114.6386894 | -1.585847896 | 0.231077 | -6.86284 | 6.75E-12 | 3.50E-10 |
| Mfumv2_1602 | 204.4075375 | -1.409393229 | 0.170282 | -8.27684 | 1.26E-16 | 9.41E-15 |
| Mfumv2_1603 | 33.05444413 | -1.728794431 | 0.338674 | -5.1046 | 3.31E-07 | 7.65E-06 |
| Mfumv2_1604 | 304.2056189 | -1.825932333 | 0.200944 | -9.08677 | 1.02E-19 | 8.20E-18 |
| Mfumv2_1605 | 721.0958321 | -1.559494853 | 0.153058 | -10.1889 | 2.22E-24 | 2.48E-22 |
| Mfumv2_1606 | 2664.082637 | -1.524399166 | 0.135927 | -11.2148 | 3.45E-29 | 4.95E-27 |
| Mfumv2_1607 | 0.675912109 | 2.530577072 | 2.108171 | 1.200366 | 0.229997 | NA |
| Mfumv2_1608 | 174.4440709 | -0.787106612 | 0.182097 | -4.32247 | 1.54E-05 | 0.000267 |
| Mfumv2_1609 | 181.3742974 | -0.960121932 | 0.165644 | -5.79631 | 6.78E-09 | 2.39E-07 |
| Mfumv2_1610 | 335.4623896 | -1.064036007 | 0.13599 | -7.82437 | 5.10E-15 | 3.53E-13 |
| Mfumv2_1611 | 159.5787795 | -0.536099162 | 0.180664 | -2.96739 | 0.003003 | 0.019914 |
| Mfumv2_1612 | 167.3314076 | -0.197782774 | 0.167497 | -1.18081 | 0.237677 | 0.476065 |
| Mfumv2_1613 | 3717.212097 | -0.532487468 | 0.139835 | -3.80797 | 0.00014 | 0.001666 |
| Mfumv2_1614 | 2845.371461 | -0.168124606 | 0.112775 | -1.49079 | 0.136016 | 0.325306 |
| Mfumv2_1615 | 103.2095451 | 0.409123099 | 0.201403 | 2.031363 | 0.042218 | 0.145483 |
| Mfumv2_1616 | 190.8894371 | -0.615524124 | 0.18118 | -3.39731 | 0.000681 | 0.006158 |
| Mfumv2_1617 | 14.95269589 | 0.60336726 | 0.494466 | 1.22024 | 0.222374 | NA |
| Mfumv2_1618 | 22.87127849 | 0.407175863 | 0.420252 | 0.968886 | 0.332602 | NA |
| Mfumv2_1619 | 152.9497769 | 0.207770744 | 0.1807 | 1.149809 | 0.250223 | 0.490502 |
| Mfumv2_1620 | 51.64751989 | -0.165814333 | 0.278835 | -0.59467 | 0.552065 | 0.766482 |
| Mfumv2_1621 | 104.650334 | -0.088499316 | 0.20542 | -0.43082 | 0.666598 | 0.835952 |
| Mfumv2_1622 | 7.610117828 | 0.260874637 | 0.631412 | 0.413161 | 0.679489 | NA |
| Mfumv2_1623 | 5.073980855 | -0.171899847 | 0.76405 | -0.22499 | 0.821991 | NA |
| Mfumv2_1624 | 1.505105239 | 2.510223474 | 1.642631 | 1.528172 | 0.12647 | NA |
| Mfumv2_1625 | 0.255553176 | 1.14483936 | 3.242772 | 0.353043 | 0.724056 | NA |
| Mfumv2_1626 | 0 | NA | NA | NA | NA | NA |
| Mfumv2_1628 | 7.375328318 | 0.476477599 | 0.681141 | 0.699528 | 0.484222 | NA |
| Mfumv2_1629 | 2.885010589 | 0.213977948 | 1.118313 | 0.19134 | 0.848259 | NA |
| Mfumv2_1630 | 95.54601665 | -0.008559703 | 0.204298 | -0.0419 | 0.96658 | 0.988892 |
| Mfumv2_1631 | 201.831977 | -0.522953591 | 0.188894 | -2.7685 | 0.005632 | 0.032482 |
| Mfumv2_1632 | 215.3697909 | -0.170609246 | 0.154299 | -1.1057 | 0.268855 | 0.507699 |
| Mfumv2_1633 | 585.5072395 | -0.126043025 | 0.142308 | -0.88571 | 0.375776 | 0.611282 |
| Mfumv2_1634 | 292.3122568 | 0.040600099 | 0.143767 | 0.282402 | 0.777635 | 0.892523 |
| Mfumv2_1635 | 1222.843486 | -0.059108404 | 0.096811 | -0.61056 | 0.541493 | 0.76074 |

| | | | | | | |
|---|---|---|---|---|---|---|
| Mfumv2_1637 | 93.98890776 | -0.055481287 | 0.204038 | -0.27192 | 0.785687 | 0.89568 |
| Mfumv2_1638 | 95.47179985 | 0.308816492 | 0.242798 | 1.271908 | 0.203406 | 0.433912 |
| Mfumv2_1639 | 169.9616115 | 0.383192069 | 0.168638 | 2.272275 | 0.02307 | 0.096467 |
| Mfumv2_1640 | 425.2751067 | 0.2188767 | 0.177093 | 1.23594 | 0.216481 | 0.447899 |
| Mfumv2_1641 | 661.2118683 | 0.161792453 | 0.119734 | 1.351262 | 0.176612 | 0.392907 |
| Mfumv2_1642 | 1415.402777 | -0.068392991 | 0.111077 | -0.61573 | 0.538075 | 0.758591 |
| Mfumv2_1643 | 1002.230013 | -0.396159324 | 0.137085 | -2.88988 | 0.003854 | 0.024501 |
| Mfumv2_1644 | 165.7030728 | -0.061536477 | 0.167658 | -0.36704 | 0.713592 | 0.859991 |
| Mfumv2_1645 | 236.3650685 | 0.059233011 | 0.148065 | 0.400047 | 0.689122 | 0.846498 |
| Mfumv2_1646 | 594.3496298 | -0.142584841 | 0.116684 | -1.22198 | 0.221717 | 0.456269 |
| Mfumv2_1647 | 15.90019383 | -0.125994702 | 0.447176 | -0.28176 | 0.77813 | NA |
| Mfumv2_1648 | 8.557350738 | 0.339164099 | 0.600315 | 0.564977 | 0.57209 | NA |
| Mfumv2_1649 | 11.14511897 | 0.56656286 | 0.551433 | 1.027437 | 0.304215 | NA |
| Mfumv2_1650 | 31.39034928 | 0.128984232 | 0.332178 | 0.388299 | 0.697795 | 0.850369 |
| Mfumv2_1651 | 89.088789 | -0.283072216 | 0.20785 | -1.3619 | 0.173228 | 0.387545 |
| Mfumv2_1652 | 156.9498187 | 0.31860078 | 0.180308 | 1.766984 | 0.077231 | 0.226506 |
| Mfumv2_1653 | 271.5233307 | 0.078118635 | 0.14272 | 0.547357 | 0.584133 | 0.790427 |
| Mfumv2_1654 | 542.4955736 | -0.100226466 | 0.128718 | -0.77865 | 0.436186 | 0.666893 |
| Mfumv2_1655 | 1101.361011 | -0.163880988 | 0.126074 | -1.29988 | 0.193644 | 0.421298 |
| Mfumv2_1656 | 559.4595199 | 0.401888862 | 0.178756 | 2.248258 | 0.02456 | 0.09944 |
| Mfumv2_1657 | 102.5951037 | 0.300463387 | 0.201255 | 1.492946 | 0.135451 | 0.324727 |
| Mfumv2_1658 | 44.99223549 | -0.274505709 | 0.310306 | -0.88463 | 0.376357 | 0.611732 |
| Mfumv2_1659 | 70.88981709 | -0.599743603 | 0.26757 | -2.24145 | 0.024997 | 0.100438 |
| Mfumv2_1660 | 95.24070839 | 0.062579403 | 0.215666 | 0.290168 | 0.771688 | 0.888026 |
| Mfumv2_1661 | 36.09533166 | 0.308538753 | 0.327109 | 0.943229 | 0.345564 | 0.588836 |
| Mfumv2_1662 | 129.8318955 | 0.636795585 | 0.18949 | 3.360574 | 0.000778 | 0.006794 |
| Mfumv2_1664 | 401.2985219 | -0.218297626 | 0.17572 | -1.2423 | 0.214125 | 0.444857 |
| Mfumv2_1666 | 11.97567447 | -0.157528136 | 0.504064 | -0.31252 | 0.754648 | NA |
| Mfumv2_1667 | 176.8888065 | 0.076747966 | 0.172142 | 0.445842 | 0.655711 | 0.832295 |
| Mfumv2_1668 | 723.7448967 | -0.687289711 | 0.14533 | -4.72916 | 2.25E-06 | 4.67E-05 |
| Mfumv2_1669 | 62.24646236 | 0.066193493 | 0.264566 | 0.250196 | 0.802436 | 0.903903 |
| Mfumv2_1670 | 433.6047824 | -0.395953569 | 0.1432 | -2.76505 | 0.005691 | 0.032483 |
| Mfumv2_1671 | 64.11067112 | -0.117424224 | 0.267152 | -0.43954 | 0.66027 | 0.835316 |
| Mfumv2_1672 | 673.3554163 | 0.292296857 | 0.168084 | 1.738989 | 0.082037 | 0.234775 |
| Mfumv2_1673 | 1.728792372 | 0.9301812 | 1.393747 | 0.667396 | 0.504519 | NA |
| Mfumv2_1674 | 2.059915635 | 0.366736342 | 1.324648 | 0.276856 | 0.781891 | NA |
| Mfumv2_1675 | 227.0226915 | -0.148666499 | 0.145541 | -1.02148 | 0.307028 | 0.548772 |
| Mfumv2_1676 | 314.2591004 | 0.126536856 | 0.133656 | 0.946738 | 0.343772 | 0.587277 |
| Mfumv2_1677 | 212.0452255 | 0.33013995 | 0.162474 | 2.031955 | 0.042158 | 0.145483 |
| Mfumv2_1678 | 1357.612495 | -0.121413519 | 0.135526 | -0.89587 | 0.370322 | 0.608143 |
| Mfumv2_1679 | 150.386473 | 0.091776652 | 0.199918 | 0.459071 | 0.646183 | 0.827395 |
| Mfumv2_1680 | 0 | NA | NA | NA | NA | NA |
| Mfumv2_1681 | 773.598856 | -0.245467009 | 0.112842 | -2.17531 | 0.029607 | 0.112732 |
| Mfumv2_1682 | 315.2755068 | -0.01410042 | 0.136662 | -0.10318 | 0.917822 | 0.967421 |
| Mfumv2_1683 | 484.2549607 | 0.07556196 | 0.122215 | 0.618269 | 0.536398 | 0.75729 |
| Mfumv2_1684 | 266.9346248 | -0.188395055 | 0.136952 | -1.37563 | 0.168935 | 0.380068 |
| Mfumv2_1685 | 366.6157929 | -0.032518481 | 0.134484 | -0.2418 | 0.808934 | 0.907397 |
| Mfumv2_1686 | 543.7788228 | 0.098443507 | 0.115997 | 0.848675 | 0.396062 | 0.627515 |
| Mfumv2_1687 | 460.3167522 | -0.041610836 | 0.116402 | -0.35747 | 0.720737 | 0.866005 |
| Mfumv2_1688 | 313.7772832 | -0.02204062 | 0.131932 | -0.16706 | 0.867323 | 0.938842 |
| Mfumv2_1690 | 531.615327 | -0.176920264 | 0.111411 | -1.588 | 0.112287 | 0.291452 |
| Mfumv2_1691 | 1067.193181 | -0.179474515 | 0.104521 | -1.71711 | 0.085958 | 0.243226 |

| | | | | | | |
|---|---|---|---|---|---|---|
| Mfumv2_1692 | 355.176436 | 0.188716491 | 0.131575 | 1.434287 | 0.15149 | 0.349419 |
| Mfumv2_1693 | 153.3710327 | 0.045462409 | 0.190032 | 0.239236 | 0.810923 | 0.907508 |
| Mfumv2_1694 | 238.0055873 | 0.078874895 | 0.149386 | 0.527993 | 0.597505 | 0.79866 |
| Mfumv2_1695 | 250.1067091 | 0.347407987 | 0.195917 | 1.773244 | 0.076188 | 0.224309 |
| Mfumv2_1696 | 463.4462735 | -0.238398558 | 0.125434 | -1.90059 | 0.057356 | 0.181461 |
| Mfumv2_1697 | 70.09370738 | 0.172707853 | 0.22332 | 0.773364 | 0.439307 | 0.669636 |
| Mfumv2_1698 | 0 | NA | NA | NA | NA | NA |
| Mfumv2_1699 | 57.75467059 | -0.186726102 | 0.276902 | -0.67434 | 0.500095 | 0.727605 |
| Mfumv2_16s_rRN/ | 233793.0092 | -0.239406713 | 0.148381 | -1.61346 | 0.106645 | 0.283025 |
| Mfumv2_1701 | 8.586139329 | 0.49491314 | 0.650555 | 0.760755 | 0.446803 | NA |
| Mfumv2_1702 | 546.432341 | 0.001295799 | 0.129027 | 0.010043 | 0.991987 | 0.995456 |
| Mfumv2_1703 | 14.3346861 | 0.145539822 | 0.463986 | 0.313673 | 0.75377 | NA |
| Mfumv2_1704 | 259.7745446 | 0.172650818 | 0.147559 | 1.17005 | 0.241981 | 0.481327 |
| Mfumv2_1705 | 122.062501 | -0.135506256 | 0.19089 | -0.70987 | 0.477787 | 0.704753 |
| Mfumv2_1706 | 513.433712 | -0.229016752 | 0.130031 | -1.76125 | 0.078196 | 0.227347 |
| Mfumv2_1707 | 528.3622504 | -0.203140659 | 0.157862 | -1.28682 | 0.198157 | 0.427601 |
| Mfumv2_1709 | 102.4650054 | 0.130499473 | 0.196053 | 0.665634 | 0.505645 | 0.731347 |
| Mfumv2_1710 | 124.9970953 | 0.535626216 | 0.186673 | 2.869326 | 0.004113 | 0.025825 |
| Mfumv2_1711 | 162.5286947 | 0.266775536 | 0.161094 | 1.656028 | 0.097716 | 0.267968 |
| Mfumv2_1712 | 459.537816 | 0.121114144 | 0.131179 | 0.923275 | 0.355864 | 0.59836 |
| Mfumv2_1713 | 3.751468352 | 0.232285792 | 0.960944 | 0.241727 | 0.808992 | NA |
| Mfumv2_1714 | 461.6501678 | -0.315308604 | 0.140285 | -2.24762 | 0.0246 | 0.09944 |
| Mfumv2_1715 | 331.0350148 | -0.275569462 | 0.149325 | -1.84543 | 0.064975 | 0.199594 |
| Mfumv2_1716 | 357.5831634 | 0.261814233 | 0.138516 | 1.890135 | 0.05874 | 0.185197 |
| Mfumv2_1717 | 103.6618181 | 0.614725287 | 0.210972 | 2.913774 | 0.003571 | 0.023067 |
| Mfumv2_1718 | 231.7673714 | -0.044505311 | 0.145193 | -0.30653 | 0.759204 | 0.884498 |
| Mfumv2_1719 | 1519.104595 | -0.035997503 | 0.095811 | -0.37571 | 0.707129 | 0.855281 |
| Mfumv2_1720 | 173.4725916 | 0.216249793 | 0.163316 | 1.324122 | 0.185462 | 0.40507 |
| Mfumv2_1721 | 86.13879054 | 0.119178244 | 0.222312 | 0.536085 | 0.591899 | 0.793988 |
| Mfumv2_1722 | 0.285988495 | -0.677928192 | 3.169635 | -0.21388 | 0.830639 | NA |
| Mfumv2_1723 | 0 | NA | NA | NA | NA | NA |
| Mfumv2_1724 | 0.140690113 | 0.283845181 | 4.080473 | 0.069562 | 0.944542 | NA |
| Mfumv2_1725 | 0 | NA | NA | NA | NA | NA |
| Mfumv2_1726 | 1.206509975 | 0.982840376 | 1.671629 | 0.587954 | 0.556563 | NA |
| Mfumv2_1727 | 217.8369303 | 0.226522466 | 0.148009 | 1.530468 | 0.125901 | 0.311041 |
| Mfumv2_1728 | 625.8339607 | 0.097224231 | 0.131325 | 0.740334 | 0.459097 | 0.687278 |
| Mfumv2_1729 | 526.7969906 | -0.492262445 | 0.122022 | -4.03423 | 5.48E-05 | 0.000777 |
| Mfumv2_1730 | 2108.917235 | -0.235885959 | 0.1392 | -1.69458 | 0.090156 | 0.252964 |
| Mfumv2_1731 | 2504.426345 | -0.201282169 | 0.136689 | -1.47256 | 0.14087 | 0.33186 |
| Mfumv2_1732 | 3040.579869 | -0.039050641 | 0.132146 | -0.29551 | 0.767604 | 0.887638 |
| Mfumv2_1733 | 3809.124633 | 0.181761591 | 0.118761 | 1.530481 | 0.125898 | 0.311041 |
| Mfumv2_1734 | 4082.721977 | 0.527109896 | 0.115573 | 4.560843 | 5.09E-06 | 9.84E-05 |
| Mfumv2_1735 | 1020.440244 | 0.44881222 | 0.128929 | 3.481072 | 0.000499 | 0.004847 |
| Mfumv2_1736 | 286.6369257 | -0.065151689 | 0.133919 | -0.4865 | 0.626613 | 0.819574 |
| Mfumv2_1737 | 909.9738092 | 0.010038825 | 0.111857 | 0.089747 | 0.928488 | 0.972652 |
| Mfumv2_1738 | 361.611869 | -0.239055583 | 0.157887 | -1.51409 | 0.130003 | 0.317732 |
| Mfumv2_1739 | 138.3686134 | 0.003515203 | 0.177684 | 0.019783 | 0.984216 | 0.992996 |
| Mfumv2_1741 | 253.1303244 | -0.043007348 | 0.143576 | -0.29954 | 0.764524 | 0.887034 |
| Mfumv2_1742 | 16.34211124 | 0.353154631 | 0.446549 | 0.790852 | 0.42903 | NA |
| Mfumv2_1743 | 1.224235104 | 0.003907563 | 1.66797 | 0.002343 | 0.998131 | NA |
| Mfumv2_1744 | 72.88803457 | 0.323226857 | 0.225792 | 1.431527 | 0.152279 | 0.350434 |
| Mfumv2_1745 | 206.5639266 | -0.273029497 | 0.171834 | -1.58892 | 0.112079 | 0.291291 |

| | | | | | | |
|---|---|---|---|---|---|---|
| Mfumv2_1746 | 294.1527333 | 0.162990854 | 0.140045 | 1.163845 | 0.244487 | 0.483537 |
| Mfumv2_1747 | 524.1212915 | 0.046063955 | 0.122102 | 0.377259 | 0.705981 | 0.855161 |
| Mfumv2_1749 | 51.15793543 | 0.368031234 | 0.261819 | 1.405671 | 0.159822 | 0.364866 |
| Mfumv2_1750 | 809.2838213 | -0.25510035 | 0.143411 | -1.77881 | 0.075272 | 0.222056 |
| Mfumv2_1751 | 1167.421192 | -0.125253258 | 0.104775 | -1.19545 | 0.231911 | 0.468721 |
| Mfumv2_1752 | 854.313536 | -0.472448872 | 0.12195 | -3.87412 | 0.000107 | 0.001361 |
| Mfumv2_1753 | 320.1499491 | 0.024524382 | 0.139557 | 0.17573 | 0.860506 | 0.936786 |
| Mfumv2_1754 | 130.6773975 | 0.283222432 | 0.176751 | 1.602381 | 0.109071 | 0.288196 |
| Mfumv2_1755 | 326.9864111 | 0.440103503 | 0.144314 | 3.049634 | 0.002291 | 0.01632 |
| Mfumv2_1756 | 597.9036452 | 0.106419551 | 0.113157 | 0.940461 | 0.346981 | 0.590453 |
| Mfumv2_1757 | 789.653002 | 0.093604109 | 0.143085 | 0.654183 | 0.512994 | 0.736672 |
| Mfumv2_1758 | 4215.672666 | 0.116151285 | 0.117625 | 0.987473 | 0.323411 | 0.566462 |
| Mfumv2_1759 | 2549.916588 | -0.607450044 | 0.145895 | -4.16362 | 3.13E-05 | 0.000496 |
| Mfumv2_1760 | 112.9336278 | -0.083889012 | 0.184644 | -0.45433 | 0.649592 | 0.829769 |
| Mfumv2_1761 | 212.1093408 | 0.081585656 | 0.16447 | 0.496052 | 0.619858 | 0.814983 |
| Mfumv2_1762 | 602.705384 | -0.078356825 | 0.108688 | -0.72093 | 0.470952 | 0.698655 |
| Mfumv2_1763 | 88.94869701 | 0.283521602 | 0.204819 | 1.384257 | 0.16628 | 0.375766 |
| Mfumv2_1764 | 176.0914614 | 0.620305589 | 0.176626 | 3.511973 | 0.000445 | 0.004468 |
| Mfumv2_1765 | 482.1973877 | -0.021878429 | 0.14343 | -0.15254 | 0.878763 | 0.945691 |
| Mfumv2_1766 | 65.15064342 | 0.145597376 | 0.242544 | 0.600293 | 0.548311 | 0.76593 |
| Mfumv2_1767 | 78.91304777 | 0.334853846 | 0.214027 | 1.56454 | 0.117691 | 0.300433 |
| Mfumv2_1768 | 92.03891742 | -0.331211417 | 0.250989 | -1.31962 | 0.18696 | 0.407821 |
| Mfumv2_1769 | 739.541375 | -0.16579032 | 0.117804 | -1.40734 | 0.159327 | 0.364149 |
| Mfumv2_1770 | 170.5399433 | -0.345603671 | 0.163955 | -2.10792 | 0.035038 | 0.127521 |
| Mfumv2_1771 | 495.24156 | -0.230094328 | 0.148397 | -1.55053 | 0.121014 | 0.304876 |
| Mfumv2_1772 | 410.2986567 | -0.323500931 | 0.134725 | -2.4012 | 0.016342 | 0.073776 |
| Mfumv2_1773 | 295.4657283 | 0.199859202 | 0.139454 | 1.43316 | 0.151812 | 0.34976 |
| Mfumv2_1774 | 3074.262194 | -0.298782315 | 0.114445 | -2.61072 | 0.009035 | 0.047518 |
| Mfumv2_1775 | 240.4607098 | 0.277276874 | 0.141848 | 1.954744 | 0.050613 | 0.166966 |
| Mfumv2_1776 | 1901.844659 | -0.026759418 | 0.108969 | -0.24557 | 0.806015 | 0.90564 |
| Mfumv2_1777 | 10.39922155 | 0.247558548 | 0.596259 | 0.415186 | 0.678005 | NA |
| Mfumv2_1779 | 217.0666892 | 0.26931601 | 0.181692 | 1.48227 | 0.138268 | 0.328735 |
| Mfumv2_1780 | 189.049811 | 0.583265628 | 0.167425 | 3.483752 | 0.000494 | 0.004845 |
| Mfumv2_1781 | 4.186614261 | 1.040163232 | 0.903287 | 1.151531 | 0.249514 | NA |
| Mfumv2_1782 | 12.82207573 | 0.074932064 | 0.499675 | 0.149962 | 0.880795 | NA |
| Mfumv2_1784 | 7.583590359 | 0.808859344 | 0.671973 | 1.203708 | 0.228703 | NA |
| Mfumv2_1785 | 175.2377494 | 1.041174146 | 0.176473 | 5.899908 | 3.64E-09 | 1.35E-07 |
| Mfumv2_1786 | 598.2434595 | 0.663083519 | 0.110213 | 6.016392 | 1.78E-09 | 7.03E-08 |
| Mfumv2_1787 | 988.0774735 | 0.258817614 | 0.147015 | 1.760481 | 0.078326 | 0.227395 |
| Mfumv2_1788 | 36621.80025 | 0.864629608 | 0.126759 | 6.821027 | 9.04E-12 | 4.43E-10 |
| Mfumv2_1789 | 19.50343619 | 1.038256682 | 0.438745 | 2.366426 | 0.017961 | NA |
| Mfumv2_1790 | 14.60053159 | -0.186406758 | 0.507152 | -0.36756 | 0.713205 | NA |
| Mfumv2_1791 | 135092.5276 | -0.714439094 | 0.141324 | -5.05534 | 4.30E-07 | 9.70E-06 |
| Mfumv2_1792 | 83669.34156 | -0.764520709 | 0.132207 | -5.78274 | 7.35E-09 | 2.50E-07 |
| Mfumv2_1793 | 245684.7111 | -0.695037011 | 0.122216 | -5.68696 | 1.29E-08 | 4.19E-07 |
| Mfumv2_1794 | 260.4600091 | 0.049124247 | 0.137085 | 0.358349 | 0.720082 | 0.866005 |
| Mfumv2_1795 | 71.21298509 | 0.297637671 | 0.243298 | 1.223344 | 0.2212 | 0.455785 |
| Mfumv2_1796 | 257.573448 | -0.100779832 | 0.169195 | -0.59564 | 0.551413 | 0.766105 |
| Mfumv2_1797 | 4.65482974 | 1.739750816 | 0.912124 | 1.907362 | 0.056474 | NA |
| Mfumv2_1798 | 132.1105513 | -0.151550078 | 0.19421 | -0.78034 | 0.435191 | 0.666666 |
| Mfumv2_1799 | 1421.470951 | -0.598140406 | 0.141838 | -4.21705 | 2.48E-05 | 0.000401 |
| Mfumv2_1800 | 164.7537059 | 0.386475688 | 0.160791 | 2.403598 | 0.016235 | 0.073624 |

| | | | | | | |
|---|---|---|---|---|---|---|
| Mfumv2_1801 | 49.15442208 | 0.174768503 | 0.277083 | 0.630745 | 0.528207 | 0.750482 |
| Mfumv2_1802 | 103.7751399 | 0.115668884 | 0.19237 | 0.601283 | 0.547651 | 0.765645 |
| Mfumv2_1803 | 75.20923677 | -0.023890552 | 0.225876 | -0.10577 | 0.915766 | 0.966776 |
| Mfumv2_1804 | 60.0974186 | 0.021091558 | 0.245498 | 0.085913 | 0.931535 | 0.973858 |
| Mfumv2_1805 | 225.3695885 | -0.145279958 | 0.160024 | -0.90787 | 0.363949 | 0.603278 |
| Mfumv2_1806 | 75.46433629 | 0.332477914 | 0.222821 | 1.49213 | 0.135665 | 0.324853 |
| Mfumv2_1807 | 399.5894728 | -0.117487511 | 0.127651 | -0.92038 | 0.357373 | 0.59836 |
| Mfumv2_1808 | 294.9100073 | -0.013862212 | 0.142804 | -0.09707 | 0.92267 | 0.969479 |
| Mfumv2_1809 | 2.835083758 | 1.406682819 | 1.162809 | 1.209728 | 0.226383 | NA |
| Mfumv2_1810 | 13.96785856 | 0.465137775 | 0.512507 | 0.907573 | 0.364104 | NA |
| Mfumv2_1811 | 902.4876074 | 0.048283172 | 0.112941 | 0.427508 | 0.66901 | 0.837408 |
| Mfumv2_1812 | 251.3386256 | -0.247186022 | 0.141292 | -1.74947 | 0.080209 | 0.23169 |
| Mfumv2_1813 | 516.1178729 | 0.238737076 | 0.122542 | 1.948207 | 0.05139 | 0.168148 |
| Mfumv2_1814 | 265.5565545 | 0.149722648 | 0.167151 | 0.895734 | 0.370395 | 0.608143 |
| Mfumv2_1815 | 232.6760173 | 0.025377199 | 0.1405 | 0.180621 | 0.856665 | 0.93483 |
| Mfumv2_1816 | 273.5806106 | -0.079459191 | 0.194933 | -0.40762 | 0.683551 | 0.844041 |
| Mfumv2_1817 | 609.2172116 | 0.015825097 | 0.177801 | 0.089004 | 0.929078 | 0.972652 |
| Mfumv2_1818 | 1591.204808 | -0.155130596 | 0.097364 | -1.5933 | 0.111093 | 0.291291 |
| Mfumv2_1819 | 771.0391932 | -0.45503696 | 0.169632 | -2.6825 | 0.007307 | 0.040332 |
| Mfumv2_1820 | 1682.695409 | -0.391409088 | 0.116915 | -3.3478 | 0.000815 | 0.007054 |
| Mfumv2_1821 | 947.330253 | -0.390510004 | 0.127412 | -3.06494 | 0.002177 | 0.015621 |
| Mfumv2_1822 | 640.116434 | 0.12074988 | 0.115673 | 1.043889 | 0.296537 | 0.53545 |
| Mfumv2_1824 | 11.05456249 | 0.536363549 | 0.53733 | 0.998202 | 0.318181 | NA |
| Mfumv2_1825 | 8.096475119 | -0.681711406 | 0.627614 | -1.0862 | 0.277392 | NA |
| Mfumv2_1826 | 1221.162421 | -0.573378128 | 0.175659 | -3.26416 | 0.001098 | 0.008894 |
| Mfumv2_1827 | 0.281380227 | 1.260342684 | 3.513739 | 0.35869 | 0.719827 | NA |
| Mfumv2_1828 | 10.88649249 | 0.380734942 | 0.550671 | 0.691402 | 0.489313 | NA |
| Mfumv2_1829 | 0.255553176 | 1.14483936 | 3.242772 | 0.353043 | 0.724056 | NA |
| Mfumv2_1830 | 3.643457451 | 0.065699715 | 0.930879 | 0.070578 | 0.943734 | NA |
| Mfumv2_1831 | 74.50820873 | 0.240286548 | 0.220542 | 1.089526 | 0.275922 | 0.516133 |
| Mfumv2_1832 | 494.2995074 | -0.150726914 | 0.158393 | -0.9516 | 0.341301 | 0.584519 |
| Mfumv2_1833 | 151.90091 | -0.003307073 | 0.163229 | -0.02026 | 0.983836 | 0.992996 |
| Mfumv2_1834 | 198.8967225 | -0.062575405 | 0.163792 | -0.38204 | 0.70243 | 0.85371 |
| Mfumv2_1835 | 120.3839398 | 0.203806359 | 0.190843 | 1.067924 | 0.285555 | 0.523113 |
| Mfumv2_1836 | 232.8522384 | 0.257050348 | 0.144482 | 1.779116 | 0.075221 | 0.222056 |
| Mfumv2_1837 | 715.7776173 | -0.013174773 | 0.132336 | -0.09956 | 0.920697 | 0.9689 |
| Mfumv2_1838 | 289.821011 | -0.113864989 | 0.131844 | -0.86363 | 0.387789 | 0.621494 |
| Mfumv2_1839 | 495.6675781 | -0.365072743 | 0.143607 | -2.54217 | 0.011017 | 0.054822 |
| Mfumv2_1840 | 1141.982078 | -0.462033915 | 0.135544 | -3.40874 | 0.000653 | 0.006014 |
| Mfumv2_1841 | 416.721923 | -0.003979865 | 0.121384 | -0.03279 | 0.973844 | 0.990165 |
| Mfumv2_1842 | 173.1183448 | 0.139666512 | 0.157583 | 0.886307 | 0.375452 | 0.611251 |
| Mfumv2_1843 | 700.5954247 | 0.100409616 | 0.112877 | 0.889549 | 0.373708 | 0.610542 |
| Mfumv2_1844 | 1328.242011 | -0.090753511 | 0.105439 | -0.86072 | 0.389392 | 0.622615 |
| Mfumv2_1845 | 412.4859358 | 0.090881551 | 0.129769 | 0.700334 | 0.483719 | 0.711319 |
| Mfumv2_1846 | 439.3834241 | 0.08034176 | 0.149921 | 0.535892 | 0.592033 | 0.793988 |
| Mfumv2_1847 | 873.3446623 | 0.000147781 | 0.109638 | 0.001348 | 0.998925 | 0.999422 |
| Mfumv2_1848 | 904.8381197 | -0.14961426 | 0.103282 | -1.44861 | 0.147448 | 0.342058 |
| Mfumv2_1849 | 279.9093567 | -0.215362565 | 0.146895 | -1.4661 | 0.142622 | 0.334337 |
| Mfumv2_1850 | 1602.081825 | -0.445258718 | 0.102416 | -4.34756 | 1.38E-05 | 0.000243 |
| Mfumv2_1851 | 830.6683899 | -0.475766684 | 0.110686 | -4.29833 | 1.72E-05 | 0.00029 |
| Mfumv2_1852 | 521.2433937 | -0.388459181 | 0.129879 | -2.99093 | 0.002781 | 0.019005 |
| Mfumv2_1853 | 396.1314095 | -0.341879651 | 0.133306 | -2.56463 | 0.010329 | 0.052399 |

| | | | | | | |
|---|---|---|---|---|---|---|
| Mfumv2_1854 | 1562.133193 | -0.171256746 | 0.09343 | -1.833 | 0.066803 | 0.203961 |
| Mfumv2_1855 | 270.2800471 | -0.026188793 | 0.145461 | -0.18004 | 0.857121 | 0.93483 |
| Mfumv2_1856 | 109.3095773 | 0.20636345 | 0.186043 | 1.109222 | 0.267335 | 0.506197 |
| Mfumv2_1857 | 193.3928114 | -0.007791457 | 0.202468 | -0.03848 | 0.969303 | 0.98909 |
| Mfumv2_1858 | 1.259349529 | 0.210274779 | 1.669495 | 0.125951 | 0.899771 | NA |
| Mfumv2_1859 | 0.373839054 | 1.700247813 | 2.80949 | 0.60518 | 0.545059 | NA |
| Mfumv2_1860 | 0.362617395 | -0.677928342 | 2.938145 | -0.23073 | 0.817522 | NA |
| Mfumv2_1861 | 2.24356031 | 0.180999459 | 1.138405 | 0.158994 | 0.873674 | NA |
| Mfumv2_1862 | 23.69355599 | 0.06810477 | 0.367397 | 0.185371 | 0.852938 | 0.932292 |
| Mfumv2_1863 | 57.93703745 | 0.679911544 | 0.264863 | 2.56703 | 0.010257 | 0.05217 |
| Mfumv2_1864 | 28.99443333 | 0.344565403 | 0.374004 | 0.921287 | 0.356901 | 0.59836 |
| Mfumv2_1865 | 13.05052605 | 0.307219552 | 0.514529 | 0.597089 | 0.550448 | NA |
| Mfumv2_1866 | 25.67807593 | 0.652740319 | 0.363443 | 1.79599 | 0.072496 | 0.215979 |
| Mfumv2_1867 | 20.2326818 | 0.172249894 | 0.434432 | 0.396495 | 0.69174 | NA |
| Mfumv2_1868 | 9.548388458 | 0.317553231 | 0.572412 | 0.554763 | 0.579056 | NA |
| Mfumv2_1869 | 6.288657264 | -0.231176843 | 0.686129 | -0.33693 | 0.736171 | NA |
| Mfumv2_1870 | 32.92064503 | 0.303840307 | 0.327237 | 0.928501 | 0.353148 | 0.597703 |
| Mfumv2_1871 | 19.21317679 | -0.049261743 | 0.422034 | -0.11672 | 0.907078 | NA |
| Mfumv2_1872 | 7.838665033 | 0.434713899 | 0.643938 | 0.675087 | 0.499621 | NA |
| Mfumv2_1873 | 30.56858308 | 0.084349482 | 0.339098 | 0.248746 | 0.803557 | 0.903963 |
| Mfumv2_1874 | 17.04283942 | 0.210265124 | 0.430306 | 0.488641 | 0.625096 | NA |
| Mfumv2_1875 | 20.66626983 | 0.69418927 | 0.404063 | 1.718022 | 0.085793 | NA |
| Mfumv2_1876 | 14.37932664 | 0.491927987 | 0.466457 | 1.054606 | 0.291606 | NA |
| Mfumv2_1877 | 101.2475219 | 0.247559279 | 0.194896 | 1.27021 | 0.20401 | 0.433912 |
| Mfumv2_1878 | 64.12991295 | 0.443797162 | 0.251314 | 1.765906 | 0.077412 | 0.226706 |
| Mfumv2_1879 | 96.61022714 | -0.001305819 | 0.20083 | -0.0065 | 0.994812 | 0.997294 |
| Mfumv2_1881 | 15.71298063 | 0.059408877 | 0.463044 | 0.128301 | 0.897911 | NA |
| Mfumv2_1882 | 314.8446118 | -0.242347169 | 0.138212 | -1.75345 | 0.079525 | 0.230211 |
| Mfumv2_1883 | 36.09761128 | 0.242040138 | 0.300932 | 0.804303 | 0.421222 | 0.654474 |
| Mfumv2_1884 | 16.64613731 | 0.040988455 | 0.433945 | 0.094455 | 0.924747 | NA |
| Mfumv2_1885 | 201.5514753 | 0.212624696 | 0.154089 | 1.379881 | 0.167623 | 0.378376 |
| Mfumv2_1886 | 340.2264484 | 0.097692994 | 0.147251 | 0.663447 | 0.507044 | 0.732316 |
| Mfumv2_1887 | 199.4618157 | -0.032336073 | 0.149354 | -0.21651 | 0.828593 | 0.915646 |
| Mfumv2_1888 | 503.5291016 | -0.079720512 | 0.150748 | -0.52883 | 0.596921 | 0.798412 |
| Mfumv2_1889 | 2.51237285 | 0.713650636 | 1.163059 | 0.613598 | 0.539481 | NA |
| Mfumv2_1890 | 4.74931977 | -0.036622565 | 0.794307 | -0.04611 | 0.963226 | NA |
| Mfumv2_1891 | 1.455975714 | 0.352387148 | 1.432384 | 0.246014 | 0.805671 | NA |
| Mfumv2_1892 | 5.902464206 | 1.440075363 | 0.796327 | 1.808397 | 0.070545 | NA |
| Mfumv2_1893 | 5.545203941 | 0.42290184 | 0.76374 | 0.553725 | 0.579767 | NA |
| Mfumv2_1894 | 528.3456491 | -0.153897177 | 0.120718 | -1.27485 | 0.202362 | 0.43339 |
| Mfumv2_1895 | 115.5622171 | 0.243937691 | 0.200563 | 1.216263 | 0.223885 | 0.45844 |
| Mfumv2_1896 | 71.13890228 | 0.219940153 | 0.232512 | 0.945929 | 0.344185 | 0.587483 |
| Mfumv2_1897 | 3.35302641 | 0.501495327 | 0.987141 | 0.508028 | 0.611434 | NA |
| Mfumv2_1898 | 2.587568196 | 0.686537415 | 1.13929 | 0.602601 | 0.546774 | NA |
| Mfumv2_1899 | 6.367815405 | 1.288819211 | 0.745482 | 1.728841 | 0.083838 | NA |
| Mfumv2_1900 | 1.997191089 | 0.965640822 | 1.315302 | 0.734159 | 0.462852 | NA |
| Mfumv2_1901 | 2.500863977 | 1.689270082 | 1.247995 | 1.353587 | 0.175868 | NA |
| Mfumv2_1902 | 11.35227324 | -0.355306187 | 0.528407 | -0.67241 | 0.501323 | NA |
| Mfumv2_1903 | 0 | NA | NA | NA | NA | NA |
| Mfumv2_1904 | 73.90215667 | 0.039519499 | 0.223157 | 0.177092 | 0.859436 | 0.936499 |
| Mfumv2_1905 | 213.4737103 | 0.118022963 | 0.150621 | 0.783575 | 0.43329 | 0.666013 |
| Mfumv2_1906 | 274.6272372 | 0.149969326 | 0.145982 | 1.027314 | 0.304273 | 0.545789 |

| | | | | | | |
|---|---|---|---|---|---|---|
| Mfumv2_1907 | 887.2492314 | -0.178398482 | 0.134172 | -1.32963 | 0.183642 | 0.403743 |
| Mfumv2_1908 | 2352.451379 | -0.390770132 | 0.10973 | -3.56118 | 0.000369 | 0.003843 |
| Mfumv2_1909 | 179.1423152 | -0.140339981 | 0.159613 | -0.87925 | 0.379266 | 0.613977 |
| Mfumv2_1910 | 908.7777796 | -0.282460407 | 0.116368 | -2.42731 | 0.015211 | 0.070091 |
| Mfumv2_1911 | 0.138978706 | 0.283845181 | 4.080473 | 0.069562 | 0.944542 | NA |
| Mfumv2_1912 | 1137.357295 | 0.015778933 | 0.106637 | 0.147969 | 0.882367 | 0.946455 |
| Mfumv2_1913 | 1056.370616 | -0.30866402 | 0.126219 | -2.44547 | 0.014466 | 0.067588 |
| Mfumv2_1914 | 5841.029633 | 0.001052325 | 0.090999 | 0.011564 | 0.990773 | 0.995243 |
| Mfumv2_1915 | 294.0812475 | -0.039985838 | 0.132297 | -0.30224 | 0.762467 | 0.885944 |
| Mfumv2_1916 | 154.2231412 | 0.002671277 | 0.16249 | 0.01644 | 0.986884 | 0.994308 |
| Mfumv2_1917 | 229.4375737 | 0.264817676 | 0.171232 | 1.546541 | 0.121974 | 0.305925 |
| Mfumv2_1918 | 18.06515367 | -0.182525702 | 0.422233 | -0.43229 | 0.665533 | NA |
| Mfumv2_1919 | 37.59847196 | 0.133097617 | 0.298586 | 0.445759 | 0.655771 | 0.832295 |
| Mfumv2_1920 | 198.326272 | 0.259041381 | 0.160444 | 1.614528 | 0.106413 | 0.283025 |
| Mfumv2_1921 | 345.890306 | -0.493702752 | 0.166949 | -2.95721 | 0.003104 | 0.020315 |
| Mfumv2_1922 | 360.9077375 | 0.21919998 | 0.127283 | 1.722152 | 0.085042 | 0.241654 |
| Mfumv2_1923 | 2145.590804 | -0.097183047 | 0.099541 | -0.97631 | 0.328912 | 0.572108 |
| Mfumv2_1924 | 318.8943166 | -0.06087102 | 0.140925 | -0.43194 | 0.665785 | 0.835454 |
| Mfumv2_1925 | 282.7308531 | -0.01320782 | 0.141762 | -0.09317 | 0.92577 | 0.972211 |
| Mfumv2_1926 | 649.4287 | -0.131792923 | 0.113971 | -1.15637 | 0.247528 | 0.486579 |
| Mfumv2_1927 | 397.2128783 | 0.064732762 | 0.135901 | 0.476322 | 0.633845 | 0.822606 |
| Mfumv2_1928 | 234.4874011 | -0.053047576 | 0.164014 | -0.32343 | 0.746368 | 0.878414 |
| Mfumv2_1929 | 511.2242759 | -0.302035379 | 0.116874 | -2.58428 | 0.009758 | 0.050397 |
| Mfumv2_1930 | 780.1270619 | -0.379199944 | 0.116911 | -3.24349 | 0.001181 | 0.009451 |
| Mfumv2_1931 | 1347.729977 | -0.215328988 | 0.10444 | -2.06174 | 0.039232 | 0.138277 |
| Mfumv2_1932 | 261.1311047 | -0.045520806 | 0.137082 | -0.33207 | 0.739837 | 0.875859 |
| Mfumv2_1933 | 828.8194328 | -0.349189464 | 0.139229 | -2.50803 | 0.012141 | 0.059057 |
| Mfumv2_1934 | 1926.025512 | -0.072979674 | 0.124557 | -0.58591 | 0.557933 | 0.771231 |
| Mfumv2_1935 | 77.51385279 | -0.076370142 | 0.229113 | -0.33333 | 0.738885 | 0.875764 |
| Mfumv2_1936 | 426.7970676 | -0.541445576 | 0.169498 | -3.19441 | 0.001401 | 0.010868 |
| Mfumv2_1937 | 611.005273 | -0.288242682 | 0.114892 | -2.50882 | 0.012113 | 0.059057 |
| Mfumv2_1938 | 85.68768835 | -0.000605527 | 0.20964 | -0.00289 | 0.997695 | 0.99869 |
| Mfumv2_1939 | 258.9678274 | 0.310482826 | 0.144398 | 2.150186 | 0.031541 | 0.117342 |
| Mfumv2_1940 | 137.1165416 | 0.072322592 | 0.191766 | 0.377139 | 0.70607 | 0.855161 |
| Mfumv2_1941 | 225.8044761 | -0.139644396 | 0.149816 | -0.9321 | 0.351283 | 0.595048 |
| Mfumv2_1942 | 60.60735564 | -0.024874198 | 0.247912 | -0.10033 | 0.920079 | 0.9689 |
| Mfumv2_1943 | 46.07253929 | 0.211381258 | 0.27251 | 0.775682 | 0.437937 | 0.668552 |
| Mfumv2_1944 | 106.4765895 | -0.220922591 | 0.225049 | -0.98167 | 0.326265 | 0.568981 |
| Mfumv2_1945 | 205.1558215 | -0.640869053 | 0.175117 | -3.65966 | 0.000253 | 0.002757 |
| Mfumv2_1946 | 34.77196049 | 0.074326829 | 0.312129 | 0.238129 | 0.811781 | 0.907551 |
| Mfumv2_1947 | 71.5165658 | 0.010070261 | 0.223161 | 0.045126 | 0.964007 | 0.988107 |
| Mfumv2_1948 | 31.70774115 | 0.413232845 | 0.332614 | 1.242378 | 0.214097 | 0.444857 |
| Mfumv2_1949 | 29.75330146 | 0.131441621 | 0.338888 | 0.387862 | 0.698118 | 0.850369 |
| Mfumv2_1950 | 1513.475748 | 3.975286808 | 0.136971 | 29.02275 | 3.40E-185 | 3.41E-182 |
| Mfumv2_1951 | 64.43820515 | 3.278251331 | 0.332305 | 9.865178 | 5.89E-23 | 6.23E-21 |
| Mfumv2_1952 | 0.536933403 | 2.200609238 | 2.28148 | 0.964553 | 0.334769 | NA |
| Mfumv2_1953 | 1.073573091 | -0.259054255 | 1.730862 | -0.14967 | 0.881027 | NA |
| Mfumv2_1954 | 0.385021631 | -0.677928376 | 2.88264 | -0.23518 | 0.814072 | NA |
| Mfumv2_1955 | 16.13257764 | 0.478895329 | 0.489197 | 0.978941 | 0.327609 | NA |
| Mfumv2_1956 | 101.5418003 | 0.465618454 | 0.244764 | 1.902318 | 0.05713 | 0.181031 |
| Mfumv2_1958 | 68.95851329 | 0.822711777 | 0.239222 | 3.439111 | 0.000584 | 0.005477 |
| Mfumv2_1959 | 496.500078 | 0.162553926 | 0.13581 | 1.196923 | 0.231336 | 0.468609 |

| | | | | | | |
|---|---|---|---|---|---|---|
| Mfumv2_1960 | 3.108139186 | 0.705700632 | 1.012937 | 0.696687 | 0.485998 | NA |
| Mfumv2_1961 | 109.3389798 | 0.475779967 | 0.204674 | 2.32457 | 0.020095 | 0.086751 |
| Mfumv2_1962 | 87.42710774 | -0.06763977 | 0.217603 | -0.31084 | 0.755922 | 0.884498 |
| Mfumv2_1963 | 3.479630585 | -0.689704872 | 0.985828 | -0.69962 | 0.484165 | NA |
| Mfumv2_1964 | 22.82309015 | 0.006033764 | 0.387352 | 0.015577 | 0.987572 | NA |
| Mfumv2_1965 | 165.7988858 | 0.448781374 | 0.171324 | 2.619487 | 0.008806 | 0.046658 |
| Mfumv2_1966 | 32.26646097 | 0.321892277 | 0.325221 | 0.989766 | 0.322289 | 0.566462 |
| Mfumv2_1967 | 65.97585556 | 0.265971805 | 0.240526 | 1.105792 | 0.268817 | 0.507699 |
| Mfumv2_1968 | 642.4826036 | -0.103310295 | 0.118378 | -0.87272 | 0.382818 | 0.616745 |
| Mfumv2_1969 | 359.8326409 | 0.09982313 | 0.136998 | 0.728646 | 0.466219 | 0.694814 |
| Mfumv2_1970 | 281.0245926 | 0.02294662 | 0.135321 | 0.169572 | 0.865347 | 0.938198 |
| Mfumv2_1971 | 189.9153175 | 0.105897509 | 0.174623 | 0.606434 | 0.544227 | 0.762981 |
| Mfumv2_1972 | 175.8814476 | -0.011648996 | 0.162777 | -0.07156 | 0.942949 | 0.978504 |
| Mfumv2_1973 | 150.3190239 | 0.096362672 | 0.178838 | 0.538827 | 0.590006 | 0.793808 |
| Mfumv2_1974 | 202.2353906 | 0.123339269 | 0.158212 | 0.779583 | 0.435636 | 0.666666 |
| Mfumv2_1975 | 70.07961006 | -0.37075534 | 0.233297 | -1.5892 | 0.112016 | 0.291291 |
| Mfumv2_1976 | 61.64457684 | -0.586281386 | 0.246585 | -2.37761 | 0.017425 | 0.077795 |
| Mfumv2_1977 | 25.98829227 | -0.062264337 | 0.358104 | -0.17387 | 0.861966 | 0.93693 |
| Mfumv2_1978 | 200.2581957 | 0.034600202 | 0.150371 | 0.230099 | 0.818015 | 0.909463 |
| Mfumv2_1979 | 194.7438337 | -0.020837389 | 0.153842 | -0.13545 | 0.892259 | 0.94894 |
| Mfumv2_1980 | 296.198653 | 0.057366389 | 0.150933 | 0.380079 | 0.703887 | 0.854415 |
| Mfumv2_1981 | 189.3548901 | 0.068658753 | 0.158185 | 0.43404 | 0.664259 | 0.835454 |
| Mfumv2_1982 | 4.899097777 | 0.08658438 | 0.973915 | 0.088903 | 0.929159 | NA |
| Mfumv2_1983 | 12.24731082 | 0.128780617 | 0.544423 | 0.236545 | 0.81301 | NA |
| Mfumv2_1984 | 46.32376549 | 0.753990365 | 0.28252 | 2.6688 | 0.007612 | 0.041557 |
| Mfumv2_1985 | 128.0894521 | 0.132796874 | 0.183164 | 0.725016 | 0.468442 | 0.697111 |
| Mfumv2_1986 | 207.9024401 | -0.317158859 | 0.148535 | -2.13524 | 0.032741 | 0.120913 |
| Mfumv2_1987 | 144.1567057 | -0.675581774 | 0.238273 | -2.83533 | 0.004578 | 0.027954 |
| Mfumv2_1988 | 1356.133995 | -0.141644992 | 0.11092 | -1.277 | 0.201602 | 0.432711 |
| Mfumv2_1989 | 1580.525625 | 0.041123481 | 0.094165 | 0.436716 | 0.662317 | 0.835454 |
| Mfumv2_1990 | 384.1559311 | 1.832975773 | 0.147258 | 12.44737 | 1.45E-35 | 2.42E-33 |
| Mfumv2_1991 | 516.2424505 | 1.29291893 | 0.137713 | 9.388474 | 6.09E-21 | 5.82E-19 |
| Mfumv2_1992 | 581.981346 | 2.34759016 | 0.156492 | 15.00131 | 7.20E-51 | 2.07E-48 |
| Mfumv2_1993 | 672.5056246 | 0.020031544 | 0.131688 | 0.152114 | 0.879097 | 0.945691 |
| Mfumv2_1994 | 113.4258629 | 0.370611209 | 0.183781 | 2.016592 | 0.043738 | 0.149185 |
| Mfumv2_1995 | 880.5431524 | -0.032992858 | 0.112057 | -0.29443 | 0.76843 | 0.887638 |
| Mfumv2_1996 | 988.0578493 | -0.305656042 | 0.112079 | -2.72716 | 0.006388 | 0.035849 |
| Mfumv2_1997 | 1142.187532 | -0.097020777 | 0.113224 | -0.85689 | 0.391504 | 0.624232 |
| Mfumv2_1998 | 1091.248252 | -0.056921036 | 0.112728 | -0.50494 | 0.6136 | 0.808872 |
| Mfumv2_1999 | 184.9507549 | 0.181116501 | 0.165998 | 1.091077 | 0.275239 | 0.516133 |
| Mfumv2_2000 | 194.7599179 | 0.243971452 | 0.162174 | 1.504379 | 0.132484 | 0.320289 |
| Mfumv2_2001 | 0.677623517 | 2.533666148 | 2.134614 | 1.186943 | 0.23525 | NA |
| Mfumv2_2002 | 0.674710794 | -0.558377147 | 2.238682 | -0.24942 | 0.803034 | NA |
| Mfumv2_2003 | 156.7603559 | -0.296354993 | 0.229494 | -1.29134 | 0.196585 | 0.42558 |
| Mfumv2_2004 | 1772.644436 | -0.350027787 | 0.101773 | -3.43928 | 0.000583 | 0.005477 |
| Mfumv2_2006 | 1027.218336 | -0.389167595 | 0.105422 | -3.69152 | 0.000223 | 0.002502 |
| Mfumv2_2007 | 648.4624173 | -0.028653258 | 0.118982 | -0.24082 | 0.809695 | 0.907508 |
| Mfumv2_2008 | 346.0288158 | -0.345729991 | 0.131147 | -2.63619 | 0.008384 | 0.044798 |
| Mfumv2_2009 | 214.4208931 | 0.234483493 | 0.159019 | 1.474567 | 0.140329 | 0.331845 |
| Mfumv2_2010 | 195.9435941 | -0.072740776 | 0.151236 | -0.48098 | 0.630533 | 0.82074 |
| Mfumv2_2011 | 1777.753128 | -0.258797043 | 0.099725 | -2.59511 | 0.009456 | 0.049472 |
| Mfumv2_2012 | 520.137279 | 0.239528239 | 0.133246 | 1.797634 | 0.072235 | 0.215979 |

| | | | | | | |
|---|---|---|---|---|---|---|
| Mfumv2_2013 | 41.285259 | -0.134131812 | 0.295419 | -0.45404 | 0.649801 | 0.829769 |
| Mfumv2_2014 | 734.8478942 | -0.036324711 | 0.123901 | -0.29318 | 0.769388 | 0.887638 |
| Mfumv2_2015 | 230.9462182 | -0.109386636 | 0.146062 | -0.7489 | 0.453915 | 0.683083 |
| Mfumv2_2016 | 637.9769321 | 0.086425688 | 0.120403 | 0.717803 | 0.472879 | 0.700084 |
| Mfumv2_2017 | 1028.19701 | 0.006183509 | 0.106687 | 0.05796 | 0.953781 | 0.981632 |
| Mfumv2_2018 | 1720.46528 | -0.245731961 | 0.108849 | -2.25755 | 0.023973 | 0.098091 |
| Mfumv2_2019 | 854.8705555 | 0.259468556 | 0.11509 | 2.25449 | 0.024165 | 0.098475 |
| Mfumv2_2020 | 252.0454332 | 0.501387975 | 0.201945 | 2.482798 | 0.013035 | 0.062689 |
| Mfumv2_2021 | 63.15990736 | 0.332102954 | 0.263332 | 1.261156 | 0.207253 | 0.437364 |
| Mfumv2_2022 | 488.5369414 | -0.248097806 | 0.115756 | -2.14328 | 0.032091 | 0.118968 |
| Mfumv2_2023 | 1427.606399 | -0.288170709 | 0.111092 | -2.59398 | 0.009487 | 0.049506 |
| Mfumv2_2024 | 765.8604357 | 0.52421258 | 0.108605 | 4.826765 | 1.39E-06 | 2.93E-05 |
| Mfumv2_2025 | 257.2655692 | 0.410272748 | 0.154553 | 2.654569 | 0.007941 | 0.043001 |
| Mfumv2_2026 | 6.526255901 | -0.067784456 | 0.68368 | -0.09915 | 0.921022 | NA |
| Mfumv2_2027 | 161.951578 | 0.045451586 | 0.162623 | 0.279491 | 0.779868 | 0.893756 |
| Mfumv2_2028 | 462.8566745 | 0.076854079 | 0.123887 | 0.620357 | 0.535023 | 0.756944 |
| Mfumv2_2030 | 238.8151192 | -0.138256139 | 0.162703 | -0.84975 | 0.395466 | 0.627515 |
| Mfumv2_2031 | 554.5375096 | -0.015210503 | 0.111347 | -0.1366 | 0.891344 | 0.948469 |
| Mfumv2_2032 | 425.7900509 | -0.006425995 | 0.11809 | -0.05442 | 0.956604 | 0.982524 |
| Mfumv2_2033 | 332.6148671 | -0.076676569 | 0.170896 | -0.44867 | 0.653667 | 0.832096 |
| Mfumv2_2034 | 11.35560869 | -0.066832237 | 0.529218 | -0.12628 | 0.899506 | NA |
| Mfumv2_2035 | 521.1593807 | 0.189052301 | 0.125092 | 1.511304 | 0.130711 | 0.318384 |
| Mfumv2_2036 | 349.6223561 | -0.851362597 | 0.191163 | -4.4536 | 8.44E-06 | 0.000157 |
| Mfumv2_2037 | 1600.721522 | -0.183208747 | 0.11753 | -1.55882 | 0.119038 | 0.303487 |
| Mfumv2_2038 | 1635.667802 | -0.193549888 | 0.119779 | -1.61589 | 0.106118 | 0.282747 |
| Mfumv2_2039 | 625.0803057 | 0.183352199 | 0.122322 | 1.498928 | 0.133892 | 0.322143 |
| Mfumv2_2040 | 66.87816108 | 0.520084978 | 0.252435 | 2.060275 | 0.039372 | 0.138299 |
| Mfumv2_2041 | 1.684778378 | -0.621196683 | 1.381333 | -0.44971 | 0.652921 | NA |
| Mfumv2_2042 | 345.7218205 | -0.521255931 | 0.12712 | -4.10051 | 4.12E-05 | 0.000624 |
| Mfumv2_2043 | 497.607562 | -1.107647655 | 0.121129 | -9.14437 | 6.00E-20 | 5.02E-18 |
| Mfumv2_2044 | 62.1014227 | -0.104332025 | 0.301956 | -0.34552 | 0.729703 | 0.871376 |
| Mfumv2_2045 | 32.75371647 | 0.381924027 | 0.32596 | 1.17169 | 0.241322 | 0.480964 |
| Mfumv2_2046 | 778.9317704 | -0.015466536 | 0.110601 | -0.13984 | 0.888785 | 0.94819 |
| Mfumv2_2047 | 735.8580547 | 0.40005835 | 0.115223 | 3.472024 | 0.000517 | 0.004965 |
| Mfumv2_2048 | 60.78502652 | 0.011626512 | 0.239747 | 0.048495 | 0.961322 | 0.986361 |
| Mfumv2_2049 | 270.9876923 | 0.169406222 | 0.145243 | 1.166366 | 0.243466 | 0.48259 |
| Mfumv2_2050 | 94.05599094 | -0.272964574 | 0.204622 | -1.33399 | 0.182206 | 0.402255 |
| Mfumv2_2051 | 430.5930466 | -0.308271944 | 0.119771 | -2.57384 | 0.010058 | 0.051427 |
| Mfumv2_2052 | 694.228711 | -0.102387632 | 0.113961 | -0.89845 | 0.368948 | 0.607278 |
| Mfumv2_2053 | 422.935261 | -0.021378198 | 0.123423 | -0.17321 | 0.862485 | 0.93693 |
| Mfumv2_2054 | 1.078936723 | 0.555387907 | 1.863775 | 0.297991 | 0.76571 | NA |
| Mfumv2_2055 | 16.89643927 | -0.48804067 | 0.447436 | -1.09075 | 0.275383 | NA |
| Mfumv2_2056 | 2.020868304 | 1.41184636 | 1.410368 | 1.001048 | 0.316803 | NA |
| Mfumv2_2057 | 2.46210386 | 1.02203479 | 1.154634 | 0.885159 | 0.376071 | NA |
| Mfumv2_2058 | 118.9385944 | 0.608861875 | 0.194596 | 3.128859 | 0.001755 | 0.012929 |
| Mfumv2_2059 | 60.92733722 | 0.06019531 | 0.247416 | 0.243296 | 0.807776 | 0.907112 |
| Mfumv2_2060 | 62.4421671 | -0.016350951 | 0.240491 | -0.06799 | 0.945794 | 0.979433 |
| Mfumv2_2061 | 101.0216894 | 0.006996045 | 0.19087 | 0.036653 | 0.970761 | 0.98916 |
| Mfumv2_2062 | 74.16647941 | 0.085603573 | 0.22752 | 0.376246 | 0.706734 | 0.855281 |
| Mfumv2_2063 | 117.3306084 | -0.297856957 | 0.195822 | -1.52106 | 0.128244 | 0.314967 |
| Mfumv2_2064 | 570.5376814 | -0.658027964 | 0.116467 | -5.6499 | 1.61E-08 | 4.89E-07 |
| Mfumv2_2065 | 83.42864804 | -1.316467967 | 0.227431 | -5.78844 | 7.10E-09 | 2.46E-07 |

| | | | | | | |
|---|---|---|---|---|---|---|
| Mfumv2_2066 | 61.10255901 | 0.10154846 | 0.247442 | 0.410392 | 0.681518 | 0.843309 |
| Mfumv2_2067 | 371.9312937 | -0.074365642 | 0.126897 | -0.58603 | 0.557856 | 0.771231 |
| Mfumv2_2068 | 214.8291227 | -0.060325044 | 0.15336 | -0.39336 | 0.694056 | 0.849183 |
| Mfumv2_2070 | 95.899487 | 0.341204798 | 0.212232 | 1.607696 | 0.107902 | 0.285606 |
| Mfumv2_2073 | 27.07283888 | 0.343479187 | 0.365844 | 0.938868 | 0.347799 | 0.59114 |
| Mfumv2_2074 | 50.32265295 | 0.485407345 | 0.265668 | 1.82712 | 0.067682 | 0.206332 |
| Mfumv2_2075 | 70.21510729 | -0.508661819 | 0.265687 | -1.91451 | 0.055555 | 0.177416 |
| Mfumv2_2076 | 82.07667166 | 0.088968858 | 0.227809 | 0.390541 | 0.696137 | 0.850369 |
| Mfumv2_2077 | 233.3756169 | -0.12015499 | 0.153232 | -0.78414 | 0.432958 | 0.666013 |
| Mfumv2_2078 | 416.4828762 | -0.033618742 | 0.134414 | -0.25011 | 0.8025 | 0.903903 |
| Mfumv2_2079 | 162.6582539 | 0.093943085 | 0.190326 | 0.493589 | 0.621596 | 0.816201 |
| Mfumv2_2080 | 15.46464305 | -0.187936796 | 0.453127 | -0.41476 | 0.678321 | NA |
| Mfumv2_2081 | 46.92223 | 0.051088097 | 0.291298 | 0.175381 | 0.86078 | 0.936786 |
| Mfumv2_2082 | 689.5898767 | -0.016273879 | 0.110206 | -0.14767 | 0.882605 | 0.946455 |
| Mfumv2_2083 | 1103.764596 | 0.006454273 | 0.108472 | 0.059502 | 0.952552 | 0.981373 |
| Mfumv2_2084 | 282.5250284 | -0.143519523 | 0.164391 | -0.87304 | 0.382641 | 0.616745 |
| Mfumv2_2085 | 842.6391341 | -0.301132423 | 0.127089 | -2.36947 | 0.017814 | 0.079177 |
| Mfumv2_2086 | 16.4152278 | -1.071441318 | 0.446017 | -2.40225 | 0.016295 | NA |
| Mfumv2_2087 | 83040.51821 | -0.619191554 | 0.120368 | -5.14416 | 2.69E-07 | 6.43E-06 |
| Mfumv2_2088 | 611.69895 | -0.231219893 | 0.117799 | -1.96283 | 0.049666 | 0.16465 |
| Mfumv2_2089 | 522.6261809 | -0.14295288 | 0.122841 | -1.16372 | 0.244537 | 0.483537 |
| Mfumv2_2090 | 2195.574992 | -0.020032324 | 0.092366 | -0.21688 | 0.828302 | 0.915646 |
| Mfumv2_2091 | 0.281380227 | 1.260342684 | 3.513739 | 0.35869 | 0.719827 | NA |
| Mfumv2_2093 | 39.25174945 | -0.176150624 | 0.37838 | -0.46554 | 0.641545 | 0.825137 |
| Mfumv2_2094 | 1200.450112 | -0.104553914 | 0.112041 | -0.93318 | 0.350728 | 0.594609 |
| Mfumv2_2095 | 3301.204642 | 0.010356642 | 0.133963 | 0.07731 | 0.938377 | 0.977463 |
| Mfumv2_2096 | 2679.688559 | -0.499256084 | 0.115048 | -4.33954 | 1.43E-05 | 0.000249 |
| Mfumv2_2097 | 268.4376593 | -0.361987986 | 0.143194 | -2.52795 | 0.011473 | 0.056532 |
| Mfumv2_2098 | 159.8288056 | -0.430662229 | 0.214964 | -2.00342 | 0.045132 | 0.151948 |
| Mfumv2_2099 | 183.0780107 | -0.240454439 | 0.165604 | -1.45198 | 0.146507 | 0.340268 |
| Mfumv2_2100 | 0.96697052 | -0.761233215 | 1.93281 | -0.39385 | 0.693693 | NA |
| Mfumv2_2101 | 3.023152906 | 1.429614946 | 1.068906 | 1.337456 | 0.181074 | NA |
| Mfumv2_2102 | 22.55918975 | 0.053817362 | 0.398533 | 0.135039 | 0.892581 | NA |
| Mfumv2_2103 | 1186.16467 | -0.032055795 | 0.116232 | -0.27579 | 0.782708 | 0.895479 |
| Mfumv2_2104 | 555.8868159 | -0.522095281 | 0.136481 | -3.8254 | 0.000131 | 0.00159 |
| Mfumv2_2105 | 589.7195053 | -0.094599845 | 0.120276 | -0.78652 | 0.431563 | 0.664885 |
| Mfumv2_2106 | 233.6625413 | -0.068003345 | 0.144986 | -0.46903 | 0.639045 | 0.823866 |
| Mfumv2_2107 | 108.8482592 | -0.01570969 | 0.191759 | -0.08192 | 0.934707 | 0.975484 |
| Mfumv2_2109 | 155.6687708 | 0.043915748 | 0.176608 | 0.248662 | 0.803622 | 0.903963 |
| Mfumv2_2110 | 128.6335765 | 0.533891296 | 0.187506 | 2.847322 | 0.004409 | 0.027254 |
| Mfumv2_2111 | 346.7199977 | 0.166442819 | 0.143358 | 1.161027 | 0.245631 | 0.48434 |
| Mfumv2_2112 | 87.94028172 | 0.393560662 | 0.210808 | 1.866917 | 0.061913 | 0.191971 |
| Mfumv2_2113 | 138.8938334 | 0.058075128 | 0.169225 | 0.343183 | 0.731461 | 0.871592 |
| Mfumv2_2114 | 208.1147397 | -0.317115569 | 0.152682 | -2.07697 | 0.037804 | 0.135864 |
| Mfumv2_2115 | 138.5200353 | 0.08118243 | 0.19686 | 0.412387 | 0.680056 | 0.842824 |
| Mfumv2_2116 | 35.58417198 | 0.201463565 | 0.337087 | 0.59766 | 0.550067 | 0.76593 |
| Mfumv2_2117 | 97.86407614 | 0.479737212 | 0.212071 | 2.262158 | 0.023688 | 0.09769 |
| Mfumv2_2118 | 162.3998279 | 0.024181278 | 0.172442 | 0.140228 | 0.88848 | 0.94819 |
| Mfumv2_2119 | 1093.630423 | -0.369703372 | 0.109634 | -3.37217 | 0.000746 | 0.006628 |
| Mfumv2_2120 | 643.8206503 | 0.104606048 | 0.138291 | 0.756418 | 0.449399 | 0.67934 |
| Mfumv2_2121 | 78.57455855 | 0.342146323 | 0.217586 | 1.572465 | 0.115843 | 0.297606 |
| Mfumv2_2122 | 182.180758 | 0.201207037 | 0.178974 | 1.124224 | 0.260918 | 0.499223 |

| | | | | | | |
|---|---|---|---|---|---|---|
| Mfumv2_2123 | 360.6413855 | 0.198335321 | 0.129539 | 1.531083 | 0.125749 | 0.311041 |
| Mfumv2_2124 | 490.5415751 | -0.05948405 | 0.127758 | -0.4656 | 0.641503 | 0.825137 |
| Mfumv2_2125 | 196.2272968 | 0.361694149 | 0.150619 | 2.401381 | 0.016333 | 0.073776 |
| Mfumv2_2127 | 1.096781134 | 0.774446423 | 1.955339 | 0.396068 | 0.692055 | NA |
| Mfumv2_2128 | 4.427800817 | 0.84103679 | 0.860518 | 0.977361 | 0.32839 | NA |
| Mfumv2_2130 | 389.6757729 | -0.914887054 | 0.149695 | -6.11169 | 9.86E-10 | 4.04E-08 |
| Mfumv2_2131 | 315.8971742 | -0.894248814 | 0.139102 | -6.42871 | 1.29E-10 | 6.01E-09 |
| Mfumv2_2132 | 241.2394961 | -0.776265961 | 0.142859 | -5.43378 | 5.52E-08 | 1.56E-06 |
| Mfumv2_2133 | 591.2094664 | -0.728556265 | 0.117053 | -6.22416 | 4.84E-10 | 2.11E-08 |
| Mfumv2_2134 | 274.8226999 | -0.924117231 | 0.157796 | -5.85641 | 4.73E-09 | 1.73E-07 |
| Mfumv2_2135 | 402.1503031 | -0.861138631 | 0.125494 | -6.86198 | 6.79E-12 | 3.50E-10 |
| Mfumv2_2136 | 5.333076335 | -0.802689919 | 0.760442 | -1.05556 | 0.29117 | NA |
| Mfumv2_2137 | 1.583732761 | -0.336480082 | 1.434245 | -0.2346 | 0.814516 | NA |
| Mfumv2_2138 | 327.6561265 | -0.21519872 | 0.128237 | -1.67813 | 0.093322 | 0.258242 |
| Mfumv2_2139 | 117.9818935 | 0.41234838 | 0.189327 | 2.177967 | 0.029408 | 0.112536 |
| Mfumv2_2140 | 1147.819812 | -0.235832259 | 0.105175 | -2.24228 | 0.024943 | 0.100423 |
| Mfumv2_2141 | 118.2536204 | -0.0766029 | 0.189022 | -0.40526 | 0.685286 | 0.845144 |
| Mfumv2_2142 | 138.2656291 | 0.158219132 | 0.175443 | 0.901828 | 0.367148 | 0.60658 |
| Mfumv2_2143 | 69.29843778 | -0.154882976 | 0.231675 | -0.66854 | 0.503791 | 0.730096 |
| Mfumv2_2144 | 42.93785064 | 0.124234671 | 0.283796 | 0.43776 | 0.66156 | 0.835454 |
| Mfumv2_2145 | 185.6188583 | 0.017219657 | 0.173971 | 0.09898 | 0.921154 | 0.9689 |
| Mfumv2_2146 | 82.96453359 | 0.100228172 | 0.226836 | 0.441852 | 0.658596 | 0.834776 |
| Mfumv2_2147 | 7.325036933 | 0.154016617 | 0.648133 | 0.237631 | 0.812167 | NA |
| Mfumv2_2148 | 8.509468969 | 0.174808401 | 0.611017 | 0.286094 | 0.774806 | NA |
| Mfumv2_2149 | 1.6042194 | -1.147911722 | 1.497296 | -0.76666 | 0.443286 | NA |
| Mfumv2_2150 | 325.960429 | -0.236062055 | 0.199027 | -1.18608 | 0.23559 | 0.473775 |
| Mfumv2_2151 | 224.5024084 | -0.337838358 | 0.176584 | -1.91319 | 0.055724 | 0.177416 |
| Mfumv2_2152 | 153.8378496 | 0.230670162 | 0.179049 | 1.288306 | 0.197639 | 0.426944 |
| Mfumv2_2153 | 142.7599127 | 0.187783941 | 0.186225 | 1.008371 | 0.313276 | 0.555492 |
| Mfumv2_2154 | 419.3615371 | -0.567692725 | 0.138868 | -4.08801 | 4.35E-05 | 0.000643 |
| Mfumv2_2155 | 43.52983006 | -0.072399569 | 0.283911 | -0.25501 | 0.798717 | 0.902079 |
| Mfumv2_2156 | 20.82214792 | 0.283824663 | 0.410206 | 0.691907 | 0.488996 | NA |
| Mfumv2_2157 | 144.0847195 | 0.079988127 | 0.179464 | 0.445706 | 0.65581 | 0.832295 |
| Mfumv2_2158 | 748.7827081 | 0.025440661 | 0.108241 | 0.235037 | 0.81418 | 0.908704 |
| Mfumv2_2159 | 172.5513295 | -0.057792195 | 0.17825 | -0.32422 | 0.745772 | 0.878227 |
| Mfumv2_2160 | 190.3663441 | 0.015368301 | 0.179275 | 0.085725 | 0.931685 | 0.973858 |
| Mfumv2_2161 | 215.784862 | -0.260033129 | 0.162323 | -1.60195 | 0.109167 | 0.288196 |
| Mfumv2_2162 | 96.87542245 | 0.253053665 | 0.199657 | 1.26744 | 0.204998 | 0.434665 |
| Mfumv2_2163 | 151.2327143 | -0.009509009 | 0.172832 | -0.05502 | 0.956124 | 0.982524 |
| Mfumv2_2164 | 2145.9447 | -0.381569839 | 0.128811 | -2.96226 | 0.003054 | 0.020133 |
| Mfumv2_2165 | 5836.36735 | -0.311665513 | 0.13375 | -2.3302 | 0.019795 | 0.085709 |
| Mfumv2_2166 | 1549.690808 | -0.072972389 | 0.151765 | -0.48082 | 0.630641 | 0.82074 |
| Mfumv2_2167 | 153.6123876 | 0.060591531 | 0.207278 | 0.29232 | 0.770042 | 0.887638 |
| Mfumv2_2168 | 70.63130565 | -0.517844019 | 0.228021 | -2.27104 | 0.023145 | 0.096467 |
| Mfumv2_2169 | 784.1828205 | -0.196378093 | 0.10859 | -1.80844 | 0.070538 | 0.213098 |
| Mfumv2_2170 | 222.6226505 | 0.025573197 | 0.147048 | 0.173911 | 0.861936 | 0.93693 |
| Mfumv2_2171 | 366.851921 | -1.14908657 | 0.200068 | -5.74347 | 9.28E-09 | 3.11E-07 |
| Mfumv2_2172 | 28.03192692 | -1.314207384 | 0.348861 | -3.76714 | 0.000165 | 0.001907 |
| Mfumv2_2173 | 13.78525905 | -0.006052921 | 0.502451 | -0.01205 | 0.990388 | NA |
| Mfumv2_2174 | 27.17765817 | 0.59638574 | 0.420386 | 1.418662 | 0.155997 | 0.357761 |
| Mfumv2_2175 | 79.63702399 | 0.604230557 | 0.221365 | 2.729562 | 0.006342 | 0.035688 |
| Mfumv2_2176 | 240.1387785 | 0.745185335 | 0.14982 | 4.973886 | 6.56E-07 | 1.43E-05 |

| | | | | | | | |
|---|---|---|---|---|---|---|---|
| Mfumv2_2177 | 76.65927737 | | 0.113732498 | 0.566625 | 0.200719 | 0.840918 | 0.924411 |
| Mfumv2_2178 | 229.7258271 | | -0.024254508 | 0.23494 | -0.10324 | 0.917775 | 0.967421 |
| Mfumv2_2179 | 8.207468962 | | -0.285819269 | 0.617457 | -0.4629 | 0.643438 | NA |
| Mfumv2_2180 | 1219.118764 | | -0.149745607 | 0.107145 | -1.3976 | 0.162235 | 0.369116 |
| Mfumv2_2181 | 637.4621707 | | 0.511108255 | 0.172555 | 2.961998 | 0.003056 | 0.020133 |
| Mfumv2_2182 | 7.956633975 | | 0.113909518 | 0.628658 | 0.181195 | 0.856215 | NA |
| Mfumv2_2183 | 0 | NA | | NA | NA | NA | NA |
| Mfumv2_2184 | 1.625102525 | | 2.640553607 | 1.648442 | 1.601848 | 0.109189 | NA |
| Mfumv2_2185 | 636.7342954 | | 0.091335229 | 0.127932 | 0.713936 | 0.475267 | 0.702067 |
| Mfumv2_2186 | 369.1008557 | | 0.842868394 | 0.198087 | 4.255033 | 2.09E-05 | 0.000341 |
| Mfumv2_2187 | 376.6078501 | | 1.279438972 | 0.136404 | 9.379744 | 6.61E-21 | 6.04E-19 |
| Mfumv2_2188 | 76.05307692 | | 0.079081508 | 0.233012 | 0.339389 | 0.734317 | 0.872254 |
| Mfumv2_2189 | 269.0859492 | | 0.68530576 | 0.150795 | 4.544624 | 5.50E-06 | 0.000105 |
| Mfumv2_2190 | 2658.194943 | | -0.114572468 | 0.08834 | -1.29695 | 0.194648 | 0.422298 |
| Mfumv2_2191 | 143.4593307 | | 0.227481713 | 0.1867 | 1.218435 | 0.223059 | 0.457963 |
| Mfumv2_2192 | 53.12884752 | | 0.550383077 | 0.275371 | 1.998698 | 0.045641 | 0.153333 |
| Mfumv2_2194 | 1512.845757 | | -0.160820854 | 0.107103 | -1.50156 | 0.133211 | 0.321018 |
| Mfumv2_2195 | 195.2601377 | | 0.367721434 | 0.165336 | 2.22408 | 0.026143 | 0.103325 |
| Mfumv2_2196 | 478.0280921 | | -0.130087672 | 0.126795 | -1.02596 | 0.304908 | 0.546441 |
| Mfumv2_2197 | 109.9010282 | | 0.380412288 | 0.191613 | 1.985318 | 0.047109 | 0.157474 |
| Mfumv2_2198 | 159.8823954 | | -0.069434523 | 0.16073 | -0.432 | 0.665745 | 0.835454 |
| Mfumv2_2199 | 137.7691713 | | -0.000797659 | 0.186524 | -0.00428 | 0.996588 | 0.998576 |
| Mfumv2_2200 | 72.43639922 | | -0.064277158 | 0.22836 | -0.28147 | 0.778347 | 0.892523 |
| Mfumv2_2201 | 68.91942838 | | 0.269626054 | 0.227741 | 1.183917 | 0.236446 | 0.474072 |
| Mfumv2_2202 | 1528.78422 | | -0.124889038 | 0.109167 | -1.14402 | 0.252617 | 0.492248 |
| Mfumv2_2203 | 717.3963076 | | -0.150968726 | 0.13004 | -1.16094 | 0.245666 | 0.48434 |
| Mfumv2_2204 | 492.6540937 | | 0.078201304 | 0.134501 | 0.58142 | 0.560958 | 0.77267 |
| Mfumv2_2205 | 4602.509442 | | -0.376408557 | 0.132333 | -2.84441 | 0.004449 | 0.02742 |
| Mfumv2_2206 | 7622.751618 | | -0.238234679 | 0.108889 | -2.18787 | 0.028679 | 0.110166 |
| Mfumv2_2207 | 82.15028373 | | 0.389355024 | 0.240704 | 1.61757 | 0.105755 | 0.282156 |
| Mfumv2_2208 | 299.5616135 | | 0.548852586 | 0.215745 | 2.543984 | 0.01096 | 0.054771 |
| Mfumv2_2209 | 274.8080998 | | -0.007209003 | 0.144279 | -0.04997 | 0.96015 | 0.985662 |
| Mfumv2_2211 | 187.2287036 | | 0.227825882 | 0.155275 | 1.467245 | 0.14231 | 0.334337 |
| Mfumv2_2212 | 534.7062827 | | -0.056221638 | 0.124273 | -0.4524 | 0.650979 | 0.829833 |
| Mfumv2_2213 | 604.4391954 | | 0.210757354 | 0.118882 | 1.772822 | 0.076258 | 0.224309 |
| Mfumv2_2214 | 130.3521349 | | 0.040322839 | 0.188074 | 0.214399 | 0.830236 | 0.916957 |
| Mfumv2_2215 | 191.13372 | | -0.405238755 | 0.167568 | -2.41836 | 0.015591 | 0.071347 |
| Mfumv2_2216 | 285.4284064 | | -0.678035932 | 0.219213 | -3.09304 | 0.001981 | 0.014369 |
| Mfumv2_2217 | 2873.915467 | | -1.681979364 | 0.152506 | -11.0289 | 2.77E-28 | 3.71E-26 |
| Mfumv2_2218 | 81.09121841 | | -0.121571083 | 0.209255 | -0.58097 | 0.56126 | 0.77267 |
| Mfumv2_2219 | 64.46634256 | | -0.216274452 | 0.243485 | -0.88824 | 0.374409 | 0.610542 |
| Mfumv2_2220 | 1835.480488 | | -1.589355235 | 0.111313 | -14.2783 | 2.99E-46 | 6.67E-44 |
| Mfumv2_2221 | 1373.073492 | | -0.210402171 | 0.102339 | -2.05593 | 0.03979 | 0.139264 |
| Mfumv2_2222 | 422.8216411 | | 0.069573767 | 0.145136 | 0.479368 | 0.631677 | 0.82074 |
| Mfumv2_2223 | 2.381050487 | | 0.859596502 | 1.195475 | 0.719042 | 0.472115 | NA |
| Mfumv2_2224 | 1.590091519 | | 0.790625634 | 1.422627 | 0.555751 | 0.578381 | NA |
| Mfumv2_2225 | 3.101574239 | | -0.295177861 | 1.051723 | -0.28066 | 0.77897 | NA |
| Mfumv2_2226 | 24.00510559 | | 0.129809511 | 0.392749 | 0.330515 | 0.741011 | 0.876167 |
| Mfumv2_2227 | 232.1586788 | | 0.079113456 | 0.147624 | 0.53591 | 0.592021 | 0.793988 |
| Mfumv2_2228 | 1231.49263 | | -0.002363892 | 0.123989 | -0.01907 | 0.984789 | 0.992996 |
| Mfumv2_2229 | 116.2828028 | | 0.285705793 | 0.184418 | 1.54923 | 0.121326 | 0.304876 |
| Mfumv2_2230 | 32.35668939 | | 0.407677901 | 0.320852 | 1.270609 | 0.203868 | 0.433912 |

| | | | | | | |
|---|---|---|---|---|---|---|
| Mfumv2_2231 | 769.0626437 | -0.396918843 | 0.176983 | -2.24269 | 0.024917 | 0.100423 |
| Mfumv2_2232 | 308.8126766 | -0.029704784 | 0.136983 | -0.21685 | 0.828326 | 0.915646 |
| Mfumv2_2233 | 32.7544375 | 0.628144241 | 0.35625 | 1.763212 | 0.077865 | 0.227347 |
| Mfumv2_2234 | 1740.416869 | 0.346883649 | 0.142564 | 2.433174 | 0.014967 | 0.069604 |
| Mfumv2_2235 | 361.0251063 | -0.364045606 | 0.140799 | -2.58557 | 0.009722 | 0.050337 |
| Mfumv2_2236 | 1122.029156 | -0.501033971 | 0.139744 | -3.58537 | 0.000337 | 0.00354 |
| Mfumv2_2237 | 1023.598008 | -0.54091862 | 0.163969 | -3.29891 | 0.000971 | 0.008058 |
| Mfumv2_2238 | 398.1568891 | -0.546377871 | 0.185358 | -2.9477 | 0.003202 | 0.020883 |
| Mfumv2_2239 | 1008.801968 | -0.660706371 | 0.16095 | -4.10503 | 4.04E-05 | 0.00062 |
| Mfumv2_2240 | 521.2575826 | -0.713340023 | 0.180427 | -3.95362 | 7.70E-05 | 0.001024 |
| Mfumv2_2241 | 622.6683974 | -0.670622719 | 0.16971 | -3.95158 | 7.76E-05 | 0.001026 |
| Mfumv2_2242 | 32.86662501 | -0.366225788 | 0.324446 | -1.12877 | 0.258993 | 0.497912 |
| Mfumv2_2243 | 247.3213426 | -0.203615378 | 0.15075 | -1.35068 | 0.176799 | 0.392907 |
| Mfumv2_2244 | 114.3375617 | 0.112785126 | 0.210097 | 0.536825 | 0.591389 | 0.793988 |
| Mfumv2_2245 | 146.1976667 | 0.113623009 | 0.166055 | 0.684248 | 0.493818 | 0.722039 |
| Mfumv2_2246 | 154.5363322 | 0.273859512 | 0.181868 | 1.505818 | 0.132114 | 0.320165 |
| Mfumv2_2247 | 402.4576711 | -0.416625841 | 0.13317 | -3.12853 | 0.001757 | 0.012929 |
| Mfumv2_2248 | 26.4005076 | -0.597667591 | 0.36806 | -1.62383 | 0.104412 | 0.280433 |
| Mfumv2_2249 | 1338.449668 | -0.70426468 | 0.114876 | -6.13064 | 8.75E-10 | 3.66E-08 |
| Mfumv2_2250 | 122.2017645 | -0.644389068 | 0.182922 | -3.52275 | 0.000427 | 0.004312 |
| Mfumv2_2251 | 0.867914164 | -0.166206782 | 1.928429 | -0.08619 | 0.931317 | NA |
| Mfumv2_2252 | 9.761484131 | -0.443186784 | 0.566886 | -0.78179 | 0.434337 | NA |
| Mfumv2_2253 | 271.852167 | 0.272088589 | 0.134628 | 2.021034 | 0.043276 | 0.148113 |
| Mfumv2_2254 | 273.5662505 | -0.164958636 | 0.162945 | -1.01236 | 0.311366 | 0.55308 |
| Mfumv2_2255 | 43.9020893 | 0.440047094 | 0.296255 | 1.485366 | 0.137447 | 0.327557 |
| Mfumv2_2256 | 219.7518165 | 0.061210426 | 0.153118 | 0.39976 | 0.689333 | 0.846498 |
| Mfumv2_2257 | 520.1547441 | -0.143939833 | 0.124171 | -1.15921 | 0.246371 | 0.484779 |
| Mfumv2_2258 | 1086.982712 | -0.683794317 | 0.135862 | -5.03301 | 4.83E-07 | 1.08E-05 |
| Mfumv2_2259 | 78.8419997 | 0.042601273 | 0.248736 | 0.171271 | 0.86401 | 0.937762 |
| Mfumv2_2260 | 1.418927005 | -0.701757981 | 1.509631 | -0.46485 | 0.642036 | NA |
| Mfumv2_2262 | 17.7201827 | 0.302655028 | 0.441248 | 0.685906 | 0.492772 | NA |
| Mfumv2_2263 | 2.095349206 | -0.655290321 | 1.189855 | -0.55073 | 0.581818 | NA |
| Mfumv2_2264 | 0.340491013 | -0.677928304 | 2.998655 | -0.22608 | 0.821141 | NA |
| Mfumv2_2265 | 0 | NA | NA | NA | NA | NA |
| Mfumv2_2266 | 1.110812127 | -0.713548712 | 1.873844 | -0.38079 | 0.703356 | NA |
| Mfumv2_2267 | 615.6087761 | 0.647912709 | 0.157728 | 4.107778 | 3.99E-05 | 0.000617 |
| Mfumv2_2268 | 254.1831335 | 0.702877279 | 0.159687 | 4.401584 | 1.07E-05 | 0.000196 |
| Mfumv2_2269 | 20.084299 | 0.40901077 | 0.412135 | 0.992419 | 0.320993 | NA |
| Mfumv2_2270 | 119.8085071 | 0.32709612 | 0.21211 | 1.542106 | 0.123048 | 0.307411 |
| Mfumv2_2271 | 573.3267697 | -0.013980833 | 0.115861 | -0.12067 | 0.903953 | 0.957896 |
| Mfumv2_2272 | 309.968607 | 0.045989836 | 0.144269 | 0.318779 | 0.749894 | 0.880222 |
| Mfumv2_2273 | 374.3713924 | -0.023731788 | 0.137306 | -0.17284 | 0.862778 | 0.93693 |
| Mfumv2_2274 | 320.5176957 | -0.16754636 | 0.186098 | -0.90031 | 0.367953 | 0.607278 |
| Mfumv2_2275 | 395.3506873 | -0.001434908 | 0.124232 | -0.01155 | 0.990784 | 0.995243 |
| Mfumv2_2276 | 69.6523621 | 0.080664806 | 0.237698 | 0.339358 | 0.73434 | 0.872254 |
| Mfumv2_2277 | 153.2728556 | -0.144548829 | 0.172377 | -0.83856 | 0.401714 | 0.634968 |
| Mfumv2_2278 | 2456.897775 | -0.466141726 | 0.117692 | -3.96069 | 7.47E-05 | 0.001001 |
| Mfumv2_2279 | 353.8039943 | -0.348053566 | 0.157751 | -2.20635 | 0.02736 | 0.107146 |
| Mfumv2_2280 | 221.8744406 | -0.089590062 | 0.1625 | -0.55132 | 0.581411 | 0.788739 |
| Mfumv2_2281 | 903.0580565 | -0.263319784 | 0.105176 | -2.50361 | 0.012293 | 0.059655 |
| Mfumv2_2282 | 107.498495 | -0.211936901 | 0.200888 | -1.055 | 0.291425 | 0.52984 |
| Mfumv2_2283 | 212.2401628 | -0.126248128 | 0.154034 | -0.81961 | 0.412438 | 0.646829 |

| | | | | | | |
|---|---|---|---|---|---|---|
| Mfumv2_2285 | 233.6411666 | 0.270156662 | 0.150459 | 1.795547 | 0.072567 | 0.215979 |
| Mfumv2_2286 | 118.3289902 | 0.545639459 | 0.182716 | 2.986272 | 0.002824 | 0.019095 |
| Mfumv2_2287 | 0.11657447 | 0.283845181 | 4.080473 | 0.069562 | 0.944542 | NA |
| Mfumv2_2288 | 929.7466399 | -0.26240139 | 0.127629 | -2.05597 | 0.039786 | 0.139264 |
| Mfumv2_2289 | 125.142322 | 0.262970338 | 0.217781 | 1.207499 | 0.22724 | 0.463477 |
| Mfumv2_2290 | 0.829241574 | 0.418343462 | 2.049175 | 0.204152 | 0.838235 | NA |
| Mfumv2_2291 | 0.169414025 | -1.639708171 | 4.080473 | -0.40184 | 0.6878 | NA |
| Mfumv2_2292 | 429.5064567 | -0.33051483 | 0.137287 | -2.40747 | 0.016063 | 0.073013 |
| Mfumv2_2293 | 54.09160273 | 0.088974933 | 0.260292 | 0.341828 | 0.732481 | 0.872002 |
| Mfumv2_2294 | 556.8320542 | 0.076881874 | 0.116861 | 0.65789 | 0.510609 | 0.73535 |
| Mfumv2_2295 | 2.880618244 | -0.280183129 | 1.107467 | -0.25299 | 0.800272 | NA |
| Mfumv2_2296 | 76.16139844 | 0.061044768 | 0.23322 | 0.261748 | 0.793516 | 0.898632 |
| Mfumv2_2297 | 10.90371652 | 0.175969896 | 0.528933 | 0.332689 | 0.739369 | NA |
| Mfumv2_2298 | 1629.857368 | -0.204558177 | 0.102866 | -1.98859 | 0.046747 | 0.156524 |
| Mfumv2_2299 | 2700.014589 | -0.096665461 | 0.105181 | -0.91904 | 0.358075 | 0.59836 |
| Mfumv2_2300 | 183.0323851 | 0.341795069 | 0.188018 | 1.817888 | 0.069081 | 0.20941 |
| Mfumv2_2301 | 71.53472117 | 0.366167334 | 0.224483 | 1.631156 | 0.102857 | 0.278116 |
| Mfumv2_2302 | 123.5622311 | 0.321871433 | 0.185214 | 1.737837 | 0.08224 | 0.23502 |
| Mfumv2_2303 | 113.1515759 | 0.406323107 | 0.192472 | 2.111082 | 0.034765 | 0.126758 |
| Mfumv2_2304 | 1445.379876 | -0.228539276 | 0.101048 | -2.26168 | 0.023717 | 0.09769 |
| Mfumv2_2305 | 70.08497402 | -0.563643768 | 0.239961 | -2.3489 | 0.018829 | 0.082956 |
| Mfumv2_2306 | 23.066166 | 0.017431112 | 0.418985 | 0.041603 | 0.966815 | NA |
| Mfumv2_2307 | 15403.5076 | -0.529267956 | 0.166867 | -3.17179 | 0.001515 | 0.01147 |
| Mfumv2_2308 | 2060.465958 | -0.526997645 | 0.166188 | -3.17109 | 0.001519 | 0.01147 |
| Mfumv2_2309 | 9258.750971 | -0.44950492 | 0.164961 | -2.72492 | 0.006432 | 0.035992 |
| Mfumv2_2310 | 1287.929821 | -0.019021574 | 0.104905 | -0.18132 | 0.856115 | 0.934825 |
| Mfumv2_2311 | 845.2794389 | 0.468299105 | 0.109502 | 4.276613 | 1.90E-05 | 0.000312 |
| Mfumv2_2312 | 81.08344431 | -0.013677786 | 0.21083 | -0.06488 | 0.948273 | 0.980261 |
| Mfumv2_2313 | 291.3647942 | 0.052624334 | 0.144967 | 0.363008 | 0.716599 | 0.863097 |
| Mfumv2_2314 | 97.67799019 | 0.209003236 | 0.212497 | 0.98356 | 0.325332 | 0.568517 |
| Mfumv2_2315 | 1970.249337 | -0.27914064 | 0.143911 | -1.93967 | 0.052419 | 0.169856 |
| Mfumv2_2316 | 56.23250884 | -0.460185053 | 0.246501 | -1.86687 | 0.06192 | 0.191971 |
| Mfumv2_2317 | 620.7956928 | -0.554361189 | 0.155382 | -3.56772 | 0.00036 | 0.003768 |
| Mfumv2_2318 | 522.0899501 | -0.164353238 | 0.13226 | -1.24265 | 0.213997 | 0.444857 |
| Mfumv2_2319 | 299.7247258 | -0.329290658 | 0.140049 | -2.35125 | 0.01871 | 0.082614 |
| Mfumv2_2320 | 351.0367467 | -0.388155112 | 0.139489 | -2.7827 | 0.005391 | 0.031792 |
| Mfumv2_2321 | 671.2585213 | -0.622351537 | 0.107138 | -5.80887 | 6.29E-09 | 2.26E-07 |
| Mfumv2_2322 | 0.648883743 | -0.64430623 | 2.136162 | -0.30162 | 0.762943 | NA |
| Mfumv2_2323 | 342.737895 | 0.310531371 | 0.183414 | 1.69306 | 0.090444 | 0.25342 |
| Mfumv2_2324 | 29.07725767 | 0.441100639 | 0.343403 | 1.284497 | 0.198968 | 0.428432 |
| Mfumv2_2325 | 254.4140429 | 0.089960137 | 0.145348 | 0.618929 | 0.535963 | 0.75729 |
| Mfumv2_2326 | 272.3588358 | 0.284422965 | 0.139386 | 2.040548 | 0.041296 | 0.14304 |
| Mfumv2_2327 | 545.1742074 | -0.049353204 | 0.115605 | -0.42691 | 0.669442 | 0.837428 |
| Mfumv2_2328 | 441.1463931 | -0.135365432 | 0.120371 | -1.12457 | 0.260773 | 0.499223 |
| Mfumv2_2329 | 114.8999769 | -0.296003798 | 0.181849 | -1.62774 | 0.10358 | 0.278943 |
| Mfumv2_2330 | 62.04494271 | 0.408096933 | 0.244067 | 1.672072 | 0.09451 | 0.26117 |
| Mfumv2_2331 | 345.4187836 | -0.806087222 | 0.14481 | -5.56652 | 2.60E-08 | 7.68E-07 |
| Mfumv2_2332 | 171.6100066 | -0.006664815 | 0.192704 | -0.03459 | 0.97241 | 0.990093 |
| Mfumv2_2333 | 154.052157 | 0.309357606 | 0.204778 | 1.510701 | 0.130865 | 0.318384 |
| Mfumv2_2334 | 438.9375373 | 0.030617625 | 0.128151 | 0.238918 | 0.811169 | 0.907508 |
| Mfumv2_2335 | 473.9351567 | -0.289090683 | 0.12917 | -2.23806 | 0.025217 | 0.101121 |
| Mfumv2_2336 | 953.6804517 | -0.106864621 | 0.109957 | -0.97188 | 0.331112 | 0.573388 |

| | | | | | | |
|---|---|---|---|---|---|---|
| Mfumv2_2337 | 389.7450264 | -0.399270365 | 0.133167 | -2.99828 | 0.002715 | 0.018617 |
| Mfumv2_2339 | 531.1946324 | -0.504047338 | 0.125998 | -4.00045 | 6.32E-05 | 0.00087 |
| Mfumv2_2340 | 16.08168029 | 0.499224412 | 0.468237 | 1.066178 | 0.286343 | NA |
| Mfumv2_2341 | 836.2485117 | -0.340264023 | 0.139651 | -2.43653 | 0.014829 | 0.069122 |
| Mfumv2_2342 | 340.8464216 | 0.278386601 | 0.142712 | 1.950688 | 0.051094 | 0.167862 |
| Mfumv2_2343 | 697.8332508 | 0.280681402 | 0.127365 | 2.203758 | 0.027541 | 0.107517 |
| Mfumv2_2344 | 877.0828791 | -0.123799367 | 0.106754 | -1.15967 | 0.246185 | 0.484779 |
| Mfumv2_2345 | 2.93845384 | 0.838103581 | 1.070627 | 0.782815 | 0.433736 | NA |
| Mfumv2_2347 | 29.8164312 | 0.380564369 | 0.3446 | 1.104365 | 0.269435 | 0.508125 |
| Mfumv2_2348 | 1903.454052 | -0.555744763 | 0.132867 | -4.18272 | 2.88E-05 | 0.000459 |
| Mfumv2_2349 | 322.8919034 | -0.145967674 | 0.136117 | -1.07237 | 0.283554 | 0.523113 |
| Mfumv2_2350 | 398.9597285 | 0.056634536 | 0.123045 | 0.460274 | 0.64532 | 0.827343 |
| Mfumv2_2351 | 456.9010424 | -0.020021638 | 0.120786 | -0.16576 | 0.868345 | 0.939349 |
| Mfumv2_2352 | 154.9267664 | -0.09065963 | 0.172226 | -0.5264 | 0.598611 | 0.799607 |
| Mfumv2_2353 | 345.4395503 | -0.320468474 | 0.157997 | -2.02832 | 0.042527 | 0.146046 |
| Mfumv2_2354 | 123.1643112 | 0.157639662 | 0.178816 | 0.881575 | 0.378007 | 0.612926 |
| Mfumv2_2355 | 55.09780775 | 0.040202832 | 0.254254 | 0.158121 | 0.874361 | 0.94339 |
| Mfumv2_2356 | 336.079722 | -0.266042869 | 0.131721 | -2.01975 | 0.043409 | 0.148315 |
| Mfumv2_2357 | 236.314098 | 0.102558101 | 0.149002 | 0.688302 | 0.491262 | 0.719349 |
| Mfumv2_2358 | 59.73373913 | 0.295239134 | 0.263209 | 1.121691 | 0.261994 | 0.499854 |
| Mfumv2_2359 | 226.8291246 | 0.341962544 | 0.146703 | 2.330987 | 0.019754 | 0.085709 |
| Mfumv2_2360 | 20.31899096 | 0.865379642 | 0.412392 | 2.09844 | 0.035866 | NA |
| Mfumv2_2361 | 236.3848958 | 0.085891724 | 0.155731 | 0.551539 | 0.581264 | 0.788739 |
| Mfumv2_2362 | 16.41729073 | 0.148327656 | 0.465306 | 0.318774 | 0.749898 | NA |
| Mfumv2_2363 | 42.1570535 | 0.142251381 | 0.28864 | 0.492833 | 0.622131 | 0.816369 |
| Mfumv2_2364 | 224.5365335 | 0.171089551 | 0.148085 | 1.15535 | 0.247947 | 0.486927 |
| Mfumv2_2365 | 216.2002324 | -0.140439954 | 0.179615 | -0.7819 | 0.434276 | 0.66651 |
| Mfumv2_2366 | 285.6581793 | -0.210106165 | 0.135614 | -1.5493 | 0.121309 | 0.304876 |
| Mfumv2_2367 | 1022.86963 | -0.044452471 | 0.102512 | -0.43363 | 0.664557 | 0.835454 |
| Mfumv2_2368 | 501.6356883 | -0.185863371 | 0.163505 | -1.13674 | 0.255646 | 0.493892 |
| Mfumv2_2369 | 1011.854885 | -0.144088728 | 0.127614 | -1.1291 | 0.258856 | 0.497912 |
| Mfumv2_2370 | 188.0748835 | 0.407863623 | 0.183429 | 2.223554 | 0.026178 | 0.103325 |
| Mfumv2_2371 | 2176.627543 | 0.024831841 | 0.089296 | 0.278084 | 0.780948 | 0.894484 |
| Mfumv2_2372 | 874.0446146 | 0.048919084 | 0.106723 | 0.458373 | 0.646684 | 0.827509 |
| Mfumv2_2373 | 774.2682278 | 0.257067587 | 0.134015 | 1.9182 | 0.055086 | 0.176222 |
| Mfumv2_2374 | 1326.269623 | -0.306384395 | 0.152946 | -2.00322 | 0.045153 | 0.151948 |
| Mfumv2_2375 | 2.8921662 | -0.030517637 | 1.0111 | -0.03018 | 0.975921 | NA |
| Mfumv2_2376 | 100.8843827 | -0.128583408 | 0.191537 | -0.67132 | 0.502014 | 0.729773 |
| Mfumv2_2377 | 139.7638978 | 0.062695076 | 0.195726 | 0.320321 | 0.748725 | 0.880198 |
| Mfumv2_2378 | 1312.216115 | -0.184505028 | 0.120183 | -1.5352 | 0.124734 | 0.309753 |
| Mfumv2_2379 | 320.1233719 | 0.250342241 | 0.136331 | 1.836279 | 0.066316 | 0.203094 |
| Mfumv2_2380 | 2386.584673 | -0.177332814 | 0.115305 | -1.53795 | 0.124061 | 0.308846 |
| Mfumv2_2381 | 3668.827676 | -0.582293086 | 0.11617 | -5.01242 | 5.38E-07 | 1.19E-05 |
| Mfumv2_2382 | 560.8957512 | -0.29618241 | 0.115624 | -2.5616 | 0.010419 | 0.052594 |
| Mfumv2_2383 | 243.9231849 | 0.124621087 | 0.143041 | 0.871226 | 0.383631 | 0.617559 |
| Mfumv2_2384 | 22.91674644 | 0.507419945 | 0.388301 | 1.306771 | 0.191291 | NA |
| Mfumv2_2385 | 18.55337246 | -0.128680093 | 0.432995 | -0.29719 | 0.766325 | NA |
| Mfumv2_2386 | 477.0392048 | -0.183305922 | 0.118615 | -1.54538 | 0.122253 | 0.306243 |
| Mfumv2_2387 | 93.10676366 | 0.218084863 | 0.200826 | 1.085942 | 0.277505 | 0.517918 |
| Mfumv2_2388 | 229.7628287 | 0.231113391 | 0.16527 | 1.398402 | 0.161992 | 0.368983 |
| Mfumv2_2389 | 238.9330639 | 0.140021317 | 0.152041 | 0.920942 | 0.357081 | 0.59836 |
| Mfumv2_2390 | 48.53004575 | -0.121713955 | 0.293976 | -0.41403 | 0.678855 | 0.842824 |

| | | | | | | |
|---|---|---|---|---|---|---|
| Mfumv2_2391 | 57.3167447 | -0.790714294 | 0.321081 | -2.46267 | 0.013791 | 0.065498 |
| Mfumv2_2392 | 87.3776283 | 0.515996539 | 0.224069 | 2.302849 | 0.021287 | 0.090799 |
| Mfumv2_2393 | 737.8544728 | 0.18679882 | 0.118352 | 1.578327 | 0.114491 | 0.294886 |
| Mfumv2_2394 | 986.0715496 | -0.181420947 | 0.119807 | -1.51428 | 0.129955 | 0.317732 |
| Mfumv2_2395 | 274.4882074 | 0.067932272 | 0.145017 | 0.468444 | 0.639467 | 0.823866 |
| Mfumv2_2396 | 3.793657816 | 0.555413237 | 0.91709 | 0.605626 | 0.544763 | NA |
| Mfumv2_2397 | 7245.487057 | -0.080442536 | 0.100639 | -0.79932 | 0.424104 | 0.656853 |
| Mfumv2_2398 | 1199.205271 | -0.074569077 | 0.128628 | -0.57973 | 0.562099 | 0.772934 |
| Mfumv2_2399 | 122.2868566 | 0.125884767 | 0.191827 | 0.65624 | 0.51167 | 0.735445 |
| Mfumv2_23s_rRN/ | 2738840.272 | -0.371487993 | 0.131696 | -2.82081 | 0.00479 | 0.028987 |
| Mfumv2_2400 | 211.9950202 | 0.348235195 | 0.15347 | 2.269078 | 0.023264 | 0.096763 |
| Mfumv2_2401 | 361.4902016 | 0.072252657 | 0.126124 | 0.572868 | 0.566734 | 0.777672 |
| Mfumv2_2402 | 42.4780906 | 0.548828244 | 0.290732 | 1.887749 | 0.05906 | 0.185682 |
| Mfumv2_2403 | 158.0603404 | 0.454094128 | 0.173401 | 2.61875 | 0.008825 | 0.046658 |
| Mfumv2_2404 | 91.11141656 | 0.072538679 | 0.203902 | 0.355752 | 0.722026 | 0.867012 |
| Mfumv2_2405 | 207.1522937 | 0.488273521 | 0.156063 | 3.128701 | 0.001756 | 0.012929 |
| Mfumv2_2406 | 10.6045171 | 0.061226092 | 0.536066 | 0.114214 | 0.909068 | NA |
| Mfumv2_2407 | 30.16099598 | 0.416542057 | 0.349796 | 1.190815 | 0.233726 | 0.470497 |
| Mfumv2_2408 | 115.1033886 | 0.168630558 | 0.197562 | 0.853559 | 0.393349 | 0.625684 |
| Mfumv2_2409 | 11.5807591 | 0.343626935 | 0.515584 | 0.666481 | 0.505104 | NA |
| Mfumv2_2410 | 17.48617981 | 0.099409602 | 0.430527 | 0.230902 | 0.817391 | NA |
| Mfumv2_2411 | 30.48975038 | 0.108474524 | 0.33106 | 0.327658 | 0.74317 | 0.87722 |
| Mfumv2_2412 | 34.9421411 | 0.237642556 | 0.315803 | 0.752504 | 0.451748 | 0.680331 |
| Mfumv2_2413 | 173.8610122 | 0.184137809 | 0.160936 | 1.144169 | 0.252553 | 0.492248 |
| Mfumv2_2414 | 363.3956449 | 0.07135921 | 0.128576 | 0.554995 | 0.578898 | 0.788261 |
| Mfumv2_2415 | 414.1498063 | -0.160806673 | 0.143576 | -1.12001 | 0.262708 | 0.500739 |
| Mfumv2_2416 | 916.170303 | -0.368519469 | 0.127744 | -2.88482 | 0.003916 | 0.024742 |
| Mfumv2_2417 | 256.8950183 | -0.062839563 | 0.178967 | -0.35112 | 0.725495 | 0.868606 |
| Mfumv2_2418 | 52.48524631 | -0.297856735 | 0.273344 | -1.08968 | 0.275856 | 0.516133 |
| Mfumv2_2419 | 121.0257821 | -0.015790388 | 0.197437 | -0.07998 | 0.936256 | 0.976096 |
| Mfumv2_2420 | 88.08440941 | -0.270908629 | 0.21673 | -1.24998 | 0.211306 | 0.442663 |
| Mfumv2_2421 | 159.4543612 | -0.038323532 | 0.16382 | -0.23394 | 0.815034 | 0.908704 |
| Mfumv2_2422 | 544.2103512 | 0.209045248 | 0.115005 | 1.81771 | 0.069108 | 0.20941 |
| Mfumv2_2423 | 19.0151404 | 0.572554212 | 0.418943 | 1.366663 | 0.171731 | NA |
| Mfumv2_2424 | 13.62136702 | 0.398252052 | 0.489368 | 0.813809 | 0.415754 | NA |
| Mfumv2_2425 | 448.206162 | 0.219422041 | 0.127198 | 1.725042 | 0.08452 | 0.240511 |
| Mfumv2_2426 | 39.33281269 | -0.579218544 | 0.302694 | -1.91355 | 0.055678 | 0.177416 |
| Mfumv2_2427 | 46.77565407 | 0.07761995 | 0.265613 | 0.29223 | 0.770111 | 0.887638 |
| Mfumv2_2428 | 26.01223224 | 0.402094572 | 0.372825 | 1.078508 | 0.280807 | 0.521056 |
| Mfumv2_2429 | 389.667275 | 0.057952886 | 0.151998 | 0.381273 | 0.703001 | 0.853886 |
| Mfumv2_2430 | 548.8244419 | 0.351005002 | 0.126677 | 2.770867 | 0.005591 | 0.032368 |
| Mfumv2_2431 | 3.778797792 | 0.730318819 | 0.974419 | 0.749491 | 0.453561 | NA |
| Mfumv2_2432 | 4.544791321 | 0.027267695 | 0.836619 | 0.032593 | 0.973999 | NA |
| Mfumv2_2433 | 293.7550492 | -0.120646092 | 0.130292 | -0.92596 | 0.354464 | 0.59836 |
| Mfumv2_2434 | 552.3762386 | -0.15298277 | 0.112552 | -1.35922 | 0.174075 | 0.389007 |
| Mfumv2_2435 | 170.8019338 | 0.260409641 | 0.159097 | 1.636801 | 0.101672 | 0.276774 |
| Mfumv2_2436 | 34.70793452 | 0.274218105 | 0.317721 | 0.863079 | 0.388094 | 0.621494 |
| Mfumv2_2437 | 35.74286085 | 1.02890016 | 0.314364 | 3.272959 | 0.001064 | 0.008692 |
| Mfumv2_2438 | 229.2583288 | 0.247538485 | 0.147503 | 1.678196 | 0.093309 | 0.258242 |
| Mfumv2_2439 | 1212.073861 | 0.201739747 | 0.144659 | 1.394586 | 0.163141 | 0.370758 |
| Mfumv2_2440 | 563.2630275 | -0.240178652 | 0.150405 | -1.59688 | 0.110293 | 0.290405 |
| Mfumv2_2441 | 506.3501279 | -0.211515475 | 0.132712 | -1.59379 | 0.110983 | 0.291291 |

| | | | | | | |
|---|---|---|---|---|---|---|
| Mfumv2_2442 | 676.2643066 | -0.068594411 | 0.1206 | -0.56878 | 0.569509 | 0.779561 |
| Mfumv2_2443 | 700.7868886 | -0.204680132 | 0.161173 | -1.26994 | 0.204105 | 0.433912 |
| Mfumv2_2444 | 79.44173555 | 0.118733751 | 0.231547 | 0.512785 | 0.608102 | 0.804793 |
| Mfumv2_2445 | 3.541159865 | 1.32204832 | 1.003408 | 1.317559 | 0.187651 | NA |
| Mfumv2_2446 | 242.3740274 | 0.165629286 | 0.150034 | 1.103944 | 0.269618 | 0.508125 |
| Mfumv2_2448 | 127.4258861 | -0.209587211 | 0.193313 | -1.08419 | 0.278281 | 0.518615 |
| Mfumv2_2449 | 129.3976455 | 0.212431897 | 0.177798 | 1.194797 | 0.232167 | 0.468766 |
| Mfumv2_2450 | 214.6696745 | 0.022918169 | 0.165918 | 0.13813 | 0.890138 | 0.94819 |
| Mfumv2_2451 | 267.8500595 | 0.105079292 | 0.142367 | 0.738086 | 0.460462 | 0.688808 |
| Mfumv2_2452 | 874.9751486 | 0.024219175 | 0.10804 | 0.224169 | 0.822626 | 0.913574 |
| Mfumv2_2453 | 1589.34821 | -0.225645198 | 0.102404 | -2.20347 | 0.027562 | 0.107517 |
| Mfumv2_2454 | 3255.234077 | -0.219522084 | 0.111046 | -1.97686 | 0.048058 | 0.160114 |
| Mfumv2_2455 | 628.8344067 | -0.324835119 | 0.146655 | -2.21496 | 0.026763 | 0.105012 |
| Mfumv2_2456 | 2428.42839 | -0.218193856 | 0.096948 | -2.25063 | 0.024409 | 0.099066 |
| Mfumv2_2457 | 1049.997606 | -0.334464842 | 0.138051 | -2.42276 | 0.015403 | 0.070812 |
| Mfumv2_2458 | 3458.884147 | -0.732144625 | 0.109508 | -6.68576 | 2.30E-11 | 1.10E-09 |
| Mfumv2_2459 | 816.0201713 | -0.699494237 | 0.152744 | -4.57953 | 4.66E-06 | 9.09E-05 |
| Mfumv2_2460 | 2222.392344 | -0.642773304 | 0.093928 | -6.84325 | 7.74E-12 | 3.89E-10 |
| Mfumv2_2461 | 594.8932989 | -0.376457483 | 0.126534 | -2.97516 | 0.002928 | 0.019545 |
| Mfumv2_2462 | 112.1029376 | -0.147516005 | 0.182765 | -0.80714 | 0.419588 | 0.652439 |
| Mfumv2_2463 | 112.4070897 | 0.357786656 | 0.209501 | 1.707806 | 0.087672 | 0.247149 |
| Mfumv2_2464 | 83.81445279 | -0.035162884 | 0.210512 | -0.16703 | 0.867343 | 0.938842 |
| Mfumv2_2465 | 204.1956734 | -0.211816307 | 0.148567 | -1.42573 | 0.153946 | 0.353863 |
| Mfumv2_2466 | 842.5870224 | -0.331120191 | 0.124913 | -2.6508 | 0.00803 | 0.043251 |
| Mfumv2_2467 | 29.73353513 | 0.608860162 | 0.371689 | 1.638089 | 0.101403 | 0.276417 |
| Mfumv2_2468 | 795.317108 | 0.742622093 | 0.139943 | 5.306621 | 1.12E-07 | 2.95E-06 |
| Mfumv2_2469 | 140.769606 | 0.593455784 | 0.179218 | 3.311366 | 0.000928 | 0.007837 |
| Mfumv2_2470 | 175.8427913 | 0.142806788 | 0.157222 | 0.908312 | 0.363713 | 0.603278 |
| Mfumv2_2471 | 665.7158197 | 0.100567881 | 0.137778 | 0.729929 | 0.465433 | 0.694177 |
| Mfumv2_2472 | 1534.62195 | -0.427100684 | 0.130441 | -3.27429 | 0.001059 | 0.008686 |
| Mfumv2_2474 | 0.340491013 | -0.677928304 | 2.998655 | -0.22608 | 0.821141 | NA |
| Mfumv2_2475 | 2.375964708 | 0.483005403 | 1.133676 | 0.426052 | 0.67007 | NA |
| Mfumv2_2476 | 110.0550854 | 0.182096112 | 0.19129 | 0.951937 | 0.341129 | 0.584519 |
| Mfumv2_2477 | 4.277600451 | 0.443264974 | 0.849813 | 0.521603 | 0.601947 | NA |
| Mfumv2_2478 | 0.923666441 | 1.568223259 | 1.908821 | 0.821566 | 0.411324 | NA |
| Mfumv2_2479 | 308.836085 | -0.152372455 | 0.13675 | -1.11424 | 0.265175 | 0.504008 |
| Mfumv2_2480 | 121.5712786 | 0.003593009 | 0.192229 | 0.018691 | 0.985087 | 0.992996 |
| Mfumv2_2481 | 92.47616315 | 0.175185618 | 0.217046 | 0.807138 | 0.419587 | 0.652439 |
| Mfumv2_2482 | 174.7281936 | 0.006275937 | 0.166048 | 0.037796 | 0.96985 | 0.98909 |
| Mfumv2_2483 | 257.1830722 | 0.036799092 | 0.146531 | 0.251134 | 0.80171 | 0.903903 |
| Mfumv2_2484 | 324.7647781 | 0.134613485 | 0.128755 | 1.045497 | 0.295793 | 0.534877 |
| Mfumv2_2485 | 556.1621834 | 0.458970132 | 0.121545 | 3.77612 | 0.000159 | 0.00185 |
| Mfumv2_2486 | 152.9921287 | 0.43558122 | 0.176308 | 2.470572 | 0.01349 | 0.06422 |
| Mfumv2_2487 | 77.85215618 | -0.110039292 | 0.212308 | -0.5183 | 0.60425 | 0.802869 |
| Mfumv2_2488 | 82.98152088 | 0.023679758 | 0.215575 | 0.109844 | 0.912533 | 0.963869 |
| Mfumv2_2489 | 59.33653836 | 0.276124167 | 0.242289 | 1.139647 | 0.254433 | 0.493394 |
| Mfumv2_2490 | 57.7738422 | 0.399582324 | 0.252938 | 1.579765 | 0.114161 | 0.294651 |
| Mfumv2_2491 | 127.7738052 | 0.072441641 | 0.180737 | 0.400812 | 0.688558 | 0.846498 |
| Mfumv2_2492 | 129.8497538 | 0.063488024 | 0.18003 | 0.352653 | 0.724349 | 0.86775 |
| Mfumv2_2493 | 494.9679286 | 0.132325575 | 0.151093 | 0.875792 | 0.381143 | 0.615528 |
| Mfumv2_2494 | 76.90167868 | 0.243197721 | 0.216065 | 1.125574 | 0.260346 | 0.499079 |
| Mfumv2_2495 | 130.4962412 | 0.119411903 | 0.178401 | 0.669345 | 0.503275 | 0.730021 |

| | | | | | | | |
|---|---|---|---|---|---|---|---|
| Mfumv2_2496 | 263.6173698 | | 0.004700224 | 0.151572 | 0.03101 | 0.975262 | 0.990165 |
| Mfumv2_2497 | 811.609866 | | -0.013209425 | 0.108949 | -0.12124 | 0.903498 | 0.957896 |
| Mfumv2_2498 | 911.1790332 | | -0.236106578 | 0.144552 | -1.63336 | 0.102392 | 0.277826 |
| Mfumv2_2499 | 1121.068271 | | -0.101527348 | 0.101574 | -0.99954 | 0.317532 | 0.560564 |
| Mfumv2_2500 | 535.2666995 | | -0.197789532 | 0.127245 | -1.55439 | 0.120091 | 0.304013 |
| Mfumv2_2501 | 1028.471875 | | -0.486751524 | 0.143236 | -3.39825 | 0.000678 | 0.006158 |
| Mfumv2_2502 | 482.0256037 | | -0.142154786 | 0.127939 | -1.11111 | 0.26652 | 0.505132 |
| Mfumv2_2503 | 1449.095924 | | -0.438931757 | 0.10194 | -4.30579 | 1.66E-05 | 0.000283 |
| Mfumv2_2504 | 316.8480791 | | 0.078461306 | 0.136045 | 0.576731 | 0.564121 | 0.775185 |
| Mfumv2_2505 | 292.6594618 | | -0.171312463 | 0.15255 | -1.12299 | 0.261442 | 0.499274 |
| Mfumv2_2506 | 164.1705581 | | 0.361823485 | 0.16826 | 2.150383 | 0.031525 | 0.117342 |
| Mfumv2_2507 | 126.5017112 | | 0.108755628 | 0.182573 | 0.595684 | 0.551386 | 0.766105 |
| Mfumv2_2508 | 175.1301488 | | -0.074958027 | 0.176146 | -0.42554 | 0.67044 | 0.838154 |
| Mfumv2_2509 | 409.3147044 | | 0.094404222 | 0.120146 | 0.785748 | 0.432015 | 0.665071 |
| Mfumv2_5s_rRNA_ | 5619.644765 | | -0.67185844 | 0.190034 | -3.53547 | 0.000407 | 0.00413 |
| Mfumv2_miscRNA | 168.8289109 | | 0.055870216 | 0.162117 | 0.344629 | 0.730373 | 0.871376 |
| Mfumv2_miscRNA | 0.169414025 | | -1.639708171 | 4.080473 | -0.40184 | 0.6878 | NA |
| Mfumv2_miscRNA | 61.29780632 | | -0.105908451 | 0.245162 | -0.43199 | 0.665746 | 0.835454 |
| Mfumv2_miscRNA | 0 | NA | | NA | NA | NA | NA |
| Mfumv2_miscRNA | 16968.39152 | | -0.095286375 | 0.139847 | -0.68136 | 0.495644 | 0.72344 |
| Mfumv2_miscRNA | 0 | NA | | NA | NA | NA | NA |
| Mfumv2_miscRNA | 4.029910259 | | -0.039025557 | 0.948477 | -0.04115 | 0.96718 | NA |
| Mfumv2_miscRNA | 15.1300946 | | -0.139907876 | 0.45832 | -0.30526 | 0.760166 | NA |
| Mfumv2_miscRNA | 76.12920332 | | -0.105576008 | 0.223572 | -0.47222 | 0.636767 | 0.823208 |
| Mfumv2_miscRNA | 0 | NA | | NA | NA | NA | NA |
| Mfumv2_miscRNA | 17.30781161 | | 0.048323976 | 0.430642 | 0.112214 | 0.910654 | NA |
| Mfumv2_miscRNA | 2.070762554 | | 2.03886788 | 1.403924 | 1.452264 | 0.146428 | NA |
| Mfumv2_miscRNA | 11.09593961 | | 0.537537715 | 0.57282 | 0.938405 | 0.348036 | NA |
| Mfumv2_miscRNA | 1.393625908 | | -0.002499561 | 1.530144 | -0.00163 | 0.998697 | NA |
| Mfumv2_miscRNA | 169.6002998 | | -0.069366362 | 0.167495 | -0.41414 | 0.678772 | 0.842824 |
| Mfumv2_miscRNA | 64.36677086 | | 0.043991341 | 0.234299 | 0.187757 | 0.851067 | 0.93126 |
| Mfumv2_miscRNA | 99.6863853 | | 0.122458022 | 0.220842 | 0.554506 | 0.579233 | 0.788261 |
| Mfumv2_miscRNA | 0 | NA | | NA | NA | NA | NA |
| Mfumv2_tmRNA1 | 38.36918339 | | -0.390493461 | 0.326284 | -1.19679 | 0.231389 | 0.468609 |
| Mfumv2_tRNA1 | 84.1076689 | | 0.24568223 | 0.22221 | 1.105632 | 0.268886 | 0.507699 |
| Mfumv2_tRNA10 | 483.0326722 | | -0.12475434 | 0.116024 | -1.07525 | 0.282263 | 0.522845 |
| Mfumv2_tRNA11 | 59.44431851 | | -0.384479428 | 0.307915 | -1.24865 | 0.211791 | 0.443218 |
| Mfumv2_tRNA12 | 0 | NA | | NA | NA | NA | NA |
| Mfumv2_tRNA13 | 807.8814242 | | -0.381688361 | 0.120154 | -3.17667 | 0.00149 | 0.011424 |
| Mfumv2_tRNA14 | 0.58806155 | | 0.863038643 | 2.301258 | 0.375029 | 0.707639 | NA |
| Mfumv2_tRNA15 | 735.1128553 | | 0.072639621 | 0.14791 | 0.491107 | 0.623351 | 0.817436 |
| Mfumv2_tRNA16 | 5.596818959 | | -0.049880676 | 0.733196 | -0.06803 | 0.94576 | NA |
| Mfumv2_tRNA17 | 2.974023381 | | 0.488060159 | 1.021767 | 0.477663 | 0.63289 | NA |
| Mfumv2_tRNA18 | 4.732605463 | | -0.010547089 | 0.830659 | -0.0127 | 0.989869 | NA |
| Mfumv2_tRNA19 | 42.97525933 | | 0.315271608 | 0.310934 | 1.01395 | 0.310606 | 0.55222 |
| Mfumv2_tRNA2 | 2.873414189 | | 0.267925685 | 1.044735 | 0.256453 | 0.797601 | NA |
| Mfumv2_tRNA20 | 6.689063036 | | 0.533238375 | 0.679738 | 0.784476 | 0.432761 | NA |
| Mfumv2_tRNA21 | 3.54136688 | | 0.646165528 | 0.964963 | 0.669627 | 0.503095 | NA |
| Mfumv2_tRNA22 | 0 | NA | | NA | NA | NA | NA |
| Mfumv2_tRNA23 | 171.210725 | | 0.394935007 | 0.174696 | 2.260696 | 0.023778 | 0.09769 |
| Mfumv2_tRNA24 | 63.71380964 | | 0.35003293 | 0.239852 | 1.45937 | 0.144463 | 0.337099 |
| Mfumv2_tRNA25 | 0.308392731 | | -0.677928239 | 3.098586 | -0.21879 | 0.826816 | NA |

| | | | | | | |
|---|---|---|---|---|---|---|
| Mfumv2_tRNA26 | 3.557389963 | -0.663836076 | 0.947849 | -0.70036 | 0.483702 | NA |
| Mfumv2_tRNA27 | 7.846337857 | 1.208280409 | 0.684154 | 1.766095 | 0.07738 | NA |
| Mfumv2_tRNA28 | 62.45472004 | 0.5391748 | 0.239561 | 2.250676 | 0.024406 | 0.099066 |
| Mfumv2_tRNA29 | 7.949290413 | 0.451569191 | 0.667445 | 0.676563 | 0.498683 | NA |
| Mfumv2_tRNA3 | 16.30639945 | 0.488635511 | 0.460179 | 1.061837 | 0.28831 | NA |
| Mfumv2_tRNA30 | 27.34524283 | -0.103936619 | 0.35783 | -0.29046 | 0.771461 | 0.888026 |
| Mfumv2_tRNA31 | 55.40763725 | 0.646902055 | 0.264188 | 2.448641 | 0.01434 | 0.067309 |
| Mfumv2_tRNA32 | 22.6230385 | 0.213625129 | 0.388376 | 0.550047 | 0.582287 | NA |
| Mfumv2_tRNA33 | 0.960372992 | 0.179301777 | 1.8578 | 0.096513 | 0.923113 | NA |
| Mfumv2_tRNA34 | 91.21432551 | 0.146833902 | 0.219208 | 0.669839 | 0.50296 | 0.730021 |
| Mfumv2_tRNA35 | 0.716002392 | -3.250576656 | 2.306894 | -1.40907 | 0.158814 | NA |
| Mfumv2_tRNA36 | 57.32143576 | 0.397866018 | 0.256263 | 1.552572 | 0.120526 | 0.304191 |
| Mfumv2_tRNA37 | 227.1389517 | 0.036632938 | 0.166283 | 0.220304 | 0.825634 | 0.914892 |
| Mfumv2_tRNA38 | 12.66836281 | 0.185674644 | 0.502952 | 0.36917 | 0.712001 | NA |
| Mfumv2_tRNA39 | 62.48531398 | -0.003276845 | 0.235767 | -0.0139 | 0.988911 | 0.994355 |
| Mfumv2_tRNA4 | 38.02293357 | 0.370557448 | 0.297393 | 1.246021 | 0.212757 | 0.443851 |
| Mfumv2_tRNA40 | 5.57875167 | -0.303250141 | 0.737594 | -0.41113 | 0.680974 | NA |
| Mfumv2_tRNA41 | 46.22525283 | 0.501262559 | 0.293551 | 1.707583 | 0.087714 | 0.247149 |
| Mfumv2_tRNA42 | 6.672254701 | -0.815019292 | 0.691756 | -1.17819 | 0.238721 | NA |
| Mfumv2_tRNA43 | 1.406836828 | 1.134744938 | 1.678902 | 0.675885 | 0.499114 | NA |
| Mfumv2_tRNA44 | 22.84789607 | -0.340273563 | 0.390287 | -0.87186 | 0.383287 | NA |
| Mfumv2_tRNA45 | 6.507478667 | 0.105029304 | 0.690359 | 0.152137 | 0.879079 | NA |
| Mfumv2_tRNA46 | 2.424062853 | -1.438989431 | 1.23042 | -1.16951 | 0.242198 | NA |
| Mfumv2_tRNA47 | 2.430467226 | 0.447459873 | 1.116221 | 0.40087 | 0.688515 | NA |
| Mfumv2_tRNA5 | 6.54024128 | -0.696305396 | 0.706208 | -0.98598 | 0.324144 | NA |
| Mfumv2_tRNA6 | 18.84219077 | -0.001501359 | 0.502897 | -0.00299 | 0.997618 | NA |
| Mfumv2_tRNA7 | 20.12387857 | 0.239064648 | 0.396329 | 0.603197 | 0.546378 | NA |
| Mfumv2_tRNA8 | 3.882884774 | 0.597682406 | 0.938827 | 0.636627 | 0.524368 | NA |
| Mfumv2_tRNA9 | 52.68711549 | -0.404207217 | 0.299083 | -1.35149 | 0.176539 | 0.392907 |

| | methane reactor | | methane/sulfide reactor | | methane/sulfide versus methane only | | | | |
|---|---|---|---|---|---|---|---|---|---|
| locus tag | TPM average | TPM stdv | TPM average | TPM stdv | log2 fold change | fold change | adjusted p value | Annotation | |
| Mfumv2_0219 | 210.45 | 6.69 | 479.12 | 78.59 | 0.85 | 1.81 | 0.00 | NAD(FAD)-dependent dehydrogenase | hcaD (SQR) |
| Mfumv2_0220 | 332.85 | 13.68 | 710.12 | 123.78 | 0.76 | 1.70 | 0.00 | Putative sulfur carrier protein AF_0556 | |
| Mfumv2_0221 | 805.59 | 110.42 | 1696.96 | 308.55 | 0.73 | 1.66 | 0.00 | Peroxiredoxin family protein | |
| Mfumv2_0303 | 14.65 | 3.56 | 29.53 | 6.00 | 0.68 | 1.60 | 0.03 | conserved protein of unknown function | |
| Mfumv2_0334 | 182.55 | 49.37 | 354.59 | 27.23 | 0.61 | 1.53 | 0.01 | DNA repair photolyase | |
| Mfumv2_0506 | 29.82 | 6.65 | 69.02 | 10.72 | 0.91 | 1.88 | 0.03 | conserved protein of unknown function | |
| Mfumv2_0636 | 11.27 | 3.52 | 27.26 | 3.90 | 0.95 | 1.93 | 0.03 | Acetyltransferase, GNAT family | wecD |
| Mfumv2_0674 | 287.50 | 93.50 | 543.12 | 48.05 | 0.62 | 1.53 | 0.00 | ABC-type transport system involved in gliding motility, ATPase component | |
| Mfumv2_0748 | 7.24 | 1.78 | 16.54 | 3.90 | 0.84 | 1.79 | 0.03 | Na+/H+ antiporter | kefB |
| Mfumv2_0768 | 38.08 | 11.22 | 76.24 | 3.21 | 0.70 | 1.63 | 0.00 | Membrane-fusion protein | acrA |
| Mfumv2_0777 | 22.85 | 7.14 | 47.84 | 5.66 | 0.74 | 1.67 | 0.01 | Transcriptional regulator LuxR family | csgD |
| Mfumv2_0791 | 13.74 | 1.81 | 26.12 | 2.45 | 0.61 | 1.52 | 0.02 | Heavy metal RND efflux outer membrane protein,CzcC family | tolC |
| Mfumv2_0900 | 25.18 | 3.89 | 48.67 | 3.62 | 0.63 | 1.55 | 0.01 | Putative Methylase involved in ubiquinone/menaquinone biosynthesis | |
| Mfumv2_0902 | 297.05 | 41.57 | 830.41 | 101.84 | 1.15 | 2.22 | 0.00 | carbonic anhydrase | |
| Mfumv2_0911 | 32.05 | 9.12 | 61.08 | 4.40 | 0.61 | 1.53 | 0.03 | protein of unknown function | |
| Mfumv2_0942 | 39.97 | 14.31 | 96.39 | 16.83 | 0.95 | 1.93 | 0.00 | Cytochrome c family protein | |
| Mfumv2_0943 | 16.49 | 4.89 | 170.10 | 23.66 | 3.06 | 8.33 | 0.00 | Sulfite oxidase or related enzyme | |
| Mfumv2_1083 | 9.80 | 3.32 | 21.13 | 0.82 | 0.79 | 1.73 | 0.02 | Glycosyltransferase | rfaG |
| Mfumv2_1143 | 68.87 | 32.53 | 143.78 | 15.81 | 0.74 | 1.67 | 0.05 | protein of unknown function | |
| Mfumv2_1181 | 27.01 | 6.89 | 52.27 | 5.25 | 0.65 | 1.57 | 0.03 | HAD superfamily hydrolase | |
| Mfumv2_1229 | 19.96 | 4.93 | 38.41 | 5.08 | 0.61 | 1.53 | 0.01 | Oligopeptide ABC transporter, periplasmic oligopeptide-binding protein oppA (TC 3.A.1.5.1) | |
| Mfumv2_1238 | 105.68 | 32.96 | 215.30 | 29.14 | 0.71 | 1.64 | 0.00 | tRNA-specific 2-thiouridylase MnmA 2 | mnmA |
| Mfumv2_1252 | 28.80 | 2.71 | 61.66 | 4.43 | 0.79 | 1.73 | 0.00 | 1-acyl-sn-glycerol-3-phosphate acyltransferase | |
| Mfumv2_1255 | 74.92 | 11.62 | 144.06 | 8.52 | 0.63 | 1.55 | 0.00 | Fe(2+) transporter FeoB | feoB |
| Mfumv2_1257 | 167.40 | 24.29 | 943.52 | 78.39 | 2.16 | 4.47 | 0.00 | conserved protein of unknown function | |
| Mfumv2_1258 | 79.90 | 18.39 | 554.24 | 79.46 | 2.44 | 5.44 | 0.00 | conserved protein of unknown function | |
| Mfumv2_1259 | 93.42 | 19.27 | 617.52 | 91.92 | 2.39 | 5.23 | 0.00 | Cytochrome c oxidase (B(O/a)3-type) chain II | |
| Mfumv2_1260 | 81.40 | 11.44 | 458.04 | 45.14 | 2.16 | 4.46 | 0.00 | Cytochrome c oxidase subunit 1 | |
| Mfumv2_1261 | 46.56 | 24.18 | 184.64 | 44.57 | 1.66 | 3.16 | 0.00 | conserved protein of unknown function | |
| Mfumv2_1269 | 24.59 | 14.09 | 82.32 | 26.93 | 1.43 | 2.70 | 0.00 | conserved protein of unknown function | |
| Mfumv2_1270 | 69.85 | 13.99 | 155.00 | 14.01 | 0.84 | 1.79 | 0.00 | L-sorbosone dehydrogenase | |
| Mfumv2_1272 | 82.58 | 6.27 | 156.96 | 7.37 | 0.60 | 1.52 | 0.00 | Rhodanese-related sulfurtransferase | |
| Mfumv2_1278 | 140.86 | 48.09 | 281.40 | 35.55 | 0.68 | 1.61 | 0.01 | conserved protein of unknown function | |
| Mfumv2_1279 | 352.71 | 63.78 | 690.72 | 57.71 | 0.65 | 1.57 | 0.00 | Putative bacterial haemoglobin | |
| Mfumv2_1289 | 92.71 | 16.09 | 353.54 | 36.94 | 1.62 | 3.07 | 0.00 | Organic hydroperoxide reductase | osmC |
| Mfumv2_1291 | 31.51 | 2.83 | 61.06 | 7.57 | 0.63 | 1.54 | 0.01 | SAM-dependent methyltransferase | smtA |
| Mfumv2_1292 | 55.15 | 8.91 | 102.71 | 8.53 | 0.58 | 1.50 | 0.00 | Hemoglobin-like flavoprotein fused to Roadblock/LC7 domain | |
| Mfumv2_1293 | 9.12 | 1.67 | 18.47 | 1.18 | 0.70 | 1.63 | 0.00 | Outer membrane receptor protein, mostly Fe transport | cirA |
| Mfumv2_1297 | 1.90 | 0.13 | 6.17 | 0.27 | 1.39 | 2.62 | 0.00 | Assimilatory nitrate reductase catalytic subunit | nasC |
| Mfumv2_1311 | 27.04 | 6.48 | 60.43 | 7.56 | 0.84 | 1.79 | 0.03 | protein of unknown function | |
| Mfumv2_1316 | 42.98 | 7.92 | 83.86 | 4.57 | 0.64 | 1.56 | 0.00 | Uracil phosphoribosyltransferase | upp |
| Mfumv2_1331 | 61.12 | 5.91 | 124.47 | 17.78 | 0.70 | 1.63 | 0.00 | Transaldolase | tal |
| Mfumv2_1342 | 55.49 | 18.57 | 105.66 | 9.28 | 0.62 | 1.54 | 0.01 | Demethylmenaquinone methyltransferase | menG |
| Mfumv2_1348 | 15.37 | 0.55 | 39.52 | 3.06 | 1.04 | 2.05 | 0.00 | Signal transduction histidine kinase | |
| Mfumv2_1350 | 22.99 | 6.95 | 47.29 | 2.19 | 0.74 | 1.67 | 0.01 | conserved protein of unknown function | |
| Mfumv2_1378 | 22.43 | 3.72 | 45.48 | 1.10 | 0.70 | 1.62 | 0.00 | DNA-directed RNA polymerase, sigma subunit (Sigma70/sigma32) | rpoD |

| | | | | | | | | | |
|---|---|---|---|---|---|---|---|---|---|
| Mfumv2_1379 | 3.49 | 0.99 | 8.88 | 2.33 | 1.04 | 2.05 | 0.05 | conserved protein of unknown function | |
| Mfumv2_1415 | 21.46 | 6.60 | 47.78 | 4.42 | 0.82 | 1.76 | 0.00 | SAM-dependent methyltransferase (Modular protein) | |
| Mfumv2_1429 | 80.28 | 2.65 | 182.90 | 30.31 | 0.86 | 1.81 | 0.00 | Predicted metal-binding, possibly nucleic acid-binding protein | |
| Mfumv2_1502 | 26.93 | 7.07 | 53.42 | 6.87 | 0.68 | 1.60 | 0.00 | transposase | |
| Mfumv2_1531 | 97.89 | 21.65 | 261.35 | 27.46 | 1.10 | 2.14 | 0.00 | conserved protein of unknown function | |
| Mfumv2_1662 | 22.69 | 4.32 | 44.36 | 7.90 | 0.64 | 1.55 | 0.01 | Na+/H+ antiporter | kefB |
| Mfumv2_1717 | 24.15 | 7.04 | 46.15 | 6.53 | 0.61 | 1.53 | 0.02 | conserved protein of unknown function | |
| Mfumv2_1764 | 44.27 | 8.99 | 85.94 | 13.87 | 0.62 | 1.54 | 0.00 | conserved protein of unknown function | |
| Mfumv2_1780 | 53.23 | 11.00 | 99.06 | 8.65 | 0.58 | 1.50 | 0.00 | TPR_REGION domain-containing protein | |
| Mfumv2_1785 | 43.36 | 9.58 | 112.47 | 6.15 | 1.04 | 2.06 | 0.00 | 6-phosphogluconolactonase | pgl |
| Mfumv2_1786 | 76.36 | 7.61 | 150.75 | 10.50 | 0.66 | 1.58 | 0.00 | Glucose-6-phosphate 1-dehydrogenase | zwf |
| Mfumv2_1788 | 8013.54 | 396.01 | 18258.13 | 746.79 | 0.86 | 1.82 | 0.00 | Particulate methane monooxygenase PmoC subunit | pmoC |
| Mfumv2_1789 | 5.52 | 0.80 | 14.07 | 3.33 | 1.04 | 2.06 | 0.00 | protein of unknown function | |
| Mfumv2_1863 | 17.39 | 4.05 | 34.55 | 4.29 | 0.68 | 1.60 | 0.05 | conserved protein of unknown function | |
| Mfumv2_1950 | 19.32 | 1.61 | 379.69 | 22.56 | 3.98 | 15.73 | 0.00 | MULTIHEME_CYTC domain-containing protein | |
| Mfumv2_1951 | 6.32 | 0.54 | 76.46 | 4.34 | 3.28 | 9.70 | 0.00 | Putative Starvation-inducible outer membrane lipoprotein | |
| Mfumv2_1958 | 94.89 | 6.91 | 209.32 | 13.56 | 0.82 | 1.77 | 0.01 | protein of unknown function | |
| Mfumv2_1984 | 13.82 | 2.75 | 29.40 | 5.23 | 0.75 | 1.69 | 0.04 | conserved exported protein of unknown function | |
| Mfumv2_1990 | 27.15 | 3.63 | 120.73 | 19.93 | 1.83 | 3.56 | 0.00 | Outer membrane protein | tolC |
| Mfumv2_1991 | 23.61 | 1.76 | 72.79 | 7.33 | 1.29 | 2.45 | 0.00 | Cation/multidrug efflux pump | acrB |
| Mfumv2_1992 | 267.43 | 42.85 | 1712.40 | 96.87 | 2.35 | 5.09 | 0.00 | conserved protein of unknown function | |
| Mfumv2_2058 | 21.58 | 4.03 | 40.95 | 6.56 | 0.61 | 1.53 | 0.01 | Phosphate-binding protein PstS | pstS |
| Mfumv2_2175 | 21.18 | 5.27 | 40.23 | 3.74 | 0.60 | 1.52 | 0.04 | Ribosomal large subunit pseudouridine synthase D | |
| Mfumv2_2176 | 45.94 | 7.00 | 95.33 | 0.71 | 0.75 | 1.68 | 0.00 | Alcohol dehydrogenase | adhT |
| Mfumv2_2186 | 46.53 | 19.24 | 103.63 | 14.82 | 0.84 | 1.79 | 0.00 | Exopolyphosphatase | ppx |
| Mfumv2_2187 | 485.15 | 89.21 | 1474.24 | 122.35 | 1.28 | 2.43 | 0.00 | protein of unknown function | |
| Mfumv2_2189 | 64.78 | 11.83 | 130.49 | 19.97 | 0.69 | 1.61 | 0.00 | Ribosomal RNA small subunit methyltransferase A | rsmA |
| Mfumv2_2267 | 151.27 | 21.01 | 299.93 | 41.50 | 0.65 | 1.57 | 0.00 | conserved membrane protein of unknown function | |
| Mfumv2_2268 | 50.37 | 3.72 | 102.33 | 5.05 | 0.70 | 1.63 | 0.00 | NosD domain-containing protein | |
| Mfumv2_2437 | 9.25 | 1.26 | 23.65 | 0.61 | 1.03 | 2.04 | 0.01 | conserved protein of unknown function | |
| Mfumv2_2468 | 123.80 | 22.22 | 256.75 | 35.14 | 0.74 | 1.67 | 0.00 | 3,4-dihydroxy-2-butanone 4-phosphate synthase / GTP cyclohydrolase-2 | ribBA |
| Mfumv2_2469 | 38.16 | 8.48 | 71.56 | 7.23 | 0.59 | 1.51 | 0.01 | conserved protein of unknown function | |

| | methane reactor | | methane/sulfide reactor | | methane/sulfide versus methane only | | | | |
|---|---|---|---|---|---|---|---|---|---|
| locus tag | TPM average | TPM stdv | TPM average | TPM stdv | log2 fold change | fold change | adjusted p value | Annotation | |
| Mfumv2_0016 | 142.68 | 32.46 | 105.44 | 14.78 | -0.76 | 1.69 | 0.00 | LSU m5C1962 methyltransferase RlmI | |
| Mfumv2_0060 | 156.14 | 20.16 | 94.25 | 1.62 | -1.06 | 2.08 | 0.00 | HD Cas3-type domain-containing protein | |
| Mfumv2_0061 | 185.99 | 24.31 | 126.68 | 5.23 | -0.89 | 1.85 | 0.00 | conserved protein of unknown function | |
| Mfumv2_0062 | 123.93 | 11.51 | 84.26 | 5.95 | -0.89 | 1.85 | 0.00 | conserved protein of unknown function | |
| Mfumv2_0063 | 89.50 | 5.81 | 63.22 | 6.91 | -0.84 | 1.78 | 0.00 | conserved protein of unknown function | |
| Mfumv2_0064 | 85.89 | 7.02 | 60.77 | 2.05 | -0.82 | 1.77 | 0.00 | conserved protein of unknown function | |
| Mfumv2_0076 | 212.80 | 100.63 | 131.70 | 4.46 | -0.98 | 1.98 | 0.02 | subunit of E1(0) component of 2-oxoglutarate dehydrogenase | sucA |
| Mfumv2_0077 | 163.31 | 47.82 | 123.08 | 7.40 | -0.72 | 1.64 | 0.00 | dihydrolipoyltranssuccinylase | sucB |
| Mfumv2_0078 | 81.80 | 23.95 | 54.47 | 6.76 | -0.90 | 1.86 | 0.00 | Dihydrolipoyl dehydrogenase 3 | lpd |
| Mfumv2_0109 | 1201.10 | 240.10 | 946.11 | 31.99 | -0.68 | 1.60 | 0.00 | conserved protein of unknown function | |
| Mfumv2_0169 | 2219.04 | 190.97 | 1380.73 | 49.56 | -1.01 | 2.01 | 0.00 | conserved protein of unknown function | |
| Mfumv2_0170 | 29.65 | 4.24 | 23.95 | 5.45 | -0.64 | 1.56 | 0.00 | Outer membrane receptor protein, mostly Fe transport | cirA |
| Mfumv2_0200 | 188.58 | 52.52 | 108.43 | 10.35 | -1.11 | 2.16 | 0.00 | Putative type I restriction enzyme HindVIIP M protein | |
| Mfumv2_0202 | 99.25 | 26.36 | 51.01 | 3.43 | -1.27 | 2.42 | 0.00 | Type I restriction-modification system, specificity subunit S | |
| Mfumv2_0203 | 55.95 | 11.48 | 27.13 | 0.18 | -1.36 | 2.57 | 0.00 | Putative type I restriction enzyme HindVIIP R protein | |
| Mfumv2_0247 | 73.97 | 12.96 | 62.00 | 7.03 | -0.59 | 1.51 | 0.01 | protein of unknown function | |
| Mfumv2_0248 | 82.36 | 16.21 | 65.46 | 5.38 | -0.67 | 1.59 | 0.00 | protein of unknown function | |
| Mfumv2_0252 | 333.97 | 48.38 | 272.05 | 23.50 | -0.61 | 1.53 | 0.00 | Peroxiredoxin | bcp |
| Mfumv2_0253 | 136.40 | 12.82 | 113.19 | 7.25 | -0.59 | 1.50 | 0.00 | ATP-dependent 6-phosphofructokinase | pfkA |
| Mfumv2_0274 | 136.35 | 60.35 | 104.27 | 4.43 | -0.68 | 1.60 | 0.01 | conserved protein of unknown function | |
| Mfumv2_0344 | 444.46 | 32.78 | 369.50 | 17.29 | -0.59 | 1.51 | 0.00 | conserved protein of unknown function | |
| Mfumv2_0358 | 577.47 | 126.71 | 404.52 | 17.09 | -0.85 | 1.80 | 0.00 | Ribosome hibernation protein YhbH | |
| Mfumv2_0423 | 176.06 | 8.15 | 127.97 | 13.74 | -0.77 | 1.71 | 0.00 | conserved protein of unknown function | |
| Mfumv2_0424 | 2057.80 | 93.43 | 1357.91 | 151.78 | -0.93 | 1.90 | 0.00 | protein of unknown function | |
| Mfumv2_0425 | 1088.48 | 144.56 | 721.49 | 84.99 | -0.92 | 1.89 | 0.00 | conserved protein of unknown function | |
| Mfumv2_0426 | 880.10 | 85.68 | 601.73 | 68.25 | -0.87 | 1.83 | 0.00 | conserved protein of unknown function | |
| Mfumv2_0427 | 908.05 | 84.09 | 619.96 | 63.45 | -0.88 | 1.84 | 0.00 | conserved protein of unknown function | |
| Mfumv2_0428 | 864.23 | 75.05 | 700.65 | 147.48 | -0.64 | 1.56 | 0.00 | conserved protein of unknown function | |
| Mfumv2_0443 | 225.03 | 28.29 | 187.01 | 16.60 | -0.60 | 1.52 | 0.00 | conserved protein of unknown function | |
| Mfumv2_0525 | 441.60 | 281.79 | 46.24 | 4.99 | -3.54 | 11.63 | 0.00 | Sulfate adenylyltransferase subunit 1 | |
| Mfumv2_0526 | 909.86 | 369.38 | 59.87 | 6.49 | -4.22 | 18.65 | 0.00 | sulfate adenylyltransferase subunit 2 | cysD |
| Mfumv2_0527 | 376.13 | 128.38 | 15.46 | 3.11 | -4.88 | 29.39 | 0.00 | phosphoadenosine phosphosulfate reductase | cysH |
| Mfumv2_0528 | 29.77 | 19.86 | 4.13 | 1.14 | -3.10 | 8.59 | 0.00 | Homocitrate synthase 1 | nifV |
| Mfumv2_0564 | 1017.54 | 74.22 | 837.65 | 59.40 | -0.61 | 1.53 | 0.00 | Ysc84 domain-containing protein | |
| Mfumv2_0567 | 231.72 | 46.62 | 171.96 | 15.57 | -0.77 | 1.70 | 0.00 | conserved protein of unknown function | |
| Mfumv2_0573 | 28.21 | 11.33 | 16.48 | 2.59 | -1.12 | 2.17 | 0.00 | Polysulphide reductase | |
| Mfumv2_0578 | 453.78 | 69.88 | 336.73 | 51.40 | -0.77 | 1.71 | 0.00 | conserved protein of unknown function | |
| Mfumv2_0658 | 813.38 | 175.43 | 514.85 | 56.30 | -0.97 | 1.96 | 0.00 | Biopolymer transport protein | tolQ |
| Mfumv2_0659 | 383.67 | 88.54 | 254.68 | 63.57 | -0.91 | 1.88 | 0.00 | Biopolymer transport protein | exbD |
| Mfumv2_0660 | 137.01 | 49.61 | 93.73 | 15.95 | -0.87 | 1.82 | 0.00 | Periplasmic protein TonB | tonB |
| Mfumv2_0814 | 1402.28 | 59.99 | 762.65 | 39.49 | -1.21 | 2.31 | 0.00 | Transcriptional regulator, GntR family | phnF |
| Mfumv2_0815 | 1374.81 | 401.82 | 50.63 | 5.18 | -5.06 | 33.40 | 0.00 | Sulfite reductase [NADPH] hemoprotein beta-component | cysI |
| Mfumv2_0823 | 629.69 | 59.85 | 514.72 | 59.02 | -0.62 | 1.53 | 0.00 | conserved protein of unknown function | |
| Mfumv2_0894 | 852.43 | 147.15 | 687.57 | 13.61 | -0.64 | 1.56 | 0.00 | putative Glutamine synthetase and cystathionine beta-lyase binding protein | |
| Mfumv2_1068 | 332.52 | 4.65 | 271.66 | 12.53 | -0.62 | 1.53 | 0.00 | Epoxyqueuosine reductase QueH | queH |
| Mfumv2_1191 | 2120.36 | 135.87 | 1752.12 | 153.39 | -0.61 | 1.52 | 0.00 | 3-hydroxyisobutyrate dehydrogenase or related beta-hydroxyacid dehydrogenase | mmsB |

| | | | | | | | | | |
|---|---|---|---|---|---|---|---|---|---|
| Mfumv2_1245 | 68.78 | 28.46 | 55.47 | 2.26 | -0.61 | 1.52 | 0.03 | succinyl-CoA synthetase subunit beta | sucC |
| Mfumv2_1355 | 485.00 | 112.47 | 397.11 | 34.29 | -0.60 | 1.52 | 0.00 | Ribosomal large subunit pseudouridine synthase D | rluD |
| Mfumv2_1357 | 4871.29 | 58.77 | 3058.76 | 134.22 | -0.99 | 1.99 | 0.00 | Ferredoxin | |
| Mfumv2_1464 | 354.35 | 41.47 | 212.94 | 34.05 | -1.06 | 2.08 | 0.00 | PqqA peptide cyclase | pqqE |
| Mfumv2_1539 | 403.32 | 39.70 | 99.20 | 7.08 | -2.35 | 5.10 | 0.00 | conserved protein of unknown function | |
| Mfumv2_1540 | 76.44 | 7.95 | 46.07 | 4.28 | -1.05 | 2.07 | 0.00 | HTH cro/C1-type domain-containing protein | |
| Mfumv2_1601 | 55.85 | 12.01 | 23.47 | 2.89 | -1.59 | 3.00 | 0.00 | conserved membrane protein of unknown function | |
| Mfumv2_1602 | 31.31 | 4.40 | 14.76 | 1.73 | -1.41 | 2.66 | 0.00 | phosphoenolpyruvate synthetase | ppsA |
| Mfumv2_1603 | 22.19 | 2.21 | 8.51 | 2.00 | -1.73 | 3.31 | 0.00 | conserved protein of unknown function | |
| Mfumv2_1604 | 97.34 | 17.70 | 35.06 | 8.35 | -1.83 | 3.55 | 0.00 | Methane monooxygenase subunit alpha | pmoB3 |
| Mfumv2_1605 | 366.98 | 47.94 | 157.35 | 17.30 | -1.56 | 2.95 | 0.00 | Methane monooxygenase subunit beta | pmoA3 |
| Mfumv2_1606 | 1170.47 | 123.49 | 513.87 | 52.26 | -1.52 | 2.88 | 0.00 | Methane monooxygenase subunit gamma | pmoC3 |
| Mfumv2_1608 | 144.79 | 16.56 | 106.14 | 14.89 | -0.79 | 1.73 | 0.00 | Lactoylglutathione lyase family protein | gloA |
| Mfumv2_1609 | 107.35 | 12.60 | 68.52 | 0.66 | -0.96 | 1.95 | 0.00 | conserved protein of unknown function | |
| Mfumv2_1610 | 73.93 | 5.03 | 44.19 | 2.55 | -1.06 | 2.09 | 0.00 | Glucose-methanol-choline (GMC) oxidoreductase:NAD binding subunit | |
| Mfumv2_1616 | 103.62 | 11.14 | 85.88 | 16.42 | -0.62 | 1.53 | 0.01 | Ribonuclease HIII | |
| Mfumv2_1668 | 86.15 | 22.80 | 66.27 | 6.24 | -0.69 | 1.61 | 0.00 | Outer membrane receptor protein, mostly Fe transport | cirA |
| Mfumv2_1759 | 1129.07 | 152.78 | 936.44 | 68.60 | -0.61 | 1.52 | 0.00 | conserved protein of unknown function | |
| Mfumv2_1791 | 33277.44 | 4505.79 | 25524.24 | 641.52 | -0.71 | 1.64 | 0.00 | Methane monooxygenase subunit alpha | pmoB1 |
| Mfumv2_1792 | 36538.89 | 3862.35 | 27028.85 | 289.90 | -0.76 | 1.70 | 0.00 | Methane monooxygenase subunit beta | pmoA1 |
| Mfumv2_1793 | 93774.56 | 5646.44 | 72542.24 | 1997.94 | -0.70 | 1.62 | 0.00 | Particulate methane monooxygenase PmoC subunit | pmoC |
| Mfumv2_1799 | 625.74 | 73.82 | 521.97 | 41.20 | -0.60 | 1.51 | 0.00 | DNA-binding response regulator, NarL family (REC-HTH domains) | citB |
| Mfumv2_1945 | 173.79 | 23.16 | 139.49 | 9.99 | -0.64 | 1.56 | 0.00 | Transcriptional regulator containing HTH domain,ArsR | arsR |
| Mfumv2_1987 | 62.11 | 28.95 | 48.32 | 6.07 | -0.68 | 1.60 | 0.03 | putative membrane transporter protein | |
| Mfumv2_2036 | 537.88 | 103.05 | 379.40 | 73.98 | -0.85 | 1.80 | 0.00 | conserved protein of unknown function | |
| Mfumv2_2043 | 80.69 | 3.68 | 47.03 | 5.20 | -1.11 | 2.15 | 0.00 | Acyl-CoA synthetase (AMP-forming)/AMP-acid ligase II | caiC |
| Mfumv2_2064 | 92.04 | 7.72 | 73.01 | 4.66 | -0.66 | 1.58 | 0.00 | Succinate dehydrogenase flavoprotein subunit | |
| Mfumv2_2065 | 44.54 | 9.48 | 22.48 | 3.37 | -1.32 | 2.49 | 0.00 | Succinate dehydrogenase cytochrome b subunit | |
| Mfumv2_2087 | 28729.42 | 1923.36 | 23516.96 | 946.55 | -0.62 | 1.54 | 0.00 | Opacity protein or related surface antigen | |
| Mfumv2_2130 | 174.12 | 12.39 | 115.53 | 4.88 | -0.91 | 1.89 | 0.00 | HNHc domain-containing protein | |
| Mfumv2_2131 | 84.98 | 4.43 | 57.52 | 5.93 | -0.89 | 1.86 | 0.00 | Putative DNA modification methylase | |
| Mfumv2_2132 | 67.44 | 5.60 | 49.33 | 3.34 | -0.78 | 1.71 | 0.00 | conserved protein of unknown function | |
| Mfumv2_2133 | 88.58 | 3.97 | 66.74 | 2.21 | -0.73 | 1.66 | 0.00 | DUF87 domain-containing protein | |
| Mfumv2_2134 | 65.68 | 8.57 | 43.18 | 4.84 | -0.92 | 1.90 | 0.00 | SIR2_2 domain-containing protein | |
| Mfumv2_2135 | 59.72 | 5.92 | 40.95 | 0.45 | -0.86 | 1.82 | 0.00 | ATPase-like | |
| Mfumv2_2171 | 361.14 | 92.10 | 205.58 | 15.98 | -1.15 | 2.22 | 0.00 | conserved exported protein of unknown function | |
| Mfumv2_2172 | 30.33 | 5.97 | 15.14 | 2.44 | -1.31 | 2.49 | 0.00 | conserved protein of unknown function | |
| Mfumv2_2216 | 83.25 | 36.76 | 63.75 | 2.07 | -0.68 | 1.60 | 0.01 | Ligand-gated ion channel, periplasmic domain | hisJ |
| Mfumv2_2217 | 1473.00 | 405.74 | 563.65 | 31.82 | -1.68 | 3.21 | 0.00 | conserved protein of unknown function | |
| Mfumv2_2220 | 731.17 | 44.63 | 304.46 | 16.38 | -1.59 | 3.01 | 0.00 | O-acetylserine sulfhydrylase A | cysK |
| Mfumv2_2239 | 256.72 | 84.65 | 200.67 | 10.65 | -0.66 | 1.58 | 0.00 | NADH-quinone oxidoreductase subunit D | nuoD |
| Mfumv2_2240 | 265.47 | 96.28 | 199.54 | 13.57 | -0.71 | 1.64 | 0.00 | NADH-quinone oxidoreductase subunit C | nuoC |
| Mfumv2_2241 | 388.57 | 122.09 | 301.78 | 32.98 | -0.67 | 1.59 | 0.00 | NADH-quinone oxidoreductase subunit B | nuoB |
| Mfumv2_2249 | 878.95 | 86.14 | 674.94 | 38.91 | -0.70 | 1.63 | 0.00 | Nucleotide excision repair protein, with UvrB/UvrC motif | |
| Mfumv2_2250 | 108.82 | 18.21 | 87.34 | 9.45 | -0.64 | 1.56 | 0.00 | conserved protein of unknown function | |
| Mfumv2_2258 | 1003.75 | 90.09 | 785.94 | 70.84 | -0.68 | 1.61 | 0.00 | conserved exported protein of unknown function | |
| Mfumv2_2321 | 469.98 | 35.58 | 381.62 | 27.03 | -0.62 | 1.54 | 0.00 | conserved protein of unknown function | |
| Mfumv2_2331 | 162.85 | 34.84 | 116.01 | 14.68 | -0.81 | 1.75 | 0.00 | Periplasmic protein TonB | tonB |

| | | | | | | | | | |
|---|---|---|---|---|---|---|---|---|---|
| Mfumv2_2381 | 1464.03 | 93.66 | 1229.59 | 93.27 | -0.58 | 1.50 | 0.00 | Opacity protein or related surface antigen | |
| Mfumv2_2458 | 720.05 | 117.80 | 540.45 | 59.92 | -0.73 | 1.66 | 0.00 | ATP synthase F1 complex subunit alpha | atpA |
| Mfumv2_2459 | 298.24 | 85.83 | 228.31 | 28.74 | -0.70 | 1.62 | 0.00 | ATP synthase gamma chain | atpG |
| Mfumv2_2460 | 499.37 | 59.49 | 399.09 | 33.47 | -0.64 | 1.56 | 0.00 | ATP synthase F1 complex subunit beta | atpD |
